# Rarity and reproductive biology drive population genomics of obligate symbioses

**DOI:** 10.1101/2025.10.20.682950

**Authors:** Stephen T. Sharrett, Julianna Paulsen, Khoi Nguyen, James C. Lendemer, Paul Spruell, Meaghan Petix, Elaine M. Larsen, Jesse E.D. Miller, Jessica L. Allen

## Abstract

Modern perspectives on obligate symbioses increasingly recognize their dynamism, complexity, and global importance. Population genomics has been key to this advancement, yet the extension of existing frameworks to reproducibly integrate across multiple genotype-phenotype-environment relationships remains elusive. We generated 962 metagenomes for six lichens, obligate symbioses between fungi and algae, with contrasting traits sampled at a landscape scale. Core neutral population metrics were primarily driven by rarity and reproductive biology, while genes underlying expression regulation were under selection in response to land use and climate across all species. Examined within a comparative framework, these results illustrate that symbiont interactions are a major factor influencing microevolutionary trajectories, especially neutral processes. We present a conceptual framework for population-scale evolution in obligate symbioses that centers on biotic interactions.

## Introduction

Obligate symbioses are foundational to the structure and function of all ecosystems. These intimate interorganism relationships, ranging from parasitisms to mutualisms, promote biodiversity, stabilize food web connectivity, and regulate biogeochemical cycling (*1–4*). Metagenomic sequencing has broadened capabilities for studying complex assemblages of organisms, allowing the reconceptualization of symbioses to encompass multilayered biotic interactions. This expansion in perspective has been pivotal in revealing the ecological and evolutionary dynamism of symbiotic relationships (*5–8*).

Rarity is an emergent ecological phenomenon that describes the distributions or demographics of species (*9*). Patterns of rarity are shaped by the interplay between life histories and the environment (*9*, *10*). Rare species are crucial to ecosystem function (*11*), but efforts to prevent extinctions are limited by an incomplete understanding of the factors that underpin rarity. Obligate symbiotic systems present unique challenges in clarifying the interactions that produce patterns of rarity. For instance, one symbiosis-specific factor that may be a determinant of rarity is the degree of symbiont specificity (*12*, *13*). Further, obligately symbiotic organisms often have complex life cycles and reproductive strategies that include combinations of symbiotic and aposymbiotic, asexual and sexual reproductive strategies that are entwined with both rarity and symbiont specificity. Despite advances in our understanding of the inherent complexity of symbioses, we still lack population-level evolutionary frameworks that simultaneously integrate across multiple biotic and abiotic interactions that are fundamental to obligate symbioses.

Leveraging the power of comparative frameworks and the emerging field of landscape genomics, we connect microevolutionary processes to rarity, reproduction, symbiont dynamics, and the environment to advance our understanding of population genomics within the context of symbiotic relationships. We generated 962 metagenomes and symbiont-specific reference genomes from six different lichens, obligate symbioses between fungi (mycobionts) and algae (photobionts), in a comparative framework of three pairs of congeners with contrasting range sizes and reproductive biologies (Fig. 1; see Materials and Methods). Here we show that core population metrics, including gene flow, genetic diversity, and symbiont specificity, are primarily driven by rarity and reproductive biology. In contrast, adaptation to land use and climate in dominant symbionts is recovered in genes underlying expression regulation in both symbionts, regardless of rarity or reproductive mode. We use our findings to develop an extended eco-evolutionary framework for obligate symbioses founded on our novel empirical data.

**Figure 1.**
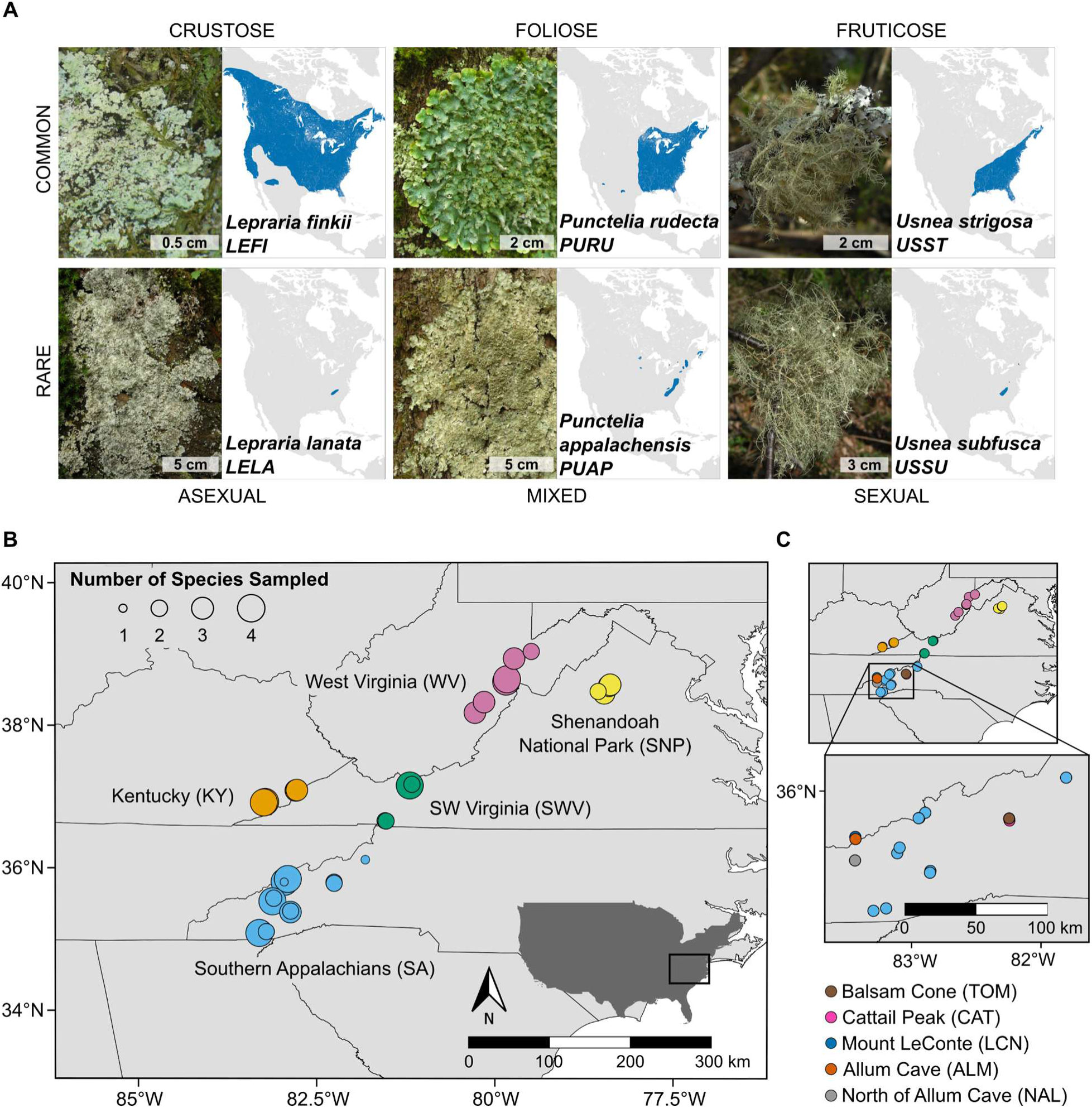
Study species and population genetic sampling sites. (**A**) Images for each study species are paired with range maps for the United States and Canada. Blue polygons represent the known ranges for each species. (**B**) Population genetic sampling sites for all study species except *Lepraria lanata.* Dots indicate sites where populations were sampled. The colors of the dots indicate landscape-level groupings for all sampling sites. The size of the dots corresponds to the number of study species sampled at each site. (**C**) Maps highlighting the population genetic sampling sites for *L. lanata.* The colors of dots represent sampling locations for the species.

### Isolation-by-distance shapes the population structure of core symbionts

Sustained restrictions in gene flow, driven by geographic distances, differences in establishment due to environmental heterogeneity, or the interaction of those factors, can lead to genetic isolation among populations (*14*). To test for isolation-by-distance (IBD) we conducted Mantel tests for each species, examining the relationship between pairwise geographic and genetic distances (Table S1). Genetic distances were quantified from neutral loci using pairwise *F_ST_* (*15*), a ratio of the variance in allele frequencies between populations in relation to total variance that is indicative of population structure (*16*). We recovered significant positive relationships between genetic and geographic distances for all three of the rare mycobiont species *Lepraria lanata* (Fig. 2G), *Punctelia appalachensis* (Fig. 2J), and *Usnea subfusca* (Fig. 2K). No significant relationships were recovered for the mycobionts *Lepraria finkii* (Fig. 2A), *Punctelia rudecta* (Fig. 2C), or *Usnea strigosa* (Fig. 2E) - all common species. Among the photobionts, we recovered significant relationships between geographic and genetic distances in algal partners associated with the rare mycobionts *P. appalachensis* (Fig. 2J) and *U. subfusca* (Fig. 2L), as well as the algal partner of one common fungal species, *P. rudecta* (Fig. 2D). Notably, there was a marginally significant trend for the *L. lanata* photobiont (Fig. 2H). To test for isolation-by-environment (IBE), we conducted partial Mantel tests using environmental differentiation as an explanatory variable for population structure, while controlling for geographic distance. Principal component analysis (PCA) of site-level environmental variables for three measures of habitat quality (light availability, stand age, and landscape structure; Table S2) and 23 climatic variables from the ClimateNA dataset (Table S3) were used to obtain environmental PCs for each species. Environmental differentiation was measured using Euclidean distances of environmental PCs. Only the rare mycobiont *P. appalachensis* and the common mycobiont *L. finkii* showed a significant relationship between population structure and environmental differentiation (Table S4).

**Figure 2.**
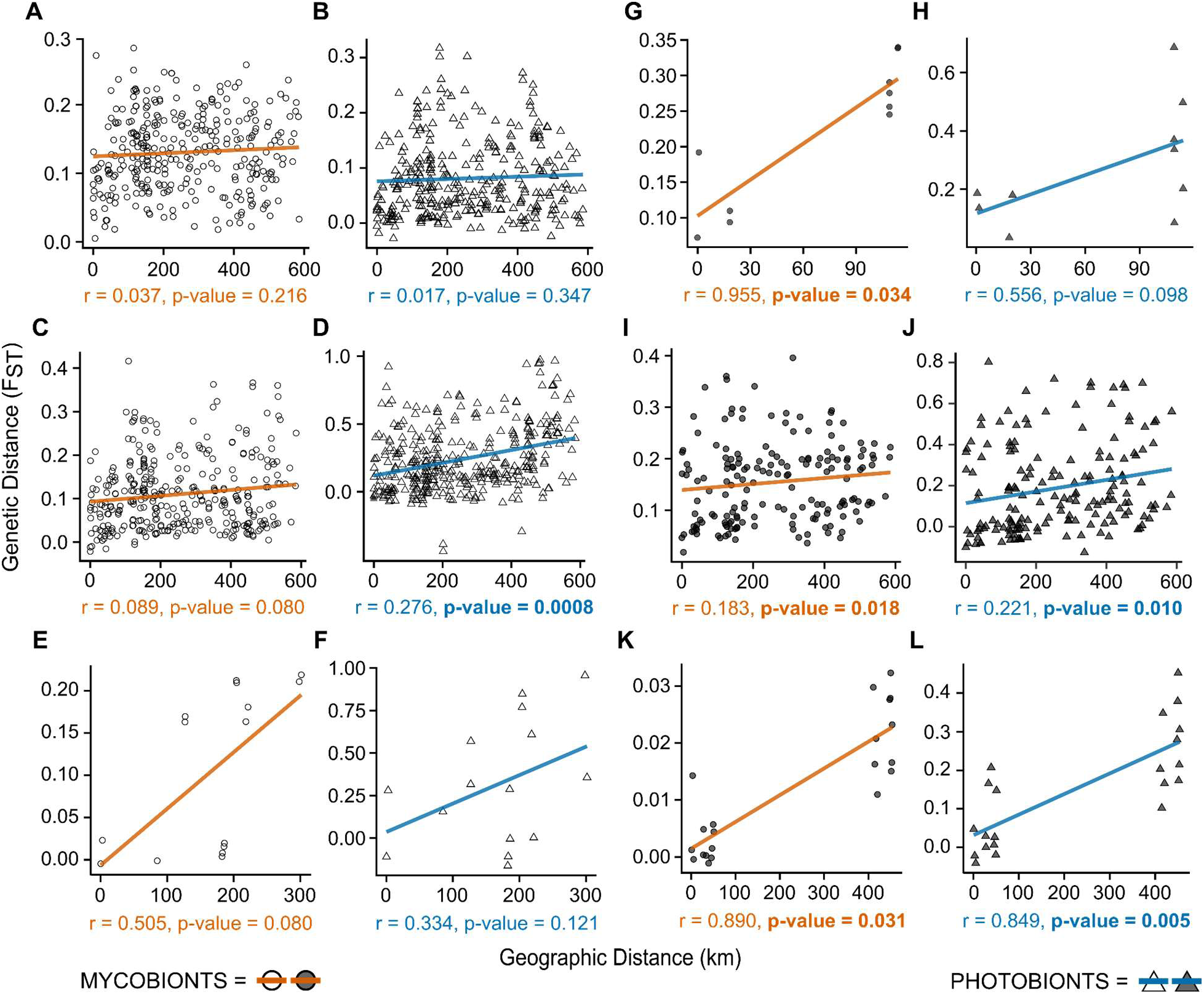
Testing for isolation-by-distance with Mantel tests. Each plot displays a Mantel test of the relationship between genetic distance and geographic distances for single lichen symbiont species. Lichen mycobiont and photobiont plots are paired with common pairs in the left column (A–F) and rare pairs in the right column (G–L). **(A)** *Lepraria finkii -* mycobiont. **(B)** *L. finkii -* photobiont. **(C)** *Punctelia rudecta* - mycobiont. **(D)** *P. rudecta -* photobiont. **(E)** *Usnea strigosa -* mycobiont. **(F)** *U. strigosa -* photobiont. **(G)** *Lepraria lanata* - mycobiont. **(H)** *L. lanata -* photobiont. **(I)** *Punctelia appalachensis* - mycobiont. **(J)** *P. appalachensis -* photobiont. **(K)** *Usnea subfusca* - mycobiont. **(L)** *U. subfusca* - photobiont. Mycobionts are indicated using circles, orange lines, and orange text. Photobionts are denoted with triangles, blue lines, and blue text. Circles and triangles for common species are empty. Circles and triangles for rare species are shaded. Pairwise genetic distances were calculated using Weir-Cockerham *F_ST_*, and geographic distance was calculated as pairwise Euclidean distances among sampling sites. Significant p-values are in bold font.

To further investigate the degree of genetic exchange among populations, we predicted the probability that each individual belonged to one pre-defined, geographically circumscribed group, or multiple groups, using discriminant analysis of principal components (DAPC). Groups were defined at the landscape level for all study species except *L. lanata* due to its extremely narrow endemism (Fig. 1B–C). We found that in the rare mycobionts and their associated photobionts single-group or one majority group belongingness was higher than in their common congeners (Fig. 3). Photobionts exhibited a greater degree of multi-group belonging than the mycobionts. Thus, while some associated photobionts exhibited significant isolation-by-distance, they generally still exhibited a greater degree of connectivity than the mycobionts. Relatively higher photobiont connectivity may reflect a true underlying pattern or may be due to photobiont datasets having been generated from organellar genomes, in contrast to our mycobiont datasets generated from whole nuclear genomes.

**Figure 3.**
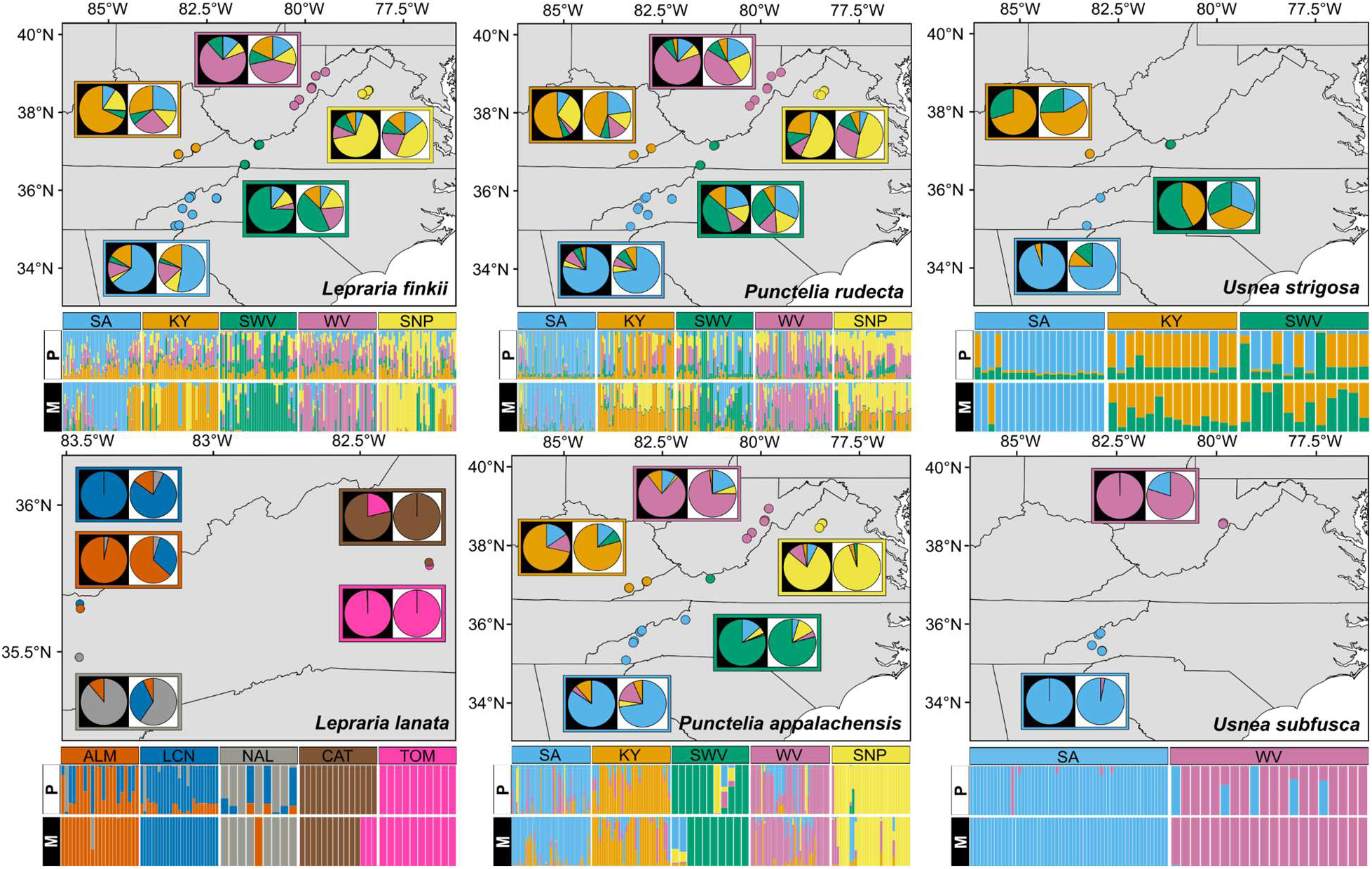
Discriminant analyses of principal components for symbionts of all six species. The bar graphs illustrate the per-individual inferred belongingness. Each vertical bar represents an individual of either the mycobiont (M) or photobiont (P). Average inferred belongingness for each region/site is shown as a pie chart outlined with the color from which the samples shown in the graph were collected. The background of the pie chart indicates the symbiont examined (mycobiont = black, photobiont = white). Regional groups were circumscribed based on geographic region, as illustrated with the dots on the maps for all species except *Lepraria lanata*. Due to the high endemism of *L. lanata,* individual sampling sites were not grouped regionally, instead each color represents a sampling site. Regional group abbreviations are as follows: SA = Southern Appalachians, KY = Kentucky, SWV = Southwestern Virginia, WV = West Virginia, and SNP = Shenandoah National Park. Site abbreviations for *L. lanata*: ALM = Allum Cave; LCN = Mount LeConte; NAL = North Allum Cave; CAT = Cattail Peak; TOM = Balsam Cone.

Our findings suggest that geographic distance is a determinant of restricted gene flow among populations of rare lichens rather than environmental barriers to establishment. Landscape genetic studies have detected IBD, IBE, or both phenomena in rare species across the web of life, including plants (*17*, *18*), invertebrates (*19*, *20*), vertebrates (*21*), and fungi (*22*). Variations in habitat types and climate along elevational and latitudinal gradients are often associated with IBE, whereas IBD has been linked to dispersal limitations and habitat fragmentation. Rarity is often associated with limited dispersal and low local abundance (*23*), which can reduce gene flow and erode genetic diversity. Comparisons of mean allelic richness (*Ar*), a measure of genetic diversity, showed that the rare mycobionts *L. lanata* and *U. subfusca,* along with the photobionts associated with all three rare mycobiont species, have significantly lower *Ar* than their common congeners and their algal partners (Table S5-6; see Supplementary Text). Maintenance of genetic diversity, along with larger effective population sizes and potential for adaptive change, are enhanced by gene flow, which buffers against bottlenecks and local extinctions (24). Gene flow enhances evolutionary potential and buffers populations against bottlenecks and local extinction by maintaining genetic diversity, larger effective population sizes, and facilitating adaptive change (*24*). Future work integrating the relative impacts of external drivers, such as habitat fragmentation, and intrinsic biological traits, such as dispersal ability and spore longevity, is essential for understanding the forces that shape population connectivity in lichens and inform the resilience of species to environmental and demographic changes.

### Rarity of one core symbiont influences genetic diversity and population differentiation of the other

We used standard population genetic metrics to compare genetic diversity and population differentiation among species, genera, and range sizes. Mean *Ar*, measured by averaging rarefied allele counts for each species’ populations, was used to quantify genetic diversity. The mean *Ar* of photobionts associated with common mycobiont species was significantly higher than the mean *Ar* of photobionts associated with rare mycobiont species (Fig. 4I). To characterize population differentiation, we used population-specific *F_ST_*, a measure of deviation from ancestral populations (*16*). Mean population-specific *F_ST_* values were not significantly different among the three mycobiont genera (Fig. 4E) or between rare and common mycobionts (Fig. 4F). However, mean population-specific *F_ST_* values were significantly higher in rare-associated than common-associated photobionts (Fig. 4L).

**Figure 4.**
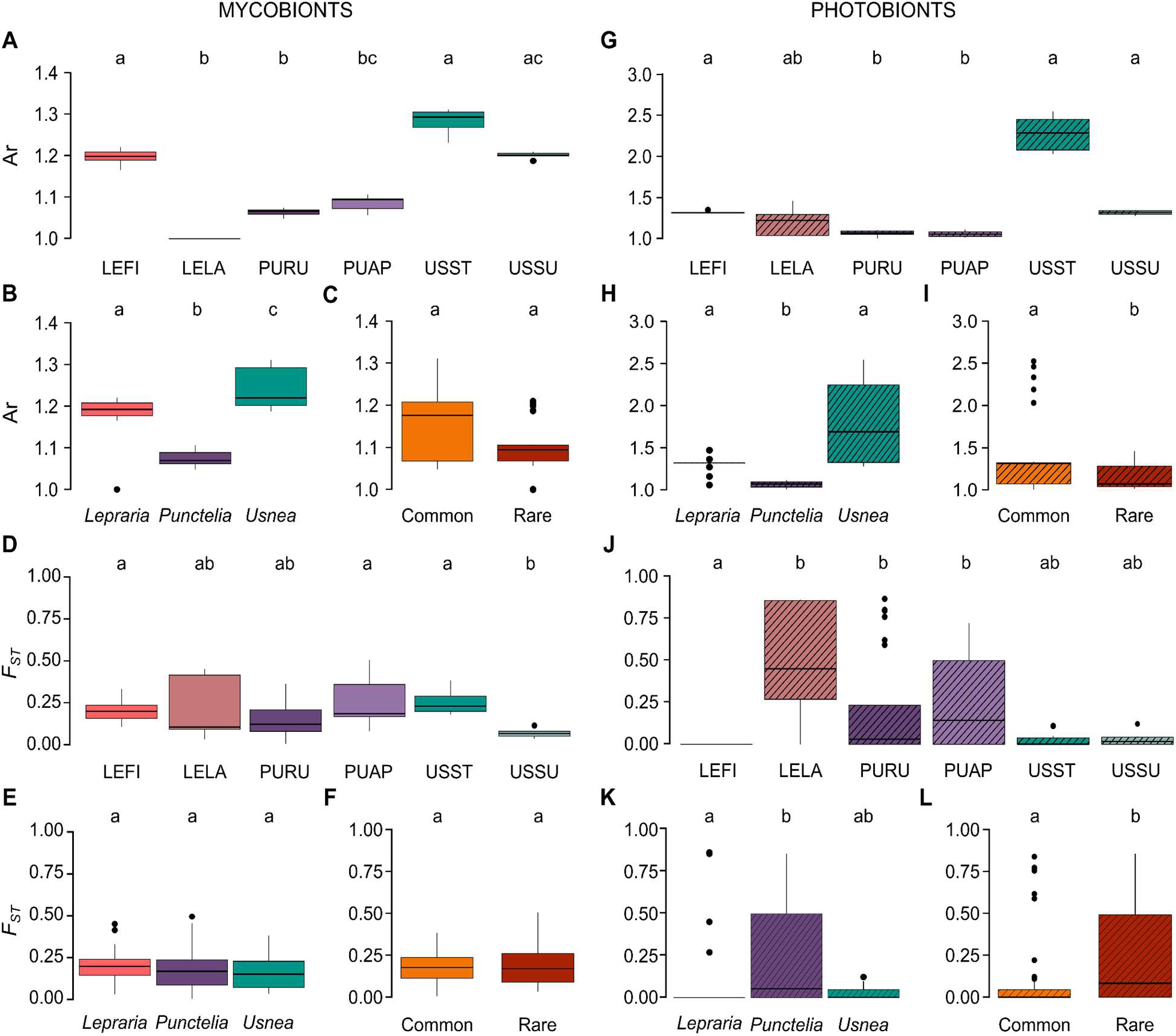
Comparisons of genetic diversity and population differentiation. (**A–C**) Comparisons of allelic richness (*Ar*), a measure of genetic diversity, among mycobiont species, and genera, and between rarity classifications. (**D–F**) Comparisons of population-specific *F_ST_*, a measure of population differentiation, among mycobiont species and genera, and between rarity classifications. (**G–I**) Comparisons of *Ar* among photobionts when grouped according to mycobiont partner species, genus, and rarity classification. (**J–L**) Comparisons of the population-specific *F_ST_* for photobionts when grouped according to mycobiont species, genus, and rarity classifications. Letters indicate significant differences in mean values based on Kruskal-Wallis Rank Sum tests and post hoc Dunn’s tests for multiple comparisons with Bonferroni corrections. Reproductive modes: LEFI/LELA (*Lepraria*) = Asexual; PURU/PUAP (*Punctelia*) = Mixed; USST/USSU (*Usnea*) = Sexual.

Symbiont dynamics comprise a spectrum of partnerships ranging from those that are taxonomically and genotypically flexible to those that are specific to a single genotype or species (*25*). Our finding that photobionts associated with rare mycobionts exhibit lower genetic diversity and greater differentiation among populations suggests a greater incidence of symbiont specificity in all rare mycobionts. This contrasts with the mycobiont comparisons, which recovered no significant differences in genetic diversity or population differentiation when data were faceted by rarity (Fig. 4F). Thus, the rarity of the mycobiont may be limited by both dispersal ability, as suggested by the tests for IBD, and symbiont specificity.

The availability and specialization of mycorrhizal symbionts have been suggested as mechanisms mediating the distributions of rare plants, though few studies have addressed species-specific interactions (*26*). Comparisons of the highly specific symbiont *Pseudotsuga japonica*, Japanese Douglas-fir, and its mycorrhizal associate *Rhizopogon togasawarius*, both endangered Japanese endemics, recovered congruent patterns of relative genetic diversity and differentiation in the two species, however genetic diversity was lower and differentiation was greater in *R. togasawarius* at all sites (*27*). In lichens, symbiont flexibility has been attributed to the ecological and biogeographic amplitude of the mycobiont by enabling selective partnerships with locally adapted algal species or genotypes across a variety of environmental conditions (*28*, *29*). Studies examining Cnidarian-Symbiodinaceae partnerships attribute thermotolerance (*30*) and range size (*12*) to symbiont flexibility and stability and overall resilience in response to bleaching events to symbiont fidelity (*31*, *32*), underscoring the benefits of both specific and flexible symbiont dynamics.

### Fungal recombination rates reflect observed reproductive phenotypes and highlight the role of asexual reproductive diversity in symbiont genetic diversity

As for many obligate symbioses, lichens have complex life cycles that involve diverse reproductive strategies and both aposymbiotic and symbiotic dispersal (*33*). We studied lichens that spanned the spectrum of observed reproductive phenotypes from strictly asexual symbiotic propagules (*Lepraria finkii*, *L. lanata;* (*34*)), to a mixed mode of frequent symbiotic asexual propagules combined with infrequent fungal sexual reproductive structures (*Punctelia appalachensis*, *P. rudecta;* (*35*)), and strictly sexual fungal reproductive structures (*Usnea strigosa, U. subfusca;* (*36*)). To assess the rates of linkage and clonality, we tested for differences in the index of association (Ia), a measure of linkage among loci and clonality, among mycobiont species, genera, and rarity classifications using methods as described above for *Ar* and *F_ST_*.

We found that the observed reproductive phenotype for each lichen was reflected in the recombination rates of the mycobiont. The Ia was nearly zero in the *Usnea* mycobionts, indicating very high rates of recombination (Fig. 5B–C; Table S7). The Ia was higher in the *Punctelia* mycobionts, and highest in the *Lepraria* mycobionts, suggesting stepwise lower rates of recombination, though the latter means were not significantly different. In the *Lepraria* and *Punctelia* mycobionts, the results of the Ia comparisons were supported by lower measures of *Ar,* which are suggestive of greater local, clonal reproduction in these genera when compared to *Usnea* mycobionts (Fig. 4B). These results collectively support expected outcomes wherein sexually reproducing species are more genetically diverse and have greater gene flow (*37*, *38*).

**Figure 5.**
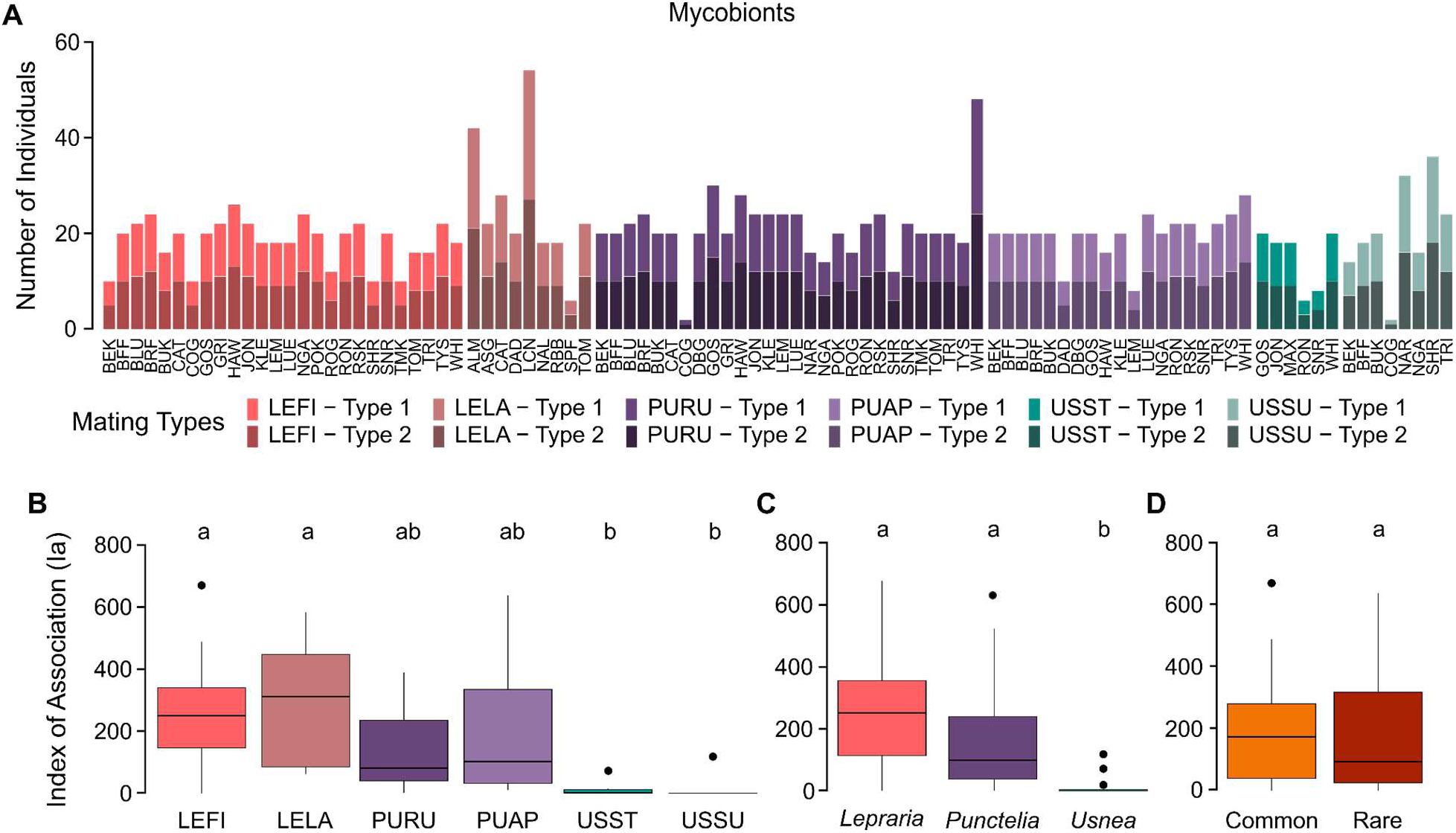
Mycobiont reproduction-related metrics. (**A**) Frequency of mating type identities present in each population for all species. Each bar represents a population. Lighter colored bars represent the presence of *MAT1-1*, and darker colored bars represent the presence of *MAT1-2*. Both mating types are present in every species at nearly equal frequencies. (**B-D**) Distribution of the population-level index of association, a metric used to infer clonal reproduction, for each mycobiont species and groupings by genus and rarity classification. Letters indicate significant differences in mean values based on Kruskal-Wallis Rank Sum tests and post hoc Dunn’s tests for multiple comparisons with Bonferroni corrections. Reproductive modes: LEFI/LELA (*Lepraria*) = Asexual; PURU/PUAP (*Punctelia*) = Mixed; USST/USSU (*Usnea*) = Sexual.

Because of the low rates of recombination in *Lepraria* and *Punctelia* mycobionts that were not significantly different from each other, we expected both genera to have similar and low photobiont genetic diversity, based on the dominance of asexual reproduction via symbiotic propagules in these taxa. However, we found that photobiont genetic diversity (*Ar*) was significantly lower in *Punctelia* than in *Lepraria* (Fig. 4H).

Subtle differences in phenotypes of structures that function as symbiotic propagules have been linked to photobiont diversity, potentially explained by the presence of a protective covering on some propagule types (*38*). We extend this hypothesis to incorporate a suite of fundamental anatomical, chemical, and developmental differences that may facilitate photobiont exchange. The lichens we studied that produce symbiotic propagules include the two major types of structures: extensions or outgrowths, versus disintegrated bundles, of the parent holobiont. Our focal *Punctelia* produce the outgrowth-type, which are covered in a protective outer layer of dense fungal cells and polysaccharides that limit adhesion, do not readily detach from the parent body, and are comparatively large. In contrast, *Lepraria* produces the bundle-type, which are composed of a loose web of fungal filaments surrounding a core of algal cells, all of which are coated in adhesive hydrophobins, readily detach from the parent body, and are comparatively small (*39*). We suggest that lichens with the bundle-type are predisposed to exchanging and incorporating a greater diversity of photobionts than the outgrowth-type, which explains the higher photobiont genetic diversity that we detected in *Lepraria* versus *Punctelia*.

For obligate symbioses, clonal co-dispersal of symbionts can retain favorable partner combinations and shift the balance of the typical evolutionary benefit of sexual reproduction towards asexuality (*40*). Lichen resynthesis during sexual reproduction promotes photobiont diversity. Symbiotic propagules have been assumed to lead to homogeneous holobiont populations that have lower long-term fitness due to extreme symbiont specificity and lower genetic diversity. This perspective overlooks the diversity of anatomy and function in asexual reproductive structures and fails to account for observed evolutionary and ecological patterns in asexually reproducing organisms (*41*). Our results suggest a complex scenario wherein asexual symbiotic co-dispersal of holobionts can lead to extreme differences in population genetic outcomes, and the genetic cost of asexuality is balanced by the benefits of symbiont reshuffling.

### Observed reproductive phenotypes do not reflect ratios of suitable mates

While reproductive morphologies were predictive of recombination rates, they were not predictive of mating type ratios. In Ascomycota, mating type genes code for transcription factors that control the expression of pheromones and pheromone receptors that are required for recognition of suitable mates (*42*). Most species in Ascomycota have bipolar breeding systems with two distinct mating types, and sexual reproduction occurs predominantly between two different mating types. We hypothesized that both mating types may not be present in *Lepraria* because of their complete lack of sexual reproductive structures, though previous research has recovered a single complete set of mating type genes from species in this genus (*43*). Our hypothesis was not supported; we instead found both mating types to be present in both *Lepraria* species at relatively even ratios (Fig. 5A). Because both species in the genus *Punctelia* reproduce sexually infrequently, we hypothesized that their mating type ratios would be skewed and that populations would frequently only host one mating type. Again, our hypothesis was not supported as both mating types were present in nearly every population, and they were present at nearly even ratios. The only genus that we hypothesized would have even mating type ratios and both mating types present in every population was *Usnea*, and our hypothesis was supported. Overall, there is no significant difference in the mating type proportions among the species, despite their significantly different reproductive morphologies and recombination rates (Fig. 5A).

The lack of congruence between mating type ratios and recombination rates that match reproductive phenotypes suggests that mating type proteins in Lecanoromycetes may not always function primarily in mate recognition and highlights the gap in the understanding of the molecular and cellular biology of mating systems. Fungal sex pheromone receptors are involved in multiple signaling pathways, including responding to plant secreted compounds (*44*), and functioning in quorum sensing and conidial germination quenching (*45*). Transcription factors encoded by the mating type genes are known to have diverse downstream targets as well (*46*, *47*). In *Ulocladium botrytis,* mating type genes control asexual traits, such as conidia size and number (*48*). These transcription factors have also been found to impact the production of carotenoids, a phytoresponsive secondary metabolite in *Fusarium verticilliodes* (*49*). While we found mating types to be present in even ratios across different populations and reproductive phenotypes, other studies of asexual fungal pathogens have found that mating type ratios can vary across populations, suggesting that different idiomorphs of the mating types may provide fitness advantages in specific contexts (*50–52*).

### Heterokaryon incompatibility and protein-protein interaction domains are under selection in mycobionts

Latent factor mixed models were used to identify loci with outlier allele frequencies, which suggests that they are under selection. Outlier loci were recovered for most species-environmental variable combinations for both mycobionts and photobionts (Fig. S1). The total number of outlier loci ranged from 25 in the mycobionts *P. appalachensis* to 5,944 in *U. strigosa,* and three in organellar genomes of photobionts associated with *P. rudecta* and *L. finkii* to 654 in organellar genomes photobionts associated with *U. strigosa* (Table S8).

To infer the function of mycobiont outlier genes and identify any function-environment relationships, we built Gene Ontologies (GO) for each species and implemented gene ontology enrichment analyses for each species-environmental variable combination and for all outliers identified in a species. Most of the significant GO terms related to regulating gene expression through DNA interactions (e.g., helicase activity, histone methyltransferase complex, pericentric heterochromatin, and regulation of DNA-templated transcription), RNA interactions (e.g., regulation of RNA biosynthetic process, regulation of RNA metabolic process), or proteins (e.g., Cul4-RING E3 ubiquitin ligase complex; Fig. S2, Table S9). Other terms included protein localization, sexual reproduction, anatomical structure morphogenesis, and multiple biosynthetic processes. It was not possible to assign a putative GO term to nearly half of the genes in each genome, so the power of our analyses were limited. Despite this inherent limitation when working with non-model organism functional genetics, it was clear that genes involved in fundamental cellular processes were under selection in every mycobiont investigated.

To make finer-scale functional genetics inferences, we used homology searches against the entire NCBI database and assigned functions or structures to the most significant outlier loci from each species-environmental variable combination. We were not able to assign putative functions to 39 of the 138 (28%) genes investigated. Of the remaining genes, there were five functions or structures wherein three or more species were discovered to have an outlier locus sharing that category (Fig. 6A). The most frequently recovered structures and functions in mycobionts were ankyrin repeat domains, histone-interacting proteins, heterokaryon incompatibility domain, secondary metabolism, and secondary metabolite biosynthesis (Fig. 6A).

**Figure 6.**
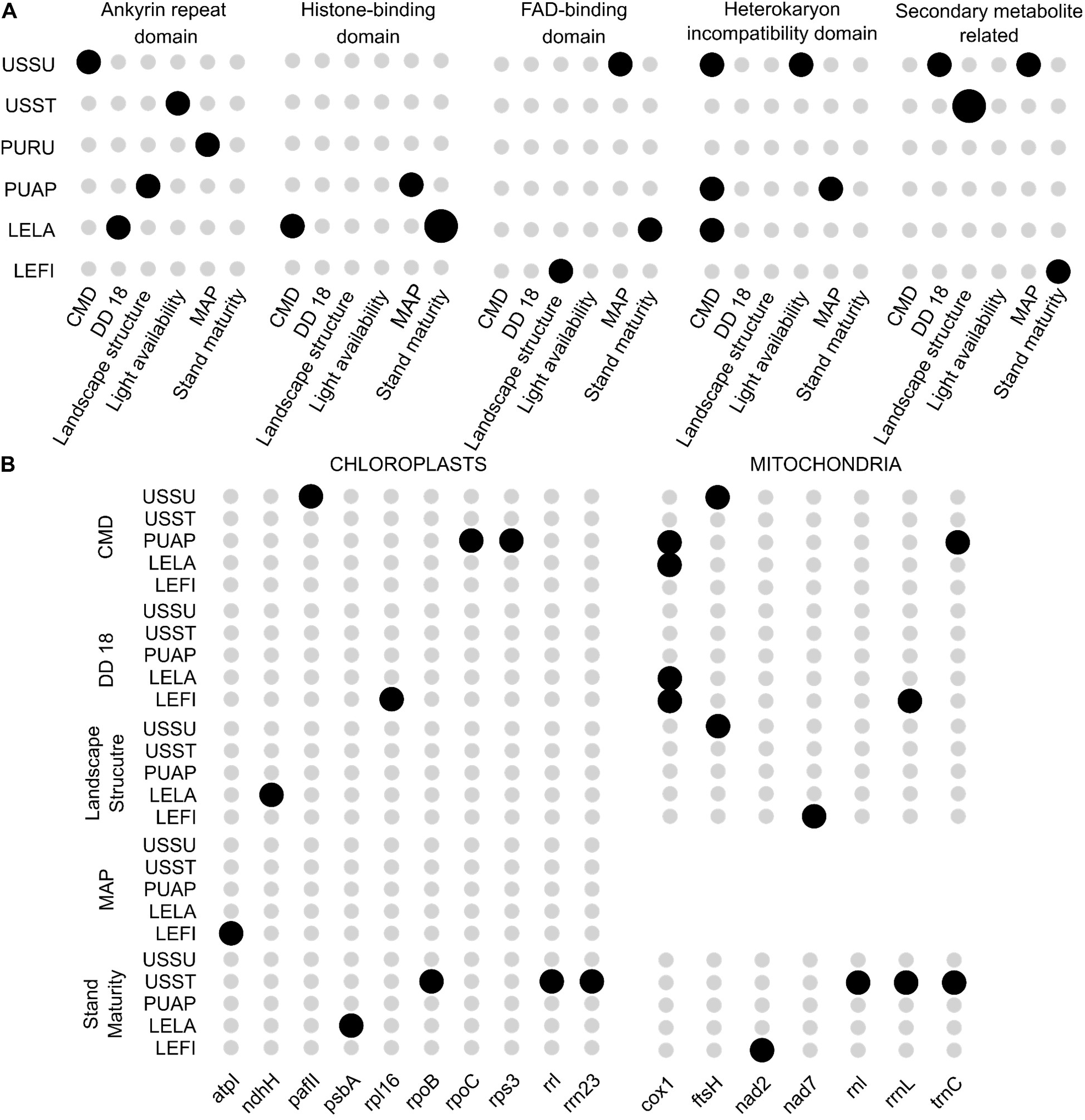
Functional genetics results. Dotplot showing all recovered species-environment-gene structure and function among the top outlier loci for the (**A**) mycobionts and (**B**) photobionts. The larger black dots indicate when two genes were found in a specific category, smaller black dots indicate one gene was found, and gray dots indicate that no genes fit that classification. White space indicates that no outlier loci were recovered for a specific environmental variable.

The recovery of GO terms related to expression regulation, intracellular movement, and signaling all suggest that fundamental cellular and genetic processes are under selection in response to environmental pressures. Further, histone-binding domains were the top outliers for multiple species and environmental relationships. Studies of differential environmental adaptation of the fungi *Suillus brevipes*, *Phellopilus nigrolimitatus*, and *Lasallia pustulata*, the latter another lichen, similarly found that genes regulating gene expression were frequently under selection (*53–55*). SNPs in helicase genes were specifically correlated with temperature variables in *S. brevipes* and *P. nigrolimitatus* (*53, 54*). Helicase SNPs were also identified in *L. pustulata* and were associated with altitude, which the authors speculated related to temperature differences and adaptation to cold shock (*55*). *Phellopilus nigrolimitatus* and *L. pustulata* showed evidence of selection in genes related to signal transduction and cellular transport and growth, providing further evidence that integral cellular processes are influenced by environmental changes (*54*, *55*).

The underlying genetics and cell biological mechanisms involved in fungal responses to stimuli and environmental pressures under short and long timeframes are complex (*56*). In fungal life cycles cells transition from a haploid state to multinucleate dikaryons as a precursor to sexual reproduction. Recovery of heterokaryon incompatibility domain-containing genes (*het*) under selection in rare but not common mycobionts further underscores the importance of reproductive processes in population genomics dynamics. Because allorecognition is core to fungal functionality, *het* genes are hypothesized to be under balancing selection that supports maintenance of genotype diversity (*57*). Secondary metabolites are essential for fungi to respond to and navigate their environment (*58*). We found four genes involved in secondary metabolism, from four different species, among the top outlier loci. Three of the genes were Type 1 Polyketide Synthases, large enzymes responsible for the biosynthesis of polyketides. Polyketides are a diverse class of secondary metabolites that includes antibacterial and antifungal compounds as well as mycotoxins (*59*). The most frequently recovered structures were ankyrin repeat domains, which is likely due to their overall high frequency among proteins, and expansion of protein families containing ankyrin domains in lichenized fungi (*60*). Ankyrin repeat domains are frequently involved in protein-protein interactions (*61*), including mediating symbiont interactions (*62*). Proteins with FAD-binding domains were also repeatedly recovered. Similar to ankyrin repeats, FAD-binding domains are present in many proteins with diverse functions, including regulating photomorphogenesis and secondary metabolite biosynthesis (*63*), and heterokaryon dynamics (*64*). Because studies of functional genetics in non-model organisms are inherently limited by the lack of genes with known functions, reciprocal field-laboratory research is essential to fundamentally disentangle adaptive responses over timescales and degrees of complexity not attainable in laboratory settings.

### Plastid-encoded functions play a critical role in symbiotic algal adaptation

Homology searches of photobiont outlier loci identified 10 chloroplast genes and seven mitochondrial genes under adaptive selection in response to variations in habitat quality and climate across all the species studied (Fig. 6B). In the photobionts associated with *L. finkii* and *L. lanata,* several chloroplast genes known to encode proteins involved in photosynthesis were under selection (*65*). These include *atpL* (adenosine triphosphate synthase or ATP synthase) in response to mean annual precipitation (MAP); *psbA* (photosystem II) in response to forest stand maturity, and *ndhH* (NADH dehydrogenase) in response to landscape structure. Similarly, a study of *Breviolum psygmophilum*, the algal symbiont of the temperate coral *Astrangia poculata*, identified genes encoding components of photosystems I and II under selection along a temperature gradient, suggesting temperature and light may be drivers of adaptive differentiation among *B. psygmophilum* populations (*66*).

Additional chloroplast genes found to be under selection were integral to self-replication and protein synthesis. These included *rpoB* and *rpoC,* which encode the β- and β’-subunits of RNA polymerase, respectively, as well as *rpl16* and *rps3,* which encode ribosomal proteins L16 (large subunit) and S3 (small subunit) (*65*, *67*). Adaptive selection on these genes varied among photobionts, but overall demonstrates that environmental variability, specifically climate moisture deficit, degree days <18 °C, and forest maturity, may drive adaptation in organellar transcription and translation (Fig. 6B). A laboratory study of the model green algae *Chlamydomonas reinhardtii* found that repressing the genes *rps12* (encoding ribosomal protein S12) and the *rpoA* (encoding the ∝-subunit of RNA polymerase) not only halted cell growth but initiated a negative feedback loop that altered nuclear gene expression, including genes involved in chloroplast biogenesis and stress response (*68*).

Among the mitochondrial genes recovered, *cox1* was the most frequent (Fig. 6B). Cytochrome c oxidase subunit I is integral to the synthesis of ATP in mitochondria during oxidative phosphorylation. Outlier loci within the *cox1* genes of the photobionts associated with *L. finkii, L. lanata,* and *P. appalachensis* were under selection in response to degree days <18 °C and climate moisture deficit, suggesting energy production adaptations were in response to temperature and drying conditions.

Organellar genomes, particularly in algae, are known to be sites of rapid and dynamic evolution (*69*, *70*). A majority of the genes we identified under selection were within chloroplast genomes, underscoring the role of plastid-encoded functions, such as photosynthesis and self-replication, in the environmental adaptation of symbiotic algae. In the context of photobionts associated with closely related mycobiont species, particularly those with vastly different range sizes, the observed adaptive variations in a core metabolic process (i.e., photosynthesis) may have driven niche divergence. Moreover, our findings highlight the sensitivity of transcription and translation to environmental variation with serious implications for the growth and survival of photobionts.

### Towards a conceptual model for the population genomics of obligate symbioses

Comparative frameworks serve as a powerful approach for identifying patterns among closely related species (*71*). Using landscape genomics and multi-level comparisons among and between pairs of congeneric mycobionts and their photobiont partners, we connected reproduction and spatial patterns of rarity to symbiont dynamics and micro-evolutionary processes. Those connections facilitated the identification of patterns in genetic variation and symbiont specificity across obligate symbioses with differing range sizes and reproductive strategies (Fig. 7A). In many cases, we recovered distinct and often contrasting patterns among and between range sizes and reproductive strategies; in other cases, those patterns were indistinct and highly species-specific. Cumulatively, our work highlights the need to develop and implement eco-evolutionary frameworks that capture the complex biotic interactions embodied by obligate symbioses and address the influence of population-level spatial and demographic patterns. Here we present a novel, extended eco-evolutionary framework for obligate symbioses, developed by drawing on established frameworks (*72*, *73*) and our own empirical data, to guide future studies of symbiotic systems (Fig. 7B). Despite the advances made by metagenomic sequencing, disparities in our knowledge of the ecology and evolution of symbiotic partnerships persist, with some symbionts left understudied. By embracing the interconnectivity of symbiotic partnerships within extended eco-evolutionary frameworks, it is possible to advance our understanding of the highly intricate and interacting biological systems that are obligate symbioses.

**Figure 7.**
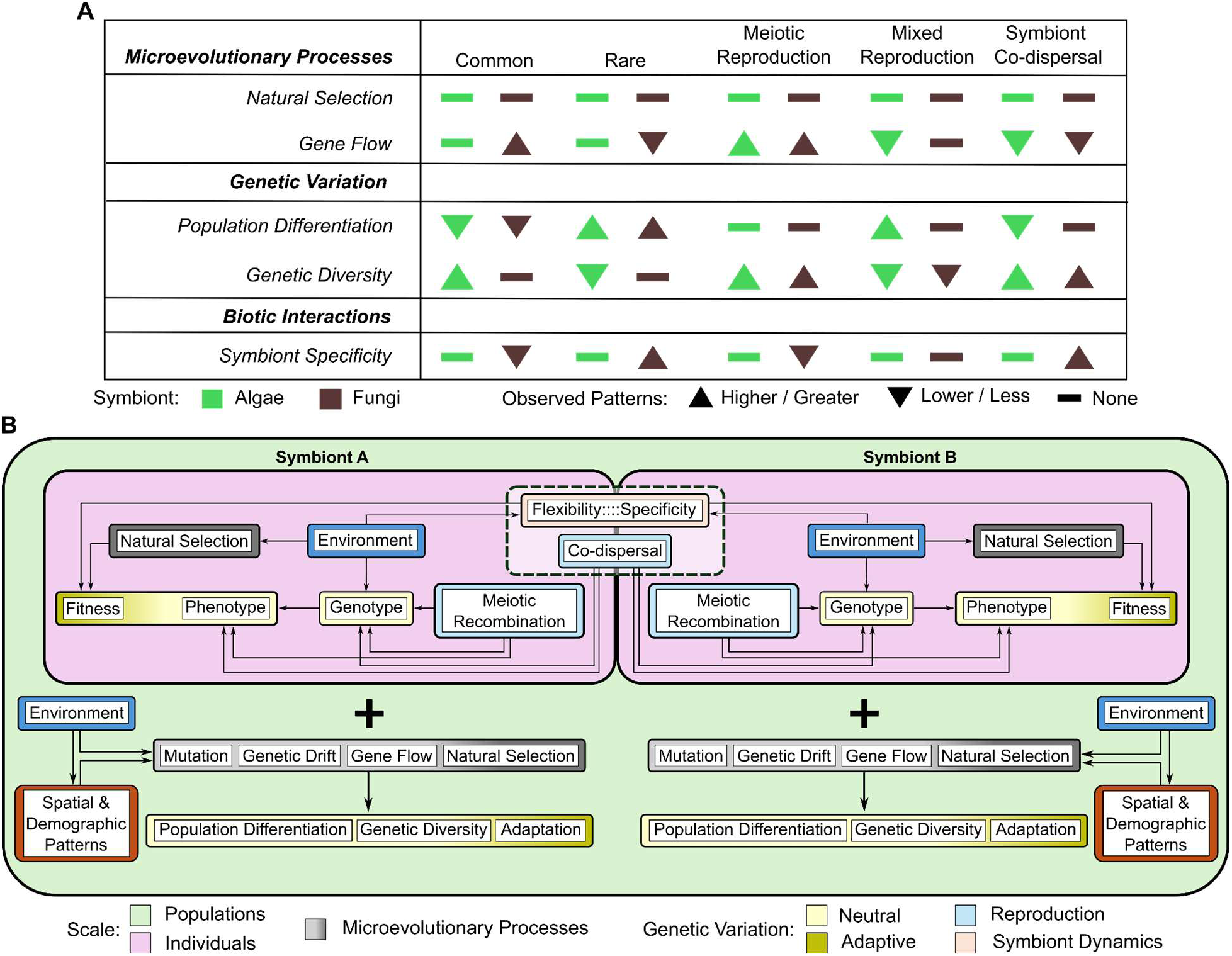
Eco-evolutionary framework for obligate symbioses using lichens as a case study. **(A)** Patterns identified with empirical data connecting organismal distributions and reproductive life history. For example, gene flow is limited in rare organisms with asexual reproduction (symbiont co-dispersal) due to isolation by distance and limited opportunities for genetic exchange among individuals. (**B**) Interactions among biotic and abiotic factors lead to detectable genetic patterns (yellow and golden boxes) in neutral loci (population differentiation and genetic diversity) and loci under selection (adaptation). Interactions among symbionts are influenced by the environment and individual genotypes leading to a phenotype with an associated fitness. At the population level, those phenotypes are acted upon by microevolutionary patterns to produce observable patterns within the organismal genomes. Symbiont dynamics and co-dispersal act within this framework as a bridge connecting the evolutionary forces influencing individuals. The magnitude of influence these factors have on the evolutionary trajectory of both symbionts is dependent on the degree of symbiont specificity versus selectivity and the mode of co-dispersal.

## Supporting information

All_supplements

## Acknowledgments

We appreciate fieldwork support from Eli Denzer, Bubba Pfeffer, Amanda Chandler, Jason Hollinger, and Seth Raynor. We also appreciate herbarium specimen digitization and processing support from Eli Denzer. Species photographs used in Figure 1A were taken by Jason Hollinger. Thanks to Olivier François and Sean Schoville for providing technical support and feedback for the implementation of latent factor mixed models.

## Funding

National Science Foundation DEB #2115191 (JLA) National Science Foundation DEB #2436848 (JCL) Eastern Washington University - Biology Department, Cheney, Washington, USA Highlands Biological Station, Highlands, North Carolina, USA

## Author contributions

JLA and JCL developed the intellectual framework and acquired funding. JLA, STS, and JCL conceptualized the manuscript. JLA and JCL executed the fieldwork. JP, STS, MP, and KN processed voucher specimens. JP, JLA, KN, and STS performed molecular laboratory work. STS, JLA, EML, and JED analyzed the data. STS and JLA created and composed figures. JLA and JCL curated publicly deposited datasets. STS, JLA, JP, EML, PS, and JCL drafted the manuscript. STS, JP, JCL, PS, MP, EML, JEDM, and JLA collaboratively reviewed, edited, and revised the manuscript. JLA, JCL, and PS supervised personnel and administered the project. JLA led the project.

## Competing interests

Authors declare that they have no competing interests.

## Data and materials availability

Sequence data are available from the National Center for Biotechnology Information. Raw reads of population-level sampling are available from BioProject PRJNA1270084, and raw reads for reference genomes are available from BioProject PRJNA1270085. Code is available on GitHub at https://github.com/jallen73/Lichen_popgen. All voucher specimens are deposited at the New York State Museum.

## Supplementary Materials

Materials and Methods

Supplementary Text

Figs. S1 to S2

Tables S1 to S15

References (*74* - *107*)

## Notes

https://www.ncbi.nlm.nih.gov/bioproject/PRJNA1270085/

https://www.ncbi.nlm.nih.gov/bioproject/PRJNA1270084/

https://github.com/jallen73/Lichen_popgen

### Competing Interest Statement

The authors have declared no competing interest.

