## Supplementary material for "Rarity and reproductive biology drive population genomics of obligate symbioses": All_supplements

**The PDF file includes:**

Materials and Methods
Supplementary Text
Figs. S1 to S2
Tables S1 to S15
References (*74 - 107*)

### Materials and Methods

Sample collecting, site characteristics, and climate data

Large, whole thalli of each study species were collected to generate reference genomes (*74*). Thallus fragments of each species were collected from 1-hectare (ha) sites spanning the central and southern Appalachian Mountains from 2018 - 2022 for population genomic data generation (Table S10, S11; Fig. 1B). Whole thalli and thallus fragments were collected using nitrile gloves and sterilized forceps or wood chisels, then cold-stored in paper bags at -20 °C before processing for DNA extractions.

Data characterizing light availability, stand maturity, and overall landscape structure for each 1-ha site were produced using protocols described (*75*) and modified for use in previous studies (*76*) (Table S12). At each site we visually estimated rock cover (RC; the percentage of rock cover), large trees (LT; the percentage of large canopy trees), canopy cover (CC; percentage of living, native canopy trees), understory (US; percentage of native understory plant species), weediness (WED; the percentage of non-native and invasive plants), leaf litter (LIT; percentage of leaf litter from native versus non-native trees), loggyness (CWD; the percentage of coarse woody debris), recruitment (REC; the percentage of native tree species recruitment), patch size (PS; the size of contiguous forest blocks), neighborhood (NH; physical connectivity between areas of native vegetation), and distance to core (D2C; distance to 50 ha core areas of natural habitat). We then summed the values for PS, NH, and D2C to create the variable landscape structure; RC and CC to create light availability; and LT, US, REC, and CWD to create stand maturity (Table S2). Those characteristics reflect anthropogenic disturbance at the site and landscape levels.

We assembled a climate dataset for this research consisting of 23 variables from the ClimateNA dataset (Table S3). Measurements of variables from ClimateNA, which comprises annual historical normals spanning 1991 - 2020 (*77*), were extracted from rasters using the extract function from the R package tidyr v1.3.1 (*78*) for all population genetic sampling sites. Values for mean annual precipitation (MAP), climate moisture deficit (CMD), and degree-days below 18°C (DD_18) were used in latent factor mixed models.

Genome generation

Preparation for DNA extractions involved manually removing plant material, debris, endophytes, and visible necrosis from thallus fragments. DNA extractions using thallus fragments and whole thalli were performed using QIAGEN DNeasyTM Plant Kits according to the manufacturer's protocols for plate-based extractions and single-sample extractions, respectively. DNA extracts were quantified using Invitrogen Qubit high-sensitivity double-stranded DNA (dsDNA HS) assays and stored at 0oC until shipment for library preparation and sequencing. Long-read sequencing was conducted at the Cold Springs Harbor Laboratory (Laurel Hollow, New York, USA) on the Oxford Nanopore Technologies PromethIon platform, and basecalling was conducted using 'Guppy' v5 in super-accurate basecaller (SUP) mode. Reference genomes were assembled with Flye v2.9.3 (*79*), and metagenomic filtering was completed according to established workflows (*80*). Annotations were completed using funannotate v1.8.17 (*81*). Short read sequencing was conducted by GENEWIZ (Azenta Life Sciences, South Plainfield, New Jersey, USA). Whole genome shotgun sequencing (WGS) at 20x coverage of 60 Megabase genomes produced 150 base pair (bp), paired-end reads using an Illumina platform. For all six mycobionts associated with target lichen species, reference genomes were sequenced, assembled, and annotated according to established protocols (*43*, *74*, *80*). Voucher specimens for reference genomes are held at the Eastern Washington University herbarium. Population genomic vouchers, along with digital images of every specimen, are held at the New York State Museum (Table S13).

Variant calling and filtration

We used 'VCFtools' to generate summary statistics for the proportion of missingness per individual and mean variant depth for all genomic datasets (*82*). For non-coding mycobiont datasets, individuals with greater than 20% missingness were removed, then filtered for locus-based missingness (threshold = 0.9) and depth using a single operation. Thresholds for maximum and minimum depth filters were determined for each species, respectively, using summary statistics. Mycobiont coding sequences displayed minimal missingness and thus were only filtered for depth using species-specific maximum and minimum thresholds. Photobiont organellar genomic (POG) datasets were filtered for missingness (threshold = 0.85) and depth, then any remaining individuals with greater than 15% missingness were removed from the dataset. Due to the limited number of variants in the POG datasets and the influence of variant count on genetic diversity metrics, POG species datasets with >6,000 variants were randomly thinned to ensure all datasets contained ~5,000 or fewer variants. All POG datasets filtered and thinned to ~5,000 variants were used to generate population genetic diversity metrics and perform discriminant analysis of principal components and Mantel tests.

Filtered non-coding mycobiont datasets were tested for linkage disequilibrium (LD) and subsequently pruned of linked loci. The squared correlation coefficient (*r2*) between all noncoding loci was calculated with PLINK using a window size of 100 variants to evaluate genomic regions spanning up to 1000 kb (*83*). The average LD across 100 kb bins was calculated using a custom script (*84*). Linkage decay was plotted using the R package 'ggplot2' v3.4.4 (*85*). Linked loci were pruned from each dataset using the independent pairwise variant pruning function. The mean *r2* was used to determine the *r2* pruning threshold for each species. Our resulting noncoding mycobiont datasets ranged between 10,000 and 20,000 variants for each species.

To generate POG and mycobiont coding datasets for explicit use in hypothesis testing for genotype-environment association (GEA) analyses, we applied additional filters. First, individuals representing duplicate multi-locus genotypes (see Multi-locus Genotypes) were removed from each species dataset. Next, each dataset was randomly thinned and filtered to include variant sites with a minor allele count (MAC) greater than or equal to a target threshold. Mycobiont coding datasets were thinned to remove any two sites within 500 bp and filtered using a MAC threshold representing 0.05 (5%) of the total number of individuals in the dataset. Subject to random thinning before this step, POG datasets were filtered using a MAC threshold representing 0.03 (3%) of the total number of individuals in the dataset before GEA analyses.

For variant counts for all datasets before and after filtering, see Table S14.

Multi-locus Genotypes

Variant call files (VCFs) were converted to genclone objects, enabling the storage of additional information on MLGs. We applied a multilocus genotype (MLG) filter using the mlg.filter function from the package 'poppr' v2.9.6 (*86*) with bitwise distance and the farthest neighbor algorithm for population-level genotypic analyses (*87*). Bitwise distance is calculated using binary variations in the fraction of alleles between samples. This metric is robust to missing data. Thresholds for collapsing MLGs (a.k.a., clonal lineage thresholds) were determined using the filter_stats function with bitwise distance, followed by the cutoff_predictor function to identify the largest gap in Euclidean genetic distances for each species. Filtered genclone objects and individuals representing unique MLGs were used in downstream analyses.

<u>Statistical analyses</u>
Discriminant analysis of principal components (DAPC) was performed on genclone objects representing unique MLGs using the package 'adegenet' v.2.1.10 (*88*). This approach is well-established for inferring genetic differentiation within populations of both sexually and asexually reproducing organisms (*89*). The optima.a.score function in 'adegenet' was used to test for bias in preliminary runs on a species-specific basis. Results were visualized using composition bar plots displaying the predicted proportions of group assignments.
Mantel tests were conducted using the mantel function in the package 'ecodist' 2.1.3 (*90*) with 10,000 permutations. Pairwise genetic distances were calculated using the pairwise.WCfst function in the R package 'hierfstat' v0.5-11 (*91*). We used the geodist function in the package 'geodist' v0.0.8 (*92*) to calculate the geodesic distance (km) between sites. To characterize environmental differences among sites, principal component analyses (PCA) were run on matrices of environmental variables for each species using the prcomp function in the 'stats' package v3.6.2 (*93*). Screeplots were used to determine the number of PCs to retain for dissimilarity indices. Dissimilarity indices for population structure, geodesic distances, and environmental differences were calculated using the vegdist function with a Euclidean dissimilarity index in the 'vegan' package v2.6-2 (*94*) and then used to perform Mantel tests. We reported Pearson's correlation coefficient (r) and two-tailed p-values obtained.
Population-specific Fst was calculated using the betas function of the 'hierfstat' package (Table S15). The function allelic.richness from the same package was used to quantify rarefied allele counts for each population, which were then averaged to obtain mean allelic richness (Table S15). Genclone objects filtered for unique MLGs were used to test for linkage using the poppr.basic.stats function from the 'poppr' package to measure the index of association for each population.

Kruskal-Wallis tests were performed using the 'stats' package v3.6.2 (*93*). The 'FSA' package v0.9.6 (95) was used for post hoc Dunn's tests with Bonferroni corrections. Non-parametric tests were chosen after using the Shapiro-Wilk tests from the 'stats' package v3.6.2 and Levene's tests from the 'car' package v3.1-1 (*96*) to verify that our datasets exhibited nonnormality and heteroscedasticity. Populations TYS and BEK contained NaN values for Fst and were removed from all datasets before comparative statistical analyses. The above statistical analyses and filtration for multi-locus genotypes were executed using R v4.2.3 (*93*).

<u>Characterizing mating types</u>
To determine the mating types of each individual, the protein sequences for Aspergillus nidulans Apn2 and Sla2 were used as queries to blast against all annotated proteins from each species'

genome. The protein numbers were noted, and, as these two genes flank the mating type genes, the protein numbers between them were also noted. The coordinates for all four genes, apn2, sla2, and both MAT genes, were extracted from the general feature format (GFF) files. These coordinates were then used to calculate the mean coverage of each section of the genome from the BAM files previously generated for variant calling. Based on the relative coverage of apn2 and sla2 compared to the MAT genes, it was evident which mating type each individual was. Specifically, if the MAT gene coverage was equal to the flanking genes, it was clearly the same mating type as the individual used to generate the reference genome. If, instead, there was no or very low coverage of the MAT genes and coverage of the flanking genes on par with the mean overall coverage across the genome, then it was ascribable to the opposite mating type. Thus, by using the ratio of the coverage of the MAT genes to the flanking genes, we assigned one or the other mating type to every individual in the dataset with high enough coverage to do so. We then ran an ANOVA to determine if the mean proportion of reference mating type differed among the species.

Latent Factor Mixed Models

Filtered coding sequences for the mycobionts and POG sequences were used to test for GEAs with five environmental variables - light availability, stand maturity, landscape structure, mean annual precipitation (MAP), climate moisture deficit (CMD), and degree-days below 18°C (heating degree-days, DD_18). We chose climate variables with the greatest variability among sites for all study species that represent biologically meaningful aspects of climate (temperature, precipitation, evapotranspiration). To account for clonal reproduction, we used datasets representing unique multilocus genotypes for each species, except for the Lepraria lanata and Usnea strigosa photobionts, with too few variants being retained after filtering for depth and MAC before modeling (*97*). Variant call files for each genomic dataset were converted to genlight objects, and the gl.impute and gl.filter functions from the package DartR v2 (*98*) were used to impute missing genotypes based on the population-level average allele frequency for that locus and remove monomorphic loci. The distribution of minor allele counts (MAC) was visualized for each dataset, and loci with disproportionately high frequency MACs were removed from the dataset. Principal coordinate analyses (PCoA) of bitwise distances for coding and noncoding datasets were used to infer population structure and select the range of latent factors (K) used in the latent factor mixed models (LFMM). Univariate LFMMs were run for each environmental variable using the lfmm2 function in the R package 'LEA' v3.2.0 (*99*), and all models were evaluated using the function lfmm2.test (*100*). The q-value function from the R package 'qvalue' v2.4.2 (*101*) was used to estimate q-values, allowing for the identification of loci with a false discovery rate (FDR) of less than 5% (0.05).

Functional genetics analyses

Functional annotations and gene ontology terms were mapped to annotated mycobiont genomes with eggNOG-mapper (*102*). Outlier loci were mapped to their respective genes to create a list of genes that included loci under selection for each species and environmental factor. Gene ontology enrichment analysis were conducted by first using the eggnogmapper outcome to create a gene ontology database for each species using AnnotationForge v3.21 (*103*). We then used the lists of genes that were associated with outlier loci to run GO enrichment analyses with q-value cutoffs of 0.1 using the enrichGO function from the R package 'clusterProfiler' v4.16.0 (*104*). Network plots of results were created with enrichGO. To investigate finer-scale function and structure of top outliers, the loci with the lowest p-values were identified for each environmental variable, and the amino acid sequence of each gene from which the loci were recovered was searched against the entire NCBI database using blastp (*105*). Functions or structures were only assigned when more than four of the resulting hits shared a structure or function, and the structure or function was in consensus across the majority of the entries. If both of these criteria were not met, a status of 'unknown function' was assigned to the gene. The results were visualized with a custom plotting function. To investigate the functional genetics of the photobionts, we extracted lists of the specific genes where top outlier loci were located. We then created dotplots using a custom plotting function.

Mapping, imaging, and figure preparation

The population sampling map (Fig. 1B) and maps of sampling locations for DAPC (Fig. 3) were generated using QGIS v3.42.0 (*106*). Species distribution maps (Fig. 1A) were generated using ArcGIS Pro v3.4.0 (*107*). Species distributions were drawn and georeferenced according to established protocols (*74*). All figures were prepared using Inkscape v1.3.

### Supplementary Text

Pairwise comparisons of genetic diversity and population differentiation

Pairwise comparisons between common and rare congeneric fungi revealed that the common fungi *Lepraria finkii* and *Usnea strigosa* have significantly higher genetic diversity, measured as allelic richness, than their rare congeners *Lepraria lanata* and *Usnea subfusca* (Table S5). Congruent analyses comparing the algal partners of common and rare congeneric fungi found that the algal partners of rare fungi all have significantly lower allelic richness than the algal partners of the common fungi studied. Analyses of population differentiation, quantified using population-specific $F_{ST}$, demonstrated that fungi in the genus *Lepraria* showed no significant difference in population differentiation, nor did algal partners of common and rare fungi in the genera *Punctelia* and *Unsea* (Table S5). Comparisons between symbionts revealed no significant difference in the population differentiation of the rare fungi *L. lanata* and *Punctelia appalachensis* and their respective algal partners (Table S6). In contrast to comparisons among the remaining symbiont pairs, the common fungi, *Punctelia rudecta,* was found to have significantly lower population differentiation than its algal partner. All algal partners exhibited significantly higher genetic diversity than their fungal partners, except for the algal partners of the common fungi *U. strigosa* and the rare fungi *P. appalachensis* (Table S6).

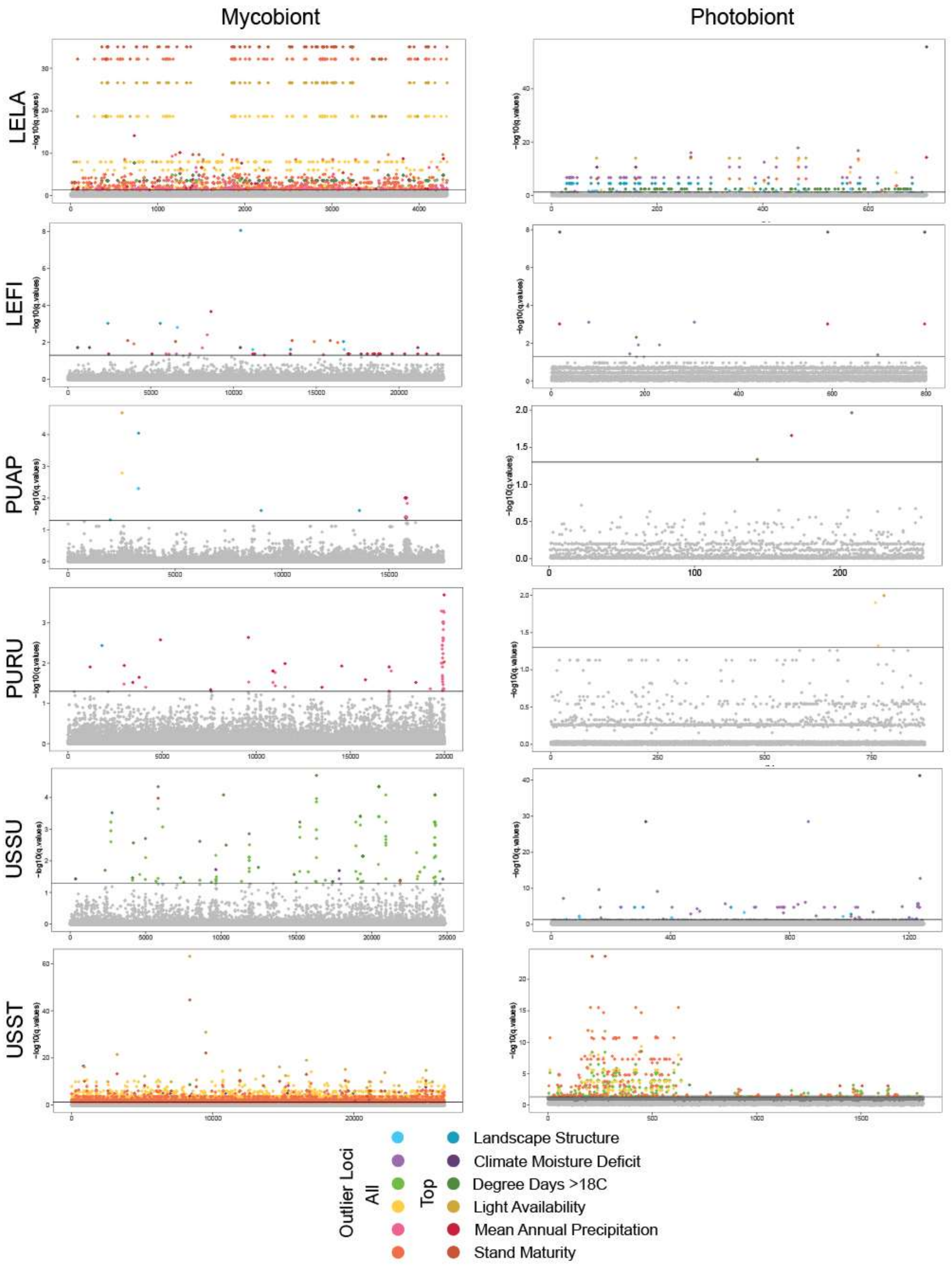

**Fig. S1**

Manhattan plots displaying outlier loci identified using latent factor mixed models (LFMM).

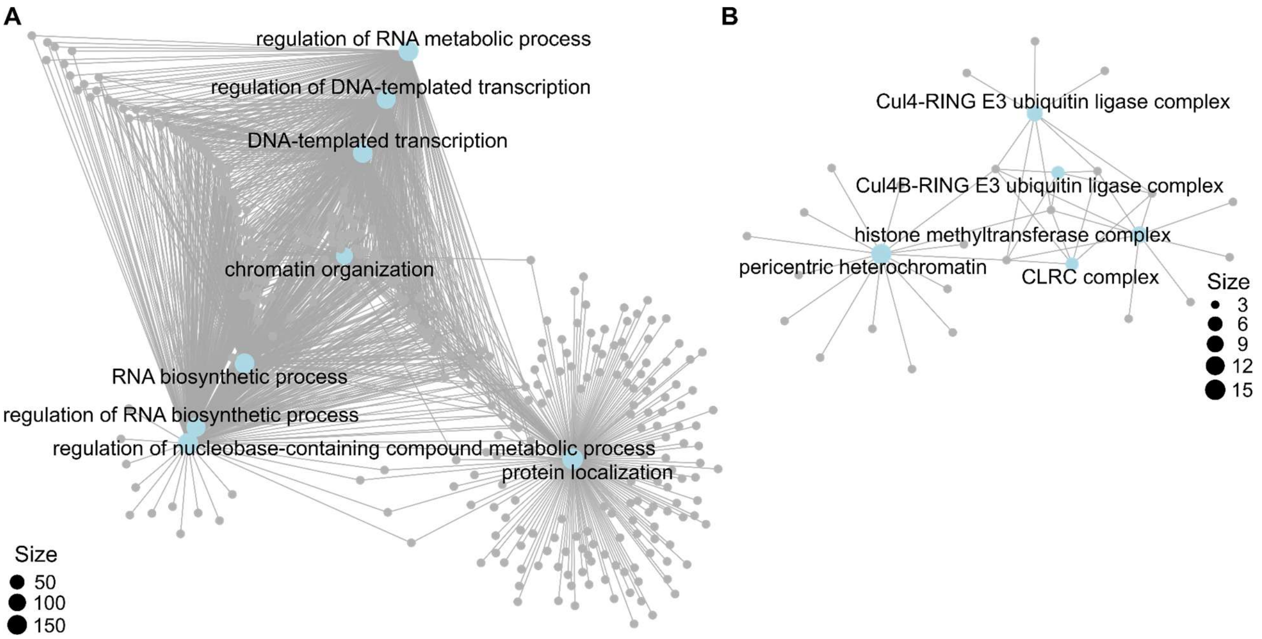

**Fig. S2**

Functional genetics results. (**A**, **B**) Gene ontology enrichment results for *Usnea strigosa* in response to light availability (left) and stand maturity (right) are shown as network plots with GO terms labeled and highlighted blue, and each gene represented by a grey dot.

| Species | Subset | Mantel r | pval1 | pval2 | pval3 | lower limit (2.55) | upper limit (97.5%) |
|---|---|---|---|---|---|---|---|
| *Lepraria finkii* | Mycobiont | 0.03731257 | 0.2165 | 0.7836 | 0.4885 | -0.02749189 | 0.11218596 |
| *Lepraria lanata* | Mycobiont | 0.9554453207 | 0.0308 | 0.987 | **0.0308** | 0.9397223504 | 0.9899320673 |
| *Punctelia appalachensis* | Mycobiont | 0.18686522 | 0.0173 | 0.9828 | **0.0173** | 0.06578272 | 0.35737683 |
| *Punctelia rudecta* | Mycobiont | 0.089820918 | 0.0801 | 0.92 | 0.0906 | -0.004895083 | 0.164970715 |
| *Usnea strigosa* | Mycobiont | 0.5056344 | 0.0809 | 0.9213 | 0.0809 | 0.2705512 | 0.6198059 |
| *Usnea subfusca* | Mycobiont | 0.8907918 | 0.0311 | 0.9695 | **0.0311** | 0.8253625 | 0.996041 |
| *Lepraria finkii* | Photobiont | 0.01705553 | 0.3472 | 0.6529 | 0.7824 | -0.02968394 | 0.08915584 |
| *Lepraria lanata* | Photobiont | 0.55688644 | 0.0954 | 0.9226 | 0.0954 | 0.07703307 | 0.9159253 |
| *Punctelia appalachensis* | Photobiont | 0.2213877 | 0.0108 | 0.9893 | **0.0108** | 0.1117613 | 0.37045 |
| *Punctelia rudecta* | Photobiont | 0.2769231 | 0.0008 | 0.9993 | **0.0008** | 0.2012101 | 0.3526072 |
| *Usnea strigosa* | Photobiont | 0.3349963 | 0.1211 | 0.8797 | 0.2227 | -0.1861635 | 0.5350934 |
| *Usnea subfusca* | Photobiont | 0.8490811 | 0.0055 | 0.995 | **0.0055** | 0.7603789 | 0.9367431 |

**Table S1.**
Results from Mantel's tests for isolation-by-distance.

| Site Code | Landscape | Region | Lat | Lon | RC | LT | CC | US | WED | REC | LIT | CWD | PS | NH | D2C | SUM | Light Availability | Stand Maturity | Landscape Structure |
|---|---|---|---|---|---|---|---|---|---|---|---|---|---|---|---|---|---|---|---|
| ALM | S. Appalachians | South | 35.65878 | -83.4379 | 2 | 4 | 3 | 25 | 15 | 10 | 5 | 5 | 10 | 5.2 | 5 | 87.2 | 5 | 59 | 20.2 |
| ASG | S. Appalachians | South | 36.10455 | -82.1293 | 1 | 3 | 2 | 20 | 15 | 10 | 5 | 3 | 8 | 3.2 | 4 | 73.2 | 3 | 51 | 15.2 |
| BEK | S. Appalachians | South | 35.8014 | -82.951 | 1 | 0 | 5 | 20 | 11 | 10 | 5 | 3 | 8 | 3.2 | 5 | 70.2 | 6 | 44 | 16.2 |
| BFF | S. Appalachians | South | 35.84115 | -82.9058 | 4 | 4 | 5 | 20 | 15 | 10 | 5 | 3 | 10 | 3.2 | 5 | 80.2 | 9 | 52 | 18.2 |
| BLU | Kentucky | Central | 37.08482 | -82.7931 | 3 | 4 | 5 | 20 | 11 | 10 | 5 | 2 | 10 | 3.2 | 5 | 75.2 | 8 | 47 | 18.2 |
| BRF | Shenandoah NP | Central | 38.4477 | -78.4664 | 3 | 3 | 5 | 20 | 15 | 10 | 5 | 3 | 8 | 4 | 5 | 78 | 8 | 51 | 17 |
| BUK | S. Appalachians | South | 35.55228 | -83.1202 | 1 | 4 | 5 | 20 | 15 | 10 | 5 | 5 | 10 | 0.4 | 5 | 79.4 | 6 | 54 | 15.4 |
| CAT | S. Appalachians | South | 35.7964 | -82.2569 | 4 | 4 | 5 | 15 | 15 | 10 | 5 | 3 | 8 | 5.2 | 5 | 75.2 | 9 | 47 | 18.2 |
| COG | S. Appalachians | South | 35.1036 | -83.2075 | 2 | 3 | 3 | 20 | 15 | 10 | 5 | 2 | 8 | 2 | 5 | 73 | 5 | 50 | 15 |
| DAD | S. Appalachians | South | 36.11211 | -81.8183 | 5 | 6 | 5 | 20 | 15 | 10 | 5 | 5 | 10 | 3.2 | 5 | 84.2 | 10 | 56 | 18.2 |
| DBG | S. Appalachians | South | 35.57191 | -83.1035 | 1 | 10 | 5 | 25 | 15 | 10 | 5 | 0 | 10 | 3.2 | 5 | 88.2 | 6 | 60 | 18.2 |
| GOS | Kentucky | Central | 36.9243 | -83.2242 | 4 | 6 | 5 | 10 | 7 | 10 | 5 | 0 | 8 | -2 | 5 | 54 | 9 | 33 | 11 |
| GRI | Shenandoah NP | Central | 38.47288 | -78.5507 | 10 | 10 | 5 | 20 | 15 | 10 | 5 | 5 | 8 | 3.2 | 5 | 86.2 | 15 | 60 | 16.2 |

| | | | | | | | | | | | | | | | | | | | |
|---|---|---|---|---|---|---|---|---|---|---|---|---|---|---|---|---|---|---|---|
| HAW | Shenandoah NP | Central | 38.55397 | -78.3969 | 8 | 2 | 5 | 15 | 15 | 10 | 5 | 2 | 8 | 3.2 | 5 | 70.2 | 13 | 44 | 16.2 |
| JON | SE. Virginia | Central | 37.17052 | -81.1642 | 2 | 0 | 5 | 15 | 11 | 10 | 5 | 3 | 10 | 3.2 | 5 | 67.2 | 7 | 39 | 18.2 |
| KLE | W. Virginia | Central | 38.93557 | -79.7305 | 2 | 9 | 5 | 15 | 9 | 10 | 5 | 5 | 10 | 3.2 | 5 | 76.2 | 7 | 48 | 18.2 |
| LCN | S. Appalachians | South | 35.65878 | -83.4379 | 1 | 4 | 5 | 20 | 11 | 5 | 5 | 0 | 8 | 3.2 | 5 | 66.2 | 6 | 40 | 16.2 |
| LEM | Kentucky | Central | 37.09032 | -82.7793 | 4 | 6 | 5 | 15 | 9 | 10 | 5 | 5 | 10 | 3.2 | 5 | 73.2 | 9 | 45 | 18.2 |
| LUE | W. Virginia | Central | 38.17587 | -80.28 | 2 | 4 | 0 | 10 | 6 | 10 | 5 | 0 | 8 | 3.2 | 4 | 50.2 | 2 | 30 | 15.2 |
| MAX | S. Appalachians | South | 35.79914 | -82.9568 | 2 | 4 | 3 | 25 | 15 | 10 | 5 | 5 | 10 | 5.2 | 5 | 87.2 | 5 | 59 | 20.2 |
| NAL | S. Appalachians | South | 35.65878 | -83.4379 | 2 | 4 | 3 | 25 | 15 | 10 | 5 | 5 | 10 | 5.2 | 5 | 87.2 | 5 | 59 | 20.2 |
| NAR | S. Appalachians | South | 35.37817 | -82.8689 | 3 | 6 | 5 | 20 | 15 | 10 | 5 | 5 | 10 | 5.2 | 5 | 86.2 | 8 | 56 | 20.2 |
| NGA | W. Virginia | Central | 38.61169 | -79.8416 | 3 | 10 | 5 | 15 | 15 | 10 | 5 | 5 | 8 | 5.2 | 5 | 83.2 | 8 | 55 | 18.2 |
| POK | W.Virginia | Central | 39.03528 | -79.4891 | 3 | 4 | 3 | 10 | 7 | 10 | 5 | 3 | 8 | 5.2 | 5 | 60.2 | 6 | 34 | 18.2 |
| RBB | S. Appalachians | South | 34.96532 | -83.2999 | 4 | 4 | 5 | 15 | 15 | 10 | 5 | 1 | 10 | 5.2 | 5 | 75.2 | 9 | 45 | 20.2 |
| ROG | SE. Virginia | Central | 36.65945 | -81.5408 | 2 | 3 | 3 | 20 | 7 | 10 | 5 | 3 | 10 | 5.2 | 5 | 71.2 | 5 | 43 | 20.2 |
| RON | SE. Virginia | Central | 37.15472 | -81.1942 | 2 | 3 | 5 | 15 | 15 | 10 | 5 | 3 | 10 | 5.2 | 5 | 76.2 | 7 | 46 | 20.2 |
| RSK | W. Virginia | Central | 38.32367 | -80.1522 | 3 | 4 | 5 | 15 | 7 | 10 | 5 | 3 | 10 | 3.2 | 5 | 67.2 | 8 | 39 | 18.2 |
| SHR | S. Appalachians | South | 35.37817 | -82.8689 | 3 | 6 | 5 | 20 | 15 | 10 | 5 | 5 | 10 | 5.2 | 5 | 86.2 | 8 | 56 | 20.2 |
| SNR | Kentucky | Central | 36.92086 | -83.2305 | 8 | 3 | 3 | 10 | 7 | 10 | 5 | 3 | 8 | 2 | 5 | 56 | 11 | 33 | 15 |

| | | | | | | | | | | | | | | | | | | | |
|---|---|---|---|---|---|---|---|---|---|---|---|---|---|---|---|---|---|---|---|
| SPF | S. Appalachians | South | 36.09362 | -82.1449 | 6 | 4 | 5 | 15 | 15 | 10 | 5 | 3 | 10 | 3.2 | 5 | 75.2 | 11 | 47 | 18.2 |
| TMK | SE. Virginia | Central | 36.65502 | -81.5264 | 4 | 3 | 3 | 15 | 11 | 10 | 5 | 3 | 10 | 5.2 | 5 | 70.2 | 7 | 42 | 20.2 |
| TOM | S. Appalachians | South | 35.7892 | -82.2558 | 3 | 5 | 4 | 15 | 15 | 10 | 5 | 5 | 8 | 5.2 | 5 | 77.2 | 7 | 50 | 18.2 |
| TRI | W. Virginia | Central | 38.64418 | -79.8303 | 1 | 6 | 5 | 15 | 11 | 10 | 5 | 1 | 8 | 5.2 | 5 | 71.2 | 6 | 43 | 18.2 |
| TYS | Shenandoah NP | Central | 38.56753 | -78.3822 | 8 | 3 | 5 | 15 | 11 | 10 | 5 | 2 | 10 | 3.2 | 5 | 69.2 | 13 | 41 | 18.2 |
| WHI | S. Appalachians | South | 35.085 | -83.3053 | 10 | 10 | 5 | 20 | 15 | 10 | 5 | 4 | 10 | 4.2 | 5 | 88.2 | 15 | 59 | 19.2 |

RC = Rock Cover
LT = Large Trees
CC = Canopy Cover
US = Understory
WED = Weediness
REC = Recruitment
LIT = Leaf Litter
CWD = Loggyness/Coarse Woody Debris
PS = Patch Size
NH = Neighborhood
D2C = Distance to core
SUM = Sum Habitat Quality Score. Sum of all variables listed above.
Light Availability = RC + CC
Stand Maturity = LT + US + REC + CWD
Landscape Structure = PS + NH + D2C

**Table S2.**
Site-level environmental variables.

| Site Code | Lat | Lon | AHM | bFFP | CMD | CMI | DD_0 | DD_18 | DD1040 | DD18 | DD5 | eFFP | EMT | Eref |
|---|---|---|---|---|---|---|---|---|---|---|---|---|---|---|
| ALM | 35.65878 | -83.4379 | 9.3 | 125 | 0 | 139.37 | 209 | 3278 | 1093 | 154 | 2243 | 281 | -26.8 | 909 |
| ASG | 36.10455 | -82.1293 | 12 | 123 | 0 | 98.22 | 239 | 3366 | 1073 | 152 | 2208 | 283 | -27.3 | 831 |
| BEK | 35.8014 | -82.951 | 14.2 | 123 | 8 | 65.47 | 163 | 2958 | 1287 | 241 | 2525 | 283 | -26.4 | 1009 |
| BFF | 35.84115 | -82.9058 | 15.4 | 121 | 46 | 53.83 | 145 | 2803 | 1418 | 307 | 2700 | 285 | -25.9 | 1042 |
| BLU | 37.08482 | -82.7931 | 16.6 | 115 | 87 | 49.26 | 128 | 2537 | 1732 | 471 | 3090 | 291 | -24.7 | 1042 |
| BRF | 38.4477 | -78.4664 | 13.6 | 123 | 14 | 77.1 | 251 | 3218 | 1321 | 282 | 2486 | 284 | -28.1 | 864 |
| BUK | 35.55228 | -83.1202 | 11.4 | 126 | 0 | 98.71 | 183 | 3151 | 1126 | 167 | 2316 | 279 | -27.2 | 977 |
| CAT | 35.7964 | -82.2569 | 10 | 124 | 0 | 129.52 | 241 | 3455 | 976 | 114 | 2092 | 285 | -27 | 797 |
| COG | 35.1036 | -83.2075 | 11.1 | 113 | 0 | 120.4 | 112 | 2640 | 1436 | 299 | 2773 | 292 | -23.6 | 963 |
| DAD | 36.11211 | -81.8183 | 11.8 | 122 | 0 | 103.09 | 228 | 3289 | 1132 | 177 | 2285 | 285 | -27 | 831 |
| DBG | 35.57191 | -83.1035 | 11.6 | 129 | 0 | 92.21 | 191 | 3193 | 1107 | 161 | 2287 | 277 | -27.8 | 971 |
| GOS | 36.9243 | -83.2242 | 16.1 | 114 | 77 | 53.66 | 120 | 2474 | 1800 | 512 | 3175 | 291 | -24.4 | 1072 |
| GRI | 38.47288 | -78.5507 | 18.3 | 119 | 148 | 33.45 | 161 | 2746 | 1656 | 459 | 2948 | 287 | -26.6 | 1051 |
| HAW | 38.55397 | -78.3969 | 13.8 | 124 | 19 | 73.95 | 260 | 3260 | 1298 | 270 | 2452 | 283 | -28.4 | 864 |
| JON | 37.17052 | -81.1642 | 19 | 121 | 148 | 29.94 | 187 | 2931 | 1478 | 354 | 2719 | 285 | -26.5 | 984 |
| KLE | 38.93557 | -79.7305 | 14 | 131 | 16 | 64.99 | 309 | 3445 | 1207 | 227 | 2316 | 275 | -29.2 | 876 |
| LCN | 35.65878 | -83.4379 | 9.3 | 125 | 0 | 139.37 | 209 | 3278 | 1093 | 154 | 2243 | 281 | -26.8 | 909 |
| LEM | 37.09032 | -82.7793 | 16.6 | 115 | 87 | 49.26 | 128 | 2537 | 1732 | 471 | 3090 | 291 | -24.7 | 1042 |
| LUE | 38.17587 | -80.28 | 14 | 128 | 26 | 69.1 | 338 | 3521 | 1172 | 210 | 2273 | 279 | -29.2 | 826 |

| | | | | | | | | | | | | | | |
|---|---|---|---|---|---|---|---|---|---|---|---|---|---|---|
| MAX | 35.79914 | -82.9568 | 14.2 | 123 | 8 | 65.47 | 163 | 2958 | 1287 | 241 | 2525 | 283 | -26.4 | 1009 |
| NAL | 35.65878 | -83.4379 | 9.3 | 125 | 0 | 139.37 | 209 | 3278 | 1093 | 154 | 2243 | 281 | -26.8 | 909 |
| NAR | 35.37817 | -82.8689 | 10.1 | 115 | 0 | 139.22 | 151 | 2892 | 1305 | 248 | 2569 | 290 | -24.8 | 861 |
| NGA | 38.61169 | -79.8416 | 12.1 | 137 | 6 | 82.82 | 446 | 4013 | 883 | 102 | 1870 | 269 | -31.4 | 799 |
| POK | 39.03528 | -79.4891 | 11.4 | 138 | 0 | 89.88 | 436 | 3937 | 935 | 118 | 1941 | 269 | -31 | 824 |
| RBB | 34.96532 | -83.2999 | 10.7 | 111 | 0 | 126.27 | 88 | 2374 | 1683 | 425 | 3102 | 294 | -23 | 1051 |
| ROG | 36.65945 | -81.5408 | 11.6 | 123 | 0 | 103.21 | 276 | 3484 | 1079 | 166 | 2179 | 281 | -27.6 | 776 |
| RON | 37.15472 | -81.1942 | 18.1 | 121 | 110 | 38.28 | 207 | 3029 | 1417 | 323 | 2631 | 284 | -26.9 | 955 |
| RSK | 38.32367 | -80.1522 | 12.5 | 132 | 3 | 83.67 | 429 | 3889 | 983 | 136 | 1999 | 274 | -30.4 | 774 |
| SHR | 35.37817 | -82.8689 | 10.1 | 115 | 0 | 139.22 | 151 | 2892 | 1305 | 248 | 2569 | 290 | -24.8 | 861 |
| SNR | 36.92086 | -83.2305 | 16.1 | 114 | 75 | 53.33 | 119 | 2466 | 1804 | 514 | 3183 | 292 | -24.3 | 1063 |
| SPF | 36.09362 | -82.1449 | 12.5 | 123 | 0 | 90.13 | 219 | 3245 | 1149 | 182 | 2313 | 284 | -27 | 867 |
| TMK | 36.65502 | -81.5264 | 11.6 | 123 | 0 | 103.21 | 276 | 3484 | 1079 | 166 | 2179 | 281 | -27.6 | 776 |
| TOM | 35.7892 | -82.2558 | 10 | 124 | 0 | 129.52 | 241 | 3455 | 976 | 114 | 2092 | 285 | -27 | 797 |
| TRI | 38.64418 | -79.8303 | 11.5 | 139 | 0 | 89.31 | 486 | 4167 | 812 | 80 | 1762 | 268 | -31.8 | 782 |
| TYS | 38.56753 | -78.3822 | 13 | 125 | 2 | 82.72 | 291 | 3396 | 1219 | 232 | 2340 | 282 | -28.7 | 833 |
| WHI | 35.085 | -83.3053 | 12.8 | 112 | 16 | 91.33 | 98 | 2490 | 1571 | 368 | 2952 | 293 | -23.4 | 1016 |

| **Site Code** | **Lat** | **Lon** | **FFP** | **MAP** | **MAT** | **MC MT** | **MSP** | **MW MT** | **NF FD** | **PAS** | **RH** | **SHM** | **TD** |
|---|---|---|---|---|---|---|---|---|---|---|---|---|---|
| ALM | 35.65878 | -83.4379 | 156 | 2087 | 9.4 | -0.1 | 968 | 18.6 | 197 | 63 | 65 | 19.2 | 18.7 |
| ASG | 36.10455 | -82.1293 | 160 | 1591 | 9.1 | -1.2 | 709 | 18.6 | 204 | 64 | 68 | 26.2 | 19.8 |

| | | | | | | | | | | | | | |
|---|---|---|---|---|---|---|---|---|---|---|---|---|---|
| BEK | 35.8014 | -82.951 | 160 | 1446 | 10.5 | 0.4 | 672 | 19.9 | 205 | 27 | 62 | 29.7 | 19.5 |
| BFF | 35.84115 | -82.9058 | 164 | 1372 | 11.1 | 0.9 | 628 | 20.7 | 211 | 21 | 61 | 33 | 19.8 |
| BLU | 37.08482 | -82.7931 | 176 | 1342 | 12.3 | 1 | 575 | 22.4 | 222 | 19 | 64 | 38.9 | 21.4 |
| BRF | 38.4477 | -78.4664 | 161 | 1461 | 9.9 | -1.5 | 703 | 20.7 | 201 | 55 | 64 | 29.4 | 22.1 |
| BUK | 35.55228 | -83.1202 | 154 | 1735 | 9.8 | 0 | 685 | 18.9 | 194 | 46 | 62 | 27.6 | 18.9 |
| CAT | 35.7964 | -82.2569 | 161 | 1878 | 8.8 | -1.1 | 783 | 17.8 | 204 | 76 | 70 | 22.8 | 19 |
| COG | 35.1036 | -83.2075 | 178 | 1949 | 11.5 | 2.2 | 791 | 20.4 | 225 | 21 | 68 | 25.8 | 18.2 |
| DAD | 36.11211 | -81.8183 | 163 | 1641 | 9.4 | -0.9 | 772 | 19 | 208 | 55 | 69 | 24.7 | 20 |
| DBG | 35.57191 | -83.1035 | 148 | 1687 | 9.6 | -0.3 | 675 | 18.8 | 185 | 48 | 60 | 27.9 | 19.1 |
| GOS | 36.9243 | -83.2242 | 177 | 1407 | 12.6 | 1.3 | 603 | 22.8 | 223 | 18 | 63 | 37.8 | 21.5 |
| GRI | 38.47288 | -78.5507 | 168 | 1187 | 11.7 | 0.3 | 568 | 22.7 | 212 | 20 | 60 | 39.9 | 22.3 |
| HAW | 38.55397 | -78.3969 | 159 | 1428 | 9.7 | -1.6 | 680 | 20.5 | 199 | 56 | 64 | 30.2 | 22.1 |
| JON | 37.17052 | -81.1642 | 164 | 1097 | 10.9 | -0.1 | 505 | 21.4 | 209 | 27 | 62 | 42.3 | 21.5 |
| KLE | 38.93557 | -79.7305 | 145 | 1364 | 9.1 | -2.4 | 656 | 19.9 | 177 | 81 | 60 | 30.4 | 22.4 |
| LCN | 35.65878 | -83.4379 | 156 | 2087 | 9.4 | -0.1 | 968 | 18.6 | 197 | 63 | 65 | 19.2 | 18.7 |
| LEM | 37.09032 | -82.7793 | 176 | 1342 | 12.3 | 1 | 575 | 22.4 | 222 | 19 | 64 | 38.9 | 21.4 |
| LUE | 38.17587 | -80.28 | 151 | 1349 | 8.9 | -3.1 | 610 | 19.6 | 188 | 95 | 64 | 32.1 | 22.7 |
| MAX | 35.79914 | -82.9568 | 160 | 1446 | 10.5 | 0.4 | 672 | 19.9 | 205 | 27 | 62 | 29.7 | 19.5 |
| NAL | 35.65878 | -83.4379 | 156 | 2087 | 9.4 | -0.1 | 968 | 18.6 | 197 | 63 | 65 | 19.2 | 18.7 |
| NAR | 35.37817 | -82.8689 | 176 | 2042 | 10.7 | 0.7 | 830 | 20.1 | 222 | 38 | 71 | 24.3 | 19.5 |
| NGA | 38.61169 | -79.8416 | 132 | 1424 | 7.2 | -4 | 648 | 17.8 | 161 | 169 | 61 | 27.5 | 21.8 |
| POK | 39.03528 | -79.4891 | 132 | 1537 | 7.5 | -3.9 | 701 | 18.3 | 162 | 178 | 60 | 26.1 | 22.2 |

| | | | | | | | | | | | | | |
|---|---|---|---|---|---|---|---|---|---|---|---|---|---|
| RBB | 34.96532 | -83.2999 | 183 | 2118 | 12.6 | 3 | 881 | 21.8 | 230 | 15 | 65 | 24.8 | 18.9 |
| ROG | 36.65945 | -81.5408 | 158 | 1624 | 8.9 | -1.6 | 733 | 19 | 199 | 83 | 69 | 26 | 20.7 |
| RON | 37.15472 | -81.1942 | 163 | 1134 | 10.5 | -0.5 | 523 | 21 | 207 | 34 | 63 | 40.2 | 21.5 |
| RSK | 38.32367 | -80.1522 | 142 | 1417 | 7.7 | -4 | 619 | 18.4 | 172 | 163 | 65 | 29.8 | 22.4 |
| SHR | 35.37817 | -82.8689 | 176 | 2042 | 10.7 | 0.7 | 830 | 20.1 | 222 | 38 | 71 | 24.3 | 19.5 |
| SNR | 36.92086 | -83.2305 | 178 | 1405 | 12.6 | 1.2 | 601 | 22.8 | 224 | 18 | 63 | 37.9 | 21.5 |
| SPF | 36.09362 | -82.1449 | 161 | 1559 | 9.5 | -0.8 | 694 | 19.1 | 206 | 53 | 67 | 27.5 | 19.9 |
| TMK | 36.65502 | -81.5264 | 158 | 1624 | 8.9 | -1.6 | 733 | 19 | 199 | 83 | 69 | 26 | 20.7 |
| TOM | 35.7892 | -82.2558 | 161 | 1878 | 8.8 | -1.1 | 783 | 17.8 | 204 | 76 | 70 | 22.8 | 19 |
| TRI | 38.64418 | -79.8303 | 129 | 1463 | 6.7 | -4.1 | 670 | 17.3 | 157 | 202 | 61 | 25.9 | 21.5 |
| TYS | 38.56753 | -78.3822 | 157 | 1480 | 9.3 | -2.1 | 719 | 20 | 196 | 70 | 65 | 27.9 | 22.1 |
| WHI | 35.085 | -83.3053 | 180 | 1727 | 12.1 | 2.6 | 701 | 21.2 | 227 | 15 | 66 | 30.3 | 18.6 |

MAT = mean annual temperature (°C)
MWMT = mean warmest month temperature (°C
MCMT = mean coldest month temperature (°C),
TD = temperature difference between MWMT and MCMT, or continentality (°C)
MAP = mean annual precipitation (mm)
MSP = May to September precipitation (mm)
AHM = annual heat-moisture index (MAT+10)/(MAP/1000))
SHM = summer heat-moisture index ((MWMT)/(MSP/1000))
DD_0 = degree-days below 0°C, chilling degree-days
DD5 = degree-days above 5°C, growing degree-days
DD_18 = degree-days below 18°C, heating degree-days
DD18 = degree-days above 18°C, cooling degree-days
NFFD = the number of frost-free days
FFP = frost-free period
bFFP = the day of the year on which FFP begins

eFFP = the day of the year on which FFP ends
PAS = precipitation as snow (mm). For individual years, it covers the period between August in the previous year and July in the current year.
EMT = extreme minimum temperature over 30 years
EXT = extreme maximum temperature over 30 years
Eref = Hargreaves reference evaporation (mm)
CMD = Hargreaves climatic moisture deficit (mm)
RH = mean annual relative humidity (%)
CMI = Hogg's climate moisture index (mm)
DD1040 = degree-days above 10°C and below 40°C

**Table S3.**
Variables from the ClimateNA dataset.

| Species | Subset | Mantel r | pval1 | pval2 | pval3 | lower limit (2.55) | upper limit (97.5%) |
|---|---|---|---|---|---|---|---|
| *Lepraria finkii* | Mycobiont | 0.3234324 | **0.0001** | 1 | **0.0001** | 0.2640937 | 0.3851078 |
| *Lepraria lanata* | Mycobiont | 0.5237657 | 0.0495 | 0.9506 | 0.1542 | 0.1763143 | 0.8859395 |
| *Punctelia appalachensis* | Mycobiont | -0.1574004 | 0.0081 | 0.992 | **0.0124** | 0.134744 | 0.2381209 |
| *Punctelia rudecta* | Mycobiont | 0.096987903 | 0.1404 | 0.8597 | 0.2968 | -0.008782784 | 0.291675962 |
| *Usnea strigosa* | Mycobiont | 0.003497767 | 0.3713 | 0.6288 | 0.9282 | -0.624300429 | 0.189020439 |
| *Usnea subfusca* | Mycobiont | -0.23031372 | 0.8436 | 0.1565 | 0.325 | -0.41019988 | -0.01215158 |
| *Lepraria finkii* | Photobiont | 0.04371511 | 0.3305 | 0.6696 | 0.6601 | -0.00483649 | 0.08786642 |
| *Lepraria lanata* | Photobiont | 0.1572747 | 0.3479 | 0.6522 | 0.6827 | -0.1827531 | 0.5728232 |
| *Punctelia appalachensis* | Photobiont | 0.1006368 | 0.2017 | 0.7984 | 0.409 | -0.1166173 | 0.2562743 |
| *Punctelia rudecta* | Photobiont | -0.09647351 | 0.8755 | 0.1246 | 0.2571 | -0.15336018 | -0.03878987 |
| *Usnea strigosa* | Photobiont | 0.002911529 | 0.3764 | 0.6237 | 0.9902 | -0.367312931 | 0.219943155 |
| *Usnea subfusca* | Photobiont | -0.35685137 | 0.9713 | 0.0288 | 0.1175 | -0.56045361 | -0.06288413 |

**Table S4.**
Results from Mantel's test of isolation-by-environment.

| | MYCOBIONTS | | | | | | PHOTOBIONTS | | | | | |
|---|---|---|---|---|---|---|---|---|---|---|---|---|
| | LEFI | LELA | PURU | PUAP | USST | USSU | LEFI | LELA | PURU | PUAP | USST | USSU |
| **Sites** | 26 | 5 | 27 | 19 | 6 | 7 | 26 | 5 | 27 | 19 | 6 | 7 |
| **Indivs.** | 227 | 81 | 291 | 181 | 45 | 76 | 237 | 81 | 289 | 184 | 45 | 78 |
| **Ave. *Ar*** | 1.19**** | 1*** | 1.06*** | 1.08**** | 1.28** | 1.2** | 1.35** | 1.21** | 1.16*** | 1.06**** | 2.27** | 1.31** |
| **Ave. $F_{ST}$** | 0.20 | 0.21 | 0.16** | 0.24** | 0.25** | 0.07** | 0*** | 0.48*** | 0.19 | 0.23 | 0.02 | 0.03 |

* indicates p<0.1; ** indicates p <0.05; *** indicates p <0.001
Gray columns denote rare species
Ave. Ar = average allelic richness using rarefied allele counts
Ave. $F_{ST}$ = average population differentiation

**Table S5.**
Summarized comparisons of genetic diversity and population differentiation between common and rare congeneric mycobionts and photobionts.

| Comparison | Subset | Metric | Shapiro Wilk: W | Shapiro Wilk: pvalue | t-test: t | t-test: df | t-test: pvalue | Wilcoxon Rank Sum: W | Wilcoxon Rank Sum: pvalue |
|---|---|---|---|---|---|---|---|---|---|
| *Lepraria finkii/ Lepraria lanata* | Mycobiont | Ar | 0.62008 | 9.33E-08 | | | | 130 | 0.0005181 |
| | | $F_{ST}$ | 0.95642 | 2.34E-01 | -0.14948 | 4.1627 | 0.8881 | | |
| *Punctelia rudecta/ Punctelia appalachensis* | Mycobiont | Ar | 0.94124 | 0.02179 | | | | 52 | 7.56E-07 |
| | | $F_{ST}$ | 0.90451 | 0.001501 | | | | 143 | 0.03721 |
| *Usnea strigosa/ Usnea subfusca* | Mycobiont | Ar | 0.82334 | 0.01308 | | | | 42 | 0.001166 |
| | | $F_{ST}$ | 0.90702 | 0.1669 | 5.3201 | 6.2452 | 0.001582 | | |
| *L. finkii/L. lanata* | Photobiont | Ar | 0.7026 | 4.09E-07 | | | | 182 | 0.01295 |
| | | $F_{ST}$ | 0.71073 | 5.49E-07 | | | | 73 | 1.01E-01 |
| *P. rudecta/P. appalachensis* | Photobiont | Ar | 0.090521 | 0.00121 | | | | 440 | 1.48E-05 |
| | | $F_{ST}$ | 0.95676 | 0.08572 | 0.054053 | 36.239 | 0.9572 | | |
| *U. strigosa/U. subfusca* | Photobiont | Ar | 0.7888 | 0.005034 | | | | 42 | 0.001166 |
| | | $F_{ST}$ | 0.97744 | 0.9647 | -0.09366685 | 10.976 | 0.8453 | | |

**Table S6.**
Statistics for t-tests and Wilcoxon Rank Sum tests comparing genetic diversity (allelic richness; *Ar*) and population differentiation (population-specific $F_{ST}$) for common and rare congeneric mycobionts and photobionts.

| Species | Subset | Site | N | MLG | eMLG | SE | H | G | lambda | E.5 | Hexp | Ia | rbarD | Corrected lambda |
|---|---|---|---|---|---|---|---|---|---|---|---|---|---|---|
| *Lepraria finkii* | Mycobiont | WHI | 9 | 8 | 8.00 | 0.00 | 2.04 | 7.36 | 0.86 | 0.95 | 0.00 | 391.92 | 0.05 | 0.97 |
| | | TYS | 11 | 10 | 9.18 | 0.39 | 2.27 | 9.31 | 0.89 | 0.96 | 0.00 | 270.20 | 0.03 | 0.98 |
| | | TRI | 7 | 6 | 6.00 | 0.00 | 1.75 | 5.44 | 0.82 | 0.94 | 0.00 | 362.29 | 0.04 | 0.95 |
| | | TOM | 8 | 8 | 8.00 | 0.00 | 2.08 | 8.00 | 0.88 | 1.00 | 0.00 | 324.35 | 0.04 | 1.00 |
| | | TMK | 5 | 5 | 5.00 | 0.00 | 1.61 | 5.00 | 0.80 | 1.00 | 0.00 | 183.57 | 0.03 | 1.00 |
| | | SNR | 10 | 10 | 10.00 | 0.00 | 2.30 | 10.00 | 0.90 | 1.00 | 0.00 | 68.62 | 0.01 | 1.00 |
| | | SHR | 5 | 5 | 5.00 | 0.00 | 1.61 | 5.00 | 0.80 | 1.00 | 0.00 | 327.41 | 0.05 | 1.00 |
| | | RSK | 10 | 7 | 7.00 | 0.00 | 1.83 | 5.56 | 0.82 | 0.87 | 0.00 | 364.04 | 0.04 | 0.91 |
| | | RON | 10 | 10 | 10.00 | 0.00 | 2.30 | 10.00 | 0.90 | 1.00 | 0.00 | 179.88 | 0.02 | 1.00 |
| | | ROG | 6 | 4 | 4.00 | 0.00 | 1.33 | 3.60 | 0.72 | 0.94 | 0.00 | 487.48 | 0.07 | 0.87 |
| | | POK | 10 | 6 | 6.00 | 0.00 | 1.50 | 3.33 | 0.70 | 0.67 | 0.00 | 675.84 | 0.08 | 0.78 |
| | | NGA | 12 | 11 | 9.32 | 0.47 | 2.37 | 10.29 | 0.90 | 0.96 | 0.00 | 78.98 | 0.01 | 0.98 |
| | | LUE | 5 | 5 | 5.00 | 0.00 | 1.61 | 5.00 | 0.80 | 1.00 | 0.00 | -0.25 | 0.00 | 1.00 |
| | | COG | 4 | 4 | 4.00 | 0.00 | 1.39 | 4.00 | 0.75 | 1.00 | 0.00 | 551.44 | 0.10 | 1.00 |
| | | CAT | 8 | 8 | 8.00 | 0.00 | 2.08 | 8.00 | 0.88 | 1.00 | 0.00 | 241.53 | 0.03 | 1.00 |
| | | GOS | 10 | 10 | 10.00 | 0.00 | 2.30 | 10.00 | 0.90 | 1.00 | 0.00 | 293.79 | 0.03 | 1.00 |
| | | BRF | 12 | 10 | 8.64 | 0.59 | 2.25 | 9.00 | 0.89 | 0.94 | 0.00 | 251.64 | 0.03 | 0.97 |
| | | HAW | 13 | 12 | 9.42 | 0.49 | 2.46 | 11.27 | 0.91 | 0.96 | 0.00 | 249.61 | 0.02 | 0.99 |
| | | BUK | 8 | 8 | 8.00 | 0.00 | 2.08 | 8.00 | 0.88 | 1.00 | 0.00 | 11.95 | 0.00 | 1.00 |

| | | | | | | | | | | | | | | |
|---|---|---|---|---|---|---|---|---|---|---|---|---|---|---|
| | | BFF | 10 | 10 | 10.00 | 0.00 | 2.30 | 10.00 | 0.90 | 1.00 | 0.00 | 57.84 | 0.01 | 1.00 |
| | | KLE | 8 | 8 | 8.00 | 0.00 | 2.08 | 8.00 | 0.88 | 1.00 | 0.00 | 123.50 | 0.01 | 1.00 |
| | | BLU | 11 | 9 | 8.27 | 0.45 | 2.10 | 7.12 | 0.86 | 0.86 | 0.00 | 322.96 | 0.04 | 0.95 |
| | | BEK | 4 | 4 | 4.00 | 0.00 | 1.39 | 4.00 | 0.75 | 1.00 | 0.00 | 29.30 | 0.00 | 1.00 |
| | | LEM | 9 | 6 | 6.00 | 0.00 | 1.74 | 5.40 | 0.81 | 0.94 | 0.00 | 351.59 | 0.04 | 0.92 |
| | | JON | 11 | 11 | 10.00 | 0.00 | 2.40 | 11.00 | 0.91 | 1.00 | 0.00 | 167.84 | 0.02 | 1.00 |
| | | GRI | 11 | 10 | 9.18 | 0.39 | 2.27 | 9.31 | 0.89 | 0.96 | 0.00 | 221.31 | 0.02 | 0.98 |
| | | **Total** | **227** | **203** | **9.94** | **0.24** | **5.26** | **177.08** | **0.99** | **0.92** | **0.00** | **50.66** | **0.00** | **1.00** |
| *Lepraria lanata* | Mycobiont | TOM | 10 | 7 | 7.00 | 0.00 | 1.75 | 4.55 | 0.78 | 0.75 | 0.00 | 582.77 | 0.18 | 0.87 |
| | | NAL | 9 | 5 | 5.00 | 0.00 | 1.52 | 4.26 | 0.77 | 0.91 | 0.00 | 446.84 | 0.09 | 0.86 |
| | | LCN | 27 | 21 | 9.14 | 0.78 | 2.97 | 17.78 | 0.94 | 0.91 | 0.00 | 85.19 | 0.01 | 0.98 |
| | | ALM | 21 | 20 | 9.79 | 0.41 | 2.98 | 19.17 | 0.95 | 0.97 | 0.00 | 61.55 | 0.01 | 1.00 |
| | | CAT | 14 | 8 | 6.49 | 0.82 | 1.91 | 5.76 | 0.83 | 0.83 | 0.00 | 311.47 | 0.08 | 0.89 |
| | | **Total** | **81** | **61** | **9.61** | **0.60** | **4.00** | **47.20** | **0.98** | **0.86** | **0.00** | **101.52** | **0.01** | **0.99** |
| *Punctelia appalachensis* | Mycobiont | WHI | 13 | 9 | 7.69 | 0.71 | 2.14 | 8.05 | 0.88 | 0.94 | 0.00 | 86.67 | 0.02 | 0.95 |
| | | TYS | 12 | 4 | 3.67 | 0.50 | 1.08 | 2.40 | 0.58 | 0.73 | 0.00 | 447.73 | 0.12 | 0.64 |
| | | TRI | 11 | 3 | 2.91 | 0.29 | 0.76 | 1.75 | 0.43 | 0.66 | 0.00 | 636.36 | 0.20 | 0.47 |
| | | SNR | 9 | 6 | 6.00 | 0.00 | 1.68 | 4.76 | 0.79 | 0.87 | 0.00 | 152.76 | 0.03 | 0.89 |

| | | | | | | | | | | | | | | |
|---|---|---|---|---|---|---|---|---|---|---|---|---|---|---|
| | | RSK | 11 | 8 | 7.55 | 0.50 | 2.02 | 7.12 | 0.86 | 0.94 | 0.00 | 102.24 | 0.02 | 0.95 |
| | | RON | 10 | 6 | 6.00 | 0.00 | 1.64 | 4.55 | 0.78 | 0.85 | 0.00 | 333.96 | 0.10 | 0.87 |
| | | DBG | 10 | 9 | 9.00 | 0.00 | 2.16 | 8.33 | 0.88 | 0.95 | 0.00 | 32.57 | 0.01 | 0.98 |
| | | HAW | 9 | 3 | 3.00 | 0.00 | 1.10 | 3.00 | 0.67 | 1.00 | 0.00 | 345.07 | 0.12 | 0.75 |
| | | DAD | 5 | 5 | 5.00 | 0.00 | 1.61 | 5.00 | 0.80 | 1.00 | 0.00 | 10.67 | 0.00 | 1.00 |
| | | NGA | 10 | 10 | 10.00 | 0.00 | 2.30 | 10.00 | 0.90 | 1.00 | 0.00 | 29.90 | 0.01 | 1.00 |
| | | BUK | 9 | 8 | 8.00 | 0.00 | 2.04 | 7.36 | 0.86 | 0.95 | 0.00 | 55.45 | 0.01 | 0.97 |
| | | BEK | 9 | 8 | 8.00 | 0.00 | 2.04 | 7.36 | 0.86 | 0.95 | 0.00 | 66.42 | 0.02 | 0.97 |
| | | KLE | 10 | 7 | 7.00 | 0.00 | 1.83 | 5.56 | 0.82 | 0.87 | 0.00 | 166.42 | 0.04 | 0.91 |
| | | BRF | 10 | 7 | 7.00 | 0.00 | 1.83 | 5.56 | 0.82 | 0.87 | 0.00 | 210.32 | 0.05 | 0.91 |
| | | GOS | 10 | 10 | 10.00 | 0.00 | 2.30 | 10.00 | 0.90 | 1.00 | 0.00 | 29.80 | 0.01 | 1.00 |
| | | BLU | 10 | 7 | 7.00 | 0.00 | 1.89 | 6.25 | 0.84 | 0.94 | 0.00 | 100.22 | 0.02 | 0.93 |
| | | BFF | 8 | 8 | 8.00 | 0.00 | 2.08 | 8.00 | 0.88 | 1.00 | 0.00 | 13.56 | 0.00 | 1.00 |
| | | LUE | 11 | 4 | 3.91 | 0.29 | 1.26 | 3.27 | 0.69 | 0.89 | 0.00 | 372.71 | 0.11 | 0.76 |
| | | LEM | 4 | 4 | 4.00 | 0.00 | 1.39 | 4.00 | 0.75 | 1.00 | 0.00 | 522.13 | 0.23 | 1.00 |
| | | **Total** | **181** | **126** | **9.72** | **0.52** | **4.67** | **83.36** | **0.99** | **0.78** | **0.00** | **17.50** | **0.00** | **0.99** |
| *Punctelia rudecta* | Mycobiont | WHI | 24 | 21 | 9.41 | 0.68 | 2.98 | 18.00 | 0.94 | 0.91 | 0.00 | 36.44 | 0.01 | 0.99 |
| | | TYS | 9 | 5 | 5.00 | 0.00 | 1.52 | 4.26 | 0.77 | 0.91 | 0.00 | 476.55 | 0.27 | 0.86 |
| | | TRI | 10 | 9 | 9.00 | 0.00 | 2.16 | 8.33 | 0.88 | 0.95 | 0.00 | 40.42 | 0.01 | 0.98 |
| | | JON | 12 | 9 | 7.82 | 0.65 | 2.09 | 7.20 | 0.86 | 0.87 | 0.00 | 251.50 | 0.07 | 0.94 |
| | | CAT | 9 | 6 | 6.00 | 0.00 | 1.58 | 3.86 | 0.74 | 0.74 | 0.00 | 241.37 | 0.08 | 0.83 |
| | | NGA | 7 | 7 | 7.00 | 0.00 | 1.95 | 7.00 | 0.86 | 1.00 | 0.00 | 71.75 | 0.02 | 1.00 |

| | | | | | | | | | | | | | | |
|---|---|---|---|---|---|---|---|---|---|---|---|---|---|---|
| | | RSK | 12 | 12 | 10.00 | 0.00 | 2.48 | 12.00 | 0.92 | 1.00 | 0.00 | 4.62 | 0.00 | 1.00 |
| | | HAW | 14 | 11 | 8.35 | 0.77 | 2.30 | 8.91 | 0.89 | 0.88 | 0.00 | 289.71 | 0.09 | 0.96 |
| | | BEK | 10 | 9 | 9.00 | 0.00 | 2.16 | 8.33 | 0.88 | 0.95 | 0.00 | 210.80 | 0.07 | 0.98 |
| | | GRI | 9 | 6 | 6.00 | 0.00 | 1.74 | 5.40 | 0.81 | 0.94 | 0.00 | 333.39 | 0.13 | 0.92 |
| | | BFF | 10 | 10 | 10.00 | 0.00 | 2.30 | 10.00 | 0.90 | 1.00 | 0.00 | 3.94 | 0.00 | 1.00 |
| | | BLU | 10 | 9 | 9.00 | 0.00 | 2.16 | 8.33 | 0.88 | 0.95 | 0.00 | 40.90 | 0.01 | 0.98 |
| | | GOS | 15 | 15 | 10.00 | 0.00 | 2.71 | 15.00 | 0.93 | 1.00 | 0.00 | 198.22 | 0.05 | 1.00 |
| | | SNR | 11 | 9 | 8.27 | 0.45 | 2.10 | 7.12 | 0.86 | 0.86 | 0.00 | 234.95 | 0.07 | 0.95 |
| | | TMK | 10 | 8 | 8.00 | 0.00 | 2.03 | 7.14 | 0.86 | 0.93 | 0.00 | 75.65 | 0.02 | 0.96 |
| | | DBG | 10 | 10 | 10.00 | 0.00 | 2.30 | 10.00 | 0.90 | 1.00 | 0.00 | 0.76 | 0.00 | 1.00 |
| | | ROG | 8 | 5 | 5.00 | 0.00 | 1.49 | 4.00 | 0.75 | 0.87 | 0.00 | 234.38 | 0.08 | 0.86 |
| | | BRF | 12 | 9 | 7.95 | 0.64 | 2.14 | 8.00 | 0.88 | 0.94 | 0.00 | 80.16 | 0.02 | 0.95 |
| | | POK | 10 | 4 | 4.00 | 0.00 | 1.22 | 2.94 | 0.66 | 0.81 | 0.00 | 388.72 | 0.16 | 0.73 |
| | | NAR | 8 | 5 | 5.00 | 0.00 | 1.49 | 4.00 | 0.75 | 0.87 | 0.00 | 225.29 | 0.08 | 0.86 |
| | | BUK | 10 | 10 | 10.00 | 0.00 | 2.30 | 10.00 | 0.90 | 1.00 | 0.00 | 5.18 | 0.00 | 1.00 |
| | | TOM | 10 | 6 | 6.00 | 0.00 | 1.61 | 4.17 | 0.76 | 0.79 | 0.00 | 225.66 | 0.08 | 0.84 |
| | | KLE | 11 | 10 | 9.18 | 0.39 | 2.27 | 9.31 | 0.89 | 0.96 | 0.00 | 80.64 | 0.02 | 0.98 |
| | | LEM | 12 | 10 | 8.64 | 0.59 | 2.25 | 9.00 | 0.89 | 0.94 | 0.00 | 304.84 | 0.11 | 0.97 |
| | | RON | 10 | 10 | 10.00 | 0.00 | 2.30 | 10.00 | 0.90 | 1.00 | 0.00 | 1.00 | 0.00 | 1.00 |
| | | SHR | 6 | 5 | 5.00 | 0.00 | 1.56 | 4.50 | 0.78 | 0.93 | 0.00 | 98.09 | 0.04 | 0.93 |
| | | LUE | 12 | 11 | 9.32 | 0.47 | 2.37 | 10.29 | 0.90 | 0.96 | 0.00 | 39.93 | 0.01 | 0.98 |
| | | **Total** | **291** | **219** | **9.87** | **0.36** | **5.26** | **156.53** | **0.99** | **0.81** | **0.00** | **46.47** | **0.00** | **1.00** |
| *Usnea strigosa* | Mycobiont | WHI | 10 | 10 | 10.00 | 0.00 | 2.30 | 10.00 | 0.90 | 1.00 | 0.00 | 0.07 | 0.00 | 1.00 |

| | | | | | | | | | | | | | | |
|---|---|---|---|---|---|---|---|---|---|---|---|---|---|---|
| | | SNR | 4 | 4 | 4.00 | 0.00 | 1.39 | 4.00 | 0.75 | 1.00 | 0.00 | 0.39 | 0.00 | 1.00 |
| | | RON | 3 | 3 | 3.00 | 0.00 | 1.10 | 3.00 | 0.67 | 1.00 | 0.00 | 0.54 | 0.00 | 1.00 |
| | | JON | 9 | 8 | 8.00 | 0.00 | 2.04 | 7.36 | 0.86 | 0.95 | 0.00 | 79.33 | 0.01 | 0.97 |
| | | GOS | 10 | 10 | 10.00 | 0.00 | 2.30 | 10.00 | 0.90 | 1.00 | 0.00 | 0.02 | 0.00 | 1.00 |
| | | MAX | 9 | 9 | 9.00 | 0.00 | 2.20 | 9.00 | 0.89 | 1.00 | 0.00 | 15.50 | 0.00 | 1.00 |
| | | **Total** | **45** | **44** | **9.95** | **0.21** | **3.78** | **43.09** | **0.98** | **0.99** | **0.00** | **33.44** | **0.00** | **1.00** |
| *Usnea subfusca* | Mycobiont | TRI | 12 | 12 | 10.00 | 0.00 | 2.48 | 12.00 | 0.92 | 1.00 | 0.00 | 0.13 | 0.00 | 1.00 |
| | | NAR | 13 | 13 | 10.00 | 0.00 | 2.56 | 13.00 | 0.92 | 1.00 | 0.00 | 0.02 | 0.00 | 1.00 |
| | | BUK | 10 | 10 | 10.00 | 0.00 | 2.30 | 10.00 | 0.90 | 1.00 | 0.00 | 0.48 | 0.00 | 1.00 |
| | | SHR | 17 | 17 | 10.00 | 0.00 | 2.83 | 17.00 | 0.94 | 1.00 | 0.00 | 0.40 | 0.00 | 1.00 |
| | | BEK | 7 | 7 | 7.00 | 0.00 | 1.95 | 7.00 | 0.86 | 1.00 | 0.00 | 0.42 | 0.00 | 1.00 |
| | | NGA | 8 | 7 | 7.00 | 0.00 | 1.91 | 6.40 | 0.84 | 0.94 | 0.00 | 125.88 | 0.02 | 0.96 |
| | | BFF | 9 | 9 | 9.00 | 0.00 | 2.20 | 9.00 | 0.89 | 1.00 | 0.00 | 0.55 | 0.00 | 1.00 |
| | | **Total** | **76** | **74** | **9.97** | **0.18** | **4.29** | **72.20** | **0.99** | **0.99** | **0.00** | **3.62** | **0.00** | **1.00** |

Pop: Population name.
N: Number of individuals observed.
MLG: Number of multilocus genotypes (MLG) observed.
eMLG: The number of expected MLG at the smallest sample size ≥ 10 based on rarefaction

SE: Standard error based on eMLG.
H: Shannon-Wiener Index of MLG diversity.
G: Stoddart and Taylor's Index of MLG diversity (Stoddart & Taylor, 1988)
lambda: Simpson's Index (Simpson, 1949).

E.5: Evenness, E5 (Pielou, 1975; Ludwig & Reynolds, 1988; Grünwald et al., 2003).
Hexp: Nei's unbiased gene diversity (Nei, 1978).
Ia: The index of association, IA (Brown, Feldman & Nevo, 1980; Smith et al., 1993).
rbarD: The standardized index of association, $\bar{r}_d$
Corrected lambda: Corrected Simpson's Index (lambda_corrected <- (N/(N - 1)) * lambda

**Table S7.**
Total and site-level indices of heterozygosity, evenness, and linkage for all mycobionts calculated using the *poppr* function from the R package poppr v2.9.6.

| **Species** | **Subset** | **Number of Loci Tested** | Latent Factors (k) | Variable | Genomic Inflation Factor (GIF) | Loci with <5% False Discovery Rate (FDR) |
|---|---|---|---|---|---|---|
| *Lepraria finkii* | Mycobiont - Coding | **1,175,991** | 6 | Light Availability | 1.577503852 | 0 |
| | | | | Stand Maturity | 1.699913701 | 8 |
| | | | | Landscape Structure | 2.253946914 | 8 |
| | | | | Mean Annual Precipitation | 1.698182236 | 27 |
| | | | | Climate Moisture Deficit | 1.733192649 | 4 |
| | | | | Heating Degree Days (DD < 18 ) | 1.775118142 | 0 |
| | | | | **Total** | | **47** |
| *Lepraria lanata* | Mycobiont - Coding | **755,604** | 2 | Light Availability | 4.467465196 | 823 |
| | | | | Stand Maturity | 12.46990769 | 570 |
| | | | | Landscape Structure | 16.22410822 | 582 |
| | | | | Mean Annual Precipitation | 0.9142275717 | 437 |
| | | | | Climate Moisture Deficit | 1.555299476 | 140 |
| | | | | Heating Degree Days (DD < 18 ) | 2.712558027 | 204 |
| | | | | | **Total** | **2756** |
| *Punctelia appalachensis* | Mycobiont - Coding | **1,204,953** | 3 | Light Availability | 1.52674672 | 2 |
| | | | | Stand Maturity | 1.685024592 | 0 |

| | | | | | | |
|---|---|---|---|---|---|---|
| | | | | Landscape Structure | 1.460216654 | 6 |
| | | | | Mean Annual Precipitation | 2.05575372 | 12 |
| | | | | Climate Moisture Deficit | 2.262605334 | 5 |
| | | | | Heating Degree Days (DD < 18 ) | 1.822793524 | 0 |
| | | | | | **Total** | **25** |
| *Punctelia rudecta* | Mycobiont - Coding | **1,610,697** | 2 | Light Availability | 1.243322367 | 0 |
| | | | | Stand Maturity | 1.408735667 | 1 |
| | | | | Landscape Structure | 1.340851459 | 1 |
| | | | | Mean Annual Precipitation | 1.03265864 | 53 |
| | | | | Climate Moisture Deficit | 1.674255461 | 0 |
| | | | | Heating Degree Days (DD < 18 ) | 1.580978104 | 0 |
| | | | | | **Total** | **55** |
| *Usnea strigosa* | Mycobiont - Coding | **2,330,686** | 2 | Light Availability | 0.343776607 | 2989 |
| | | | | Stand Maturity | 0.4895049831 | 2511 |
| | | | | Landscape Structure | 1.134757624 | 16 |
| | | | | Mean Annual Precipitation | 1.083241413 | 2 |
| | | | | Climate Moisture Deficit | 1.256321846 | 381 |
| | | | | Heating Degree Days (DD < 18 ) | 0.885941202 | 45 |
| | | | | | **Total** | **5944** |

| | | | | | | |
|---|---|---|---|---|---|---|
| *Usnea subfusca* | Mycobiont - Coding | **1,441,249** | 2 | Light Availability | 1.06086828 | 1 |
| | | | | Stand Maturity | 1.139412086 | 3 |
| | | | | Landscape Structure | 1.009521742 | 0 |
| | | | | Mean Annual Precipitation | 1.21542871 | 0 |
| | | | | Climate Moisture Deficit | 1.225793525 | 4 |
| | | | | Heating Degree Days (DD < 18 ) | 1.454714897 | 124 |
| | | | | | **Total** | **132** |
| *Lepraria finkii* | Photobiont - LFMM | **14,948** | 2 | Light Availability | 1.638110726 | 0 |
| | | | | Stand Maturity | 1.04901531 | 0 |
| | | | | Landscape Structure | 1.166443988 | 0 |
| | | | | Mean Annual Precipitation | 1.355730521 | 1 |
| | | | | Climate Moisture Deficit | 1.776109912 | 0 |
| | | | | Heating Degree Days (DD < 18 ) | 1.06822607 | 2 |
| | | | | | Total | 3 |
| *Lepraria lanata* | Photobiont - LFMM | **1,188** | 4 | Light Availability | 1.005630739 | 18 |
| | | | | Stand Maturity | 5.7678095 | 13 |
| | | | | Landscape Structure | 6.18986378 | 58 |

| | | | | | | |
|---|---|---|---|---|---|---|
| | | | | Mean Annual Precipitation | 1.475338906 | 1 |
| | | | | Climate Moisture Deficit | 0.2718481707 | 52 |
| | | | | Heating Degree Days (DD < 18 ) | 8.16228878 | 67 |
| | | | | | **Total** | **209** |
| *Punctelia appalachensis* | Photobiont - LFMM | **15,671** | 6 | Light Availability | 3.960898729 | 0 |
| | | | | Stand Maturity | 1.302345855 | 0 |
| | | | | Landscape Structure | 2.869627721 | 0 |
| | | | | Mean Annual Precipitation | 1.406719335 | 3 |
| | | | | Climate Moisture Deficit | 0.9261903861 | 9 |
| | | | | Heating Degree Days (DD < 18 ) | 2.176201618 | 1 |
| | | | | | **Total** | **13** |
| *Punctelia rudecta* | Photobiont - LFMM | **17,068** | 6 | Light Availability | 1.157705074 | 3 |
| | | | | Stand Maturity | 1.777702518 | 0 |
| | | | | Landscape Structure | 2.496738728 | 0 |
| | | | | Mean Annual Precipitation | 2.423944088 | 0 |
| | | | | Climate Moisture Deficit | 1.419913442 | 0 |
| | | | | Heating Degree Days (DD < 18 ) | 4.548958426 | 0 |
| | | | | | **Total** | **3** |

| | | | | | | |
|---|---|---|---|---|---|---|
| *Usnea strigosa* | Photobiont - LFMM | **17,421** | 3 | Light Availability | 0.1651699907 | 133 |
| | | | | Stand Maturity | 0.3453677384 | 100 |
| | | | | Landscape Structure | 1.060855856 | 201 |
| | | | | Mean Annual Precipitation | 1.964251761 | 0 |
| | | | | Climate Moisture Deficit | 1.596817763 | 0 |
| | | | | Heating Degree Days (DD < 18 ) | 0.1301311466 | 1739 |
| | | | | | **Total** | 2173 |
| *Usnea subfusca* | Photobiont - LFMM | **22,971** | 3 | Light Availability | 0.8976104703 | 0 |
| | | | | Stand Maturity | 1.731232252 | 0 |
| | | | | Landscape Structure | 0.52935582 | 11 |
| | | | | Mean Annual Precipitation | 2.406094175 | 0 |
| | | | | Climate Moisture Deficit | 0.1680688845 | 48 |
| | | | | Heating Degree Days (DD < 18 ) | 3.909459959 | 0 |
| | | | | | **Total** | 59 |

**Table S8.**
Results from latent factor mixed models (LFMM).

| Species | Var | ID | ONTOLOGY | Description | GeneRatio | BgRatio | RichFactor | Fold Enrichment | zScore | pvalue | p.adjust | qvalue | Count |
|---|---|---|---|---|---|---|---|---|---|---|---|---|---|
| USST | Light_Availability | GO:0009889 | BP | regulation of biosynthetic process | 208/845 | 668/3259 | 0.3113772455 | 1.200921234 | 3.445209405 | 0.0004006494092 | 0.08242693845 | 0.08072031254 | 208 |
| USST | Light_Availability | GO:0031326 | BP | regulation of cellular biosynthetic process | 208/845 | 667/3259 | 0.311844078 | 1.202721716 | 3.472809444 | 0.0003636047063 | 0.08014886598 | 0.07848940691 | 208 |
| USST | Light_Availability | GO:0003676 | MF | nucleic acid binding | 199/729 | 635/2806 | 0.3133858268 | 1.206256008 | 3.500060149 | 0.0003297282674 | 0.05976324847 | 0.05822307037 | 199 |
| USST | Light_Availability | GO:0010556 | BP | regulation of macromolecule biosynthetic process | 198/845 | 629/3259 | 0.3147853736 | 1.214065719 | 3.535328986 | 0.0002951123587 | 0.08014886598 | 0.07848940691 | 198 |
| USST | Light_Availability | GO:0033036 | BP | macromolecule localization | 194/845 | 616/3259 | 0.3149350649 | 1.214643049 | 3.49940224 | 0.000337160582 | 0.08014886598 | 0.07848940691 | 194 |
| USST | Light_Availability | GO:0010468 | BP | regulation of gene expression | 185/845 | 583/3259 | 0.3173241852 | 1.22385742 | 3.528563831 | 0.0003080257987 | 0.08014886598 | 0.07848940691 | 185 |
| USST | Light_Availability | GO:0070727 | BP | cellular macromolecule localization | 185/845 | 584/3259 | 0.3167808219 | 1.221761774 | 3.499181569 | 0.0003417835202 | 0.08014886598 | 0.07848940691 | 185 |
| USST | Light_Availability | GO:0008104 | BP | protein localization | 183/845 | 574/3259 | 0.318815331 | 1.229608478 | 3.585147121 | 0.0002524010702 | 0.08014886598 | 0.07848940691 | 183 |
| USST | Light_Availability | GO:0019219 | BP | regulation of nucleobase-containing compound metabolic process | 178/845 | 536/3259 | 0.3320895522 | 1.280804557 | 4.207246786 | 2.32E-05 | 0.03729559498 | 0.03652340048 | 178 |

| | | | | | | | | | | | | | |
|---|---|---|---|---|---|---|---|---|---|---|---|---|---|
| USST | Light_Availability | GO:0032774 | BP | RNA biosynthetic process | 159/845 | 479/3259 | 0.3319415449 | 1.280233722 | 3.928267146 | 7.37E-05 | 0.04294456225 | 0.04205540752 | 159 |
| USST | Light_Availability | GO:0051252 | BP | regulation of RNA metabolic process | 159/845 | 480/3259 | 0.33125 | 1.277566568 | 3.895639516 | 8.35E-05 | 0.04294456225 | 0.04205540752 | 159 |
| USST | Light_Availability | GO:0006351 | BP | DNA-templated transcription | 157/845 | 468/3259 | 0.3354700855 | 1.293842614 | 4.063432485 | 4.37E-05 | 0.03729559498 | 0.03652340048 | 157 |
| USST | Light_Availability | GO:2001141 | BP | regulation of RNA biosynthetic process | 144/845 | 424/3259 | 0.3396226415 | 1.309858211 | 4.046734091 | 4.83E-05 | 0.03729559498 | 0.03652340048 | 144 |
| USST | Light_Availability | GO:0006355 | BP | regulation of DNA-templated transcription | 143/845 | 419/3259 | 0.3412887828 | 1.316284193 | 4.102613369 | 3.89E-05 | 0.03729559498 | 0.03652340048 | 143 |
| USST | Light_Availability | GO:0000785 | CC | chromatin | 97/818 | 275/3178 | 0.3527272727 | 1.370375639 | 3.782809757 | 0.0001535849162 | 0.03551891067 | 0.03305174991 | 97 |
| USST | Light_Availability | GO:0006325 | BP | chromatin organization | 91/845 | 255/3259 | 0.3568627451 | 1.376349925 | 3.702944654 | 0.000209423118 | 0.08014886598 | 0.07848940691 | 91 |
| USST | Light_Availability | GO:0010628 | BP | positive regulation of gene expression | 81/845 | 227/3259 | 0.3568281938 | 1.376216667 | 3.47633502 | 0.0004767254456 | 0.09194842032 | 0.09004465488 | 81 |
| USST | Light_Availability | GO:0140657 | MF | ATP-dependent activity | 78/729 | 220/2806 | 0.3545454545 | 1.364683876 | 3.33753317 | 0.0007573887095 | 0.09151780239 | 0.08915926738 | 78 |
| USST | Light_Availability | GO:0016887 | MF | ATP hydrolysis activity | 73/729 | 200/2806 | 0.365 | 1.404924554 | 3.5197665 | 0.0004255826637 | 0.06170948623 | 0.06011915101 | 73 |

| | | | | | | | | | | | | | |
|---|---|---|---|---|---|---|---|---|---|---|---|---|---|
| USST | Light_Availability | GO:0004386 | MF | helicase activity | 30/729 | 65/2806 | 0.4615384615 | 1.776511554 | 3.751993104 | 0.000305099426 | 0.05976324847 | 0.05822307037 | 30 |
| USST | Light_Availability | GO:0008186 | MF | ATP-dependent activity, acting on RNA | 24/729 | 44/2806 | 0.5454545455 | 2.099513655 | 4.354387525 | 4.56E-05 | 0.03309321674 | 0.0322403607 | 24 |
| USST | Light_Availability | GO:0003724 | MF | RNA helicase activity | 23/729 | 43/2806 | 0.5348837209 | 2.058825406 | 4.144575806 | 9.89E-05 | 0.0358530998 | 0.03492911792 | 23 |
| LEFI | Stand_Maturity | GO:0071944 | CC | cell periphery | 19/46 | 522/3241 | 0.03639846743 | 2.564509412 | 4.681928936 | 3.58E-05 | 0.006796639412 | 0.006438921548 | 19 |
| USST | Light_Availability | GO:0005721 | CC | pericentric heterochromatin | 17/818 | 29/3178 | 0.5862068966 | 2.277463958 | 4.068090278 | 0.0001670168213 | 0.03551891067 | 0.03305174991 | 17 |
| USST | Stand_Maturity | GO:0005721 | CC | pericentric heterochromatin | 15/721 | 29/3178 | 0.5172413793 | 2.279879478 | 3.750208429 | 0.0005737415595 | 0.07456508795 | 0.07085342026 | 15 |
| USST | Light_Availability | GO:0035097 | CC | histone methyltransferase complex | 9/818 | Nov-78 | 0.8181818182 | 3.178706379 | 4.260889521 | 0.0001567153006 | 0.03551891067 | 0.03305174991 | 9 |
| USST | Stand_Maturity | GO:0035097 | CC | histone methyltransferase complex | 9/721 | Nov-78 | 0.8181818182 | 3.60635481 | 4.690071996 | 5.37E-05 | 0.03363920346 | 0.03196472619 | 9 |
| USST | Stand_Maturity | GO:0080008 | CC | Cul4-RING E3 ubiquitin ligase complex | 8/721 | Nov-78 | 0.7272727273 | 3.20564872 | 3.969011148 | 0.0005767189393 | 0.07456508795 | 0.07085342026 | 8 |
| USST | DD_18 | GO:0003676 | MF | nucleic acid binding | 11-Jul | 635/2806 | 0.01102362205 | 2.81202577 | 3.256065928 | 0.004100433134 | 0.09694503894 | 0.07176960662 | 7 |
| USST | Light_Availability | GO:0007535 | BP | donor selection | 6/845 | Jun-59 | 1 | 3.856804734 | 4.143328926 | 0.0002998527021 | 0.08014886598 | 0.07848940691 | 6 |
| USST | Stand_Maturity | GO:0031465 | CC | Cul4B-RING E3 ubiquitin | 5/721 | May-78 | 1 | 4.40776699 | 4.130413385 | 0.0005946179262 | 0.07456508795 | 0.07085342026 | 5 |

| | | | | | | | | | | | | | |
|---|---|---|---|---|---|---|---|---|---|---|---|---|---|
| | | | | ligase complex | | | | | | | | | |
| USST | Stand_ Maturity | GO:0043494 | CC | CLRC complex | 5/721 | May-78 | 1 | 4.40776699 | 4.130413385 | 0.0005946179262 | 0.07456508795 | 0.07085342026 | 5 |
| LEFI | Light_Availability | GO:0048584 | BP | positive regulation of response to stimulus | 16-May | 112/3321 | 0.04464285714 | 9.266183036 | 6.191166327 | 0.0001294716256 | 0.07587037263 | 0.06855181863 | 5 |
| USST | DD_18 | GO:0044877 | MF | protein-containing complex binding | 11-Apr | 191/2806 | 0.02094240838 | 5.342217991 | 3.899099597 | 0.004696560247 | 0.09694503894 | 0.07176960662 | 4 |
| USST | DD_18 | GO:2001251 | BP | negative regulation of chromosome organization | 13-Apr | 71/3259 | 0.05633802817 | 14.12351029 | 7.074455068 | 0.0001275561005 | 0.06071670386 | 0.05330502307 | 4 |
| USST | DD_18 | GO:0003690 | MF | double-stranded DNA binding | 11-Mar | 124/2806 | 0.02419354839 | 6.171554252 | 3.694659416 | 0.01071108163 | 0.09694503894 | 0.07176960662 | 3 |
| USST | DD_18 | GO:0043565 | MF | sequence-specific DNA binding | 11-Mar | 128/2806 | 0.0234375 | 5.978693182 | 3.616487188 | 0.01168817398 | 0.09694503894 | 0.07176960662 | 3 |
| USSU | Light_Availability | GO:0044283 | BP | small molecule biosynthetic process | 5-Mar | 315/3377 | 0.009523809524 | 6.432380952 | 3.898390497 | 0.006971073843 | 0.07817230055 | 0.05043374229 | 3 |
| USST | DD_18 | GO:1990837 | MF | sequence-specific double-stranded DNA binding | 11-Mar | 86/2806 | 0.03488372093 | 8.898520085 | 4.666404365 | 0.003838821184 | 0.09694503894 | 0.07176960662 | 3 |
| PURU | Light_Availability | GO:0000003 | BP | reproduction | 2-Feb | 411/3395 | 0.004866180049 | 8.260340633 | 3.811162666 | 0.01462426547 | 0.04891823018 | 0.02243618076 | 2 |

| | | | | | | | | | | | | | |
|---|---|---|---|---|---|---|---|---|---|---|---|---|---|
| PURU | Light_Availability | GO:0000902 | BP | cell morphogenesis | 2-Feb | 76/3395 | 0.02631578947 | 44.67105263 | 9.347076929 | 0.0004946787322 | 0.01535760499 | 0.007043713567 | 2 |
| USST | DD_18 | GO:0003688 | MF | DNA replication origin binding | 11-Feb | 26/2806 | 0.07692307692 | 19.62237762 | 5.983720154 | 0.004314486495 | 0.09694503894 | 0.07176960662 | 2 |
| USSU | MAP | GO:0004553 | MF | hydrolase activity, hydrolyzing O-glycosyl compounds | 4-Feb | 40/2917 | 0.05 | 36.4625 | 8.367253497 | 0.001081367903 | 0.01232875263 | 0.007209796862 | 2 |
| PURU | Light_Availability | GO:0005856 | CC | cytoskeleton | 2-Feb | 212/3301 | 0.009433962264 | 15.57075472 | 5.399105747 | 0.004106377314 | 0.02144441486 | 0.008645004873 | 2 |
| USSU | MAP | GO:0005976 | BP | polysaccharide metabolic process | 4-Feb | 81/3377 | 0.02469135802 | 20.84567901 | 6.224979344 | 0.003304788982 | 0.0737671937 | 0.06560774897 | 2 |
| USSU | Light_Availability | GO:0006520 | BP | amino acid metabolic process | 5-Feb | 191/3377 | 0.01047120419 | 7.072251309 | 3.326488093 | 0.02841291883 | 0.097253234 | 0.06274402193 | 2 |
| PURU | Light_Availability | GO:0007010 | BP | cytoskeleton organization | 2-Feb | 224/3395 | 0.008928571429 | 15.15625 | 5.321733213 | 0.004335121409 | 0.0346901797 | 0.01591053354 | 2 |
| PURU | Light_Availability | GO:0007163 | BP | establishment or maintenance of cell polarity | 2-Feb | 128/3395 | 0.015625 | 26.5234375 | 7.145762364 | 0.00141078903 | 0.02468880802 | 0.01132343827 | 2 |
| PURU | Light_Availability | GO:0008360 | BP | regulation of cell shape | 2-Feb | 56/3395 | 0.03571428571 | 60.625 | 10.92177394 | 0.0002673000869 | 0.01433700466 | 0.006575618678 | 2 |
| USSU | Light_Availability | GO:0008652 | BP | amino acid biosynthetic process | 5-Feb | 105/3377 | 0.01904761905 | 12.8647619 | 4.755426435 | 0.009006833276 | 0.07817230055 | 0.05043374229 | 2 |
| PURU | Light_Availability | GO:0009653 | BP | anatomical structure | 2-Feb | 197/3395 | 0.01015228426 | 17.23350254 | 5.69882215 | 0.003350971089 | 0.03296539071 | 0.01511946491 | 2 |

| | | | | | | | | | | | | | |
|---|---|---|---|---|---|---|---|---|---|---|---|---|---|
| | | | | morphogenesis | | | | | | | | | |
| USSU | Light_Availability | GO:0016053 | BP | organic acid biosynthetic process | 5-Feb | 176/3377 | 0.01136363636 | 7.675 | 3.501927938 | 0.02433580342 | 0.09274981838 | 0.05983859251 | 2 |
| USSU | MAP | GO:0016798 | MF | hydrolase activity, acting on glycosyl bonds | 4-Feb | 52/2917 | 0.03846153846 | 28.04807692 | 7.291707931 | 0.00182817248 | 0.01232875263 | 0.007209796862 | 2 |
| PURU | Light_Availability | GO:0019953 | BP | sexual reproduction | 2-Feb | 350/3395 | 0.005714285714 | 9.7 | 4.171945374 | 0.01060087845 | 0.04787493491 | 0.02195767692 | 2 |
| PURU | Light_Availability | GO:0022414 | BP | reproductive process | 2-Feb | 383/3395 | 0.005221932115 | 8.864229765 | 3.966496576 | 0.01269727484 | 0.04891823018 | 0.02243618076 | 2 |
| PURU | Light_Availability | GO:0022603 | BP | regulation of anatomical structure morphogenesis | 2-Feb | 80/3395 | 0.025 | 42.4375 | 9.104912152 | 0.0005484858925 | 0.01535760499 | 0.007043713567 | 2 |
| PURU | Light_Availability | GO:0022604 | BP | regulation of cell morphogenesis | 2-Feb | 60/3395 | 0.03333333333 | 56.58333333 | 10.54511404 | 0.0003072215284 | 0.01433700466 | 0.006575618678 | 2 |
| PURU | Light_Availability | GO:0032502 | BP | developmental process | 2-Feb | 315/3395 | 0.006349206349 | 10.77777778 | 4.422817999 | 0.00858397779 | 0.04787493491 | 0.02195767692 | 2 |
| USSU | Light_Availability | GO:0046394 | BP | carboxylic acid biosynthetic process | 5-Feb | 176/3377 | 0.01136363636 | 7.675 | 3.501927938 | 0.02433580342 | 0.09274981838 | 0.05983859251 | 2 |
| PURU | Light_Availability | GO:0048856 | BP | anatomical structure development | 2-Feb | 228/3395 | 0.008771929825 | 14.89035088 | 5.271516803 | 0.004491682888 | 0.0346901797 | 0.01591053354 | 2 |

| | | | | | | | | | | | | | |
|---|---|---|---|---|---|---|---|---|---|---|---|---|---|
| PURU | Light_Availability | GO:0050793 | BP | regulation of developmental process | 2-Feb | 103/3395 | 0.01941747573 | 32.96116505 | 7.996322253 | 0.0009117710106 | 0.01823542021 | 0.008363613782 | 2 |
| PURU | Light_Availability | GO:0051128 | BP | regulation of cellular component organization | 2-Feb | 420/3395 | 0.004761904762 | 8.083333333 | 3.764417874 | 0.01527255496 | 0.04972459756 | 0.02280601843 | 2 |
| PURU | Light_Availability | GO:0051641 | BP | cellular localization | 2-Feb | 794/3395 | 0.002518891688 | 4.27581864 | 2.55999379 | 0.05464394847 | 0.0921705155 | 0.04227369508 | 2 |
| PURU | Light_Availability | GO:0065008 | BP | regulation of biological quality | 2-Feb | 513/3395 | 0.003898635478 | 6.617933723 | 3.352488468 | 0.02279479598 | 0.05897398759 | 0.02704821987 | 2 |
| PURU | Light_Availability | GO:0071963 | BP | establishment or maintenance of cell polarity regulating cell shape | 2-Feb | 44/3395 | 0.04545454545 | 77.15909091 | 12.34354381 | 0.0001641986248 | 0.01433700466 | 0.006575618678 | 2 |
| PURU | Light_Availability | GO:0097435 | BP | supramolecular fiber organization | 2-Feb | 95/3395 | 0.02105263158 | 35.73684211 | 8.336315721 | 0.0007749966804 | 0.01808325588 | 0.008293824124 | 2 |
| USSU | Light_Availability | GO:1901605 | BP | alpha-amino acid metabolic process | 5-Feb | 140/3377 | 0.01428571429 | 9.648571429 | 4.024199316 | 0.01571523254 | 0.07817230055 | 0.05043374229 | 2 |
| USSU | Light_Availability | GO:1901607 | BP | alpha-amino acid biosynthetic process | 5-Feb | 96/3377 | 0.02083333333 | 14.07083333 | 5.002397706 | 0.007563016879 | 0.07817230055 | 0.05043374229 | 2 |
| PUAP | Light_Availability | GO:0000124 | CC | SAGA complex | 2-Jan | 14/3225 | 0.07142857143 | 115.1785714 | 10.66377153 | 0.008664666167 | 0.07348017774 | 0.04687730637 | 1 |
| USSU | Light_Availability | GO:0000162 | BP | tryptophan biosynthetic process | 5-Jan | May-77 | 0.2 | 135.08 | 11.55174673 | 0.007385493319 | 0.07817230055 | 0.05043374229 | 1 |

| | | | | | | | | | | | | | |
|---|---|---|---|---|---|---|---|---|---|---|---|---|---|
| PUAP | Light_Availability | GO:0000182 | MF | rDNA binding | 1-Jan | Jun-63 | 0.1666666667 | 477.1666667 | 21.82124347 | 0.002095703807 | 0.03580160671 | 0.02389837675 | 1 |
| PURU | Light_Availability | GO:0000226 | BP | microtubule cytoskeleton organization | 2-Jan | 93/3395 | 0.01075268817 | 18.25268817 | 4.095328085 | 0.05404391185 | 0.0921705155 | 0.04227369508 | 1 |
| PURU | Light_Availability | GO:0000399 | CC | cellular bud neck septin structure | 2-Jan | 14/3301 | 0.07142857143 | 117.8928571 | 10.79028585 | 0.00846557058 | 0.03315681811 | 0.01336669039 | 1 |
| PURU | Light_Availability | GO:0000749 | BP | response to pheromone triggering conjugation with cellular fusion | 2-Jan | 41/3395 | 0.0243902439 | 41.40243902 | 6.318252381 | 0.0240108378 | 0.05897398759 | 0.02704821987 | 1 |
| PURU | Light_Availability | GO:0000750 | BP | pheromone-dependent signal transduction involved in conjugation with cellular fusion | 2-Jan | 25/3395 | 0.04 | 67.9 | 8.150047852 | 0.01467546906 | 0.04891823018 | 0.02243618076 | 1 |
| PURU | Light_Availability | GO:0000775 | CC | chromosome, centromeric region | 2-Jan | 96/3301 | 0.01041666667 | 17.19270833 | 3.963892819 | 0.0573269808 | 0.08981226993 | 0.03620651419 | 1 |
| PURU | Light_Availability | GO:0000776 | CC | kinetochore | 2-Jan | 47/3301 | 0.02127659574 | 35.11702128 | 5.799520274 | 0.02827774871 | 0.06122171426 | 0.02468064629 | 1 |
| PURU | Light_Availability | GO:0000779 | CC | condensed chromosome, centromeric region | 2-Jan | 52/3301 | 0.01923076923 | 31.74038462 | 5.500686869 | 0.03126215196 | 0.06122171426 | 0.02468064629 | 1 |
| PURU | Light_Availability | GO:0000793 | CC | condensed chromosome | 2-Jan | 80/3301 | 0.0125 | 20.63125 | 4.376010897 | 0.04788998742 | 0.0776148072 | 0.03128928398 | 1 |

| | | | | | | | | | | | | | |
|---|---|---|---|---|---|---|---|---|---|---|---|---|---|
| PURU | Light_Availability | GO:0000794 | CC | condensed nuclear chromosome | 2-Jan | 65/3301 | 0.01538461538 | 25.39230769 | 4.88974399 | 0.03900011934 | 0.07050021573 | 0.0284211396 | 1 |
| PURU | Light_Availability | GO:0000922 | CC | spindle pole | 2-Jan | 13/3301 | 0.07692307692 | 126.9615385 | 11.20274726 | 0.007862080361 | 0.03315681811 | 0.01336669039 | 1 |
| PURU | Light_Availability | GO:0000940 | CC | outer kinetochore | 2-Jan | 13/3301 | 0.07692307692 | 126.9615385 | 11.20274726 | 0.007862080361 | 0.03315681811 | 0.01336669039 | 1 |
| PUAP | Light_Availability | GO:0001173 | BP | DNA-templated transcriptional start site selection | 1-Jan | Aug-23 | 0.125 | 415.375 | 20.35620299 | 0.002407463136 | 0.04975423814 | 0.02998769871 | 1 |
| PUAP | Light_Availability | GO:0001174 | BP | transcriptional start site selection at RNA polymerase II promoter | 1-Jan | Aug-23 | 0.125 | 415.375 | 20.35620299 | 0.002407463136 | 0.04975423814 | 0.02998769871 | 1 |
| PUAP | Light_Availability | GO:0001178 | BP | regulation of transcriptional start site selection at RNA polymerase II promoter | 1-Jan | Mar-23 | 0.3333333333 | 1107.666667 | 33.26659987 | 0.0009027986759 | 0.0373156786 | 0.02249077403 | 1 |
| PURU | Light_Availability | GO:0001403 | BP | invasive growth in response to glucose limitation | 2-Jan | 30/3395 | 0.03333333333 | 56.58333333 | 7.423208653 | 0.017597545 | 0.05241821916 | 0.02404143886 | 1 |
| PUAP | Light_Availability | GO:0003678 | MF | DNA helicase activity | 1-Jan | 27/2863 | 0.03703703704 | 106.037037 | 10.24875783 | 0.009430667132 | 0.06444289207 | 0.04301707815 | 1 |

| | | | | | | | | | | | | | | |
|---|---|---|---|---|---|---|---|---|---|---|---|---|---|---|
| USST | Landscape_Structure | GO:0003714 | MF | transcription corepressor activity | 5-Jan | Nov-06 | 0.09090909091 | 51.01818182 | 7.021472802 | 0.0194615467 | 0.0551606654 | 0.02903192916 | | 1 |
| USST | Landscape_Structure | GO:0004497 | MF | monooxygenase activity | 5-Jan | Dec-06 | 0.08333333333 | 46.76666667 | 6.711530764 | 0.02121564054 | 0.0551606654 | 0.02903192916 | | 1 |
| USSU | MAP | GO:0004556 | MF | alpha-amylase activity | 4-Jan | Mar-17 | 0.3333333333 | 243.0833333 | 15.5429789 | 0.004109584211 | 0.01232875263 | 0.007209796862 | | 1 |
| USSU | Stand_Maturity | GO:0004812 | MF | aminoacyl-tRNA ligase activity | 1-Jan | 29/2917 | 0.03448275862 | 100.5862069 | 9.979288897 | 0.009941720946 | 0.02982516284 | 0.01046496942 | | 1 |
| USST | Landscape_Structure | GO:0004857 | MF | enzyme inhibitor activity | 5-Jan | 25/2806 | 0.04 | 22.448 | 4.550416939 | 0.04379131839 | 0.09445441501 | 0.04971285001 | | 1 |
| LEFI | Landscape_Structure | GO:0004857 | MF | enzyme inhibitor activity | 2-Jan | 21/2855 | 0.04761904762 | 67.97619048 | 8.154949248 | 0.01465948796 | 0.05703233276 | 0.03694402122 | | 1 |
| USST | Landscape_Structure | GO:0004866 | MF | endopeptidase inhibitor activity | 5-Jan | Apr-06 | 0.25 | 140.3 | 11.77719666 | 0.007112348453 | 0.0551606654 | 0.02903192916 | | 1 |
| PUAP | Landscape_Structure | GO:0005085 | MF | guanyl-nucleotide exchange factor activity | 1-Jan | 39/2863 | 0.02564102564 | 73.41025641 | 8.509421626 | 0.01362207475 | 0.07055536151 | 0.03713440079 | | 1 |
| PURU | Light_Availability | GO:0005096 | MF | GTPase activator activity | 2-Jan | 36/2917 | 0.02777777778 | 40.51388889 | 6.247681386 | 0.02453476216 | 0.07360428649 | 0.04519561451 | | 1 |
| USST | DD_18 | GO:0005254 | MF | chloride channel activity | 11-Jan | Mar-06 | 0.3333333333 | 85.03030303 | 9.133913143 | 0.01171863108 | 0.09694503894 | 0.07176960662 | | 1 |
| LEFI | Landscape_Structure | GO:0005637 | CC | nuclear inner membrane | 3-Jan | Sep-41 | 0.1111111111 | 120.037037 | 10.88334502 | 0.008310207146 | 0.07479186431 | 0.06123310528 | | 1 |
| PUAP | Landscape_Structure | GO:0005768 | CC | endosome | 1-Jan | 123/3225 | 0.008130081301 | 26.2195122 | 5.021903244 | 0.03813953488 | 0.07204134367 | 0.03568611451 | | 1 |

| | | | | | | | | | | | | | | |
|---|---|---|---|---|---|---|---|---|---|---|---|---|---|---|
| PUAP | Landscape_Structure | GO:0005769 | CC | early endosome | 1-Jan | Nov-25 | 0.09090909091 | 293.1818182 | 17.09332671 | 0.003410852713 | 0.02899224806 | 0.01436148511 | | 1 |
| PUAP | Landscape_Structure | GO:0005794 | CC | Golgi apparatus | 1-Jan | 175/3225 | 0.005714285714 | 18.42857143 | 4.174754056 | 0.05426356589 | 0.07704233751 | 0.038163387 | | 1 |
| PUAP | Landscape_Structure | GO:0005802 | CC | trans-Golgi network | 1-Jan | 30/3225 | 0.03333333333 | 107.5 | 10.31988372 | 0.009302325581 | 0.04480620155 | 0.02219502244 | | 1 |
| PURU | Light_Availability | GO:0005815 | CC | microtubule organizing center | 2-Jan | 75/3301 | 0.01333333333 | 22.00666667 | 4.530398494 | 0.04493128804 | 0.07542037635 | 0.030404631 | | 1 |
| PURU | Light_Availability | GO:0005816 | CC | spindle pole body | 2-Jan | 68/3301 | 0.01470588235 | 24.27205882 | 4.773832975 | 0.04078139774 | 0.07098984051 | 0.02861852473 | | 1 |
| PURU | Light_Availability | GO:0005819 | CC | spindle | 2-Jan | 50/3301 | 0.02 | 33.01 | 5.614912613 | 0.03006894146 | 0.06122171426 | 0.02468064629 | | 1 |
| PURU | Light_Availability | GO:0005828 | CC | kinetochore microtubule | 2-Jan | Apr-01 | 0.25 | 412.625 | 20.27930541 | 0.002422406433 | 0.02134098483 | 0.008603308552 | | 1 |
| USST | Landscape_Structure | GO:0005847 | CC | mRNA cleavage and polyadenylation specificity factor complex | 4-Jan | 14/3178 | 0.07142857143 | 56.75 | 7.420373982 | 0.01751326134 | 0.09457161126 | 0.07374004777 | | 1 |
| USST | Landscape_Structure | GO:0005849 | CC | mRNA cleavage factor complex | 4-Jan | 17/3178 | 0.05882352941 | 46.73529412 | 6.711179132 | 0.0212359734 | 0.09556188031 | 0.07451218738 | | 1 |
| PURU | Light_Availability | GO:0005874 | CC | microtubule | 2-Jan | 30/3301 | 0.03333333333 | 55.01666667 | 7.31693077 | 0.0180964446 | 0.05315830602 | 0.02143000019 | | 1 |

| | | | | | | | | | | | | | |
|---|---|---|---|---|---|---|---|---|---|---|---|---|---|
| PURU | Light_Availability | GO:0005876 | CC | spindle microtubule | 2-Jan | 19/3301 | 0.05263157895 | 86.86842105 | 9.241060197 | 0.01148026769 | 0.03854089867 | 0.01553720439 | 1 |
| PURU | Light_Availability | GO:0005880 | CC | nuclear microtubule | 2-Jan | May-01 | 0.2 | 330.1 | 18.13009555 | 0.003027549044 | 0.02134098483 | 0.008603308552 | 1 |
| USSU | Light_Availability | GO:0006166 | BP | purine ribonucleoside salvage | 5-Jan | Apr-77 | 0.25 | 168.85 | 12.93259296 | 0.005911896961 | 0.07817230055 | 0.05043374229 | 1 |
| PUAP | Light_Availability | GO:0006275 | BP | regulation of DNA replication | 1-Jan | 50/3323 | 0.02 | 66.46 | 8.090735442 | 0.0150466446 | 0.06930054598 | 0.04176858034 | 1 |
| PUAP | Light_Availability | GO:0006352 | BP | DNA-templated transcription initiation | 1-Jan | 78/3323 | 0.01282051282 | 42.6025641 | 6.450004969 | 0.02347276557 | 0.07967401648 | 0.04802084185 | 1 |
| PUAP | Light_Availability | GO:0006353 | BP | DNA-templated transcription termination | 1-Jan | 34/3323 | 0.02941176471 | 97.73529412 | 9.835410216 | 0.01023171833 | 0.06930054598 | 0.04176858034 | 1 |
| PUAP | Light_Availability | GO:0006354 | BP | DNA-templated transcription elongation | 1-Jan | 75/3323 | 0.01333333333 | 44.30666667 | 6.580780096 | 0.0225699669 | 0.07967401648 | 0.04802084185 | 1 |
| PUAP | Light_Availability | GO:0006360 | BP | transcription by RNA polymerase I | 1-Jan | 45/3323 | 0.02222222222 | 73.84444444 | 8.53489569 | 0.01354198014 | 0.06930054598 | 0.04176858034 | 1 |
| PUAP | Light_Availability | GO:0006363 | BP | termination of RNA polymerase I transcription | 1-Jan | Mar-23 | 0.3333333333 | 1107.666667 | 33.26659987 | 0.0009027986759 | 0.0373156786 | 0.02249077403 | 1 |
| PUAP | Light_Availability | GO:0006367 | BP | transcription initiation at RNA polymerase II promoter | 1-Jan | 71/3323 | 0.01408450704 | 46.8028169 | 6.767777841 | 0.02136623533 | 0.07967401648 | 0.04802084185 | 1 |

| | | | | | | | | | | | | | |
|---|---|---|---|---|---|---|---|---|---|---|---|---|---|
| PUAP | Light_Availability | GO:0006368 | BP | transcription elongation by RNA polymerase II | 1-Jan | 66/3323 | 0.01515151515 | 50.34848485 | 7.024847674 | 0.01986157087 | 0.07967401648 | 0.04802084185 | 1 |
| PUAP | Light_Availability | GO:0006369 | BP | termination of RNA polymerase II transcription | 1-Jan | 22/3323 | 0.04545454545 | 151.0454545 | 12.24930425 | 0.006620523623 | 0.06219279767 | 0.03748462339 | 1 |
| USSU | Stand_Maturity | GO:0006418 | BP | tRNA aminoacylation for protein translation | 1-Jan | 30/3377 | 0.03333333333 | 112.5666667 | 10.56251233 | 0.008883624519 | 0.06122867361 | 0.04373476686 | 1 |
| PUAP | Light_Availability | GO:0006473 | BP | protein acetylation | 1-Jan | 51/3323 | 0.01960784314 | 65.15686275 | 8.009797922 | 0.01534757749 | 0.06930054598 | 0.04176858034 | 1 |
| USSU | Light_Availability | GO:0006553 | BP | lysine metabolic process | 5-Jan | Jul-77 | 0.1428571429 | 96.48571429 | 9.736769947 | 0.01032744348 | 0.07817230055 | 0.05043374229 | 1 |
| USSU | Light_Availability | GO:0006568 | BP | tryptophan metabolic process | 5-Jan | Dec-77 | 0.08333333333 | 56.28333333 | 7.386432749 | 0.0176518098 | 0.07817230055 | 0.05043374229 | 1 |
| USSU | Light_Availability | GO:0006576 | BP | biogenic amine metabolic process | 5-Jan | 19/3377 | 0.05263157895 | 35.54736842 | 5.814252177 | 0.02783300486 | 0.097253234 | 0.06274402193 | 1 |
| USSU | Light_Availability | GO:0006586 | BP | indolalkylamine metabolic process | 5-Jan | Dec-77 | 0.08333333333 | 56.28333333 | 7.386432749 | 0.0176518098 | 0.07817230055 | 0.05043374229 | 1 |
| PURU | Light_Availability | GO:0007015 | BP | actin filament organization | 2-Jan | 65/3395 | 0.01538461538 | 26.11538462 | 4.96310407 | 0.03793057661 | 0.07479268628 | 0.0343034125 | 1 |
| PURU | Light_Availability | GO:0007017 | BP | microtubule-based process | 2-Jan | 105/3395 | 0.009523809524 | 16.16666667 | 3.832355861 | 0.0609079698 | 0.09915250897 | 0.04547596276 | 1 |

| | | | | | | | | | | | | | |
|---|---|---|---|---|---|---|---|---|---|---|---|---|---|
| PURU | Light_Availability | GO:0007080 | BP | mitotic metaphase chromosome alignment | 2-Jan | 28/3395 | 0.03571428571 | 60.625 | 7.690681687 | 0.01642923534 | 0.05111317661 | 0.02344288551 | 1 |
| PURU | Light_Availability | GO:0007114 | BP | cell budding | 2-Jan | 40/3395 | 0.025 | 42.4375 | 6.399650638 | 0.02342867904 | 0.05897398759 | 0.02704821987 | 1 |
| PURU | Light_Availability | GO:0007117 | BP | budding cell bud growth | 2-Jan | 22/3395 | 0.04545454545 | 77.15909091 | 8.699692618 | 0.01292014063 | 0.04891823018 | 0.02243618076 | 1 |
| PURU | Light_Availability | GO:0007118 | BP | budding cell apical bud growth | 2-Jan | 15/3395 | 0.06666666667 | 113.1666667 | 10.56890709 | 0.008818299295 | 0.04787493491 | 0.02195767692 | 1 |
| PURU | Light_Availability | GO:0007119 | BP | budding cell isotropic bud growth | 2-Jan | May-95 | 0.2 | 339.5 | 18.38750539 | 0.002943772385 | 0.03170216415 | 0.01454009032 | 1 |
| PURU | Light_Availability | GO:0007124 | BP | pseudohyphal growth | 2-Jan | 37/3395 | 0.02702702703 | 45.87837838 | 6.663105029 | 0.02168116133 | 0.05837235743 | 0.02677228424 | 1 |
| PURU | Light_Availability | GO:0007127 | BP | meiosis I | 2-Jan | 96/3395 | 0.01041666667 | 17.68229167 | 4.025122996 | 0.05576226955 | 0.09293711592 | 0.04262529377 | 1 |
| PURU | Light_Availability | GO:0007186 | BP | G protein-coupled receptor signaling pathway | 2-Jan | 35/3395 | 0.02857142857 | 48.5 | 6.857044693 | 0.02051528167 | 0.05837235743 | 0.02677228424 | 1 |
| PURU | Light_Availability | GO:0007264 | BP | small GTPase-mediated signal transduction | 2-Jan | 87/3395 | 0.01149425287 | 19.51149425 | 4.246169626 | 0.05060251002 | 0.08855439253 | 0.04061517251 | 1 |
| PURU | Light_Availability | GO:0008017 | MF | microtubule binding | 2-Jan | 25/2917 | 0.04 | 58.34 | 7.540810493 | 0.01707035951 | 0.07360428649 | 0.04519561451 | 1 |

| | | | | | | | | | | | | | |
|---|---|---|---|---|---|---|---|---|---|---|---|---|---|
| PURU | Light_Availability | GO:0008047 | MF | enzyme activator activity | 2-Jan | 107/2917 | 0.009345794393 | 13.63084112 | 3.48626567 | 0.07202962812 | 0.09603950417 | 0.05897162537 | 1 |
| PURU | Light_Availability | GO:0008092 | MF | cytoskeletal protein binding | 2-Jan | 69/2917 | 0.01449275362 | 21.13768116 | 4.4335692 | 0.04675726654 | 0.09603950417 | 0.05897162537 | 1 |
| PUAP | Light_Availability | GO:0008156 | BP | negative regulation of DNA replication | 1-Jan | 25/3323 | 0.04 | 132.92 | 11.48564321 | 0.007523322299 | 0.06219279767 | 0.03748462339 | 1 |
| LEFI | Landscape_Structure | GO:0008168 | MF | methyltransferase activity | 2-Jan | 90/2855 | 0.01111111111 | 15.86111111 | 3.792416824 | 0.06206424265 | 0.1158007585 | 0.07501263711 | 1 |
| LEFI | Landscape_Structure | GO:0008170 | MF | N-methyltransferase activity | 2-Jan | 39/2855 | 0.02564102564 | 36.6025641 | 5.926364398 | 0.02713860904 | 0.0705603835 | 0.04570713101 | 1 |
| LEFI | Landscape_Structure | GO:0008173 | MF | RNA methyltransferase activity | 2-Jan | 32/2855 | 0.03125 | 44.609375 | 6.567355244 | 0.02229506748 | 0.0664090219 | 0.0430179899 | 1 |
| LEFI | Landscape_Structure | GO:0008175 | MF | tRNA methyltransferase activity | 2-Jan | 17/2855 | 0.05882352941 | 83.97058824 | 9.083088185 | 0.01187554997 | 0.05703233276 | 0.03694402122 | 1 |
| USSU | MAP | GO:0008194 | MF | UDP-glycosyltransferase activity | 4-Jan | 23/2917 | 0.04347826087 | 31.70652174 | 5.477706928 | 0.03118403824 | 0.0801875269 | 0.04689329059 | 1 |
| USSU | Light_Availability | GO:0008252 | MF | nucleotidase activity | 3-Jan | Mar-17 | 0.3333333333 | 324.1111111 | 17.96293754 | 0.003083245755 | 0.02626526032 | 0.01382382122 | 1 |
| USST | DD_18 | GO:0008308 | MF | voltage-gated monoatomic anion channel activity | 11-Jan | Mar-06 | 0.3333333333 | 85.03030303 | 9.133913143 | 0.01171863108 | 0.09694503894 | 0.07176960662 | 1 |
| PURU | Light_Availability | GO:0008361 | BP | regulation of cell size | 2-Jan | 29/3395 | 0.03448275862 | 58.53448276 | 7.553515822 | 0.01701347696 | 0.05178014726 | 0.02374878935 | 1 |

| | | | | | | | | | | | | | |
|---|---|---|---|---|---|---|---|---|---|---|---|---|---|
| PURU | Light_Availability | GO:0008608 | BP | attachment of spindle microtubules to kinetochore | 2-Jan | 37/3395 | 0.02702702703 | 45.87837838 | 6.663105029 | 0.02168116133 | 0.05837235743 | 0.02677228424 | 1 |
| LEFI | Landscape_Structure | GO:0008757 | MF | S-adenosylmethionine-dependent methyltransferase activity | 2-Jan | 75/2855 | 0.01333333333 | 19.03333333 | 4.189626815 | 0.05185827002 | 0.112359585 | 0.07278353687 | 1 |
| USSU | Light_Availability | GO:0009073 | BP | aromatic amino acid family biosynthetic process | 5-Jan | Dec-77 | 0.08333333333 | 56.28333333 | 7.386432749 | 0.0176518098 | 0.07817230055 | 0.05043374229 | 1 |
| USSU | Light_Availability | GO:0009085 | BP | lysine biosynthetic process | 5-Jan | Jul-77 | 0.1428571429 | 96.48571429 | 9.736769947 | 0.01032744348 | 0.07817230055 | 0.05043374229 | 1 |
| USSU | Light_Availability | GO:0009309 | BP | amine biosynthetic process | 5-Jan | Sep-77 | 0.1111111111 | 75.04444444 | 8.563871269 | 0.01326241392 | 0.07817230055 | 0.05043374229 | 1 |
| USSU | Stand_Maturity | GO:0009895 | BP | negative regulation of catabolic process | 1-Jan | 25/3377 | 0.04 | 135.08 | 11.57929186 | 0.007403020432 | 0.06122867361 | 0.04373476686 | 1 |
| PUAP | Landscape_Structure | GO:0010008 | CC | endosome membrane | 1-Jan | 34/3225 | 0.02941176471 | 94.85294118 | 9.687772767 | 0.01054263566 | 0.04480620155 | 0.02219502244 | 1 |
| PUAP | Light_Availability | GO:0010639 | BP | negative regulation of organelle organization | 1-Jan | 118/3323 | 0.008474576271 | 28.16101695 | 5.211623255 | 0.03551008125 | 0.09572282772 | 0.05769372469 | 1 |
| PUAP | Landscape_Structure | GO:0012506 | CC | vesicle membrane | 1-Jan | 69/3225 | 0.01449275362 | 46.73913043 | 6.76307108 | 0.02139534884 | 0.05798449612 | 0.02872297022 | 1 |

| | | | | | | | | | | | | | |
|---|---|---|---|---|---|---|---|---|---|---|---|---|---|
| USST | DD_18 | GO:0015108 | MF | chloride transmembrane transporter activity | 11-Jan | Mar-06 | 0.3333333333 | 85.03030303 | 9.133913143 | 0.01171863108 | 0.09694503894 | 0.07176960662 | 1 |
| PURU | Light_Availability | GO:0015631 | MF | tubulin binding | 2-Jan | 33/2917 | 0.0303030303 | 44.1969697 | 6.535850382 | 0.02250183753 | 0.07360428649 | 0.04519561451 | 1 |
| PURU | Light_Availability | GO:0016049 | BP | cell growth | 2-Jan | 62/3395 | 0.01612903226 | 27.37903226 | 5.088807632 | 0.03619607676 | 0.07452133451 | 0.03417895793 | 1 |
| USSU | MAP | GO:0016160 | MF | amylase activity | 4-Jan | Mar-17 | 0.3333333333 | 243.0833333 | 15.5429789 | 0.004109584211 | 0.01232875263 | 0.007209796862 | 1 |
| LEFI | Landscape_Structure | GO:0016423 | MF | tRNA (guanine) methyltransferase activity | 2-Jan | May-55 | 0.2 | 285.5 | 16.85527875 | 0.003500172431 | 0.05703233276 | 0.03694402122 | 1 |
| PUAP | Light_Availability | GO:0016567 | BP | protein ubiquitination | 1-Jan | 96/3323 | 0.01041666667 | 34.61458333 | 5.797808494 | 0.02888955763 | 0.08529297967 | 0.0514074835 | 1 |
| USST | Landscape_Structure | GO:0016604 | CC | nuclear body | 4-Jan | Dec-78 | 0.08333333333 | 66.20833333 | 8.032919407 | 0.01502556046 | 0.09457161126 | 0.07374004777 | 1 |
| USST | Landscape_Structure | GO:0016705 | MF | oxidoreductase activity, acting on paired donors, with incorporation or reduction of molecular oxygen | 5-Jan | 27/2806 | 0.03703703704 | 20.78518519 | 4.363878082 | 0.0472272075 | 0.09445441501 | 0.04971285001 | 1 |
| USST | Landscape_Structure | GO:0016709 | MF | oxidoreductase activity, acting on paired donors, with | 5-Jan | Sep-06 | 0.1111111111 | 62.35555556 | 7.787961483 | 0.01594581455 | 0.0551606654 | 0.02903192916 | 1 |

| | | | | | | | | | | | | | | |
|---|---|---|---|---|---|---|---|---|---|---|---|---|---|---|
| | | | | incorporation or reduction of molecular oxygen, NAD(P)H as one donor, and incorporation of one atom of oxygen | | | | | | | | | | |
| LEFI | Landscape_Structure | GO:0016741 | MF | transferase activity, transferring one-carbon groups | 2-Jan | 97/2855 | 0.01030927835 | 14.71649485 | 3.63850501 | 0.06680812992 | 0.1158007585 | 0.07501263711 | | 1 |
| USSU | Light_Availability | GO:0016830 | MF | carbon-carbon lyase activity | 3-Jan | 25/2917 | 0.04 | 38.89333333 | 6.104404217 | 0.02550028781 | 0.05100057563 | 0.02684240822 | | 1 |
| USSU | Light_Availability | GO:0016831 | MF | carboxy-lyase activity | 3-Jan | 13/2917 | 0.07692307692 | 74.79487179 | 8.554786318 | 0.01331494962 | 0.03550653233 | 0.01868764859 | | 1 |
| USSU | Light_Availability | GO:0016832 | MF | aldehyde-lyase activity | 3-Jan | Apr-17 | 0.25 | 243.0833333 | 15.5429789 | 0.004109584211 | 0.02626526032 | 0.01382382122 | | 1 |
| USSU | Light_Availability | GO:0016833 | MF | oxo-acid-lyase activity | 3-Jan | May-17 | 0.2 | 194.4666667 | 13.89009063 | 0.005135218007 | 0.02626526032 | 0.01382382122 | | 1 |
| USSU | Light_Availability | GO:0016853 | MF | isomerase activity | 3-Jan | 54/2917 | 0.01851851852 | 18.00617284 | 4.046713032 | 0.05453310386 | 0.0969477402 | 0.05102512642 | | 1 |
| USSU | Light_Availability | GO:0016860 | MF | intramolecular oxidoreductase activity | 3-Jan | 17/2917 | 0.05882352941 | 57.19607843 | 7.454887348 | 0.0173879481 | 0.03974388137 | 0.0209178323 | | 1 |
| USSU | Light_Availability | GO:0016861 | MF | intramolecular oxidoreductase activity, interconvertin | 3-Jan | Jul-17 | 0.1428571429 | 138.9047619 | 11.71902287 | 0.007184372584 | 0.02626526032 | 0.01382382122 | | 1 |

| | | | | | | | | | | | | | |
|---|---|---|---|---|---|---|---|---|---|---|---|---|---|
| | | | | g aldoses and ketoses | | | | | | | | | |
| USSU | Stand_Maturity | GO:0016874 | MF | ligase activity | 1-Jan | 84/2917 | 0.0119047619 | 34.72619048 | 5.80742546 | 0.02879670895 | 0.04319506342 | 0.0151561626 | 1 |
| USSU | Stand_Maturity | GO:0016875 | MF | ligase activity, forming carbon-oxygen bonds | 1-Jan | 29/2917 | 0.03448275862 | 100.5862069 | 9.979288897 | 0.009941720946 | 0.02982516284 | 0.01046496942 | 1 |
| PUAP | Light_Availability | GO:0018205 | BP | peptidyl-lysine modification | 1-Jan | 89/3323 | 0.01123595506 | 37.33707865 | 6.02802444 | 0.02678302738 | 0.08100232673 | 0.04882143631 | 1 |
| PUAP | Light_Availability | GO:0018394 | BP | peptidyl-lysine acetylation | 1-Jan | 45/3323 | 0.02222222222 | 73.84444444 | 8.53489569 | 0.01354198014 | 0.06930054598 | 0.04176858034 | 1 |
| PURU | Light_Availability | GO:0019236 | BP | response to pheromone | 2-Jan | 50/3395 | 0.02 | 33.95 | 5.697985596 | 0.02924245593 | 0.06498323541 | 0.02980434105 | 1 |
| USSU | Light_Availability | GO:0019878 | BP | lysine biosynthetic process via aminoadipic acid | 5-Jan | Jun-77 | 0.1666666667 | 112.5666667 | 10.53108565 | 0.008857341638 | 0.07817230055 | 0.05043374229 | 1 |
| PUAP | Landscape_Structure | GO:0019898 | CC | extrinsic component of membrane | 1-Jan | 77/3225 | 0.01298701299 | 41.88311688 | 6.393990685 | 0.02387596899 | 0.05798449612 | 0.02872297022 | 1 |
| PUAP | Landscape_Structure | GO:0019899 | MF | enzyme binding | 1-Jan | 152/2863 | 0.006578947368 | 18.83552632 | 4.223212795 | 0.05309116312 | 0.08494586098 | 0.04470834789 | 1 |
| PURU | Light_Availability | GO:0019954 | BP | asexual reproduction | 2-Jan | 76/3395 | 0.01315789474 | 22.33552632 | 4.566521615 | 0.04427704439 | 0.08156297651 | 0.03740858321 | 1 |
| PURU | Light_Availability | GO:0030010 | BP | establishment of cell polarity | 2-Jan | 64/3395 | 0.015625 | 26.5234375 | 5.004040575 | 0.03735258357 | 0.07470516714 | 0.03426327215 | 1 |

| | | | | | | | | | | | | | |
|---|---|---|---|---|---|---|---|---|---|---|---|---|---|
| PUAP | Landscape_Structure | GO:0030234 | MF | enzyme regulator activity | 1-Jan | 215/2863 | 0.004651162791 | 13.31627907 | 3.509455666 | 0.07509605309 | 0.08582406068 | 0.04517055825 | 1 |
| USST | Landscape_Structure | GO:0030414 | MF | peptidase inhibitor activity | 5-Jan | May-06 | 0.2 | 112.24 | 10.5168168 | 0.008884096582 | 0.0551606654 | 0.02903192916 | 1 |
| PUAP | Light_Availability | GO:0030466 | BP | silent mating-type cassette heterochromatin formation | 1-Jan | 50/3323 | 0.02 | 66.46 | 8.090735442 | 0.0150466446 | 0.06930054598 | 0.04176858034 | 1 |
| USSU | MAP | GO:0030515 | MF | snoRNA binding | 4-Jan | 27/2917 | 0.03703703704 | 27.00925926 | 5.030538274 | 0.03653198397 | 0.08219696393 | 0.04806839996 | 1 |
| LEFI | Landscape_Structure | GO:0030544 | MF | Hsp70 protein binding | 2-Jan | 20/2855 | 0.05 | 71.375 | 8.360802613 | 0.01396387164 | 0.05703233276 | 0.03694402122 | 1 |
| PUAP | Landscape_Structure | GO:0030659 | CC | cytoplasmic vesicle membrane | 1-Jan | 69/3225 | 0.01449275362 | 46.73913043 | 6.76307108 | 0.02139534884 | 0.05798449612 | 0.02872297022 | 1 |
| PUAP | Landscape_Structure | GO:0030695 | MF | GTPase regulator activity | 1-Jan | 82/2863 | 0.01219512195 | 34.91463415 | 5.823627233 | 0.02864128537 | 0.07055536151 | 0.03713440079 | 1 |
| PURU | Light_Availability | GO:0030695 | MF | GTPase regulator activity | 2-Jan | 83/2917 | 0.01204819277 | 17.57228916 | 4.011539785 | 0.05610763826 | 0.09603950417 | 0.05897162537 | 1 |
| PURU | Light_Availability | GO:0030865 | BP | cortical cytoskeleton organization | 2-Jan | 72/3395 | 0.01388888889 | 23.57638889 | 4.700397017 | 0.04197166793 | 0.07942344181 | 0.03642729286 | 1 |
| PUAP | Light_Availability | GO:0030874 | CC | nucleolar chromatin | 2-Jan | Jul-25 | 0.1428571429 | 230.3571429 | 15.13040774 | 0.0043370458 | 0.07348017774 | 0.04687730637 | 1 |
| LEFI | Landscape_Structure | GO:0031072 | MF | heat shock protein binding | 2-Jan | 22/2855 | 0.04545454545 | 64.88636364 | 7.963194412 | 0.01535485882 | 0.05703233276 | 0.03694402122 | 1 |
| PURU | Light_Availability | GO:0031106 | BP | septin ring organization | 2-Jan | 20/3395 | 0.05 | 84.875 | 9.132498918 | 0.01174905382 | 0.04891823018 | 0.02243618076 | 1 |

| | | | | | | | | | | | | | |
|---|---|---|---|---|---|---|---|---|---|---|---|---|---|
| PURU | Light_Availability | GO:0031122 | BP | cytoplasmic microtubule organization | 2-Jan | 16/3395 | 0.0625 | 106.09375 | 10.22873133 | 0.009404797342 | 0.04787493491 | 0.02195767692 | 1 |
| PURU | Light_Availability | GO:0031134 | BP | GTPase regulator activity | 2-Jan | 17/3395 | 0.05882352941 | 99.85294118 | 9.918893346 | 0.009991121819 | 0.04787493491 | 0.02195767692 | 1 |
| PURU | Light_Availability | GO:0031137 | BP | regulation of conjugation with cellular fusion | 2-Jan | 79/3395 | 0.01265822785 | 21.48734177 | 4.47271148 | 0.04600425424 | 0.08360448871 | 0.03834491587 | 1 |
| PURU | Light_Availability | GO:0031139 | BP | positive regulation of conjugation with cellular fusion | 2-Jan | 70/3395 | 0.01428571429 | 24.25 | 4.771502321 | 0.04081793827 | 0.07936821329 | 0.03640196249 | 1 |
| USSU | Stand_Maturity | GO:0031330 | BP | negative regulation of cellular catabolic process | 1-Jan | 23/3377 | 0.04347826087 | 146.826087 | 12.07584726 | 0.006810778798 | 0.06122867361 | 0.04373476686 | 1 |
| PUAP | Light_Availability | GO:0031400 | BP | negative regulation of protein modification process | 1-Jan | 45/3323 | 0.02222222222 | 73.84444444 | 8.53489569 | 0.01354198014 | 0.06930054598 | 0.04176858034 | 1 |
| PUAP | Landscape_Structure | GO:0031410 | CC | cytoplasmic vesicle | 1-Jan | 188/3225 | 0.005319148936 | 17.15425532 | 4.019235664 | 0.05829457364 | 0.07704233751 | 0.038163387 | 1 |
| PUAP | Light_Availability | GO:0031490 | MF | chromatin DNA binding | 1-Jan | Aug-63 | 0.125 | 357.875 | 18.89113549 | 0.002794271743 | 0.03580160671 | 0.02389837675 | 1 |
| PUAP | Light_Availability | GO:0031507 | BP | heterochromatin formation | 1-Jan | 84/3323 | 0.0119047619 | 39.55952381 | 6.209631536 | 0.02527836293 | 0.08037223084 | 0.04844166714 | 1 |

| | | | | | | | | | | | | | |
|---|---|---|---|---|---|---|---|---|---|---|---|---|---|
| USSU | MAP | GO:0031671 | BP | primary cell septum biogenesis | 4-Jan | Apr-77 | 0.25 | 211.0625 | 14.47416099 | 0.004731620199 | 0.08848129771 | 0.07869431488 | 1 |
| PUAP | Landscape_Structure | GO:0031901 | CC | early endosome membrane | 1-Jan | Feb-25 | 0.5 | 1612.5 | 40.14349262 | 0.0006201550388 | 0.01054263566 | 0.005222358221 | 1 |
| PUAP | Light_Availability | GO:0031934 | CC | mating-type region heterochromatin | 2-Jan | 18/3225 | 0.05555555556 | 89.58333333 | 9.386877552 | 0.01113336026 | 0.07348017774 | 0.04687730637 | 1 |
| PUAP | Landscape_Structure | GO:0031982 | CC | vesicle | 1-Jan | 190/3225 | 0.005263157895 | 16.97368421 | 3.996709173 | 0.05891472868 | 0.07704233751 | 0.038163387 | 1 |
| PURU | Light_Availability | GO:0032005 | BP | signal transduction involved in positive regulation of conjugation with cellular fusion | 2-Jan | 28/3395 | 0.03571428571 | 60.625 | 7.690681687 | 0.01642923534 | 0.05111317661 | 0.02344288551 | 1 |
| PURU | Light_Availability | GO:0032156 | CC | septin cytoskeleton | 2-Jan | 23/3301 | 0.04347826087 | 71.76086957 | 8.383655318 | 0.01388872059 | 0.04351799118 | 0.01754364706 | 1 |
| PURU | Light_Availability | GO:0032161 | CC | cleavage apparatus septin structure | 2-Jan | 16/3301 | 0.0625 | 103.15625 | 10.0841212 | 0.00967200022 | 0.0349680008 | 0.01409684243 | 1 |
| PURU | Light_Availability | GO:0032176 | CC | split septin rings | 2-Jan | Apr-01 | 0.25 | 412.625 | 20.27930541 | 0.002422406433 | 0.02134098483 | 0.008603308552 | 1 |
| PURU | Light_Availability | GO:0032177 | CC | cellular bud neck split septin rings | 2-Jan | Apr-01 | 0.25 | 412.625 | 20.27930541 | 0.002422406433 | 0.02134098483 | 0.008603308552 | 1 |

| | | | | | | | | | | | | | |
|---|---|---|---|---|---|---|---|---|---|---|---|---|---|
| PURU | Light_Availability | GO:0032185 | BP | septin cytoskeleton organization | 2-Jan | 20/3395 | 0.05 | 84.875 | 9.132498918 | 0.01174905382 | 0.04891823018 | 0.02243618076 | 1 |
| PURU | Light_Availability | GO:0032231 | BP | regulation of actin filament bundle assembly | 2-Jan | Aug-95 | 0.125 | 212.1875 | 14.51725787 | 0.004707952959 | 0.0346901797 | 0.01591053354 | 1 |
| PUAP | Light_Availability | GO:0032392 | BP | DNA geometric change | 1-Jan | 42/3323 | 0.02380952381 | 79.11904762 | 8.838498041 | 0.01263918146 | 0.06930054598 | 0.04176858034 | 1 |
| PUAP | Light_Availability | GO:0032446 | BP | protein modification by small protein conjugation | 1-Jan | 114/3323 | 0.008771929825 | 29.14912281 | 5.305574691 | 0.03430634968 | 0.09453305246 | 0.05697662755 | 1 |
| USST | Landscape_Structure | GO:0032451 | MF | demethylase activity | 5-Jan | Sep-06 | 0.1111111111 | 62.35555556 | 7.787961483 | 0.01594581455 | 0.0551606654 | 0.02903192916 | 1 |
| PUAP | Light_Availability | GO:0032508 | BP | DNA duplex unwinding | 1-Jan | 35/3323 | 0.02857142857 | 94.94285714 | 9.692412349 | 0.01053265122 | 0.06930054598 | 0.04176858034 | 1 |
| PURU | Light_Availability | GO:0032535 | BP | regulation of cellular component size | 2-Jan | 59/3395 | 0.01694915254 | 28.77118644 | 5.223798375 | 0.03446001477 | 0.07309700103 | 0.0335256922 | 1 |
| PURU | Light_Availability | GO:0032878 | BP | regulation of establishment or maintenance of cell polarity | 2-Jan | 25/3395 | 0.04 | 67.9 | 8.150047852 | 0.01467546906 | 0.04891823018 | 0.02243618076 | 1 |
| PURU | Light_Availability | GO:0032956 | BP | regulation of actin cytoskeleton organization | 2-Jan | 73/3395 | 0.01369863014 | 23.25342466 | 4.665921829 | 0.0425482724 | 0.07942344181 | 0.03642729286 | 1 |

| | | | | | | | | | | | | | | |
|---|---|---|---|---|---|---|---|---|---|---|---|---|---|---|
| PURU | Light_Availability | GO:0032970 | BP | regulation of actin filament-based process | 2-Jan | 73/3395 | 0.01369863014 | 23.25342466 | 4.665921829 | 0.0425482724 | 0.07942344181 | 0.03642729286 | | 1 |
| USSU | MAP | GO:0033201 | MF | alpha-1,4-glucan synthase activity | 4-Jan | Mar-17 | 0.3333333333 | 243.0833333 | 15.5429789 | 0.004109584211 | 0.01232875263 | 0.007209796862 | | 1 |
| USSU | Stand_Maturity | GO:0033615 | BP | mitochondrial proton-transporting ATP synthase complex assembly | 1-Jan | Jun-77 | 0.1666666667 | 562.8333333 | 23.70302372 | 0.001776724904 | 0.02901984009 | 0.02072845721 | | 1 |
| PUAP | Light_Availability | GO:0034728 | BP | nucleosome organization | 1-Jan | 68/3323 | 0.01470588235 | 48.86764706 | 6.918644886 | 0.02046343665 | 0.07967401648 | 0.04802084185 | | 1 |
| PUAP | Light_Availability | GO:0035064 | MF | methylated histone binding | 1-Jan | Oct-63 | 0.1 | 286.3 | 16.89082591 | 0.003492839679 | 0.03580160671 | 0.02389837675 | | 1 |
| PURU | Light_Availability | GO:0036267 | BP | invasive filamentous growth | 2-Jan | 36/3395 | 0.02777777778 | 47.15277778 | 6.758076138 | 0.02109830829 | 0.05837235743 | 0.02677228424 | | 1 |
| USSU | Light_Availability | GO:0042278 | BP | purine nucleoside metabolic process | 5-Jan | 17/3377 | 0.05882352941 | 39.72941176 | 6.163652717 | 0.02493274688 | 0.09274981838 | 0.05983859251 | | 1 |
| PUAP | Light_Availability | GO:0042393 | MF | histone binding | 1-Jan | 33/2863 | 0.0303030303 | 86.75757576 | 9.260538632 | 0.01152637094 | 0.06751160122 | 0.04506551044 | | 1 |
| USSU | Light_Availability | GO:0042401 | BP | biogenic amine biosynthetic process | 5-Jan | Sep-77 | 0.1111111111 | 75.04444444 | 8.563871269 | 0.01326241392 | 0.07817230055 | 0.05043374229 | | 1 |
| USSU | Light_Availability | GO:0042430 | BP | indole-containing compound | 5-Jan | Dec-77 | 0.08333333333 | 56.28333333 | 7.386432749 | 0.0176518098 | 0.07817230055 | 0.05043374229 | | 1 |

| | | | | | | | | | | | | | |
|---|---|---|---|---|---|---|---|---|---|---|---|---|---|
| | | | | metabolic process | | | | | | | | | |
| USSU | Light_Availability | GO:0042435 | BP | indole-containing compound biosynthetic process | 5-Jan | May-77 | 0.2 | 135.08 | 11.55174673 | 0.007385493319 | 0.07817230055 | 0.05043374229 | 1 |
| USSU | Light_Availability | GO:0042451 | BP | purine nucleoside biosynthetic process | 5-Jan | Aug-77 | 0.125 | 84.425 | 9.095637457 | 0.01179580038 | 0.07817230055 | 0.05043374229 | 1 |
| USSU | Light_Availability | GO:0042455 | BP | ribonucleoside biosynthetic process | 5-Jan | 20/3377 | 0.05 | 33.77 | 5.659241233 | 0.02928054357 | 0.097253234 | 0.06274402193 | 1 |
| PURU | Light_Availability | GO:0042729 | CC | DASH complex | 2-Jan | Jun-01 | 0.1666666667 | 275.0833333 | 16.5428889 | 0.003632508055 | 0.02134098483 | 0.008603308552 | 1 |
| USSU | Stand_Maturity | GO:0043038 | BP | amino acid activation | 1-Jan | 32/3377 | 0.03125 | 105.53125 | 10.22405252 | 0.009475866153 | 0.06122867361 | 0.04373476686 | 1 |
| USSU | Stand_Maturity | GO:0043039 | BP | tRNA aminoacylation | 1-Jan | 32/3377 | 0.03125 | 105.53125 | 10.22405252 | 0.009475866153 | 0.06122867361 | 0.04373476686 | 1 |
| PURU | Light_Availability | GO:0043060 | BP | meiotic metaphase I homologous chromosome alignment | 2-Jan | 18/3395 | 0.05555555556 | 94.30555556 | 9.635121872 | 0.01057727272 | 0.04787493491 | 0.02195767692 | 1 |
| PURU | Light_Availability | GO:0043087 | BP | regulation of GTPase activity | 2-Jan | 48/3395 | 0.02083333333 | 35.36458333 | 5.820802073 | 0.02808108913 | 0.06444840127 | 0.02955904119 | 1 |
| USSU | Light_Availability | GO:0043101 | BP | purine-containing compound salvage | 5-Jan | Jul-77 | 0.1428571429 | 96.48571429 | 9.736769947 | 0.01032744348 | 0.07817230055 | 0.05043374229 | 1 |

| | | | | | | | | | | | | | |
|---|---|---|---|---|---|---|---|---|---|---|---|---|---|
| USSU | Light_Availability | GO:0043174 | BP | nucleoside salvage | 5-Jan | Oct-77 | 0.1 | 67.54 | 8.113414736 | 0.01472728563 | 0.07817230055 | 0.05043374229 | 1 |
| PUAP | Light_Availability | GO:0043254 | BP | regulation of protein-containing complex assembly | 1-Jan | 83/3323 | 0.01204819277 | 40.03614458 | 6.247891211 | 0.02497743003 | 0.08037223084 | 0.04844166714 | 1 |
| USST | DD_18 | GO:0043295 | MF | glutathione binding | 11-Jan | Mar-06 | 0.3333333333 | 85.03030303 | 9.133913143 | 0.01171863108 | 0.09694503894 | 0.07176960662 | 1 |
| USSU | Stand_Maturity | GO:0043461 | BP | proton-transporting ATP synthase complex assembly | 1-Jan | Jun-77 | 0.1666666667 | 562.8333333 | 23.70302372 | 0.001776724904 | 0.02901984009 | 0.02072845721 | 1 |
| USSU | Stand_Maturity | GO:0043487 | BP | regulation of RNA stability | 1-Jan | 63/3377 | 0.01587301587 | 53.6031746 | 7.252804603 | 0.01865561149 | 0.09949659461 | 0.07106899615 | 1 |
| USSU | Stand_Maturity | GO:0043488 | BP | regulation of mRNA stability | 1-Jan | 63/3377 | 0.01587301587 | 53.6031746 | 7.252804603 | 0.01865561149 | 0.09949659461 | 0.07106899615 | 1 |
| USSU | Stand_Maturity | GO:0043489 | BP | RNA stabilization | 1-Jan | Jun-77 | 0.1666666667 | 562.8333333 | 23.70302372 | 0.001776724904 | 0.02901984009 | 0.02072845721 | 1 |
| PUAP | Light_Availability | GO:0043543 | BP | protein acylation | 1-Jan | 57/3323 | 0.01754385965 | 58.29824561 | 7.569560464 | 0.01715317484 | 0.07334460967 | 0.04420600413 | 1 |
| PURU | Light_Availability | GO:0043547 | BP | positive regulation of GTPase activity | 2-Jan | 44/3395 | 0.02272727273 | 38.57954545 | 6.090734015 | 0.02575627266 | 0.06111657918 | 0.02803091226 | 1 |
| USST | Landscape_Structure | GO:0044613 | CC | nuclear pore central transport channel | 4-Jan | Mar-78 | 0.3333333333 | 264.8333333 | 16.2275729 | 0.003772394882 | 0.045247363 | 0.03528059493 | 1 |
| USST | Landscape_Structure | GO:0044614 | CC | nuclear pore cytoplasmic filaments | 4-Jan | Apr-78 | 0.25 | 198.625 | 14.03794573 | 0.005027484778 | 0.045247363 | 0.03528059493 | 1 |

| | | | | | | | | | | | | | |
|---|---|---|---|---|---|---|---|---|---|---|---|---|---|
| PURU | Light_Availability | GO:0044732 | CC | mitotic spindle pole body | 2-Jan | 50/3301 | 0.02 | 33.01 | 5.614912613 | 0.03006894146 | 0.06122171426 | 0.02468064629 | 1 |
| PURU | Light_Availability | GO:0045132 | BP | meiotic chromosome segregation | 2-Jan | 57/3395 | 0.01754385965 | 29.78070175 | 5.319546722 | 0.03330177225 | 0.07172689408 | 0.03289729729 | 1 |
| PURU | Light_Availability | GO:0045143 | BP | homologous chromosome segregation | 2-Jan | 41/3395 | 0.0243902439 | 41.40243902 | 6.318252381 | 0.0240108378 | 0.05897398759 | 0.02704821987 | 1 |
| PUAP | Light_Availability | GO:0045815 | BP | transcription initiation-coupled chromatin remodeling | 1-Jan | 19/3323 | 0.05263157895 | 174.8947368 | 13.18691536 | 0.005717724947 | 0.06219279767 | 0.03748462339 | 1 |
| USSU | Light_Availability | GO:0046128 | BP | purine ribonucleoside metabolic process | 5-Jan | 13/3377 | 0.07692307692 | 51.95384615 | 7.087010761 | 0.01911146536 | 0.08078937629 | 0.05212217825 | 1 |
| USSU | Light_Availability | GO:0046129 | BP | purine ribonucleoside biosynthetic process | 5-Jan | Aug-77 | 0.125 | 84.425 | 9.095637457 | 0.01179580038 | 0.07817230055 | 0.05043374229 | 1 |
| USSU | Light_Availability | GO:0046219 | BP | indolalkylamine biosynthetic process | 5-Jan | May-77 | 0.2 | 135.08 | 11.55174673 | 0.007385493319 | 0.07817230055 | 0.05043374229 | 1 |
| PUAP | Light_Availability | GO:0046695 | CC | SLIK (SAGA-like) complex | 2-Jan | 13/3225 | 0.07692307692 | 124.0384615 | 11.07151777 | 0.008047011753 | 0.07348017774 | 0.04687730637 | 1 |
| USSU | Light_Availability | GO:0046912 | MF | acyltransferase activity, acyl groups converted into alkyl on transfer | 3-Jan | Aug-17 | 0.125 | 121.5416667 | 10.95266837 | 0.00820789385 | 0.02626526032 | 0.01382382122 | 1 |

| | | | | | | | | | | | | | |
|---|---|---|---|---|---|---|---|---|---|---|---|---|---|
| USSU | MAP | GO:0047657 | MF | alpha-1,3-glucan synthase activity | 4-Jan | Mar-17 | 0.3333333333 | 243.0833333 | 15.5429789 | 0.004109584211 | 0.01232875263 | 0.007209796862 | 1 |
| USSU | Stand_Maturity | GO:0048255 | BP | mRNA stabilization | 1-Jan | Jun-77 | 0.1666666667 | 562.8333333 | 23.70302372 | 0.001776724904 | 0.02901984009 | 0.02072845721 | 1 |
| PURU | Light_Availability | GO:0050000 | BP | chromosome localization | 2-Jan | 56/3395 | 0.01785714286 | 30.3125 | 5.369299765 | 0.03272239063 | 0.07158022951 | 0.03283003008 | 1 |
| PURU | Light_Availability | GO:0051010 | MF | microtubule plus-end binding | 2-Jan | Jul-17 | 0.1428571429 | 208.3571429 | 14.38503768 | 0.004794513784 | 0.05753416541 | 0.0353279963 | 1 |
| PURU | Light_Availability | GO:0051017 | BP | actin filament bundle assembly | 2-Jan | 21/3395 | 0.04761904762 | 80.83333333 | 8.908413313 | 0.01233468401 | 0.04891823018 | 0.02243618076 | 1 |
| PUAP | Landscape_Structure | GO:0051020 | MF | GTPase binding | 1-Jan | 76/2863 | 0.01315789474 | 37.67105263 | 6.055662856 | 0.02654558156 | 0.07055536151 | 0.03713440079 | 1 |
| PUAP | Light_Availability | GO:0051052 | BP | regulation of DNA metabolic process | 1-Jan | 108/3323 | 0.009259259259 | 30.76851852 | 5.456053383 | 0.03250075233 | 0.0915930293 | 0.05520462717 | 1 |
| PUAP | Light_Availability | GO:0051053 | BP | negative regulation of DNA metabolic process | 1-Jan | 50/3323 | 0.02 | 66.46 | 8.090735442 | 0.0150466446 | 0.06930054598 | 0.04176858034 | 1 |
| LEFI | Landscape_Structure | GO:0051087 | MF | protein-folding chaperone binding | 2-Jan | 33/2855 | 0.0303030303 | 43.25757576 | 6.46359506 | 0.02298773835 | 0.0664090219 | 0.0430179899 | 1 |
| PUAP | Light_Availability | GO:0051248 | BP | negative regulation of protein metabolic process | 1-Jan | 78/3323 | 0.01282051282 | 42.6025641 | 6.450004969 | 0.02347276557 | 0.07967401648 | 0.04802084185 | 1 |

| | | | | | | | | | | | | | |
|---|---|---|---|---|---|---|---|---|---|---|---|---|---|
| PURU | Light_Availability | GO:0051303 | BP | establishment of chromosome localization | 2-Jan | 47/3395 | 0.02127659574 | 36.11702128 | 5.885086656 | 0.02750014537 | 0.06416700585 | 0.02942998013 | 1 |
| PURU | Light_Availability | GO:0051310 | BP | metaphase chromosome alignment | 2-Jan | 40/3395 | 0.025 | 42.4375 | 6.399650638 | 0.02342867904 | 0.05897398759 | 0.02704821987 | 1 |
| PURU | Light_Availability | GO:0051311 | BP | meiotic metaphase chromosome alignment | 2-Jan | 18/3395 | 0.05555555556 | 94.30555556 | 9.635121872 | 0.01057727272 | 0.04787493491 | 0.02195767692 | 1 |
| PURU | Light_Availability | GO:0051315 | BP | attachment of mitotic spindle microtubules to kinetochore | 2-Jan | 24/3395 | 0.04166666667 | 70.72916667 | 8.321846611 | 0.01409053315 | 0.04891823018 | 0.02243618076 | 1 |
| PURU | Light_Availability | GO:0051316 | BP | attachment of meiotic spindle microtubules to kinetochore | 2-Jan | 18/3395 | 0.05555555556 | 94.30555556 | 9.635121872 | 0.01057727272 | 0.04787493491 | 0.02195767692 | 1 |
| PURU | Light_Availability | GO:0051345 | BP | positive regulation of hydrolase activity | 2-Jan | 80/3395 | 0.0125 | 21.21875 | 4.442592882 | 0.04657964371 | 0.08360448871 | 0.03834491587 | 1 |
| PURU | Light_Availability | GO:0051455 | BP | spindle attachment to meiosis I kinetochore | 2-Jan | 17/3395 | 0.05882352941 | 99.85294118 | 9.918893346 | 0.009991121819 | 0.04787493491 | 0.02195767692 | 1 |
| PURU | Light_Availability | GO:0051493 | BP | regulation of cytoskeleton organization | 2-Jan | 93/3395 | 0.01075268817 | 18.25268817 | 4.095328085 | 0.05404391185 | 0.0921705155 | 0.04227369508 | 1 |
| PURU | Light_Availability | GO:0051666 | BP | actin cortical patch localization | 2-Jan | 24/3395 | 0.04166666667 | 70.72916667 | 8.321846611 | 0.01409053315 | 0.04891823018 | 0.02243618076 | 1 |

| | | | | | | | | | | | | | |
|---|---|---|---|---|---|---|---|---|---|---|---|---|---|
| PUAP | Light_Availability | GO:0060260 | BP | regulation of transcription initiation by RNA polymerase II | 1-Jan | 23/3323 | 0.04347826087 | 144.4782609 | 11.97824114 | 0.006921456515 | 0.06219279767 | 0.03748462339 | 1 |
| PURU | Light_Availability | GO:0060341 | BP | regulation of cellular localization | 2-Jan | 99/3395 | 0.0101010101 | 17.14646465 | 3.958042278 | 0.05747906511 | 0.09467140136 | 0.04342071792 | 1 |
| PURU | Light_Availability | GO:0060583 | BP | regulation of actin cortical patch localization | 2-Jan | Apr-95 | 0.25 | 424.375 | 20.56696919 | 0.002355365051 | 0.02747925893 | 0.01260326913 | 1 |
| LEFI | Landscape_Structure | GO:0060589 | MF | nucleoside-triphosphatase regulator activity | 2-Jan | 96/2855 | 0.01041666667 | 14.86979167 | 3.659491986 | 0.0661311681 | 0.1158007585 | 0.07501263711 | 1 |
| PUAP | Landscape_Structure | GO:0060589 | MF | nucleoside-triphosphatase regulator activity | 1-Jan | 101/2863 | 0.009900990099 | 28.34653465 | 5.229391423 | 0.03527768075 | 0.07055536151 | 0.03713440079 | 1 |
| PURU | Light_Availability | GO:0060589 | MF | nucleoside-triphosphatase regulator activity | 2-Jan | 102/2917 | 0.009803921569 | 14.29901961 | 3.580716939 | 0.06872371553 | 0.09603950417 | 0.05897162537 | 1 |
| LEFI | Landscape_Structure | GO:0060590 | MF | ATPase regulator activity | 2-Jan | 19/2855 | 0.05263157895 | 75.13157895 | 8.582583433 | 0.01326800987 | 0.05703233276 | 0.03694402122 | 1 |
| USSU | Stand_Maturity | GO:0061013 | BP | regulation of mRNA catabolic process | 1-Jan | 64/3377 | 0.015625 | 52.765625 | 7.194833216 | 0.01895173231 | 0.09949659461 | 0.07106899615 | 1 |
| USST | Landscape_Structure | GO:0061134 | MF | peptidase regulator activity | 5-Jan | Oct-06 | 0.1 | 56.12 | 7.376247932 | 0.01770493894 | 0.0551606654 | 0.02903192916 | 1 |

| | | | | | | | | | | | | | |
|---|---|---|---|---|---|---|---|---|---|---|---|---|---|
| USST | Landscape_Structure | GO:0061135 | MF | endopeptidase regulator activity | 5-Jan | Jul-06 | 0.1428571429 | 80.17142857 | 8.859535093 | 0.01242000506 | 0.0551606654 | 0.02903192916 | 1 |
| PURU | Light_Availability | GO:0061572 | BP | actin filament bundle organization | 2-Jan | 23/3395 | 0.04347826087 | 73.80434783 | 8.504649853 | 0.01350542367 | 0.04891823018 | 0.02243618076 | 1 |
| USSU | Stand_ Maturity | GO:0070071 | BP | proton-transporting two-sector ATPase complex assembly | 1-Jan | Nov-77 | 0.09090909091 | 307 | 17.49285568 | 0.00325732899 | 0.0342019544 | 0.02442996743 | 1 |
| PUAP | Light_Availability | GO:0070461 | CC | SAGA-type complex | 2-Jan | 17/3225 | 0.05882352941 | 94.85294118 | 9.663568504 | 0.01051647527 | 0.07348017774 | 0.04687730637 | 1 |
| USST | Landscape_Structure | GO:0070566 | MF | adenylyltransferase activity | 5-Jan | Sep-06 | 0.1111111111 | 62.35555556 | 7.787961483 | 0.01594581455 | 0.0551606654 | 0.02903192916 | 1 |
| USSU | MAP | GO:0070595 | BP | (1->3)-alpha-glucan metabolic process | 4-Jan | Mar-77 | 0.3333333333 | 281.4166667 | 16.73073247 | 0.003550292745 | 0.0737671937 | 0.06560774897 | 1 |
| USSU | MAP | GO:0070596 | BP | (1->3)-alpha-glucan biosynthetic process | 4-Jan | Mar-77 | 0.3333333333 | 281.4166667 | 16.73073247 | 0.003550292745 | 0.0737671937 | 0.06560774897 | 1 |
| USSU | MAP | GO:0070597 | BP | cell wall (1->3)-alpha-glucan metabolic process | 4-Jan | Mar-77 | 0.3333333333 | 281.4166667 | 16.73073247 | 0.003550292745 | 0.0737671937 | 0.06560774897 | 1 |
| USSU | MAP | GO:0070598 | BP | cell wall (1->3)-alpha-glucan biosynthetic process | 4-Jan | Mar-77 | 0.3333333333 | 281.4166667 | 16.73073247 | 0.003550292745 | 0.0737671937 | 0.06560774897 | 1 |

| | | | | | | | | | | | | | |
|---|---|---|---|---|---|---|---|---|---|---|---|---|---|
| USSU | MAP | GO:0070599 | BP | fungal-type cell wall (1->3)-alpha-glucan metabolic process | 4-Jan | Mar-77 | 0.3333333333 | 281.4166667 | 16.73073247 | 0.003550292745 | 0.0737671937 | 0.06560774897 | 1 |
| USSU | MAP | GO:0070600 | BP | fungal-type cell wall (1->3)-alpha-glucan biosynthetic process | 4-Jan | Mar-77 | 0.3333333333 | 281.4166667 | 16.73073247 | 0.003550292745 | 0.0737671937 | 0.06560774897 | 1 |
| USST | Landscape_Structure | GO:0070628 | MF | proteasome binding | 5-Jan | Aug-06 | 0.125 | 70.15 | 8.273860674 | 0.01418417082 | 0.0551606654 | 0.02903192916 | 1 |
| USSU | MAP | GO:0070629 | BP | (1->4)-alpha-glucan metabolic process | 4-Jan | Mar-77 | 0.3333333333 | 281.4166667 | 16.73073247 | 0.003550292745 | 0.0737671937 | 0.06560774897 | 1 |
| USSU | MAP | GO:0070630 | BP | (1->4)-alpha-glucan biosynthetic process | 4-Jan | Mar-77 | 0.3333333333 | 281.4166667 | 16.73073247 | 0.003550292745 | 0.0737671937 | 0.06560774897 | 1 |
| PURU | Light_Availability | GO:0070650 | BP | actin filament bundle distribution | 2-Jan | Jun-95 | 0.1666666667 | 282.9166667 | 16.77797641 | 0.003532006148 | 0.03296539071 | 0.01511946491 | 1 |
| PURU | Light_Availability | GO:0070783 | BP | growth of unicellular organism as a thread of attached cells | 2-Jan | 63/3395 | 0.01587301587 | 26.94444444 | 5.045929063 | 0.03677441695 | 0.07461475903 | 0.03422180677 | 1 |
| PUAP | Light_Availability | GO:0070828 | BP | heterochromatin organization | 1-Jan | 87/3323 | 0.01149425287 | 38.1954023 | 6.09880335 | 0.0261811616 | 0.08100232673 | 0.04882143631 | 1 |

| | | | | | | | | | | | | | |
|---|---|---|---|---|---|---|---|---|---|---|---|---|---|
| PUAP | Light_Availability | GO:0071103 | BP | DNA conformation change | 1-Jan | 106/3323 | 0.009433962264 | 31.3490566 | 5.508997786 | 0.03189888655 | 0.0915930293 | 0.05520462717 | 1 |
| PURU | Light_Availability | GO:0071444 | BP | cellular response to pheromone | 2-Jan | 49/3395 | 0.02040816327 | 34.64285714 | 5.758467634 | 0.02866185932 | 0.06472032749 | 0.02968375922 | 1 |
| PURU | Light_Availability | GO:0071964 | BP | establishment of cell polarity regulating cell shape | 2-Jan | Apr-95 | 0.25 | 424.375 | 20.56696919 | 0.002355365051 | 0.02747925893 | 0.01260326913 | 1 |
| PURU | Light_Availability | GO:0072686 | CC | mitotic spindle | 2-Jan | 33/3301 | 0.0303030303 | 50.01515152 | 6.966699938 | 0.01989700091 | 0.05500935545 | 0.02217622392 | 1 |
| PURU | Light_Availability | GO:0090066 | BP | regulation of anatomical structure size | 2-Jan | 60/3395 | 0.01666666667 | 28.29166667 | 5.177698425 | 0.03503887567 | 0.07321556111 | 0.03358006938 | 1 |
| PUAP | Light_Availability | GO:0090329 | BP | regulation of DNA-templated DNA replication | 1-Jan | 48/3323 | 0.02083333333 | 69.22916667 | 8.260094834 | 0.01444477881 | 0.06930054598 | 0.04176858034 | 1 |
| PURU | Light_Availability | GO:0097574 | CC | lateral part of cell | 2-Jan | Jun-01 | 0.1666666667 | 275.0833333 | 16.5428889 | 0.003632508055 | 0.02134098483 | 0.008603308552 | 1 |
| PURU | Light_Availability | GO:0097575 | CC | lateral cell cortex | 2-Jan | Jun-01 | 0.1666666667 | 275.0833333 | 16.5428889 | 0.003632508055 | 0.02134098483 | 0.008603308552 | 1 |
| PUAP | Landscape_Structure | GO:0097708 | CC | intracellular vesicle | 1-Jan | 188/3225 | 0.005319148936 | 17.15425532 | 4.019235664 | 0.05829457364 | 0.07704233751 | 0.038163387 | 1 |
| PUAP | Landscape_Structure | GO:0098772 | MF | molecular function regulator activity | 1-Jan | 215/2863 | 0.004651162791 | 13.31627907 | 3.509455666 | 0.07509605309 | 0.08582406068 | 0.04517055825 | 1 |

| | | | | | | | | | | | | | |
|---|---|---|---|---|---|---|---|---|---|---|---|---|---|
| PUAP | Landscape_Structure | GO:0098791 | CC | Golgi apparatus subcompartment | 1-Jan | 95/3225 | 0.01052631579 | 33.94736842 | 5.739979828 | 0.02945736434 | 0.06259689922 | 0.03100775194 | 1 |
| PURU | Light_Availability | GO:0099081 | CC | supramolecular polymer | 2-Jan | 41/3301 | 0.0243902439 | 40.25609756 | 6.226888119 | 0.02469040603 | 0.05802245417 | 0.02339091098 | 1 |
| PURU | Light_Availability | GO:0099512 | CC | supramolecular fiber | 2-Jan | 41/3301 | 0.0243902439 | 40.25609756 | 6.226888119 | 0.02469040603 | 0.05802245417 | 0.02339091098 | 1 |
| PURU | Light_Availability | GO:0099513 | CC | polymeric cytoskeletal fiber | 2-Jan | 40/3301 | 0.025 | 41.2625 | 6.307193341 | 0.02409187299 | 0.05802245417 | 0.02339091098 | 1 |
| PURU | Light_Availability | GO:0099738 | CC | cell cortex region | 2-Jan | 64/3301 | 0.015625 | 25.7890625 | 4.93014338 | 0.03840599267 | 0.07050021573 | 0.0284211396 | 1 |
| PURU | Light_Availability | GO:0110053 | BP | regulation of actin filament organization | 2-Jan | 34/3395 | 0.02941176471 | 49.92647059 | 6.960301902 | 0.01993208148 | 0.05813523764 | 0.02666353005 | 1 |
| PUAP | Light_Availability | GO:0140030 | MF | modification-dependent protein binding | 1-Jan | 26/2863 | 0.03846153846 | 110.1153846 | 10.44583097 | 0.009081383165 | 0.06444289207 | 0.04301707815 | 1 |
| PUAP | Light_Availability | GO:0140034 | MF | methylation-dependent protein binding | 1-Jan | Oct-63 | 0.1 | 286.3 | 16.89082591 | 0.003492839679 | 0.03580160671 | 0.02389837675 | 1 |
| USSU | Stand_Maturity | GO:0140098 | MF | catalytic activity, acting on RNA | 1-Jan | 184/2917 | 0.005434782609 | 15.85326087 | 3.853992848 | 0.06307850531 | 0.07569420638 | 0.02655937066 | 1 |
| LEFI | Landscape_Structure | GO:0140101 | MF | catalytic activity, acting on a tRNA | 2-Jan | 73/2855 | 0.01369863014 | 19.55479452 | 4.251381838 | 0.05049330095 | 0.112359585 | 0.07278353687 | 1 |
| USSU | Stand_Maturity | GO:0140101 | MF | catalytic activity, acting on a tRNA | 1-Jan | 69/2917 | 0.01449275362 | 42.27536232 | 6.42459044 | 0.02365443949 | 0.04319506342 | 0.0151561626 | 1 |

| | | | | | | | | | | | | | |
|---|---|---|---|---|---|---|---|---|---|---|---|---|---|
| USSU | Stand_ Maturity | GO:0140640 | MF | catalytic activity, acting on a nucleic acid | 1-Jan | 283/2917 | 0.003533568905 | 10.30742049 | 3.050806532 | 0.09701748372 | 0.09701748372 | 0.03404122236 | 1 |
| PURU | Light_Availability | GO:0140677 | MF | molecular function activator activity | 2-Jan | 107/2917 | 0.009345794393 | 13.63084112 | 3.48626567 | 0.07202962812 | 0.09603950417 | 0.05897162537 | 1 |
| USST | Landscape_Structure | GO:0140678 | MF | molecular function inhibitor activity | 5-Jan | 25/2806 | 0.04 | 22.448 | 4.550416939 | 0.04379131839 | 0.09445441501 | 0.04971285001 | 1 |
| LEFI | Landscape_Structure | GO:0140678 | MF | molecular function inhibitor activity | 2-Jan | 21/2855 | 0.04761904762 | 67.97619048 | 8.154949248 | 0.01465948796 | 0.05703233276 | 0.03694402122 | 1 |
| PUAP | Light_Availability | GO:0140719 | BP | constitutive heterochromatin formation | 1-Jan | 52/3323 | 0.01923076923 | 63.90384615 | 7.931194497 | 0.01564851038 | 0.06930054598 | 0.04176858034 | 1 |
| PUAP | Light_Availability | GO:0141137 | BP | positive regulation of gene expression, epigenetic | 1-Jan | 19/3323 | 0.05263157895 | 174.8947368 | 13.18691536 | 0.005717724947 | 0.06219279767 | 0.03748462339 | 1 |
| USST | DD_18 | GO:1900750 | MF | oligopeptide binding | 11-Jan | Mar-06 | 0.3333333333 | 85.03030303 | 9.133913143 | 0.01171863108 | 0.09694503894 | 0.07176960662 | 1 |
| PUAP | Light_Availability | GO:1901983 | BP | regulation of protein acetylation | 1-Jan | Nov-23 | 0.09090909091 | 302.0909091 | 17.35197133 | 0.003310261812 | 0.05130905808 | 0.03092481429 | 1 |
| PUAP | Light_Availability | GO:1901984 | BP | negative regulation of protein acetylation | 1-Jan | Apr-23 | 0.25 | 830.75 | 28.80538144 | 0.001203731568 | 0.0373156786 | 0.02249077403 | 1 |
| PUAP | Light_Availability | GO:1902275 | BP | regulation of chromatin organization | 1-Jan | 70/3323 | 0.01428571429 | 47.47142857 | 6.816995568 | 0.02106530244 | 0.07967401648 | 0.04802084185 | 1 |

| | | | | | | | | | | | | | | |
|---|---|---|---|---|---|---|---|---|---|---|---|---|---|---|
| USSU | Stand_ Maturity | GO:190 2369 | BP | negative regulation of RNA catabolic process | 1-Jan | Jul-77 | 0.1428571429 | 482.4285714 | 21.94148061 | 0.002072 845721 | 0.029019 84009 | 0.02072 845721 | | 1 |
| USSU | Stand_ Maturity | GO:190 2373 | BP | negative regulation of mRNA catabolic process | 1-Jan | Jul-77 | 0.1428571429 | 482.4285714 | 21.94148061 | 0.002072 845721 | 0.029019 84009 | 0.02072 845721 | | 1 |
| PURU | Light_A vailabili ty | GO:190 2903 | BP | regulation of supramolecula r fiber organization | 2-Jan | 42/3395 | 0.0238095238 1 | 40.41666667 | 6.239743685 | 0.024592 82299 | 0.059361 98653 | 0.02722 617428 | | 1 |
| USSU | Stand_ Maturity | GO:190 3312 | BP | negative regulation of mRNA metabolic process | 1-Jan | Oct-77 | 0.1 | 337.7 | 18.34938691 | 0.002961 208173 | 0.034201 9544 | 0.02442 996743 | | 1 |
| PUAP | Light_A vailabili ty | GO:190 5268 | BP | negative regulation of chromatin organization | 1-Jan | 34/3323 | 0.0294117647 1 | 97.73529412 | 9.835410216 | 0.010231 71833 | 0.069300 54598 | 0.04176 858034 | | 1 |
| USST | Landsca pe_Stru cture | GO:199 0251 | CC | nuclear exosome focus | 4-Jan | Mar-78 | 0.3333333333 | 264.8333333 | 16.2275729 | 0.003772 394882 | 0.045247 363 | 0.03528 059493 | | 1 |
| PURU | Light_A vailabili ty | GO:199 0498 | CC | mitotic spindle microtubule | 2-Jan | May-01 | 0.2 | 330.1 | 18.13009555 | 0.003027 549044 | 0.021340 98483 | 0.00860 3308552 | | 1 |
| PURU | Light_A vailabili ty | GO:199 0758 | BP | mitotic sister chromatid biorientation | 2-Jan | Nov-95 | 0.0909090909 1 | 154.3181818 | 12.36385243 | 0.006470 571389 | 0.043137 14259 | 0.01978 47045 | | 1 |
| PUAP | Light_A vailabili ty | GO:200 0104 | BP | negative regulation of DNA- templated | 1-Jan | 24/3323 | 0.0416666666 7 | 138.4583333 | 11.72426259 | 0.007222 389407 | 0.062192 79767 | 0.03748 462339 | | 1 |

| | | | | | | | | | | | | | |
|---|---|---|---|---|---|---|---|---|---|---|---|---|---|
| | | | | DNA replication | | | | | | | | | |
| PURU | Light_Availability | GO:2000114 | BP | regulation of establishment of cell polarity | 2-Jan | Aug-95 | 0.125 | 212.1875 | 14.51725787 | 0.004707952959 | 0.0346901797 | 0.01591053354 | 1 |
| PUAP | Light_Availability | GO:2000142 | BP | regulation of DNA-templated transcription initiation | 1-Jan | 23/3323 | 0.04347826087 | 144.4782609 | 11.97824114 | 0.006921456515 | 0.06219279767 | 0.03748462339 | 1 |
| PURU | Light_Availability | GO:2000243 | BP | positive regulation of reproductive process | 2-Jan | 83/3395 | 0.01204819277 | 20.45180723 | 4.355447931 | 0.0483047707 | 0.08560339111 | 0.0392617057 | 1 |
| PUAP | Light_Availability | GO:2000756 | BP | regulation of peptidyl-lysine acetylation | 1-Jan | Oct-23 | 0.1 | 332.3 | 18.20164828 | 0.00300932892 | 0.05130905808 | 0.03092481429 | 1 |
| PUAP | Light_Availability | GO:2000757 | BP | negative regulation of peptidyl-lysine acetylation | 1-Jan | Apr-23 | 0.25 | 830.75 | 28.80538144 | 0.001203731568 | 0.0373156786 | 0.02249077403 | 1 |
| PURU | Light_Availability | GO:2000769 | BP | regulation of establishment or maintenance of cell polarity regulating cell shape | 2-Jan | 18/3395 | 0.05555555556 | 94.30555556 | 9.635121872 | 0.01057727272 | 0.04787493491 | 0.02195767692 | 1 |
| PURU | Light_Availability | GO:2000771 | BP | positive regulation of establishment or maintenance of cell polarity | 2-Jan | Sep-95 | 0.1111111111 | 188.6111111 | 13.68092054 | 0.005295666007 | 0.03706966205 | 0.01700187507 | 1 |

| | | | | | | | | | | | | | |
|---|---|---|---|---|---|---|---|---|---|---|---|---|---|
| | | | | regulating cell shape | | | | | | | | | |
| PURU | Light_Availability | GO:2000782 | BP | regulation of establishment of cell polarity regulating cell shape | 2-Jan | Apr-95 | 0.25 | 424.375 | 20.56696919 | 0.002355365051 | 0.02747925893 | 0.01260326913 | 1 |
| PURU | Light_Availability | GO:2000784 | BP | positive regulation of establishment of cell polarity regulating cell shape | 2-Jan | Apr-95 | 0.25 | 424.375 | 20.56696919 | 0.002355365051 | 0.02747925893 | 0.01260326913 | 1 |
| PUAP | Light_Availability | GO:2001251 | BP | negative regulation of chromosome organization | 1-Jan | 79/3323 | 0.01265822785 | 42.06329114 | 6.408064539 | 0.02377369847 | 0.07967401648 | 0.04802084185 | 1 |

**Table S9.**
Gene ontology terms.

| Site Code | Site Name | State | County | Locality Data | Habitat Data | Latitude | Longitude |
|---|---|---|---|---|---|---|---|
| ALM | GSMNP - LeConte | TN | Sevier | Great Smoky Mountains National Park, Mount LeConte, N-facing slopes above Trillium Gap Trail, 0.4 mi N of jct w/ Rainbow Falls Trail/Boulevard Trail at LeConte Lodge | Spruce (Picea) – fir (Abies) forest with abundant hardwoods (Betula alleghaniensis, Prunus pensylvanica, Sorbus), Viburnum understory and non-calcareous rock outcrops | 35.65878 | -83.43787 |
| ASG | CNF - Cloudland | TN | Carter | Cherokee National Forest, Roan Mountain, Appalachian Trail 0.1 mi E of Cloudland Hotel Site, just N of Toll House Gap | Spruce (Picea rubens) – fir (Abies fraseri) forest with sparse hardwoods (Betula alleghaniensis, Sorbus) and Rhododendron catawbiense | 36.10455 | -82.1293 |
| BEK | PNF - Buckeye Ridge | NC | Madison | Pisgah National Forest, Bald Mountains, N-slope of Roaring Fork Mountain, spur of Buckeye Ridge | Northern hardwood (Acer saccharum, Betula alleghaniensis, Fagus, Ilex montana, Magnolia acuminata, M. fraseri, Quercus rubra) forest | 35.8014 | -82.95096 |
| BFF | PNF - Bluff Mountain | NC | Madison | Pisgah National Forest, Bald Mountains, N-Facing slopes of Bluff Mountain summit, NE of Appalachian Trail 1 mi E of Catpen Gap, 3.4 mi E of Lemon Gap | Mature mixed hardwood (Aesculus, Betula alleghaniensis, Acer saccharum Amelanchier, Prunus, Tilia) forest with non-calcareous rock outcrops and Rhododendron stands | 35.84115 | -82.90576 |
| BLU | BBP - Blueberry Cliff | KY | Letcher | Pine Mountain, Bad Branch Nature Preserve, headwaters of Presley House Branch, | Mixed hardwood (Acer saccharum, Carya, Liriodendron, Oxydendrum, | 37.08482 | -82.79308 |

| | | | | | | | |
|---|---|---|---|---|---|---|---|
| | | | | Pine Mountain Trail ~1.5 mi E of US119 | Quercus alba, Q. rubra, Q. prinus, Tilia) forest with Rhododendron-Kalmia thickets and sandstone outcrops | | |
| BRF | SNP - Bearfence Mountain | VA | Page | Shenandoah National Park, Central District, Bearfence Mountain, W-slopes below Appalachian Trail, 0.3 mi S of jct w/ spur trail to Skyline Drive/Bearfence Parking Area | Quercus rubra dominated mixed hardwood (Q. rubra, Betula lenta, Tilia) forest with Kalmia-Ilex montana-Hamamelis understory and shaded schist boulders and outcrops | 38.4477 | -78.46635 |
| BUK | GSMNP - Buck Knob | NC | Haywood | Great Smoky Mountains National Park, Cataloochee Divide, N-slope of Little Bald Knob, Hemphill Bald Trail ~3 mi E of Polls Gap | Mature northern hardwood (Acer pennsylvanicum, A. saccharum, A. spicatum, Aesculus, Betula alleghaniensis, Fagus, Prunus) and scattered conifers (Picea rubens, dead Tsuga) | 35.552276 | -83.120161 |
| CAT | PNF - Cattail Peak | NC | Yancey | Pisgah National Forest, summit of Cattail Peak, Black Mountain Crest Trail 3 mi N of Mount Mitchell, ~8 mi S of US19E | Abies dominated Abies-Picea forest with sparse Sorbus and small rock outcrops | 35.7964 | -82.2569 |
| COG | NNF - Cole Gap | NC | Macon | Nantahala National Forest, N-facing slopes above FS4535 0.3 mi W of jct w/ SR1538/Buck Creek Rd. at Cole Gap | Ruderal mixed hardwoods (Acer, Betula lenta, Carya, Fraxinus, Halesia, Liriodendron, Prunus, Robinia, Quercus) with occasional large trees and shaded non-calcareous rock outcrops | 35.1036 | -83.2075 |

| | | | | | | | |
|---|---|---|---|---|---|---|---|
| DAD | GMSP - Shanty Spring | NC | Avery | Grandfather Mountain State Park, Grandfather Mountain, N-slopes N of Profile Trail, ~0.3 mi W of jct w/ Grandfather Trail | Mature northern hardwood (Acer pensylvanicum, A. spicatum, Aesculus, Betula alleghaniensis) – spruce (Picea) forest with large non-calcareous rock outcrops | 36.11211 | -81.81832 |
| DBG | GSMNP - Double Gap | NC | Haywood | Great Smoky Mountains National Park, Cataloochee Divide, McClue Ridge, Double Gap Trail ~0.1 mi W of Double Gap | Mature northern hardwood (Acer pensylvanicum, A. saccharum, A. spicatum, Aesculus, Betula alleghaniensis, Fagus, Prunus) forest with dead Tsuga and occasional schist boulders | 35.57191 | -83.1035 |
| GOS | KSF - Goss Park | KY | Harlan | E-slopes of Pine Mountain, Kentenia State Forest, vicinity of Goss Park, E of Little Shepherd Trail/SR1679 at jct w/ Putney Lookout Tower Rd., between Spruce Ridge Branch and Hi Lewis Branch | Large sandstone outcrops with mixed hardwood (Acer saccharum, Betula lenta, Carya, Liriodendron, Quercus alba, Q. rubra, Q. prinus, Sassafras) – conifer (Pinus rigida, P. strobus, P. virginiana, Tsuga) forest and Rhododendron-Kalmia thickets | 36.9243 | -83.22419 |
| GRI | SNP - Grindstone | VA | Page | Shenandoah National Park, Central District, S facing slopes of Grindstone Mountain above Deep Run, ~0.2 mi S of summit | Massive S-facing quartz talus slope with occasional conifers (Pinus pungens, P. rigida, P. strobus) – hardwoods (Acer rubrum, Amelanchier, Betula lenta, Quercus rubra) and Kalmia-Vaccinium | 38.47288 | -78.550732 |
| HAW | SNP - Hawksbill | VA | Madison | Shenandoah National Park, Central District, SW slopes of Hawksbill Mountain, | Basalt outcrops and talus in mixed Abies balsamea – hardwood (Betula | 38.55397 | -78.39687 |

| | | | | | | | |
|---|---|---|---|---|---|---|---|
| | | | | between Lower Hawksbill Trail and Hawksbill Fire Road ~0.7 mi W of Hawksbill Gap | alleghaniensis, Ilex montana, Quercus rubra, Sorbus) forest with sparse Kalmia and Vaccinium | | |
| JON | JNF - Ron Jon | VA | Bland | Jefferson National Forest, E slopes of Round Mountain, S of Round Mountain Rd./VA688, ~1.5 mi W of US52/Bastian | Mixed hardwood (Acer, Betula lenta, Carya, Oxydendrum, Quercus alba, Q. rubra, Q. prinus) – Pinus strobus forest with ericaceous understory (Vaccinium, sparse Kalmia and Rhododendron maximum) | 37.170518 | -81.164224 |
| KLE | MNF - Bickle Knob | WV | Randolph | Monongahela National Forest, Bickle Knob, saddle and W-facing slopes below observation tower, terminus of FSR91A | Open mature mixed northern hardwood (Acer saccharum, Betula alleghaniensis, Fagus, Prunus, Quercus rubra) forest with Crataegus and Ilex montana, and small sandstone outcrops | 38.93557 | -79.730504 |
| LEM | BBP - Lemon Squeezer | KY | Letcher | Pine Mountain, Bad Branch Nature Preserve, W headwaters of Bad Branch, Pine Mountain Trail ~2.2 mi E of US119 | Mixed hardwood (Acer saccharum, Carya, Liriodendron, Oxydendrum, Quercus alba, Q. rubra, Q. prinus, Tilia) forest with Rhododendron-Kalmia thickets | 37.09032 | -82.77928 |
| LUE | MNF - Blue Knob | WV | Pocahontas | Monongahela National Forest, Spruce Mountain, W-facing slopes below summit of Blue Knob, ~1 mi S of WV39 | Mixed hardwood (Acer, Betula alleghaniensis, Fagus, Quercus rubra) forest with scattered Ilex montana-Crataegus-Acer pensylvanicum, with sandstone outcrops and boulders | 38.175865 | -80.279981 |

| | | | | | | | |
|---|---|---|---|---|---|---|---|
| MAX | PNF - Max Patch | NC | Madison | Pisgah National Forest, Max Patch Mountain, N-slopes of Max Patch below summit, 0.4 mi E of Max Patch Rd./CR-1182 | Mixed hardwood (Acer saccharum dominant, A. pensylvanicum, Aesculus, Quercus rubra) forest on edge of grass bald, with granitic rock outcrops in shade | 35.79914 | -82.95679 |
| MLC/ LCN | GSMNP - LeConte | TN | Sevier | Great Smoky Mountains National Park, Mount LeConte, N-facing slopes above Trillium Gap Trail, 0.4 mi N of jct w/ Rainbow Falls Trail/Boulevard Trail at LeConte Lodge | Spruce (Picea) – fir (Abies) forest with abundant hardwoods (Betula alleghaniensis, Prunus pensylvanica, Sorbus), Viburnum understory and non-calcareous rock outcrops | 35.65878 | -83.43787 |
| NAL | GSMNP - LeConte | TN | Sevier | Great Smoky Mountains National Park, Mount LeConte, N-facing slopes above Trillium Gap Trail, 0.4 mi N of jct w/ Rainbow Falls Trail/Boulevard Trail at LeConte Lodge | Spruce (Picea) – fir (Abies) forest with abundant hardwoods (Betula alleghaniensis, Prunus pensylvanica, Sorbus), Viburnum understory and non-calcareous rock outcrops | 35.65878 | -83.43787 |
| NAR | PNF - Stairs Mountain | NC | Haywood | Pisgah National Forest, Balsam Mountains, Shining Rock Wilderness, Shining Rock Ledge, E slopes of Stairs Mountain summit, Art Loeb Trail 1 mi N of Shining Rock Gap | Mature northern hardwood (Amelanchier, Betula alleghaniensis, Quercus rubra) – spruce (Picea) forest with ericaceous understory (Rhododendron catawbiense, R. maximum) and schist outcrops | 35.37817 | -82.86893 |
| NGA | MNF - Not Gudineer | WV | Pocahontas | Monongahela National Forest, Shavers Mountain, E-facing slopes of Gaudineer Knob, FSR27 ~1.25 mi N of jct w/ US250 | Mixed hardwood (Acer pensylvanicum, A. saccharum, Betula alleghaniensis, Fagus, Fraxinus) – Picea forest with sandstone boulders and mature | 38.611691 | -79.841629 |

| | | | | | | | |
|---|---|---|---|---|---|---|---|
| | | | | | trees | | |
| POK | MNF - Pointy Knob | WV | Tucker | Monongahela National Forest, W-facing slopes of Canaan Mountain, E of Pointy Knob, Pointy Knob Trail ~2.5 mi S of FSR13 | Regenerating burn with open hardwood (Acer saccharum, Prunus) – spruce (Picea rubens) stands, Ilex montana understory and conglomerate boulders | 39.03528 | -79.48914 |
| RBB | CHNF - Rabun Bald | GA | Rabun | Chattahoochee National Forest, Rabun Bald, jct of Bartram Trail ~1.7 mi S of Beegum Gap | Grass bald with mixed hardwoods (Quercus rubra dominant, Acer pensylvanica, Amelanchier), Kalmia-Rhododendron thickets, schist outcrops, and observation tower | 34.96532 | -83.2999 |
| ROG | JNF - Mount Rogers | VA | Grayson | Jefferson National Forest, Mount Rogers National Recreation Area, Lewis Fork Wilderness, S-facing slopes of Mount Rogers, Mount Rogers Spur Trail 0.25 mi E of summit, ~0.25 mi W of jct w/ Appalachian Trail | Dense to open Spruce (Picea rubens) – fir (Abies fraseri) forest with sparse hardwoods (Betula alleghaniensis, Prunus, Sorbus) and Rubus in openings | 36.659449 | -81.540821 |
| RON | JNF - Round Mountain | VA | Bland | Jefferson National Forest, Round Mountain, vicinity of summit, Round Mountain Rd//VA688, ~4 mi W of US52/Bastian | Mixed hardwood (Acer, Betula lenta, Carya, Fagus, Oxydendrum, Quercus alba, Q. rubra) – conifer (Pinus strobus, Tsuga) forest with Ilex montana understory | 37.154716 | -81.194163 |
| RSK | MNF - Red Spruce Knob | WV | Pocahontas | Monongahela National Forest, S-facing slopes of Red Spruce Knob, N of Red Lick Mountain, 0.1 mi W of | Mixed hardwood (Acer, Betula lenta, Fagus, Quercus rubra) – Picea forest with sandstone outcrops and boulders. | 38.323665 | -80.152203 |

| | | | | | | | |
|---|---|---|---|---|---|---|---|
| | | | | WV150 and FSR451 | | | |
| SHR | PNF - Stairs Mountain | NC | Haywood | Pisgah National Forest, Balsam Mountains, Shining Rock Wilderness, Shining Rock Ledge, E slopes of Stairs Mountain summit, Art Loeb Trail 1 mi N of Shining Rock Gap | Mature northern hardwood (Amelanchier, Betula alleghaniensis, Quercus rubra) – spruce (Picea) forest with ericaceous understory (Rhododendron catawbiense, R. maximum) and schist outcrops | 35.37817 | -82.86893 |
| SNR | KSF - Snout Rock | KY | Harlan | E-slopes of Pine Mountain, Kentenia State Forest, S of Little Shepherd Trail/SR1679 between Mill Cliff Branch and Hi Lewis Branch | Massive sandstone outcrops with mixed hardwood (Acer saccharum, Betula lenta, Liriodendron, Quercus alba, Q. rubra, Q. prinus, Sassafras) – conifer (P. strobus, P. virginiana) forest and Rhododendron-Kalmia thickets | 36.920858 | -83.230543 |
| SPF | PNF - Roan High Bluff | NC | Mitchell | Pisgah National Forest, Roan Mountain, S slope of Roan High Bluff, ~0.2 mi E of overlook & ~0.3 mi W of parking area. | Fir (Abies) dominated forest with sparse Picea, Sorbus and Rhododendron catawbiense amid large granitic rock outcrops | 36.09362 | -82.14485 |
| TMK | JNF - Toms Knob | VA | Grayson | Jefferson National Forest, Mount Rogers National Recreation Area, Lewis Fork Wilderness, E-facing slopes of Pine Mountain, NW of Appalachian Trail ~0.6 E of Thomas Knob Shelter | Beech (Fagus)-spruce (Picea rubens)-fir (Abies fraseri) forest with hardwoods (Betula alleghaniensis, Prunus, Sorbus) and Rhododendron-Vaccinium thickets | 36.655015 | -81.526364 |
| TOM | PNF - Blasam Cone | NC | Yancey | Mount Mitchell State Park, vicinity of summit of Balsam | Spruce (Picea) –fir (Abies) forest with sparse Betula, non- | 35.7892 | -82.2558 |

| | | | | | | | |
|---|---|---|---|---|---|---|---|
| | | | | Cone, Black Mountain Crest Trail ~2 mi N of Mount Mitchell and terminus of NC128 | calcareous rock outcrops and understory of Rhododendron and Sorbus | | |
| TRI | MNF - Trillium Knob | WV | Pocahontas | Monongahela National Forest, Shavers Mountain, E-facing slopes of Trillium Knob above Old Road Run, S of Glade Run headwaters, Allegheny Trail ~1.5 mi N of FSR27 | Mixed hardwood (Acer pensylvanicum, A. saccharum, Betula alleghaniensis, Fagus, Fraxinus, Magnolia tripetala, Ostrya, Tilia) – Picea forest with sandstone boulders and remnant mature trees | 38.644182 | -79.83029 |
| TYS | SNP - Bettys Rock | VA | Page | Shenandoah National Park, Central District, Bettys Rock, ~0.3 mi N of Crescent Rock, upper W-facing slopes of Timber Hollow | Large exposed outcrop, cliff and associated talus, surrounded by hardwood forest (Quercus rubra dominant, Acer saccharum, Betula alleghaniensis, Carya, Fraxinus, Quercus alba) with Ilex montana-Acer pensylvanicum-Hamamelis understory and occasional thickets of Kalmia-Vaccinium-stunted Q. rubra | 38.56753 | -78.38223 |
| WHI | NNF - Whiterock Mountain | NC | Macon | Nantahala National Forest, summit of Whiterock Mountain, spur trail 0.3 mi N of jct w/ Bartram Trail, 1.3 mi NW of Jones Gap parking area at terminus of FS 4522 | Extensive granite dome with Kalmia-Rhododendron shrub balds and mixed hardwood (Carya, Quercus alba dominant) forest | 35.085 | -83.3053 |

**Table S10.**
Population genetic sampling sites with locality and habitat data.

| Site Code | Landscape | Region | *Lepraria finkii* | *Lepraria lanata* | *Puntelia appalachensis* | *Punctelia rudecta* | *Usnea strigosa* | *Unsea subfusca* | **Total Individuals Sampled** |
|---|---|---|---|---|---|---|---|---|---|
| ALM | S.Appalachians | South | | 21 | | | | | **21** |
| ASG | S.Appalachians | South | | 11 | | | | | **11** |
| BEK | S.Appalachians | South | 5 | | 10 | 10 | | 7 | **32** |
| BFF | S.Appalachians | South | 10 | | 10 | 10 | | 9 | **39** |
| BLU | Kentucky | Central | 11 | | 10 | 11 | | | **32** |
| BRF | ShenandoahNP | Central | 12 | | 10 | 12 | | | **34** |
| BUK | S.Appalachians | South | 8 | | 10 | 10 | | 10 | **38** |
| CAT | S.Appalachians | South | 10 | 14 | | 10 | | | **34** |
| COG | S.Appalachians | South | 5 | | | | | 1 | **6** |
| DAD | S.Appalachians | South | | 10 | 5 | | | | **15** |
| DBG | S.Appalachians | South | | | 10 | 10 | | | **20** |
| GOS | Kentucky | Central | 10 | | 10 | 15 | 10 | | **45** |
| GRI | ShenandoahNP | Central | 11 | | | 10 | | | **21** |
| HAW | ShenandoahNP | Central | 13 | | 8 | 14 | | | **35** |
| JON | SE.Virginia | Central | 11 | | | 12 | 9 | | **32** |
| KLE | W.Virginia | Central | 9 | | 10 | 12 | | | **31** |
| LEM | Kentucky | Central n | 9 | | 4 | 12 | | | **25** |
| LUE | W.Virginia | Central | 9 | | 12 | 12 | | | **33** |
| MAX | S.Appalachians | South | | | | | 9 | | **9** |

| | | | | | | | | | |
|---|---|---|---|---|---|---|---|---|---|
| LCN | S.Appalachians | South | | 27 | | | | | **27** |
| NAL | S.Appalachians | South | | 9 | | | | | **9** |
| NAR | S.Appalachians | South | | | | 8 | | 16 | **24** |
| NGA | W.Virginia | Central | 12 | | 10 | 7 | | 8 | **37** |
| POK | W.Virginia | Central | 10 | | | 10 | | | **20** |
| RBB | S.Appalachians | South | | 9 | | | | | **9** |
| ROG | SE.Virginia | Central | 6 | | | 8 | | | **14** |
| RON | SE.Virginia | Central | 10 | | 11 | 11 | 3 | | **35** |
| RSK | W.Virginia | Central | 11 | | 11 | 12 | | | **34** |
| SHR | S.Appalachians | South | 5 | | | 6 | | 18 | **29** |
| SNR | Kentucky | Central | 10 | | 9 | 11 | 4 | | **34** |
| SPF | S.Appalachians | South | | 3 | | | | | **3** |
| TMK | SE.Virginia | Central | 5 | | | 10 | | | **15** |
| TOM | S.Appalachians | South | 8 | 11 | | 10 | | | **29** |
| TRI | W.Virginia | Central | 8 | | 11 | 10 | | 12 | **41** |
| TYS | ShenandoahNP | Central | 11 | | 12 | 9 | | | **32** |
| WHI | S.Appalachians | South | 9 | | 14 | 24 | 10 | | **57** |
| **Total Individuals Sampled for Each Species** | | | **238** | **115** | **187** | **296** | **45** | **81** | **962** |

**Table S11.**
Sampling intensities for each species at population genetic sampling sites.

| Variable | Description | | | | Score |
|---|---|---|---|---|---|
| **Rock Cover (RC)** | **The percentage of exposed rock** | | | | **0-5** |
| Percent exposed rock | | | | | |
| No exposed rock present (0%) | 0 | | | | |
| Few exposed rocks present (1-20%) | 2 | | | | |
| Moderate number of exposed rocks (21-50%) | 3 | | | | |
| High number of exposed rock (51-80%) | 4 | | | | |
| Very high number of expsoed rock (81-100%) | 5 | | | | |
| **Large Trees (LT)** | **The percentage of living, mature canopy trees** | | | | **0-10** |
| Percent mature canopy trees | >70% alive | 30-70% alive | <30% alive | | |
| None | 0 | 0 | 0 | | |
| 0-20 | 3 | 2 | 1 | | |
| 20-40 | 4 | 3 | 2 | | |
| 40-70 | 6 | 9 | 4 | | |
| **Canopy Cover (CC)** | **The percentage of living, native canopy trees** | | | | **0-5** |
| Percent native canopy cover | >70% alive | 30-70% alive | <30% alive | | |
| <50% | 0 | 0 | 0 | | |
| 50-90% | 3 | 2 | 1 | | |
| >90% | 5 | 4 | 3 | | |
| **Understory (US)** | **The percentage of native understory plant species and the extent of anthropogenic modification** | | | | **0-25** |
| Percent native understory | Degree of anthropogenic modification | | | | |
| | 0 | N/A | | 0 | |
| 0-50% | | N/A | | 5 | |

| | | | | | |
|---|---|---|---|---|---|
| 50-90% | | >50% | | 10 | |
| 50-90% | | <50% | | 15 | |
| >90% | | >50% | | 15 | |
| >90% | | <50% | | 20 | |
| >90% | | | 0 | 25 | |
| **Weediness (WED)** | **The percentage of non-native and invasive plants** | | | | **0-15** |
| Percent weed cover | No high-threat taxa | <50% high-threat taxa | >50% high-threat taxa | | |
| >50% | 4 | 2 | 0 | | |
| 20-50% | 7 | 6 | 4 | | |
| 5-25% | 11 | 9 | 7 | | |
| <5% | 15 | 13 | 11 | | |
| **Recruitment (REC)** | **The percentage of native tree species recruitment** | | | | **0-10** |
| Part 1 | Part 2 | | | | |
| No native species recruiting (0%) | Anthrop. impacts | 0 | | | |
| No native species recruiting (0%) | Natural impacts | 5 | | | |
| Native species recruiting (>0%) | <50% recruitment | 5 | | | |
| Native species recruiting (>0%) | >50% recruitment | 10 | | | |
| **Litter (LIT)** | **The percentage of leaf litter from native versus non-native tree species** | | | | **0-5** |
| Percent litter | >50% from non-natives | <50% from non-natives | | | |
| <10% | 0 | 0 | | | |
| 10-100% | 3 | 5 | | | |
| **Loggyness (CWD)** | **The percentage of coarse woody debris** | | | | **0-5** |
| Percent coarse woody debris | | | | | |

| | | | | | |
|---|---|---|---|---|---|
| No log present (0%) | 0 | | | | |
| Few logs present (1-20%) | 2 | | | | |
| Moderate number of logs (21-50%) | 3 | | | | |
| High number of logs (51-80%) | 4 | | | | |
| Very high number of logs (81-100%) | 5 | | | | |
| **Patch size (PS)** | **The equivalent to the size of contiguous forest blocks or the degree of fragmentation** | | | | **0-10** |
| Estimated size of forest block | | | | | |
| <2 hectares | N/A | 1 | | | |
| 2-5 hectares | N/A | 2 | | | |
| 5-10 hectares | N/A | 4 | | | |
| 10-20 hectares | N/A | 6 | | | |
| >20 hectares | <50% disturbed | 8 | | | |
| >20 hectares | >50% disturbed | 10 | | | |
| **Neighborhood (NH)** | **The extent of physical connectivity between areas of native vegetation within nested 100 m, 1 km, and 5 km radii** | | | | **0-5** |
| Connected w/in 100 m of plot center | If yes: | | 2.4 | | |
| Connected w/in 1 km of plot center | If yes: | | 1.6 | | |
| Connected w/in 5 km of plot center | If yes: | | 1.2 | | |
| | | Sum | | | |
| Is >50% of a 5 km radius from plot center significantly disturbed? | If yes: | | -2 | | |
| | | Sum | | | |
| **Distance to Core (D2C)** | **The distance to 50 ha core area of natural habitat** | | | | **0-5** |

| | |
|---|---|
| Distance to nearest 50 ha core area of natural habitat | |
| <5 km from center of plot | 0 |
| 1-5 km from center of plot | 2 |
| <1 km from center of plot | 4 |
| No distance | 5 |

**Table S12.**
Habitat quality metric descriptions adapted from Boggess et al. (2024) with scoring metrics for habitat variables.

| Sample ID | Taxon | Collection Date | Collector | Location | Herbarium | Latitude/ Longitude | Specimen Barcode |
|---|---|---|---|---|---|---|---|
| LEFI_WHI_ 20b | Lepraria finkii | 2022-08-16 | CAM-NSF-Lichen Genomics Team; Denzer, Eli | USA: Nantahala National Forest, summit of Whiterock Mountain, spur trail 0.3 mi N of jct w/ Bartram Trail, 1.3 mi NW of Jones Gap parking area at terminus of FS 4522 | New York State Museum | 35.085 N 83.3053 W | NYSM0000191874 |
| LEFI_WHI_ 20a | Lepraria finkii | 2022-08-16 | CAM-NSF-Lichen Genomics Team; Denzer, Eli | USA: Nantahala National Forest, summit of Whiterock Mountain, spur trail 0.3 mi N of jct w/ Bartram Trail, 1.3 mi NW of Jones Gap parking area at terminus of FS 4522 | New York State Museum | 35.085 N 83.3053 W | NYSM0000191873 |
| LEFI_WHI_ 14c | Lepraria finkii | 2022-08-16 | CAM-NSF-Lichen Genomics Team; Denzer, Eli | USA: Nantahala National Forest, summit of Whiterock Mountain, spur trail 0.3 mi N of jct w/ Bartram Trail, 1.3 mi NW of Jones Gap parking area at terminus of FS 4522 | New York State Museum | 35.085 N 83.3053 W | NYSM0000191872 |
| LEFI_WHI_ 14b | Lepraria finkii | 2022-08-16 | CAM-NSF-Lichen Genomics Team; Denzer, Eli | USA: Nantahala National Forest, summit of Whiterock Mountain, spur trail 0.3 mi N of jct w/ Bartram Trail, 1.3 mi NW of Jones Gap parking area at terminus of FS 4522 | New York State Museum | 35.085 N 83.3053 W | NYSM0000191871 |

| | | | | | | | |
|---|---|---|---|---|---|---|---|
| LEFI_WHI_ 14a | Lepraria finkii | 2022-08-16 | CAM-NSF-Lichen Genomics Team; Denzer, Eli | USA: Nantahala National Forest, summit of Whiterock Mountain, spur trail 0.3 mi N of jct w/ Bartram Trail, 1.3 mi NW of Jones Gap parking area at terminus of FS 4522 | New York State Museum | 35.085 N 83.3053 W | NYSM0000191870 |
| LEFI_WHI_ 13b | Lepraria finkii | 2022-08-16 | CAM-NSF-Lichen Genomics Team; Denzer, Eli | USA: Nantahala National Forest, summit of Whiterock Mountain, spur trail 0.3 mi N of jct w/ Bartram Trail, 1.3 mi NW of Jones Gap parking area at terminus of FS 4522 | New York State Museum | 35.085 N 83.3053 W | NYSM0000191869 |
| LEFI_WHI_ 13a | Lepraria finkii | 2022-08-16 | CAM-NSF-Lichen Genomics Team; Denzer, Eli | USA: Nantahala National Forest, summit of Whiterock Mountain, spur trail 0.3 mi N of jct w/ Bartram Trail, 1.3 mi NW of Jones Gap parking area at terminus of FS 4522 | New York State Museum | 35.085 N 83.3053 W | NYSM0000191868 |
| LEFI_WHI_ 05A | Lepraria finkii | 2022-07-30 | CAM-NSF-Lichen Genomics Team; Denzer, Eli | USA: Nantahala National Forest, summit of Whiterock Mountain, spur trail 0.3 mi N of jct w/ Bartram Trail, 1.3 mi NW of Jones Gap parking area at terminus of FS 4522 | New York State Museum | 35.085 N 83.3053 W | NYSM0000191867 |
| LEFI_WHI_ 01B | Lepraria finkii | 2022-07-30 | CAM-NSF-Lichen Genomics Team; Denzer, Eli | USA: Nantahala National Forest, summit of Whiterock Mountain, spur trail 0.3 mi N of jct w/ Bartram Trail, 1.3 mi NW of Jones Gap parking area at terminus of FS 4522 | New York State Museum | 35.085 N 83.3053 W | NYSM0000191866 |

| | | | | | | | |
|---|---|---|---|---|---|---|---|
| LEFI_TYS_ 04b | Lepraria finkii | 2022-05-03 | CAM-NSF-Lichen Genomics Team; Allen, Jessica L. | USA: Shenandoah National Park, Central District, Bettys Rock, ~0.3 mi N of Crescent Rock, upper W-facing slopes of Timber Hollow | New York State Museum | 38.56753 N 78.38223 W | NYSM0000191865 |
| LEFI_TYS_ 04a | Lepraria finkii | 2022-05-03 | CAM-NSF-Lichen Genomics Team; Allen, Jessica L. | USA: Shenandoah National Park, Central District, Bettys Rock, ~0.3 mi N of Crescent Rock, upper W-facing slopes of Timber Hollow | New York State Museum | 38.56753 N 78.38223 W | NYSM0000191864 |
| LEFI_TYS_ 03c | Lepraria finkii | 2022-05-03 | CAM-NSF-Lichen Genomics Team; Allen, Jessica L. | USA: Shenandoah National Park, Central District, Bettys Rock, ~0.3 mi N of Crescent Rock, upper W-facing slopes of Timber Hollow | New York State Museum | 38.56753 N 78.38223 W | NYSM0000191863 |
| LEFI_TYS_ 03b | Lepraria finkii | 2022-05-03 | CAM-NSF-Lichen Genomics Team; Allen, Jessica L. | USA: Shenandoah National Park, Central District, Bettys Rock, ~0.3 mi N of Crescent Rock, upper W-facing slopes of Timber Hollow | New York State Museum | 38.56753 N 78.38223 W | NYSM0000191861 |
| LEFI_TYS_ 03a | Lepraria finkii | 2022-05-03 | CAM-NSF-Lichen Genomics Team; Allen, Jessica L. | USA: Shenandoah National Park, Central District, Bettys Rock, ~0.3 mi N of Crescent Rock, upper W-facing slopes of Timber Hollow | New York State Museum | 38.56753 N 78.38223 W | NYSM0000191862 |
| LEFI_TYS_ 02c | Lepraria finkii | 2022-05-03 | CAM-NSF-Lichen Genomics Team; Allen, Jessica L. | USA: Shenandoah National Park, Central District, Bettys Rock, ~0.3 mi N of Crescent Rock, upper W-facing slopes of Timber Hollow | New York State Museum | 38.56753 N 78.38223 W | NYSM0000191860 |

| | | | | | | | |
|---|---|---|---|---|---|---|---|
| LEFI_TYS_02b | Lepraria finkii | 2022-05-03 | CAM-NSF-Lichen Genomics Team; Allen, Jessica L. | USA: Shenandoah National Park, Central District, Bettys Rock, ~0.3 mi N of Crescent Rock, upper W-facing slopes of Timber Hollow | New York State Museum | 38.56753 N 78.38223 W | NYSM0000191859 |
| LEFI_TYS_02a | Lepraria finkii | 2022-05-03 | CAM-NSF-Lichen Genomics Team; Allen, Jessica L. | USA: Shenandoah National Park, Central District, Bettys Rock, ~0.3 mi N of Crescent Rock, upper W-facing slopes of Timber Hollow | New York State Museum | 38.56753 N 78.38223 W | NYSM0000191858 |
| LEFI_TYS_01c | Lepraria finkii | 2022-05-03 | CAM-NSF-Lichen Genomics Team; Allen, Jessica L. | USA: Shenandoah National Park, Central District, Bettys Rock, ~0.3 mi N of Crescent Rock, upper W-facing slopes of Timber Hollow | New York State Museum | 38.56753 N 78.38223 W | NYSM0000191857 |
| LEFI_TYS_01b | Lepraria finkii | 2022-05-03 | CAM-NSF-Lichen Genomics Team; Allen, Jessica L. | USA: Shenandoah National Park, Central District, Bettys Rock, ~0.3 mi N of Crescent Rock, upper W-facing slopes of Timber Hollow | New York State Museum | 38.56753 N 78.38223 W | NYSM0000191856 |
| LEFI_TYS_01a | Lepraria finkii | 2022-05-03 | CAM-NSF-Lichen Genomics Team; Allen, Jessica L. | USA: Shenandoah National Park, Central District, Bettys Rock, ~0.3 mi N of Crescent Rock, upper W-facing slopes of Timber Hollow | New York State Museum | 38.56753 N 78.38223 W | NYSM0000191855 |
| LEFI_TRI_14c | Lepraria finkii | 2022-04-28 | CAM-NSF-Lichen Genomics Team; Allen, Jessica L. | USA: Monongahela National Forest, Shavers Mountain, E-facing slopes of Trillium Knob above Old Road Run, S of Glade | New York State Museum | 38.644182 N 79.83029 W | NYSM0000191854 |

| | | | | | | | |
|---|---|---|---|---|---|---|---|
| | | | | Run headwaters, Allegheny Trail ~1.5 mi N of FSR27 | | | |
| LEFI_TRI_12c | Lepraria finkii | 2022-04-28 | CAM-NSF-Lichen Genomics Team; Allen, Jessica L. | USA: Monongahela National Forest, Shavers Mountain, E-facing slopes of Trillium Knob above Old Road Run, S of Glade Run headwaters, Allegheny Trail ~1.5 mi N of FSR27 | New York State Museum | 38.644182 N 79.83029 W | NYSM0000191851 |
| LEFI_TRI_12b | Lepraria finkii | 2022-04-28 | CAM-NSF-Lichen Genomics Team; Allen, Jessica L. | USA: Monongahela National Forest, Shavers Mountain, E-facing slopes of Trillium Knob above Old Road Run, S of Glade Run headwaters, Allegheny Trail ~1.5 mi N of FSR27 | New York State Museum | 38.644182 N 79.83029 W | NYSM0000191850 |
| LEFI_TRI_12a | Lepraria finkii | 2022-04-28 | CAM-NSF-Lichen Genomics Team; Allen, Jessica L. | USA: Monongahela National Forest, Shavers Mountain, E-facing slopes of Trillium Knob above Old Road Run, S of Glade Run headwaters, Allegheny Trail ~1.5 mi N of FSR27 | New York State Museum | 38.644182 N 79.83029 W | NYSM0000191849 |
| LEFI_TRI_11a | Lepraria finkii | 2022-04-28 | CAM-NSF-Lichen Genomics Team; Allen, Jessica L. | USA: Monongahela National Forest, Shavers Mountain, E-facing slopes of Trillium Knob above Old Road Run, S of Glade Run headwaters, Allegheny Trail ~1.5 mi N of FSR27 | New York State Museum | 38.644182 N 79.83029 W | NYSM0000191848 |

| | | | | | | | |
|---|---|---|---|---|---|---|---|
| LEFI_TRI_10c | Lepraria finkii | 2022-04-28 | CAM-NSF-Lichen Genomics Team; Allen, Jessica L. | USA: Monongahela National Forest, Shavers Mountain, E-facing slopes of Trillium Knob above Old Road Run, S of Glade Run headwaters, Allegheny Trail ~1.5 mi N of FSR27 | New York State Museum | 38.644182 N 79.83029 W | NYSM0000191847 |
| LEFI_TRI_10b | Lepraria finkii | 2022-04-28 | CAM-NSF-Lichen Genomics Team; Allen, Jessica L. | USA: Monongahela National Forest, Shavers Mountain, E-facing slopes of Trillium Knob above Old Road Run, S of Glade Run headwaters, Allegheny Trail ~1.5 mi N of FSR27 | New York State Museum | 38.644182 N 79.83029 W | NYSM0000191846 |
| LEFI_TRI_10a | Lepraria finkii | 2022-04-28 | CAM-NSF-Lichen Genomics Team; Allen, Jessica L. | USA: Monongahela National Forest, Shavers Mountain, E-facing slopes of Trillium Knob above Old Road Run, S of Glade Run headwaters, Allegheny Trail ~1.5 mi N of FSR27 | New York State Museum | 38.644182 N 79.83029 W | NYSM0000191845 |
| LEFI_TOM_19a | Lepraria finkii | 2021-10-27 | CAM-NSF-Lichen Genomics Team; Allen, Jessica L. | USA: Mount Mitchell State Park, vicinity of summit of Balsam Cone, Black Mountain Crest Trail ~2 mi N of Mount Mitchell and terminus of NC128 | New York State Museum | 35.7892 N 82.2558 W | NYSM0000191844 |
| LEFI_TOM_18a | Lepraria finkii | 2021-10-27 | CAM-NSF-Lichen Genomics Team; Allen, Jessica L. | USA: Mount Mitchell State Park, vicinity of summit of Balsam Cone, Black Mountain Crest Trail ~2 mi N of Mount Mitchell and terminus of NC128 | New York State Museum | 35.7892 N 82.2558 W | NYSM0000191843 |

| | | | | | | | |
|---|---|---|---|---|---|---|---|
| LEFI_TOM_ 16b | Lepraria finkii | 2021-10-27 | CAM-NSF-Lichen Genomics Team; Allen, Jessica L. | USA: Mount Mitchell State Park, vicinity of summit of Balsam Cone, Black Mountain Crest Trail ~2 mi N of Mount Mitchell and terminus of NC128 | New York State Museum | 35.7892 N 82.2558 W | NYSM0000191842 |
| LEFI_TOM_ 16a | Lepraria finkii | 2021-10-27 | CAM-NSF-Lichen Genomics Team; Allen, Jessica L. | USA: Mount Mitchell State Park, vicinity of summit of Balsam Cone, Black Mountain Crest Trail ~2 mi N of Mount Mitchell and terminus of NC128 | New York State Museum | 35.7892 N 82.2558 W | NYSM0000191841 |
| LEFI_TOM_ 15a | Lepraria finkii | 2021-10-27 | CAM-NSF-Lichen Genomics Team; Allen, Jessica L. | USA: Mount Mitchell State Park, vicinity of summit of Balsam Cone, Black Mountain Crest Trail ~2 mi N of Mount Mitchell and terminus of NC128 | New York State Museum | 35.7892 N 82.2558 W | NYSM0000191840 |
| LEFI_TOM_ 14b | Lepraria finkii | 2021-10-27 | CAM-NSF-Lichen Genomics Team; Allen, Jessica L. | USA: Mount Mitchell State Park, vicinity of summit of Balsam Cone, Black Mountain Crest Trail ~2 mi N of Mount Mitchell and terminus of NC128 | New York State Museum | 35.7892 N 82.2558 W | NYSM0000191839 |
| LEFI_TOM_ 14a | Lepraria finkii | 2021-10-27 | CAM-NSF-Lichen Genomics Team; Allen, Jessica L. | USA: Mount Mitchell State Park, vicinity of summit of Balsam Cone, Black Mountain Crest Trail ~2 mi N of Mount Mitchell and terminus of NC128 | New York State Museum | 35.7892 N 82.2558 W | NYSM0000191838 |
| LEFI_TOM_ 09a | Lepraria finkii | 2021-10-27 | CAM-NSF-Lichen Genomics Team; Allen, Jessica L. | USA: Mount Mitchell State Park, vicinity of summit of Balsam Cone, Black Mountain Crest Trail ~2 mi N of Mount Mitchell and terminus of NC128 | New York State Museum | 35.7892 N 82.2558 W | NYSM0000191837 |

| | | | | | | | |
|---|---|---|---|---|---|---|---|
| LEFI_TMK_09b | Lepraria finkii | 2022-04-15 | CAM-NSF-Lichen Genomics Team; Allen, Jessica L. | USA: Jefferson National Forest, Mount Rogers National Recreation Area, Lewis Fork Wilderness, E-facing slopes of Pine Mountain, NW of Appalachian Trail ~0.6 E of Thomas Knob Shelter | New York State Museum | 36.655015 N 81.526364 W | NYSM0000191835 |
| LEFI_TMK_09a | Lepraria finkii | 2022-04-15 | CAM-NSF-Lichen Genomics Team; Allen, Jessica L. | USA: Jefferson National Forest, Mount Rogers National Recreation Area, Lewis Fork Wilderness, E-facing slopes of Pine Mountain, NW of Appalachian Trail ~0.6 E of Thomas Knob Shelter | New York State Museum | 36.655015 N 81.526364 W | NYSM0000191834 |
| LEFI_TMK_08a | Lepraria finkii | 2022-04-15 | CAM-NSF-Lichen Genomics Team; Allen, Jessica L. | USA: Jefferson National Forest, Mount Rogers National Recreation Area, Lewis Fork Wilderness, E-facing slopes of Pine Mountain, NW of Appalachian Trail ~0.6 E of Thomas Knob Shelter | New York State Museum | 36.655015 N 81.526364 W | NYSM0000191833 |
| LEFI_TMK_07a | Lepraria finkii | 2022-04-15 | CAM-NSF-Lichen Genomics Team; Allen, Jessica L. | USA: Jefferson National Forest, Mount Rogers National Recreation Area, Lewis Fork Wilderness, E-facing slopes of Pine Mountain, NW of Appalachian Trail ~0.6 E of Thomas Knob Shelter | New York State Museum | 36.655015 N 81.526364 W | NYSM0000191832 |

| | | | | | | | |
|---|---|---|---|---|---|---|---|
| LEFI_TMK_05a | Lepraria finkii | 2022-04-15 | CAM-NSF-Lichen Genomics Team; Allen, Jessica L. | USA: Jefferson National Forest, Mount Rogers National Recreation Area, Lewis Fork Wilderness, E-facing slopes of Pine Mountain, NW of Appalachian Trail ~0.6 E of Thomas Knob Shelter | New York State Museum | 36.655015 N 81.526364 W | NYSM0000191831 |
| LEFI_SNR_13b | Lepraria finkii | 2022-04-12 | CAM-NSF-Lichen Genomics Team; Allen, Jessica L. | USA: E-slopes of Pine Mountain, Kentenia State Forest, S of Little Shepherd Trail/SR1679 between Mill Cliff Branch and Hi Lewis Branch | New York State Museum | 36.920858 N 83.230543 W | NYSM0000194993 |
| LEFI_SNR_13a | Lepraria finkii | 2022-04-12 | CAM-NSF-Lichen Genomics Team; Allen, Jessica L. | USA: E-slopes of Pine Mountain, Kentenia State Forest, S of Little Shepherd Trail/SR1679 between Mill Cliff Branch and Hi Lewis Branch | New York State Museum | 36.920858 N 83.230543 W | NYSM0000194992 |
| LEFI_SNR_09a | Lepraria finkii | 2022-04-12 | CAM-NSF-Lichen Genomics Team; Allen, Jessica L. | USA: E-slopes of Pine Mountain, Kentenia State Forest, S of Little Shepherd Trail/SR1679 between Mill Cliff Branch and Hi Lewis Branch | New York State Museum | 36.920858 N 83.230543 W | NYSM0000194991 |
| LEFI_SNR_08b | Lepraria finkii | 2022-04-12 | CAM-NSF-Lichen Genomics Team; Allen, Jessica L. | USA: E-slopes of Pine Mountain, Kentenia State Forest, S of Little Shepherd Trail/SR1679 between Mill Cliff Branch and Hi Lewis Branch | New York State Museum | 36.920858 N 83.230543 W | NYSM0000194990 |

| | | | | | | | |
|---|---|---|---|---|---|---|---|
| LEFI_SNR_08a | Lepraria finkii | 2022-04-12 | CAM-NSF-Lichen Genomics Team; Allen, Jessica L. | USA: E-slopes of Pine Mountain, Kentenia State Forest, S of Little Shepherd Trail/SR1679 between Mill Cliff Branch and Hi Lewis Branch | New York State Museum | 36.920858 N 83.230543 W | NYSM0000194989 |
| LEFI_SNR_06b | Lepraria finkii | 2022-04-12 | CAM-NSF-Lichen Genomics Team; Allen, Jessica L. | USA: E-slopes of Pine Mountain, Kentenia State Forest, S of Little Shepherd Trail/SR1679 between Mill Cliff Branch and Hi Lewis Branch | New York State Museum | 36.920858 N 83.230543 W | NYSM0000194988 |
| LEFI_SNR_06a | Lepraria finkii | 2022-04-12 | CAM-NSF-Lichen Genomics Team; Allen, Jessica L. | USA: E-slopes of Pine Mountain, Kentenia State Forest, S of Little Shepherd Trail/SR1679 between Mill Cliff Branch and Hi Lewis Branch | New York State Museum | 36.920858 N 83.230543 W | NYSM0000194987 |
| LEFI_SNR_01d | Lepraria finkii | 2022-04-12 | CAM-NSF-Lichen Genomics Team; Allen, Jessica L. | USA: E-slopes of Pine Mountain, Kentenia State Forest, S of Little Shepherd Trail/SR1679 between Mill Cliff Branch and Hi Lewis Branch | New York State Museum | 36.920858 N 83.230543 W | NYSM0000194986 |
| LEFI_SNR_01c | Lepraria finkii | 2022-04-12 | CAM-NSF-Lichen Genomics Team; Allen, Jessica L. | USA: E-slopes of Pine Mountain, Kentenia State Forest, S of Little Shepherd Trail/SR1679 between Mill Cliff Branch and Hi Lewis Branch | New York State Museum | 36.920858 N 83.230543 W | NYSM0000194985 |
| LEFI_SNR_01a | Lepraria finkii | 2022-04-12 | CAM-NSF-Lichen Genomics Team; Allen, Jessica L. | USA: E-slopes of Pine Mountain, Kentenia State Forest, S of Little Shepherd Trail/SR1679 between Mill Cliff Branch and Hi Lewis Branch | New York State Museum | 36.920858 N 83.230543 W | NYSM0000194983 |

| | | | | | | | |
|---|---|---|---|---|---|---|---|
| LEFI_SHR_06c | Lepraria finkii | 2022-08-04 | CAM-NSF-Lichen Genomics Team; Denzer, Eli | USA: Pisgah National Forest, Balsam Mountains, Shining Rock Wilderness, Shining Rock Ledge, E slopes of Stairs Mountain summit, Art Loeb Trail 1 mi N of Shining Rock Gap | New York State Museum | 35.37817 N 82.86893 W | NYSM0000191830 |
| LEFI_SHR_06b | Lepraria finkii | 2022-08-04 | CAM-NSF-Lichen Genomics Team; Denzer, Eli | USA: Pisgah National Forest, Balsam Mountains, Shining Rock Wilderness, Shining Rock Ledge, E slopes of Stairs Mountain summit, Art Loeb Trail 1 mi N of Shining Rock Gap | New York State Museum | 35.37817 N 82.86893 W | NYSM0000191829 |
| LEFI_SHR_06a | Lepraria finkii | 2022-08-04 | CAM-NSF-Lichen Genomics Team; Denzer, Eli | USA: Pisgah National Forest, Balsam Mountains, Shining Rock Wilderness, Shining Rock Ledge, E slopes of Stairs Mountain summit, Art Loeb Trail 1 mi N of Shining Rock Gap | New York State Museum | 35.37817 N 82.86893 W | |
| LEFI_SHR_05b | Lepraria finkii | 2022-08-04 | CAM-NSF-Lichen Genomics Team; Denzer, Eli | USA: Pisgah National Forest, Balsam Mountains, Shining Rock Wilderness, Shining Rock Ledge, E slopes of Stairs Mountain summit, Art Loeb Trail 1 mi N of Shining Rock Gap | New York State Museum | 35.37817 N 82.86893 W | |
| LEFI_SHR_05a | Lepraria finkii | 2022-08-04 | CAM-NSF-Lichen Genomics Team; Denzer, Eli | USA: Pisgah National Forest, Balsam Mountains, Shining Rock Wilderness, Shining Rock Ledge, E slopes of Stairs Mountain summit, Art Loeb Trail 1 mi N of Shining Rock Gap | New York State Museum | 35.37817 N 82.86893 W | NYSM0000191828 |

| | | | | | | | |
|---|---|---|---|---|---|---|---|
| LEFI_RSK_07c | Lepraria finkii | 2022-04-26 | CAM-NSF-Lichen Genomics Team; Allen, Jessica L. | USA: Monongahela National Forest, S-facing slopes of Red Spruce Knob, N of Red Lick Mountain, 0.1 mi W of WV150 and FSR451 | New York State Museum | 38.323665 N 80.152203 W | NYSM0000191827 |
| LEFI_RSK_07b | Lepraria finkii | 2022-04-26 | CAM-NSF-Lichen Genomics Team; Allen, Jessica L. | USA: Monongahela National Forest, S-facing slopes of Red Spruce Knob, N of Red Lick Mountain, 0.1 mi W of WV150 and FSR451 | New York State Museum | 38.323665 N 80.152203 W | NYSM0000191826 |
| LEFI_RSK_06c | Lepraria finkii | 2022-04-26 | CAM-NSF-Lichen Genomics Team; Allen, Jessica L. | USA: Monongahela National Forest, S-facing slopes of Red Spruce Knob, N of Red Lick Mountain, 0.1 mi W of WV150 and FSR451 | New York State Museum | 38.323665 N 80.152203 W | NYSM0000191825 |
| LEFI_RSK_06b | Lepraria finkii | 2022-04-26 | CAM-NSF-Lichen Genomics Team; Allen, Jessica L. | USA: Monongahela National Forest, S-facing slopes of Red Spruce Knob, N of Red Lick Mountain, 0.1 mi W of WV150 and FSR451 | New York State Museum | 38.323665 N 80.152203 W | NYSM0000191824 |
| LEFI_RSK_06a | Lepraria finkii | 2022-04-26 | CAM-NSF-Lichen Genomics Team; Allen, Jessica L. | USA: Monongahela National Forest, S-facing slopes of Red Spruce Knob, N of Red Lick Mountain, 0.1 mi W of WV150 and FSR451 | New York State Museum | 38.323665 N 80.152203 W | NYSM0000191823 |
| LEFI_RSK_05c | Lepraria finkii | 2022-04-26 | CAM-NSF-Lichen Genomics Team; Allen, Jessica L. | USA: Monongahela National Forest, S-facing slopes of Red Spruce Knob, N of Red Lick Mountain, 0.1 mi W of WV150 and FSR451 | New York State Museum | 38.323665 N 80.152203 W | NYSM0000191822 |

| | | | | | | | |
|---|---|---|---|---|---|---|---|
| LEFI_RSK_05b | Lepraria finkii | 2022-04-26 | CAM-NSF-Lichen Genomics Team; Allen, Jessica L. | USA: Monongahela National Forest, S-facing slopes of Red Spruce Knob, N of Red Lick Mountain, 0.1 mi W of WV150 and FSR451 | New York State Museum | 38.323665 N 80.152203 W | NYSM0000191821 |
| LEFI_RSK_05a | Lepraria finkii | 2022-04-26 | CAM-NSF-Lichen Genomics Team; Allen, Jessica L. | USA: Monongahela National Forest, S-facing slopes of Red Spruce Knob, N of Red Lick Mountain, 0.1 mi W of WV150 and FSR451 | New York State Museum | 38.323665 N 80.152203 W | NYSM0000191820 |
| LEFI_RSK_03c | Lepraria finkii | 2022-04-26 | CAM-NSF-Lichen Genomics Team; Allen, Jessica L. | USA: Monongahela National Forest, S-facing slopes of Red Spruce Knob, N of Red Lick Mountain, 0.1 mi W of WV150 and FSR451 | New York State Museum | 38.323665 N 80.152203 W | NYSM0000191819 |
| LEFI_RSK_03b | Lepraria finkii | 2022-04-26 | CAM-NSF-Lichen Genomics Team; Allen, Jessica L. | USA: Monongahela National Forest, S-facing slopes of Red Spruce Knob, N of Red Lick Mountain, 0.1 mi W of WV150 and FSR451 | New York State Museum | 38.323665 N 80.152203 W | NYSM0000191818 |
| LEFI_RSK_03a | Lepraria finkii | 2022-04-26 | CAM-NSF-Lichen Genomics Team; Allen, Jessica L. | USA: Monongahela National Forest, S-facing slopes of Red Spruce Knob, N of Red Lick Mountain, 0.1 mi W of WV150 and FSR451 | New York State Museum | 38.323665 N 80.152203 W | NYSM0000191817 |
| LEFI_RON_04a | Lepraria finkii | 2022-04-19 | CAM-NSF-Lichen Genomics Team; Allen, Jessica L. | USA: Jefferson National Forest, Round Mountain, vicinity of summit, Round Mountain Rd./VA688, ~4 mi W of US52/Bastian | New York State Museum | 37.154716 N 81.194163 W | NYSM0000191816 |

| | | | | | | | |
|---|---|---|---|---|---|---|---|
| LEFI_RON_ 03c | Lepraria finkii | 2022-04-19 | CAM-NSF-Lichen Genomics Team; Allen, Jessica L. | USA: Jefferson National Forest, Round Mountain, vicinity of summit, Round Mountain Rd./VA688, ~4 mi W of US52/Bastian | New York State Museum | 37.154716 N 81.194163 W | NYSM0000191815 |
| LEFI_RON_ 03b | Lepraria finkii | 2022-04-19 | CAM-NSF-Lichen Genomics Team; Allen, Jessica L. | USA: Jefferson National Forest, Round Mountain, vicinity of summit, Round Mountain Rd./VA688, ~4 mi W of US52/Bastian | New York State Museum | 37.154716 N 81.194163 W | NYSM0000191814 |
| LEFI_RON_ 03a | Lepraria finkii | 2022-04-19 | CAM-NSF-Lichen Genomics Team; Allen, Jessica L. | USA: Jefferson National Forest, Round Mountain, vicinity of summit, Round Mountain Rd./VA688, ~4 mi W of US52/Bastian | New York State Museum | 37.154716 N 81.194163 W | NYSM0000191813 |
| LEFI_RON_ 02c | Lepraria finkii | 2022-04-19 | CAM-NSF-Lichen Genomics Team; Allen, Jessica L. | USA: Jefferson National Forest, Round Mountain, vicinity of summit, Round Mountain Rd./VA688, ~4 mi W of US52/Bastian | New York State Museum | 37.154716 N 81.194163 W | NYSM0000191812 |
| LEFI_RON_ 02b | Lepraria finkii | 2022-04-19 | CAM-NSF-Lichen Genomics Team; Allen, Jessica L. | USA: Jefferson National Forest, Round Mountain, vicinity of summit, Round Mountain Rd./VA688, ~4 mi W of US52/Bastian | New York State Museum | 37.154716 N 81.194163 W | NYSM0000191811 |
| LEFI_RON_ 02a | Lepraria finkii | 2022-04-19 | CAM-NSF-Lichen Genomics Team; Allen, Jessica L. | USA: Jefferson National Forest, Round Mountain, vicinity of summit, Round Mountain Rd./VA688, ~4 mi W of US52/Bastian | New York State Museum | 37.154716 N 81.194163 W | NYSM0000191810 |

| | | | | | | | |
|---|---|---|---|---|---|---|---|
| LEFI_RON_ 01c | Lepraria finkii | 2022-04-19 | CAM-NSF-Lichen Genomics Team; Allen, Jessica L. | USA: Jefferson National Forest, Round Mountain, vicinity of summit, Round Mountain Rd./VA688, ~4 mi W of US52/Bastian | New York State Museum | 37.154716 N 81.194163 W | NYSM0000191809 |
| LEFI_RON_ 01b | Lepraria finkii | 2022-04-19 | CAM-NSF-Lichen Genomics Team; Allen, Jessica L. | USA: Jefferson National Forest, Round Mountain, vicinity of summit, Round Mountain Rd./VA688, ~4 mi W of US52/Bastian | New York State Museum | 37.154716 N 81.194163 W | NYSM0000191808 |
| LEFI_RON_ 01a | Lepraria finkii | 2022-04-19 | CAM-NSF-Lichen Genomics Team; Allen, Jessica L. | USA: Jefferson National Forest, Round Mountain, vicinity of summit, Round Mountain Rd./VA688, ~4 mi W of US52/Bastian | New York State Museum | 37.154716 N 81.194163 W | NYSM0000191807 |
| LEFI_ROG_ 05b | Lepraria finkii | 2022-04-16 | CAM-NSF-Lichen Genomics Team; Allen, Jessica L. | USA: Jefferson National Forest, Mount Rogers National Recreation Area, Lewis Fork Wilderness, S-facing slopes of Mount Rogers, Mount Rogers Spur Trail 0.25 mi E of summit, ~0.25 mi W of jct w/ Appalachian Trail | New York State Museum | 36.659449 N 81.540821 W | NYSM0000191806 |
| LEFI_ROG_ 05a | Lepraria finkii | 2022-04-16 | CAM-NSF-Lichen Genomics Team; Allen, Jessica L. | USA: Jefferson National Forest, Mount Rogers National Recreation Area, Lewis Fork Wilderness, S-facing slopes of Mount Rogers, Mount Rogers Spur Trail 0.25 mi E of summit, ~0.25 mi W of jct w/ Appalachian Trail | New York State Museum | 36.659449 N 81.540821 W | NYSM0000191805 |

| | | | | | | | |
|---|---|---|---|---|---|---|---|
| LEFI_ROG_02c03a | Lepraria finkii | 2022-04-16 | CAM-NSF-Lichen Genomics Team; Allen, Jessica L. | USA: Jefferson National Forest, Mount Rogers National Recreation Area, Lewis Fork Wilderness, S-facing slopes of Mount Rogers, Mount Rogers Spur Trail 0.25 mi E of summit, ~0.25 mi W of jct w/ Appalachian Trail | New York State Museum | 36.659449 N 81.540821 W | NYSM0000191803 |
| LEFI_ROG_02b | Lepraria finkii | 2022-04-16 | CAM-NSF-Lichen Genomics Team; Allen, Jessica L. | USA: Jefferson National Forest, Mount Rogers National Recreation Area, Lewis Fork Wilderness, S-facing slopes of Mount Rogers, Mount Rogers Spur Trail 0.25 mi E of summit, ~0.25 mi W of jct w/ Appalachian Trail | New York State Museum | 36.659449 N 81.540821 W | NYSM0000191802 |
| LEFI_ROG_02a | Lepraria finkii | 2022-04-16 | CAM-NSF-Lichen Genomics Team; Allen, Jessica L. | USA: Jefferson National Forest, Mount Rogers National Recreation Area, Lewis Fork Wilderness, S-facing slopes of Mount Rogers, Mount Rogers Spur Trail 0.25 mi E of summit, ~0.25 mi W of jct w/ Appalachian Trail | New York State Museum | 36.659449 N 81.540821 W | NYSM0000191801 |
| LEFI_ROG_01a | Lepraria finkii | 2022-04-16 | CAM-NSF-Lichen Genomics Team; Allen, Jessica L. | USA: Jefferson National Forest, Mount Rogers National Recreation Area, Lewis Fork Wilderness, S-facing slopes of Mount Rogers, Mount Rogers Spur Trail 0.25 mi E of summit, ~0.25 mi W of jct w/ Appalachian Trail | New York State Museum | 36.659449 N 81.540821 W | NYSM0000191800 |

| | | | | | | | |
|---|---|---|---|---|---|---|---|
| LEFI_POK_05a | Lepraria finkii | 2022-04-24 | CAM-NSF-Lichen Genomics Team; Allen, Jessica L. | USA: Monongahela National Forest, W-facing slopes of Canaan Mountain, E of Pointy Knob, Pointy Knob Trail ~2.5 mi S of FSR13 | New York State Museum | 39.035283 N 79.489141 W | NYSM0000194982 |
| LEFI_POK_04c | Lepraria finkii | 2022-04-24 | CAM-NSF-Lichen Genomics Team; Allen, Jessica L. | USA: Monongahela National Forest, W-facing slopes of Canaan Mountain, E of Pointy Knob, Pointy Knob Trail ~2.5 mi S of FSR13 | New York State Museum | 39.035283 N 79.489141 W | NYSM0000194981 |
| LEFI_POK_04b | Lepraria finkii | 2022-04-24 | CAM-NSF-Lichen Genomics Team; Allen, Jessica L. | USA: Monongahela National Forest, W-facing slopes of Canaan Mountain, E of Pointy Knob, Pointy Knob Trail ~2.5 mi S of FSR13 | New York State Museum | 39.035283 N 79.489141 W | NYSM0000194980 |
| LEFI_POK_04a | Lepraria finkii | 2022-04-24 | CAM-NSF-Lichen Genomics Team; Allen, Jessica L. | USA: Monongahela National Forest, W-facing slopes of Canaan Mountain, E of Pointy Knob, Pointy Knob Trail ~2.5 mi S of FSR13 | New York State Museum | 39.035283 N 79.489141 W | NYSM0000194979 |
| LEFI_POK_02c | Lepraria finkii | 2022-04-24 | CAM-NSF-Lichen Genomics Team; Allen, Jessica L. | USA: Monongahela National Forest, W-facing slopes of Canaan Mountain, E of Pointy Knob, Pointy Knob Trail ~2.5 mi S of FSR13 | New York State Museum | 39.035283 N 79.489141 W | NYSM0000194978 |
| LEFI_POK_02b | Lepraria finkii | 2022-04-24 | CAM-NSF-Lichen Genomics Team; Allen, Jessica L. | USA: Monongahela National Forest, W-facing slopes of Canaan Mountain, E of Pointy Knob, Pointy Knob Trail ~2.5 mi S of FSR13 | New York State Museum | 39.035283 N 79.489141 W | NYSM0000194977 |

| | | | | | | | |
|---|---|---|---|---|---|---|---|
| LEFI_POK_02a | Lepraria finkii | 2022-04-24 | CAM-NSF-Lichen Genomics Team; Allen, Jessica L. | USA: Monongahela National Forest, W-facing slopes of Canaan Mountain, E of Pointy Knob, Pointy Knob Trail ~2.5 mi S of FSR13 | New York State Museum | 39.035283 N 79.489141 W | NYSM0000194976 |
| LEFI_POK_01c | Lepraria finkii | 2022-04-24 | CAM-NSF-Lichen Genomics Team; Allen, Jessica L. | USA: Monongahela National Forest, W-facing slopes of Canaan Mountain, E of Pointy Knob, Pointy Knob Trail ~2.5 mi S of FSR13 | New York State Museum | 39.035283 N 79.489141 W | NYSM0000194975 |
| LEFI_POK_01b | Lepraria finkii | 2022-04-24 | CAM-NSF-Lichen Genomics Team; Allen, Jessica L. | USA: Monongahela National Forest, W-facing slopes of Canaan Mountain, E of Pointy Knob, Pointy Knob Trail ~2.5 mi S of FSR13 | New York State Museum | 39.035283 N 79.489141 W | NYSM0000194974 |
| LEFI_POK_01a | Lepraria finkii | 2022-04-24 | CAM-NSF-Lichen Genomics Team; Allen, Jessica L. | USA: Monongahela National Forest, W-facing slopes of Canaan Mountain, E of Pointy Knob, Pointy Knob Trail ~2.5 mi S of FSR13 | New York State Museum | 39.035283 N 79.489141 W | NYSM0000194973 |
| LEFI_NGA_05a | Lepraria finkii | 2022-04-28 | CAM-NSF-Lichen Genomics Team; Allen, Jessica L. | USA: Monongahela National Forest, Shavers Mountain, E-facing slopes of Gaudineer Knob, FSR27 ~1.25 mi N of jct w/ US250 | New York State Museum | 38.611691 N 79.841629 W | NYSM0000191799 |
| LEFI_NGA_04c | Lepraria finkii | 2022-04-28 | CAM-NSF-Lichen Genomics Team; Allen, Jessica L. | USA: Monongahela National Forest, Shavers Mountain, E-facing slopes of Gaudineer Knob, FSR27 ~1.25 mi N of jct w/ US250 | New York State Museum | 38.611691 N 79.841629 W | NYSM0000191798 |

| | | | | | | | |
|---|---|---|---|---|---|---|---|
| LEFI_NGA_04a | Lepraria finkii | 2022-04-28 | CAM-NSF-Lichen Genomics Team; Allen, Jessica L. | USA: Monongahela National Forest, Shavers Mountain, E-facing slopes of Gaudineer Knob, FSR27 ~1.25 mi N of jct w/ US250 | New York State Museum | 38.611691 N 79.841629 W | NYSM0000191796 |
| LEFI_NGA_03c | Lepraria finkii | 2022-04-28 | CAM-NSF-Lichen Genomics Team; Allen, Jessica L. | USA: Monongahela National Forest, Shavers Mountain, E-facing slopes of Gaudineer Knob, FSR27 ~1.25 mi N of jct w/ US250 | New York State Museum | 38.611691 N 79.841629 W | NYSM0000191795 |
| LEFI_NGA_03b | Lepraria finkii | 2022-04-28 | CAM-NSF-Lichen Genomics Team; Allen, Jessica L. | USA: Monongahela National Forest, Shavers Mountain, E-facing slopes of Gaudineer Knob, FSR27 ~1.25 mi N of jct w/ US250 | New York State Museum | 38.611691 N 79.841629 W | NYSM0000191794 |
| LEFI_NGA_03a | Lepraria finkii | 2022-04-28 | CAM-NSF-Lichen Genomics Team; Allen, Jessica L. | USA: Monongahela National Forest, Shavers Mountain, E-facing slopes of Gaudineer Knob, FSR27 ~1.25 mi N of jct w/ US250 | New York State Museum | 38.611691 N 79.841629 W | |
| LEFI_NGA_02c | Lepraria finkii | 2022-04-28 | CAM-NSF-Lichen Genomics Team; Allen, Jessica L. | USA: Monongahela National Forest, Shavers Mountain, E-facing slopes of Gaudineer Knob, FSR27 ~1.25 mi N of jct w/ US250 | New York State Museum | 38.611691 N 79.841629 W | NYSM0000191793 |
| LEFI_NGA_02b | Lepraria finkii | 2022-04-28 | CAM-NSF-Lichen Genomics Team; Allen, Jessica L. | USA: Monongahela National Forest, Shavers Mountain, E-facing slopes of Gaudineer Knob, FSR27 ~1.25 mi N of jct w/ US250 | New York State Museum | 38.611691 N 79.841629 W | NYSM0000191792 |

| | | | | | | | |
|---|---|---|---|---|---|---|---|
| LEFI_NGA_ 02a | Lepraria finkii | 2022-04-28 | CAM-NSF-Lichen Genomics Team; Allen, Jessica L. | USA: Monongahela National Forest, Shavers Mountain, E-facing slopes of Gaudineer Knob, FSR27 ~1.25 mi N of jct w/ US250 | New York State Museum | 38.611691 N 79.841629 W | NYSM0000191791 |
| LEFI_NGA_ 01c | Lepraria finkii | 2022-04-28 | CAM-NSF-Lichen Genomics Team; Allen, Jessica L. | USA: Monongahela National Forest, Shavers Mountain, E-facing slopes of Gaudineer Knob, FSR27 ~1.25 mi N of jct w/ US250 | New York State Museum | 38.611691 N 79.841629 W | NYSM0000191790 |
| LEFI_NGA_ 01b | Lepraria finkii | 2022-04-28 | CAM-NSF-Lichen Genomics Team; Allen, Jessica L. | USA: Monongahela National Forest, Shavers Mountain, E-facing slopes of Gaudineer Knob, FSR27 ~1.25 mi N of jct w/ US250 | New York State Museum | 38.611691 N 79.841629 W | NYSM0000191789 |
| LEFI_NGA_ 01a | Lepraria finkii | 2022-04-28 | CAM-NSF-Lichen Genomics Team; Allen, Jessica L. | USA: Monongahela National Forest, Shavers Mountain, E-facing slopes of Gaudineer Knob, FSR27 ~1.25 mi N of jct w/ US250 | New York State Museum | 38.611691 N 79.841629 W | NYSM0000191788 |
| LEFI_LUE_ 09a | Lepraria finkii | 2022-04-26 | CAM-NSF-Lichen Genomics Team; Allen, Jessica L. | USA: Monongahela National Forest, Spruce Mountain, W-facing slopes below summit of Blue Knob, ~1 mi S of WV39 | New York State Museum | 38.175865 N 80.279981 W | NYSM0000191787 |
| LEFI_LUE_ 08a | Lepraria finkii | 2022-04-26 | CAM-NSF-Lichen Genomics Team; Allen, Jessica L. | USA: Monongahela National Forest, Spruce Mountain, W-facing slopes below summit of Blue Knob, ~1 mi S of WV39 | New York State Museum | 38.175865 N 80.279981 W | NYSM0000191786 |

| | | | | | | | |
|---|---|---|---|---|---|---|---|
| LEFI_LUE_ 07b | Lepraria finkii | 2022-04-26 | CAM-NSF-Lichen Genomics Team; Allen, Jessica L. | USA: Monongahela National Forest, Spruce Mountain, W-facing slopes below summit of Blue Knob, ~1 mi S of WV39 | New York State Museum | 38.175865 N 80.279981 W | NYSM0000191785 |
| LEFI_LUE_ 07a | Lepraria finkii | 2022-04-26 | CAM-NSF-Lichen Genomics Team; Allen, Jessica L. | USA: Monongahela National Forest, Spruce Mountain, W-facing slopes below summit of Blue Knob, ~1 mi S of WV39 | New York State Museum | 38.175865 N 80.279981 W | NYSM0000191784 |
| LEFI_LUE_ 06b | Lepraria finkii | 2022-04-26 | CAM-NSF-Lichen Genomics Team; Allen, Jessica L. | USA: Monongahela National Forest, Spruce Mountain, W-facing slopes below summit of Blue Knob, ~1 mi S of WV39 | New York State Museum | 38.175865 N 80.279981 W | NYSM0000191783 |
| LEFI_LUE_ 05a | Lepraria finkii | 2022-04-26 | CAM-NSF-Lichen Genomics Team; Allen, Jessica L. | USA: Monongahela National Forest, Spruce Mountain, W-facing slopes below summit of Blue Knob, ~1 mi S of WV39 | New York State Museum | 38.175865 N 80.279981 W | NYSM0000191782 |
| LEFI_LUE_ 03c | Lepraria finkii | 2022-04-26 | CAM-NSF-Lichen Genomics Team; Allen, Jessica L. | USA: Monongahela National Forest, Spruce Mountain, W-facing slopes below summit of Blue Knob, ~1 mi S of WV39 | New York State Museum | 38.175865 N 80.279981 W | NYSM0000191780 |
| LEFI_LUE_ 03b | Lepraria finkii | 2022-04-26 | CAM-NSF-Lichen Genomics Team; Allen, Jessica L. | USA: Monongahela National Forest, Spruce Mountain, W-facing slopes below summit of Blue Knob, ~1 mi S of WV39 | New York State Museum | 38.175865 N 80.279981 W | NYSM0000191779 |

| | | | | | | | |
|---|---|---|---|---|---|---|---|
| LEFI_LUE_03a | Lepraria finkii | 2022-04-26 | CAM-NSF-Lichen Genomics Team; Allen, Jessica L. | USA: Monongahela National Forest, Spruce Mountain, W-facing slopes below summit of Blue Knob, ~1 mi S of WV39 | New York State Museum | 38.175865 N 80.279981 W | NYSM0000191778 |
| LEFI_LEM_07b | Lepraria finkii | 2022-04-13 | CAM-NSF-Lichen Genomics Team; Allen, Jessica L. | USA: Pine Mountain, Bad Branch Nature Preserve, W headwaters of Bad Branch, Pine Mountain Trail ~2.2 mi E of US119 | New York State Museum | 37.09032 N 82.77928 W | NYSM0000191777 |
| LEFI_LEM_07a | Lepraria finkii | 2022-04-13 | CAM-NSF-Lichen Genomics Team; Allen, Jessica L. | USA: Pine Mountain, Bad Branch Nature Preserve, W headwaters of Bad Branch, Pine Mountain Trail ~2.2 mi E of US119 | New York State Museum | 37.09032 N 82.77928 W | NYSM0000191776 |
| LEFI_LEM_06c | Lepraria finkii | 2022-04-13 | CAM-NSF-Lichen Genomics Team; Allen, Jessica L. | USA: Pine Mountain, Bad Branch Nature Preserve, W headwaters of Bad Branch, Pine Mountain Trail ~2.2 mi E of US119 | New York State Museum | 37.09032 N 82.77928 W | NYSM0000191775 |
| LEFI_LEM_06b | Lepraria finkii | 2022-04-13 | CAM-NSF-Lichen Genomics Team; Allen, Jessica L. | USA: Pine Mountain, Bad Branch Nature Preserve, W headwaters of Bad Branch, Pine Mountain Trail ~2.2 mi E of US119 | New York State Museum | 37.09032 N 82.77928 W | NYSM0000191774 |
| LEFI_LEM_04a | Lepraria finkii | 2022-04-13 | CAM-NSF-Lichen Genomics Team; Allen, Jessica L. | USA: Pine Mountain, Bad Branch Nature Preserve, W headwaters of Bad Branch, Pine Mountain Trail ~2.2 mi E of US119 | New York State Museum | 37.09032 N 82.77928 W | NYSM0000191772 |

| | | | | | | | |
|---|---|---|---|---|---|---|---|
| LEFI_LEM_03b | Lepraria finkii | 2022-04-13 | CAM-NSF-Lichen Genomics Team; Allen, Jessica L. | USA: Pine Mountain, Bad Branch Nature Preserve, W headwaters of Bad Branch, Pine Mountain Trail ~2.2 mi E of US119 | New York State Museum | 37.09032 N 82.77928 W | NYSM0000191771 |
| LEFI_LEM_03a | Lepraria finkii | 2022-04-13 | CAM-NSF-Lichen Genomics Team; Allen, Jessica L. | USA: Pine Mountain, Bad Branch Nature Preserve, W headwaters of Bad Branch, Pine Mountain Trail ~2.2 mi E of US119 | New York State Museum | 37.09032 N 82.77928 W | NYSM0000191770 |
| LEFI_LEM_02b | Lepraria finkii | 2022-04-13 | CAM-NSF-Lichen Genomics Team; Allen, Jessica L. | USA: Pine Mountain, Bad Branch Nature Preserve, W headwaters of Bad Branch, Pine Mountain Trail ~2.2 mi E of US119 | New York State Museum | 37.09032 N 82.77928 W | NYSM0000191769 |
| LEFI_LEM_02a | Lepraria finkii | 2022-04-13 | CAM-NSF-Lichen Genomics Team; Allen, Jessica L. | USA: Pine Mountain, Bad Branch Nature Preserve, W headwaters of Bad Branch, Pine Mountain Trail ~2.2 mi E of US119 | New York State Museum | 37.09032 N 82.77928 W | NYSM0000191768 |
| LEFI_KLE_07c | Lepraria finkii | 2022-04-24 | CAM-NSF-Lichen Genomics Team; Allen, Jessica L. | USA: Monongahela National Forest, Bickle Knob, saddle and W-facing slopes below observation tower, terminus of FSR91A | New York State Museum | 38.93557 N 79.730504 W | NYSM0000191767 |
| LEFI_KLE_07b | Lepraria finkii | 2022-04-24 | CAM-NSF-Lichen Genomics Team; Allen, Jessica L. | USA: Monongahela National Forest, Bickle Knob, saddle and W-facing slopes below observation tower, terminus of FSR91A | New York State Museum | 38.93557 N 79.730504 W | NYSM0000191766 |

| | | | | | | | |
|---|---|---|---|---|---|---|---|
| LEFI_KLE_07a | Lepraria finkii | 2022-04-24 | CAM-NSF-Lichen Genomics Team; Allen, Jessica L. | USA: Monongahela National Forest, Bickle Knob, saddle and W-facing slopes below observation tower, terminus of FSR91A | New York State Museum | 38.93557 N 79.730504 W | NYSM0000191765 |
| LEFI_KLE_06c | Lepraria finkii | 2022-04-24 | CAM-NSF-Lichen Genomics Team; Allen, Jessica L. | USA: Monongahela National Forest, Bickle Knob, saddle and W-facing slopes below observation tower, terminus of FSR91A | New York State Museum | 38.93557 N 79.730504 W | NYSM0000191764 |
| LEFI_KLE_06b | Lepraria finkii | 2022-04-24 | CAM-NSF-Lichen Genomics Team; Allen, Jessica L. | USA: Monongahela National Forest, Bickle Knob, saddle and W-facing slopes below observation tower, terminus of FSR91A | New York State Museum | 38.93557 N 79.730504 W | NYSM0000191763 |
| LEFI_KLE_06a | Lepraria finkii | 2022-04-24 | CAM-NSF-Lichen Genomics Team; Allen, Jessica L. | USA: Monongahela National Forest, Bickle Knob, saddle and W-facing slopes below observation tower, terminus of FSR91A | New York State Museum | 38.93557 N 79.730504 W | NYSM0000191762 |
| LEFI_KLE_02c | Lepraria finkii | 2022-04-24 | CAM-NSF-Lichen Genomics Team; Allen, Jessica L. | USA: Monongahela National Forest, Bickle Knob, saddle and W-facing slopes below observation tower, terminus of FSR91A | New York State Museum | 38.93557 N 79.730504 W | NYSM0000191761 |
| LEFI_KLE_02b | Lepraria finkii | 2022-04-24 | CAM-NSF-Lichen Genomics Team; Allen, Jessica L. | USA: Monongahela National Forest, Bickle Knob, saddle and W-facing slopes below observation tower, terminus of FSR91A | New York State Museum | 38.93557 N 79.730504 W | NYSM0000191760 |

| | | | | | | | |
|---|---|---|---|---|---|---|---|
| LEFI_KLE_02a | Lepraria finkii | 2022-04-24 | CAM-NSF-Lichen Genomics Team; Allen, Jessica L. | USA: Monongahela National Forest, Bickle Knob, saddle and W-facing slopes below observation tower, terminus of FSR91A | New York State Museum | 38.93557 N 79.730504 W | NYSM0000191759 |
| LEFI_JON_09c | Lepraria finkii | 2022-04-19 | CAM-NSF-Lichen Genomics Team; Allen, Jessica L. | USA: Jefferson National Forest, E slopes of Round Mountain, S of Round Mountain Rd./VA688, ~1.5 mi W of US52/Bastian | New York State Museum | 37.170518 N 81.164224 W | NYSM0000191758 |
| LEFI_JON_09b | Lepraria finkii | 2022-04-19 | CAM-NSF-Lichen Genomics Team; Allen, Jessica L. | USA: Jefferson National Forest, E slopes of Round Mountain, S of Round Mountain Rd./VA688, ~1.5 mi W of US52/Bastian | New York State Museum | 37.170518 N 81.164224 W | NYSM0000191757 |
| LEFI_JON_09a | Lepraria finkii | 2022-04-19 | CAM-NSF-Lichen Genomics Team; Allen, Jessica L. | USA: Jefferson National Forest, E slopes of Round Mountain, S of Round Mountain Rd./VA688, ~1.5 mi W of US52/Bastian | New York State Museum | 37.170518 N 81.164224 W | NYSM0000191756 |
| LEFI_JON_08b | Lepraria finkii | 2022-04-19 | CAM-NSF-Lichen Genomics Team; Allen, Jessica L. | USA: Jefferson National Forest, E slopes of Round Mountain, S of Round Mountain Rd./VA688, ~1.5 mi W of US52/Bastian | New York State Museum | 37.170518 N 81.164224 W | NYSM0000191755 |
| LEFI_JON_08a | Lepraria finkii | 2022-04-19 | CAM-NSF-Lichen Genomics Team; Allen, Jessica L. | USA: Jefferson National Forest, E slopes of Round Mountain, S of Round Mountain Rd./VA688, ~1.5 mi W of US52/Bastian | New York State Museum | 37.170518 N 81.164224 W | NYSM0000191754 |

| | | | | | | | |
|---|---|---|---|---|---|---|---|
| LEFI_JON_07a | Lepraria finkii | 2022-04-19 | CAM-NSF-Lichen Genomics Team; Allen, Jessica L. | USA: Jefferson National Forest, E slopes of Round Mountain, S of Round Mountain Rd./VA688, ~1.5 mi W of US52/Bastian | New York State Museum | 37.170518 N 81.164224 W | NYSM0000191753 |
| LEFI_JON_06b | Lepraria finkii | 2022-04-19 | CAM-NSF-Lichen Genomics Team; Allen, Jessica L. | USA: Jefferson National Forest, E slopes of Round Mountain, S of Round Mountain Rd./VA688, ~1.5 mi W of US52/Bastian | New York State Museum | 37.170518 N 81.164224 W | NYSM0000191752 |
| LEFI_JON_06a | Lepraria finkii | 2022-04-19 | CAM-NSF-Lichen Genomics Team; Allen, Jessica L. | USA: Jefferson National Forest, E slopes of Round Mountain, S of Round Mountain Rd./VA688, ~1.5 mi W of US52/Bastian | New York State Museum | 37.170518 N 81.164224 W | NYSM0000191751 |
| LEFI_JON_04c | Lepraria finkii | 2022-04-19 | CAM-NSF-Lichen Genomics Team; Allen, Jessica L. | USA: Jefferson National Forest, E slopes of Round Mountain, S of Round Mountain Rd./VA688, ~1.5 mi W of US52/Bastian | New York State Museum | 37.170518 N 81.164224 W | NYSM0000191750 |
| LEFI_JON_04b | Lepraria finkii | 2022-04-19 | CAM-NSF-Lichen Genomics Team; Allen, Jessica L. | USA: Jefferson National Forest, E slopes of Round Mountain, S of Round Mountain Rd./VA688, ~1.5 mi W of US52/Bastian | New York State Museum | 37.170518 N 81.164224 W | NYSM0000191749 |
| LEFI_JON_04a | Lepraria finkii | 2022-04-19 | CAM-NSF-Lichen Genomics Team; Allen, Jessica L. | USA: Jefferson National Forest, E slopes of Round Mountain, S of Round Mountain Rd./VA688, ~1.5 mi W of US52/Bastian | New York State Museum | 37.170518 N 81.164224 W | NYSM0000191748 |

| | | | | | | | |
|---|---|---|---|---|---|---|---|
| LEFI_HAW _05b | Lepraria finkii | 2022-05-03 | CAM-NSF-Lichen Genomics Team; Allen, Jessica L. | USA: Shenandoah National Park, Central District, SW slopes of Hawksbill Mountain, between Lower Hawksbill Trail and Hawksbill Fire Road ~0.7 mi W of Hawksbill Gap | New York State Museum | 38.55397 N 78.39687 W | NYSM0000191747 |
| LEFI_HAW _05a | Lepraria finkii | 2022-05-03 | CAM-NSF-Lichen Genomics Team; Allen, Jessica L. | USA: Shenandoah National Park, Central District, SW slopes of Hawksbill Mountain, between Lower Hawksbill Trail and Hawksbill Fire Road ~0.7 mi W of Hawksbill Gap | New York State Museum | 38.55397 N 78.39687 W | NYSM0000191746 |
| LEFI_HAW _04c | Lepraria finkii | 2022-05-03 | CAM-NSF-Lichen Genomics Team; Allen, Jessica L. | USA: Shenandoah National Park, Central District, SW slopes of Hawksbill Mountain, between Lower Hawksbill Trail and Hawksbill Fire Road ~0.7 mi W of Hawksbill Gap | New York State Museum | 38.55397 N 78.39687 W | NYSM0000191745 |
| LEFI_HAW _04b | Lepraria finkii | 2022-05-03 | CAM-NSF-Lichen Genomics Team; Allen, Jessica L. | USA: Shenandoah National Park, Central District, SW slopes of Hawksbill Mountain, between Lower Hawksbill Trail and Hawksbill Fire Road ~0.7 mi W of Hawksbill Gap | New York State Museum | 38.55397 N 78.39687 W | NYSM0000191744 |
| LEFI_HAW _04a | Lepraria finkii | 2022-05-03 | CAM-NSF-Lichen Genomics Team; Allen, Jessica L. | USA: Shenandoah National Park, Central District, SW slopes of Hawksbill Mountain, between Lower Hawksbill Trail and Hawksbill Fire Road ~0.7 mi W of Hawksbill Gap | New York State Museum | 38.55397 N 78.39687 W | NYSM0000191743 |

| | | | | | | | |
|---|---|---|---|---|---|---|---|
| LEFI_HAW _03c | Lepraria finkii | 2022-05-03 | CAM-NSF-Lichen Genomics Team; Allen, Jessica L. | USA: Shenandoah National Park, Central District, SW slopes of Hawksbill Mountain, between Lower Hawksbill Trail and Hawksbill Fire Road ~0.7 mi W of Hawksbill Gap | New York State Museum | 38.55397 N 78.39687 W | NYSM0000191742 |
| LEFI_HAW _03b | Lepraria finkii | 2022-05-03 | CAM-NSF-Lichen Genomics Team; Allen, Jessica L. | USA: Shenandoah National Park, Central District, SW slopes of Hawksbill Mountain, between Lower Hawksbill Trail and Hawksbill Fire Road ~0.7 mi W of Hawksbill Gap | New York State Museum | 38.55397 N 78.39687 W | NYSM0000194972 |
| LEFI_HAW _03a | Lepraria finkii | 2022-05-03 | CAM-NSF-Lichen Genomics Team; Allen, Jessica L. | USA: Shenandoah National Park, Central District, SW slopes of Hawksbill Mountain, between Lower Hawksbill Trail and Hawksbill Fire Road ~0.7 mi W of Hawksbill Gap | New York State Museum | 38.55397 N 78.39687 W | NYSM0000191741 |
| LEFI_HAW _02c | Lepraria finkii | 2022-05-03 | CAM-NSF-Lichen Genomics Team; Allen, Jessica L. | USA: Shenandoah National Park, Central District, SW slopes of Hawksbill Mountain, between Lower Hawksbill Trail and Hawksbill Fire Road ~0.7 mi W of Hawksbill Gap | New York State Museum | 38.55397 N 78.39687 W | NYSM0000191740 |
| LEFI_HAW _02a | Lepraria finkii | 2022-05-03 | CAM-NSF-Lichen Genomics Team; Allen, Jessica L. | USA: Shenandoah National Park, Central District, SW slopes of Hawksbill Mountain, between Lower Hawksbill Trail and Hawksbill Fire Road ~0.7 mi W of Hawksbill Gap | New York State Museum | 38.55397 N 78.39687 W | NYSM0000191739 |

| | | | | | | | |
|---|---|---|---|---|---|---|---|
| LEFI_HAW _01c | Lepraria finkii | 2022-05-03 | CAM-NSF-Lichen Genomics Team; Allen, Jessica L. | USA: Shenandoah National Park, Central District, SW slopes of Hawksbill Mountain, between Lower Hawksbill Trail and Hawksbill Fire Road ~0.7 mi W of Hawksbill Gap | New York State Museum | 38.55397 N 78.39687 W | NYSM0000191738 |
| LEFI_HAW _01b | Lepraria finkii | 2022-05-03 | CAM-NSF-Lichen Genomics Team; Allen, Jessica L. | USA: Shenandoah National Park, Central District, SW slopes of Hawksbill Mountain, between Lower Hawksbill Trail and Hawksbill Fire Road ~0.7 mi W of Hawksbill Gap | New York State Museum | 38.55397 N 78.39687 W | NYSM0000191737 |
| LEFI_HAW _01a | Lepraria finkii | 2022-05-03 | CAM-NSF-Lichen Genomics Team; Allen, Jessica L. | USA: Shenandoah National Park, Central District, SW slopes of Hawksbill Mountain, between Lower Hawksbill Trail and Hawksbill Fire Road ~0.7 mi W of Hawksbill Gap | New York State Museum | 38.55397 N 78.39687 W | NYSM0000191736 |
| LEFI_GRI_0 7d | Lepraria finkii | 2022-05-02 | CAM-NSF-Lichen Genomics Team; Allen, Jessica L. | USA: Shenandoah National Park, Central District, S facing slopes of Grindstone Mountain above Deep Run, ~0.2 mi S of summit | New York State Museum | 38.47288 N 78.550732 W | NYSM0000191733 |
| LEFI_GRI_0 7c | Lepraria finkii | 2022-05-02 | CAM-NSF-Lichen Genomics Team; Allen, Jessica L. | USA: Shenandoah National Park, Central District, S facing slopes of Grindstone Mountain above Deep Run, ~0.2 mi S of summit | New York State Museum | 38.47288 N 78.550732 W | NYSM0000191732 |

| | | | | | | | |
|---|---|---|---|---|---|---|---|
| LEFI_GRI_07b | Lepraria finkii | 2022-05-02 | CAM-NSF-Lichen Genomics Team; Allen, Jessica L. | USA: Shenandoah National Park, Central District, S facing slopes of Grindstone Mountain above Deep Run, ~0.2 mi S of summit | New York State Museum | 38.47288 N 78.550732 W | NYSM0000191731 |
| LEFI_GRI_07a | Lepraria finkii | 2022-05-02 | CAM-NSF-Lichen Genomics Team; Allen, Jessica L. | USA: Shenandoah National Park, Central District, S facing slopes of Grindstone Mountain above Deep Run, ~0.2 mi S of summit | New York State Museum | 38.47288 N 78.550732 W | NYSM0000191730 |
| LEFI_GRI_06c | Lepraria finkii | 2022-05-02 | CAM-NSF-Lichen Genomics Team; Allen, Jessica L. | USA: Shenandoah National Park, Central District, S facing slopes of Grindstone Mountain above Deep Run, ~0.2 mi S of summit | New York State Museum | 38.47288 N 78.550732 W | NYSM0000191729 |
| LEFI_GRI_06b | Lepraria finkii | 2022-05-02 | CAM-NSF-Lichen Genomics Team; Allen, Jessica L. | USA: Shenandoah National Park, Central District, S facing slopes of Grindstone Mountain above Deep Run, ~0.2 mi S of summit | New York State Museum | 38.47288 N 78.550732 W | NYSM0000191728 |
| LEFI_GRI_06a | Lepraria finkii | 2022-05-02 | CAM-NSF-Lichen Genomics Team; Allen, Jessica L. | USA: Shenandoah National Park, Central District, S facing slopes of Grindstone Mountain above Deep Run, ~0.2 mi S of summit | New York State Museum | 38.47288 N 78.550732 W | NYSM0000191727 |
| LEFI_GRI_05b | Lepraria finkii | 2022-05-02 | CAM-NSF-Lichen Genomics Team; Allen, Jessica L. | USA: Shenandoah National Park, Central District, S facing slopes of Grindstone Mountain above Deep Run, ~0.2 mi S of summit | New York State Museum | 38.47288 N 78.550732 W | NYSM0000191726 |

| | | | | | | | |
|---|---|---|---|---|---|---|---|
| LEFI_GRI_05a | Lepraria finkii | 2022-05-02 | CAM-NSF-Lichen Genomics Team; Allen, Jessica L. | USA: Shenandoah National Park, Central District, S facing slopes of Grindstone Mountain above Deep Run, ~0.2 mi S of summit | New York State Museum | 38.47288 N 78.550732 W | NYSM0000191725 |
| LEFI_GRI_04b | Lepraria finkii | 2022-05-02 | CAM-NSF-Lichen Genomics Team; Allen, Jessica L. | USA: Shenandoah National Park, Central District, S facing slopes of Grindstone Mountain above Deep Run, ~0.2 mi S of summit | New York State Museum | 38.47288 N 78.550732 W | NYSM0000191724 |
| LEFI_GRI_04a | Lepraria finkii | 2022-04-11 | CAM-NSF-Lichen Genomics Team; Allen, Jessica L. | USA: E-slopes of Pine Mountain, Kentenia State Forest, vicinity of Goss Park, E of Little Shepherd Trail/SR1679 at jct w/ Putney Lookout Tower Rd., between Spruce Ridge Branch and Hi Lewis Branch | New York State Museum | 36.9243 N 83.22419 W | NYSM0000194971 |
| LEFI_GOS_11b | Lepraria finkii | 2022-04-11 | CAM-NSF-Lichen Genomics Team; Allen, Jessica L. | USA: E-slopes of Pine Mountain, Kentenia State Forest, vicinity of Goss Park, E of Little Shepherd Trail/SR1679 at jct w/ Putney Lookout Tower Rd., between Spruce Ridge Branch and Hi Lewis Branch | New York State Museum | 36.9243 N 83.22419 W | NYSM0000194970 |
| LEFI_GOS_11a | Lepraria finkii | 2022-04-11 | CAM-NSF-Lichen Genomics Team; Allen, Jessica L. | USA: E-slopes of Pine Mountain, Kentenia State Forest, vicinity of Goss Park, E of Little Shepherd Trail/SR1679 at jct w/ Putney Lookout Tower Rd., between Spruce Ridge Branch and Hi Lewis Branch | New York State Museum | 36.9243 N 83.22419 W | NYSM0000194968 |

| | | | | | | | |
|---|---|---|---|---|---|---|---|
| LEFI_GOS_08b | Lepraria finkii | 2022-04-11 | CAM-NSF-Lichen Genomics Team; Allen, Jessica L. | USA: E-slopes of Pine Mountain, Kentenia State Forest, vicinity of Goss Park, E of Little Shepherd Trail/SR1679 at jct w/ Putney Lookout Tower Rd., between Spruce Ridge Branch and Hi Lewis Branch | New York State Museum | 36.9243 N 83.22419 W | NYSM0000194969 |
| LEFI_GOS_08a | Lepraria finkii | 2022-04-11 | CAM-NSF-Lichen Genomics Team; Allen, Jessica L. | USA: E-slopes of Pine Mountain, Kentenia State Forest, vicinity of Goss Park, E of Little Shepherd Trail/SR1679 at jct w/ Putney Lookout Tower Rd., between Spruce Ridge Branch and Hi Lewis Branch | New York State Museum | 36.9243 N 83.22419 W | NYSM0000194967 |
| LEFI_GOS_07c | Lepraria finkii | 2022-04-11 | CAM-NSF-Lichen Genomics Team; Allen, Jessica L. | USA: E-slopes of Pine Mountain, Kentenia State Forest, vicinity of Goss Park, E of Little Shepherd Trail/SR1679 at jct w/ Putney Lookout Tower Rd., between Spruce Ridge Branch and Hi Lewis Branch | New York State Museum | 36.9243 N 83.22419 W | NYSM0000194966 |
| LEFI_GOS_07b | Lepraria finkii | 2022-04-11 | CAM-NSF-Lichen Genomics Team; Allen, Jessica L. | USA: E-slopes of Pine Mountain, Kentenia State Forest, vicinity of Goss Park, E of Little Shepherd Trail/SR1679 at jct w/ Putney Lookout Tower Rd., between Spruce Ridge Branch and Hi Lewis Branch | New York State Museum | 36.9243 N 83.22419 W | NYSM0000194965 |

| | | | | | | | |
|---|---|---|---|---|---|---|---|
| LEFI_GOS_07a | Lepraria finkii | 2022-04-11 | CAM-NSF-Lichen Genomics Team; Allen, Jessica L. | USA: E-slopes of Pine Mountain, Kentenia State Forest, vicinity of Goss Park, E of Little Shepherd Trail/SR1679 at jct w/ Putney Lookout Tower Rd., between Spruce Ridge Branch and Hi Lewis Branch | New York State Museum | 36.9243 N 83.22419 W | NYSM0000194964 |
| LEFI_GOS_03c | Lepraria finkii | 2022-04-11 | CAM-NSF-Lichen Genomics Team; Allen, Jessica L. | USA: E-slopes of Pine Mountain, Kentenia State Forest, vicinity of Goss Park, E of Little Shepherd Trail/SR1679 at jct w/ Putney Lookout Tower Rd., between Spruce Ridge Branch and Hi Lewis Branch | New York State Museum | 36.9243 N 83.22419 W | NYSM0000194963 |
| LEFI_GOS_03b | Lepraria finkii | 2022-04-11 | CAM-NSF-Lichen Genomics Team; Allen, Jessica L. | USA: E-slopes of Pine Mountain, Kentenia State Forest, vicinity of Goss Park, E of Little Shepherd Trail/SR1679 at jct w/ Putney Lookout Tower Rd., between Spruce Ridge Branch and Hi Lewis Branch | New York State Museum | 36.9243 N 83.22419 W | NYSM0000194962 |
| LEFI_GOS_03a | Lepraria finkii | 2022-08-01 | CAM-NSF-Lichen Genomics Team; Denzer, Eli | USA: Nantahala National Forest, N-facing slopes above FS4535 0.3 mi W of jct w/ SR1538/Buck Creek Rd. at Cole Gap | New York State Museum | 35.1036 N 83.2075 W | NYSM0000191723 |
| LEFI_COG_15a | Lepraria finkii | 2022-08-01 | CAM-NSF-Lichen Genomics Team; Denzer, Eli | USA: Nantahala National Forest, N-facing slopes above FS4535 0.3 mi W of jct w/ SR1538/Buck Creek Rd. at Cole Gap | New York State Museum | 35.1036 N 83.2075 W | |

| | | | | | | | |
|---|---|---|---|---|---|---|---|
| LEFI_COG_13b | Lepraria finkii | 2022-08-01 | CAM-NSF-Lichen Genomics Team; Denzer, Eli | USA: Nantahala National Forest, N-facing slopes above FS4535 0.3 mi W of jct w/ SR1538/Buck Creek Rd. at Cole Gap | New York State Museum | 35.1036 N 83.2075 W | NYSM0000191722 |
| LEFI_COG_13a | Lepraria finkii | 2022-08-01 | CAM-NSF-Lichen Genomics Team; Denzer, Eli | USA: Nantahala National Forest, N-facing slopes above FS4535 0.3 mi W of jct w/ SR1538/Buck Creek Rd. at Cole Gap | New York State Museum | 35.1036 N 83.2075 W | NYSM0000191721 |
| LEFI_COG_11b | Lepraria finkii | 2022-08-01 | CAM-NSF-Lichen Genomics Team; Denzer, Eli | USA: Nantahala National Forest, N-facing slopes above FS4535 0.3 mi W of jct w/ SR1538/Buck Creek Rd. at Cole Gap | New York State Museum | 35.1036 N 83.2075 W | NYSM0000191720 |
| LEFI_COG_11a | Lepraria finkii | 2022-08-01 | CAM-NSF-Lichen Genomics Team; Denzer, Eli | USA: Nantahala National Forest, N-facing slopes above FS4535 0.3 mi W of jct w/ SR1538/Buck Creek Rd. at Cole Gap | New York State Museum | 35.1036 N 83.2075 W | NYSM0000191719 |
| LEFI_CAT_18c | Lepraria finkii | 2021-10-27 | CAM-NSF-Lichen Genomics Team; Allen, Jessica L. | USA: Pisgah National Forest, summit of Cattail Peak, Black Mountain Crest Trail 3 mi N of Mount Mitchell, ~8 mi S of US19E | New York State Museum | 35.7964 N 82.2569 W | NYSM0000191718 |
| LEFI_CAT_18b | Lepraria finkii | 2021-10-27 | CAM-NSF-Lichen Genomics Team; Allen, Jessica L. | USA: Pisgah National Forest, summit of Cattail Peak, Black Mountain Crest Trail 3 mi N of Mount Mitchell, ~8 mi S of US19E | New York State Museum | 35.7964 N 82.2569 W | NYSM0000191717 |

| | | | | | | | |
|---|---|---|---|---|---|---|---|
| LEFI_CAT_ 18a | Lepraria finkii | 2021-10-27 | CAM-NSF-Lichen Genomics Team; Allen, Jessica L. | USA: Pisgah National Forest, summit of Cattail Peak, Black Mountain Crest Trail 3 mi N of Mount Mitchell, ~8 mi S of US19E | New York State Museum | 35.7964 N 82.2569 W | NYSM0000191716 |
| LEFI_CAT_ 17b | Lepraria finkii | 2021-10-27 | CAM-NSF-Lichen Genomics Team; Allen, Jessica L. | USA: Pisgah National Forest, summit of Cattail Peak, Black Mountain Crest Trail 3 mi N of Mount Mitchell, ~8 mi S of US19E | New York State Museum | 35.7964 N 82.2569 W | NYSM0000191715 |
| LEFI_CAT_ 17a | Lepraria finkii | 2021-10-27 | CAM-NSF-Lichen Genomics Team; Allen, Jessica L. | USA: Pisgah National Forest, summit of Cattail Peak, Black Mountain Crest Trail 3 mi N of Mount Mitchell, ~8 mi S of US19E | New York State Museum | 35.7964 N 82.2569 W | NYSM0000191714 |
| LEFI_CAT_ 16b | Lepraria finkii | 2021-10-27 | CAM-NSF-Lichen Genomics Team; Allen, Jessica L. | USA: Pisgah National Forest, summit of Cattail Peak, Black Mountain Crest Trail 3 mi N of Mount Mitchell, ~8 mi S of US19E | New York State Museum | 35.7964 N 82.2569 W | NYSM0000191713 |
| LEFI_CAT_ 16a | Lepraria finkii | 2021-10-27 | CAM-NSF-Lichen Genomics Team; Allen, Jessica L. | USA: Pisgah National Forest, summit of Cattail Peak, Black Mountain Crest Trail 3 mi N of Mount Mitchell, ~8 mi S of US19E | New York State Museum | 35.7964 N 82.2569 W | NYSM0000191710 |
| LEFI_CAT_ 15c | Lepraria finkii | 2021-10-27 | CAM-NSF-Lichen Genomics Team; Allen, Jessica L. | USA: Pisgah National Forest, summit of Cattail Peak, Black Mountain Crest Trail 3 mi N of Mount Mitchell, ~8 mi S of US19E | New York State Museum | 35.7964 N 82.2569 W | NYSM0000191712 |

| | | | | | | | |
|---|---|---|---|---|---|---|---|
| LEFI_CAT_ 15b | Lepraria finkii | 2021-10-27 | CAM-NSF-Lichen Genomics Team; Allen, Jessica L. | USA: Pisgah National Forest, summit of Cattail Peak, Black Mountain Crest Trail 3 mi N of Mount Mitchell, ~8 mi S of US19E | New York State Museum | 35.7964 N 82.2569 W | NYSM0000191711 |
| LEFI_CAT_ 15a | Lepraria finkii | 2021-10-27 | CAM-NSF-Lichen Genomics Team; Allen, Jessica L. | USA: Pisgah National Forest, summit of Cattail Peak, Black Mountain Crest Trail 3 mi N of Mount Mitchell, ~8 mi S of US19E | New York State Museum | 35.7964 N 82.2569 W | NYSM0000191709 |
| LEFI_BUK_ 21a | Lepraria finkii | 2021-10-24 | CAM-NSF-Lichen Genomics Team; Allen, Jessica L. | USA: Great Smoky Mountains National Park, Cataloochee Divide, N-slope of Little Bald Knob, Hemphill Bald Trail ~3 mi E of Polls Gap | New York State Museum | 35.552276 N 83.120161 W | NYSM0000191708 |
| LEFI_BUK_ 16a | Lepraria finkii | 2021-10-24 | CAM-NSF-Lichen Genomics Team; Allen, Jessica L. | USA: Great Smoky Mountains National Park, Cataloochee Divide, N-slope of Little Bald Knob, Hemphill Bald Trail ~3 mi E of Polls Gap | New York State Museum | 35.552276 N 83.120161 W | NYSM0000191707 |
| LEFI_BUK_ 14a | Lepraria finkii | 2021-10-24 | CAM-NSF-Lichen Genomics Team; Allen, Jessica L. | USA: Great Smoky Mountains National Park, Cataloochee Divide, N-slope of Little Bald Knob, Hemphill Bald Trail ~3 mi E of Polls Gap | New York State Museum | 35.552276 N 83.120161 W | NYSM0000191706 |
| LEFI_BUK_ 13a | Lepraria finkii | 2021-10-24 | CAM-NSF-Lichen Genomics Team; Allen, Jessica L. | USA: Great Smoky Mountains National Park, Cataloochee Divide, N-slope of Little Bald Knob, Hemphill Bald Trail ~3 mi E of Polls Gap | New York State Museum | 35.552276 N 83.120161 W | NYSM0000191705 |

| | | | | | | | |
|---|---|---|---|---|---|---|---|
| LEFI_BUK_12a | Lepraria finkii | 2021-10-24 | CAM-NSF-Lichen Genomics Team; Allen, Jessica L. | USA: Great Smoky Mountains National Park, Cataloochee Divide, N-slope of Little Bald Knob, Hemphill Bald Trail ~3 mi E of Polls Gap | New York State Museum | 35.552276 N 83.120161 W | NYSM0000191704 |
| LEFI_BUK_09a | Lepraria finkii | 2021-10-24 | CAM-NSF-Lichen Genomics Team; Allen, Jessica L. | USA: Great Smoky Mountains National Park, Cataloochee Divide, N-slope of Little Bald Knob, Hemphill Bald Trail ~3 mi E of Polls Gap | New York State Museum | 35.552276 N 83.120161 W | NYSM0000191703 |
| LEFI_BUK_08a | Lepraria finkii | 2021-10-24 | CAM-NSF-Lichen Genomics Team; Allen, Jessica L. | USA: Great Smoky Mountains National Park, Cataloochee Divide, N-slope of Little Bald Knob, Hemphill Bald Trail ~3 mi E of Polls Gap | New York State Museum | 35.552276 N 83.120161 W | NYSM0000191702 |
| LEFI_BUK_07a | Lepraria finkii | 2021-10-24 | CAM-NSF-Lichen Genomics Team; Allen, Jessica L. | USA: Great Smoky Mountains National Park, Cataloochee Divide, N-slope of Little Bald Knob, Hemphill Bald Trail ~3 mi E of Polls Gap | New York State Museum | 35.552276 N 83.120161 W | NYSM0000191701 |
| LEFI_BRF_06c | Lepraria finkii | 2022-05-03 | CAM-NSF-Lichen Genomics Team; Allen, Jessica L. | USA: Shenandoah National Park, Central District, Bearfence Mountain, W-slopes below Appalachian Trail, 0.3 mi S of jct w/ spur trail to Skyline Drive/Bearfence Parking Area | New York State Museum | 38.4477 N 78.46635 W | NYSM0000194961 |

| | | | | | | | |
|---|---|---|---|---|---|---|---|
| LEFI_BRF_06b | Lepraria finkii | 2022-05-03 | CAM-NSF-Lichen Genomics Team; Allen, Jessica L. | USA: Shenandoah National Park, Central District, Bearfence Mountain, W-slopes below Appalachian Trail, 0.3 mi S of jct w/ spur trail to Skyline Drive/Bearfence Parking Area | New York State Museum | 38.4477 N 78.46635 W | NYSM0000194960 |
| LEFI_BRF_06a | Lepraria finkii | 2022-05-03 | CAM-NSF-Lichen Genomics Team; Allen, Jessica L. | USA: Shenandoah National Park, Central District, Bearfence Mountain, W-slopes below Appalachian Trail, 0.3 mi S of jct w/ spur trail to Skyline Drive/Bearfence Parking Area | New York State Museum | 38.4477 N 78.46635 W | NYSM0000191700 |
| LEFI_BRF_05c | Lepraria finkii | 2022-05-03 | CAM-NSF-Lichen Genomics Team; Allen, Jessica L. | USA: Shenandoah National Park, Central District, Bearfence Mountain, W-slopes below Appalachian Trail, 0.3 mi S of jct w/ spur trail to Skyline Drive/Bearfence Parking Area | New York State Museum | 38.4477 N 78.46635 W | NYSM0000191699 |
| LEFI_BRF_05b | Lepraria finkii | 2022-05-03 | CAM-NSF-Lichen Genomics Team; Allen, Jessica L. | USA: Shenandoah National Park, Central District, Bearfence Mountain, W-slopes below Appalachian Trail, 0.3 mi S of jct w/ spur trail to Skyline Drive/Bearfence Parking Area | New York State Museum | 38.4477 N 78.46635 W | NYSM0000191698 |
| LEFI_BRF_05a | Lepraria finkii | 2022-05-03 | CAM-NSF-Lichen Genomics Team; Allen, Jessica L. | USA: Shenandoah National Park, Central District, Bearfence Mountain, W-slopes below Appalachian Trail, 0.3 mi S of jct w/ spur trail to Skyline Drive/Bearfence Parking Area | New York State Museum | 38.4477 N 78.46635 W | NYSM0000191697 |

| | | | | | | | |
|---|---|---|---|---|---|---|---|
| LEFI_BRF_03c | Lepraria finkii | 2022-05-03 | CAM-NSF-Lichen Genomics Team; Allen, Jessica L. | USA: Shenandoah National Park, Central District, Bearfence Mountain, W-slopes below Appalachian Trail, 0.3 mi S of jct w/ spur trail to Skyline Drive/Bearfence Parking Area | New York State Museum | 38.4477 N 78.46635 W | |
| LEFI_BRF_03b | Lepraria finkii | 2022-05-03 | CAM-NSF-Lichen Genomics Team; Allen, Jessica L. | USA: Shenandoah National Park, Central District, Bearfence Mountain, W-slopes below Appalachian Trail, 0.3 mi S of jct w/ spur trail to Skyline Drive/Bearfence Parking Area | New York State Museum | 38.4477 N 78.46635 W | |
| LEFI_BRF_03a | Lepraria finkii | 2022-05-03 | CAM-NSF-Lichen Genomics Team; Allen, Jessica L. | USA: Shenandoah National Park, Central District, Bearfence Mountain, W-slopes below Appalachian Trail, 0.3 mi S of jct w/ spur trail to Skyline Drive/Bearfence Parking Area | New York State Museum | 38.4477 N 78.46635 W | |
| LEFI_BRF_02c | Lepraria finkii | 2022-05-03 | CAM-NSF-Lichen Genomics Team; Allen, Jessica L. | USA: Shenandoah National Park, Central District, Bearfence Mountain, W-slopes below Appalachian Trail, 0.3 mi S of jct w/ spur trail to Skyline Drive/Bearfence Parking Area | New York State Museum | 38.4477 N 78.46635 W | NYSM0000191696 |
| LEFI_BRF_02b | Lepraria finkii | 2022-05-03 | CAM-NSF-Lichen Genomics Team; Allen, Jessica L. | USA: Shenandoah National Park, Central District, Bearfence Mountain, W-slopes below Appalachian Trail, 0.3 mi S of jct w/ spur trail to Skyline Drive/Bearfence Parking Area | New York State Museum | 38.4477 N 78.46635 W | NYSM0000191695 |

| | | | | | | | |
|---|---|---|---|---|---|---|---|
| LEFI_BRF_02a | Lepraria finkii | 2022-05-03 | CAM-NSF-Lichen Genomics Team; Allen, Jessica L. | USA: Shenandoah National Park, Central District, Bearfence Mountain, W-slopes below Appalachian Trail, 0.3 mi S of jct w/ spur trail to Skyline Drive/Bearfence Parking Area | New York State Museum | 38.4477 N 78.46635 W | NYSM0000191694 |
| LEFI_BLU_08b | Lepraria finkii | 2022-04-13 | CAM-NSF-Lichen Genomics Team; Allen, Jessica L. | USA: Pine Mountain, Bad Branch Nature Preserve, headwaters of Presley House Branch, Pine Mountain Trail ~1.5 mi E of US119 | New York State Museum | 37.08482 N 82.79308 W | NYSM0000194959 |
| LEFI_BLU_08a | Lepraria finkii | 2022-04-13 | CAM-NSF-Lichen Genomics Team; Allen, Jessica L. | USA: Pine Mountain, Bad Branch Nature Preserve, headwaters of Presley House Branch, Pine Mountain Trail ~1.5 mi E of US119 | New York State Museum | 37.08482 N 82.79308 W | NYSM0000194958 |
| LEFI_BLU_07a | Lepraria finkii | 2022-04-13 | CAM-NSF-Lichen Genomics Team; Allen, Jessica L. | USA: Pine Mountain, Bad Branch Nature Preserve, headwaters of Presley House Branch, Pine Mountain Trail ~1.5 mi E of US119 | New York State Museum | 37.08482 N 82.79308 W | NYSM0000194957 |
| LEFI_BLU_06b | Lepraria finkii | 2022-04-13 | CAM-NSF-Lichen Genomics Team; Allen, Jessica L. | USA: Pine Mountain, Bad Branch Nature Preserve, headwaters of Presley House Branch, Pine Mountain Trail ~1.5 mi E of US119 | New York State Museum | 37.08482 N 82.79308 W | NYSM0000194956 |

| | | | | | | | |
|---|---|---|---|---|---|---|---|
| LEFI_BLU_06a | Lepraria finkii | 2022-04-13 | CAM-NSF-Lichen Genomics Team; Allen, Jessica L. | USA: Pine Mountain, Bad Branch Nature Preserve, headwaters of Presley House Branch, Pine Mountain Trail ~1.5 mi E of US119 | New York State Museum | 37.08482 N 82.79308 W | NYSM0000194955 |
| LEFI_BLU_05a | Lepraria finkii | 2022-04-13 | CAM-NSF-Lichen Genomics Team; Allen, Jessica L. | USA: Pine Mountain, Bad Branch Nature Preserve, headwaters of Presley House Branch, Pine Mountain Trail ~1.5 mi E of US119 | New York State Museum | 37.08482 N 82.79308 W | |
| LEFI_BLU_04a | Lepraria finkii | 2022-04-13 | CAM-NSF-Lichen Genomics Team; Allen, Jessica L. | USA: Pine Mountain, Bad Branch Nature Preserve, headwaters of Presley House Branch, Pine Mountain Trail ~1.5 mi E of US119 | New York State Museum | 37.08482 N 82.79308 W | NYSM0000194954 |
| LEFI_BLU_03c | Lepraria finkii | 2022-04-13 | CAM-NSF-Lichen Genomics Team; Allen, Jessica L. | USA: Pine Mountain, Bad Branch Nature Preserve, headwaters of Presley House Branch, Pine Mountain Trail ~1.5 mi E of US119 | New York State Museum | 37.08482 N 82.79308 W | NYSM0000194953 |
| LEFI_BLU_03b | Lepraria finkii | 2022-04-13 | CAM-NSF-Lichen Genomics Team; Allen, Jessica L. | USA: Pine Mountain, Bad Branch Nature Preserve, headwaters of Presley House Branch, Pine Mountain Trail ~1.5 mi E of US119 | New York State Museum | 37.08482 N 82.79308 W | NYSM0000194952 |
| LEFI_BLU_03a | Lepraria finkii | 2022-04-13 | CAM-NSF-Lichen Genomics Team; Allen, Jessica L. | USA: Pine Mountain, Bad Branch Nature Preserve, headwaters of Presley House Branch, Pine Mountain Trail ~1.5 mi E of US119 | New York State Museum | 37.08482 N 82.79308 W | NYSM0000194951 |

| | | | | | | | |
|---|---|---|---|---|---|---|---|
| LEFI_BLU_02a | Lepraria finkii | 2022-04-13 | CAM-NSF-Lichen Genomics Team; Allen, Jessica L. | USA: Pine Mountain, Bad Branch Nature Preserve, headwaters of Presley House Branch, Pine Mountain Trail ~1.5 mi E of US119 | New York State Museum | 37.08482 N 82.79308 W | NYSM0000194950 |
| LEFI_BFF_28a | Lepraria finkii | 2021-10-26 | CAM-NSF-Lichen Genomics Team; Allen, Jessica L. | USA: Pisgah National Forest, Bald Mountains, N-Facing slopes of Bluff Mountain summit, NE of Appalachian Trail 1 mi E of Catpen Gap, 3.4 mi E of Lemon Gap | New York State Museum | 35.84115 N 82.90576 W | NYSM0000191693 |
| LEFI_BFF_27a | Lepraria finkii | 2021-10-27 | CAM-NSF-Lichen Genomics Team; Allen, Jessica L. | USA: Pisgah National Forest, Bald Mountains, N-Facing slopes of Bluff Mountain summit, NE of Appalachian Trail 1 mi E of Catpen Gap, 3.4 mi E of Lemon Gap | New York State Museum | 35.84115 N 82.90576 W | |
| LEFI_BFF_23a | Lepraria finkii | 2021-10-26 | CAM-NSF-Lichen Genomics Team; Allen, Jessica L. | USA: Pisgah National Forest, Bald Mountains, N-Facing slopes of Bluff Mountain summit, NE of Appalachian Trail 1 mi E of Catpen Gap, 3.4 mi E of Lemon Gap | New York State Museum | 35.84115 N 82.90576 W | NYSM0000191692 |
| LEFI_BFF_21a | Lepraria finkii | 2021-10-26 | CAM-NSF-Lichen Genomics Team; Allen, Jessica L. | USA: Pisgah National Forest, Bald Mountains, N-Facing slopes of Bluff Mountain summit, NE of Appalachian Trail 1 mi E of Catpen Gap, 3.4 mi E of Lemon Gap | New York State Museum | 35.84115 N 82.90576 W | NYSM0000191690 |

| | | | | | | | |
|---|---|---|---|---|---|---|---|
| LEFI_BFF_20a | Lepraria finkii | 2021-10-26 | CAM-NSF-Lichen Genomics Team; Allen, Jessica L. | USA: Pisgah National Forest, Bald Mountains, N-Facing slopes of Bluff Mountain summit, NE of Appalachian Trail 1 mi E of Catpen Gap, 3.4 mi E of Lemon Gap | New York State Museum | 35.84115 N 82.90576 W | |
| LEFI_BFF_13a | Lepraria finkii | 2021-10-26 | CAM-NSF-Lichen Genomics Team; Allen, Jessica L. | USA: Pisgah National Forest, Bald Mountains, N-Facing slopes of Bluff Mountain summit, NE of Appalachian Trail 1 mi E of Catpen Gap, 3.4 mi E of Lemon Gap | New York State Museum | 35.84115 N 82.90576 W | NYSM0000191689 |
| LEFI_BFF_12a | Lepraria finkii | 2021-10-26 | CAM-NSF-Lichen Genomics Team; Allen, Jessica L. | USA: Pisgah National Forest, Bald Mountains, N-Facing slopes of Bluff Mountain summit, NE of Appalachian Trail 1 mi E of Catpen Gap, 3.4 mi E of Lemon Gap | New York State Museum | 35.84115 N 82.90576 W | NYSM0000191688 |
| LEFI_BFF_06a | Lepraria finkii | 2021-10-26 | CAM-NSF-Lichen Genomics Team; Allen, Jessica L. | USA: Pisgah National Forest, Bald Mountains, N-Facing slopes of Bluff Mountain summit, NE of Appalachian Trail 1 mi E of Catpen Gap, 3.4 mi E of Lemon Gap | New York State Museum | 35.84115 N 82.90576 W | NYSM0000191687 |
| LEFI_BFF_03a | Lepraria finkii | 2021-10-26 | CAM-NSF-Lichen Genomics Team; Allen, Jessica L. | USA: Pisgah National Forest, Bald Mountains, N-Facing slopes of Bluff Mountain summit, NE of Appalachian Trail 1 mi E of Catpen Gap, 3.4 mi E of Lemon Gap | New York State Museum | 35.84115 N 82.90576 W | NYSM0000191686 |

| | | | | | | | |
|---|---|---|---|---|---|---|---|
| LEFI_BFF_02a | Lepraria finkii | 2021-10-26 | CAM-NSF-Lichen Genomics Team; Allen, Jessica L. | USA: Pisgah National Forest, Bald Mountains, N-Facing slopes of Bluff Mountain summit, NE of Appalachian Trail 1 mi E of Catpen Gap, 3.4 mi E of Lemon Gap | New York State Museum | 35.84115 N 82.90576 W | NYSM0000191684 |
| LEFI_BEK_24a | Lepraria finkii | 2021-10-26 | CAM-NSF-Lichen Genomics Team; Allen, Jessica L. | USA: Pisgah National Forest, Bald Mountains, N-slope of Roaring Fork Mountain, spur of Buckeye Ridge | New York State Museum | 35.8014 N 82.95096 W | NYSM0000191683 |
| LEFI_BEK_23a | Lepraria finkii | 2021-10-26 | CAM-NSF-Lichen Genomics Team; Allen, Jessica L. | USA: Pisgah National Forest, Bald Mountains, N-slope of Roaring Fork Mountain, spur of Buckeye Ridge | New York State Museum | 35.8014 N 82.95096 W | NYSM0000191682 |
| LEFI_BEK_21a | Lepraria finkii | 2021-10-26 | CAM-NSF-Lichen Genomics Team; Allen, Jessica L. | USA: Pisgah National Forest, Bald Mountains, N-slope of Roaring Fork Mountain, spur of Buckeye Ridge | New York State Museum | 35.8014 N 82.95096 W | NYSM0000191681 |
| LEFI_BEK_20a | Lepraria finkii | 2021-10-26 | CAM-NSF-Lichen Genomics Team; Allen, Jessica L. | USA: Pisgah National Forest, Bald Mountains, N-slope of Roaring Fork Mountain, spur of Buckeye Ridge | New York State Museum | 35.8014 N 82.95096 W | NYSM0000191680 |

| | | | | | | | |
|---|---|---|---|---|---|---|---|
| LEFI_BEK_13a | Lepraria finkii | 2021-10-26 | CAM-NSF-Lichen Genomics Team; Allen, Jessica L. | USA: Pisgah National Forest, Bald Mountains, N-slope of Roaring Fork Mountain, spur of Buckeye Ridge | New York State Museum | 35.8014 N 82.95096 W | NYSM0000191677 |
| PUAP_WHI_19A | Punctelia appalachensis | 2022-08-04 | CAM-NSF-Lichen Genomics Team; Denzer, Eli | USA: Nantahala National Forest, summit of Whiterock Mountain, spur trail 0.3 mi N of jct w/ Bartram Trail, 1.3 mi NW of Jones Gap parking area at terminus of FS 4522 | New York State Museum | 35.085 N 83.3053 W | NYSM0000194427 |
| PUAP_WHI_17C | Punctelia appalachensis | 2022-08-04 | CAM-NSF-Lichen Genomics Team; Denzer, Eli | USA: Nantahala National Forest, summit of Whiterock Mountain, spur trail 0.3 mi N of jct w/ Bartram Trail, 1.3 mi NW of Jones Gap parking area at terminus of FS 4522 | New York State Museum | 35.085 N 83.3053 W | NYSM0000194426 |
| PUAP_WHI_17b | Punctelia appalachensis | 2022-08-04 | CAM-NSF-Lichen Genomics Team; Denzer, Eli | USA: Nantahala National Forest, summit of Whiterock Mountain, spur trail 0.3 mi N of jct w/ Bartram Trail, 1.3 mi NW of Jones Gap parking area at terminus of FS 4522 | New York State Museum | 35.085 N 83.3053 W | NYSM0000194425 |
| PUAP_WHI_17A | Punctelia appalachensis | 2022-08-04 | CAM-NSF-Lichen Genomics Team; Denzer, Eli | USA: Nantahala National Forest, summit of Whiterock Mountain, spur trail 0.3 mi N of jct w/ Bartram Trail, 1.3 mi NW of Jones Gap parking area at terminus of FS 4522 | New York State Museum | 35.085 N 83.3053 W | NYSM0000194424 |

| | | | | | | | |
|---|---|---|---|---|---|---|---|
| PUAP_WHI_16A | Punctelia appalachensis | 2022-08-04 | CAM-NSF-Lichen Genomics Team; Denzer, Eli | USA: Nantahala National Forest, summit of Whiterock Mountain, spur trail 0.3 mi N of jct w/ Bartram Trail, 1.3 mi NW of Jones Gap parking area at terminus of FS 4522 | New York State Museum | 35.085 N 83.3053 W | NYSM0000194423 |
| PUAP_WHI_14B | Punctelia appalachensis | 2022-08-04 | CAM-NSF-Lichen Genomics Team; Denzer, Eli | USA: Nantahala National Forest, summit of Whiterock Mountain, spur trail 0.3 mi N of jct w/ Bartram Trail, 1.3 mi NW of Jones Gap parking area at terminus of FS 4522 | New York State Museum | 35.085 N 83.3053 W | NYSM0000194421 |
| PUAP_WHI_14A | Punctelia appalachensis | 2022-08-04 | CAM-NSF-Lichen Genomics Team; Denzer, Eli | USA: Nantahala National Forest, summit of Whiterock Mountain, spur trail 0.3 mi N of jct w/ Bartram Trail, 1.3 mi NW of Jones Gap parking area at terminus of FS 4522 | New York State Museum | 35.085 N 83.3053 W | NYSM0000194420 |
| PUAP_WHI_13A | Punctelia appalachensis | 2022-08-04 | CAM-NSF-Lichen Genomics Team; Denzer, Eli | USA: Nantahala National Forest, summit of Whiterock Mountain, spur trail 0.3 mi N of jct w/ Bartram Trail, 1.3 mi NW of Jones Gap parking area at terminus of FS 4522 | New York State Museum | 35.085 N 83.3053 W | NYSM0000194419 |
| PUAP_WHI_12B | Punctelia appalachensis | 2022-08-04 | CAM-NSF-Lichen Genomics Team; Denzer, Eli | USA: Nantahala National Forest, summit of Whiterock Mountain, spur trail 0.3 mi N of jct w/ Bartram Trail, 1.3 mi NW of Jones Gap parking area at terminus of FS 4522 | New York State Museum | 35.085 N 83.3053 W | NYSM0000194418 |

| | | | | | | | |
|---|---|---|---|---|---|---|---|
| PUAP_WHI_12A | Punctelia appalachensis | 2022-08-04 | CAM-NSF-Lichen Genomics Team; Denzer, Eli | USA: Nantahala National Forest, summit of Whiterock Mountain, spur trail 0.3 mi N of jct w/ Bartram Trail, 1.3 mi NW of Jones Gap parking area at terminus of FS 4522 | New York State Museum | 35.085 N 83.3053 W | NYSM0000194417 |
| PUAP_WHI_06b | Punctelia appalachensis | 2022-08-04 | CAM-NSF-Lichen Genomics Team; Denzer, Eli | USA: Nantahala National Forest, summit of Whiterock Mountain, spur trail 0.3 mi N of jct w/ Bartram Trail, 1.3 mi NW of Jones Gap parking area at terminus of FS 4522 | New York State Museum | 35.085 N 83.3053 W | NYSM0000194416 |
| PUAP_WHI_06a | Punctelia appalachensis | 2022-08-04 | CAM-NSF-Lichen Genomics Team; Denzer, Eli | USA: Nantahala National Forest, summit of Whiterock Mountain, spur trail 0.3 mi N of jct w/ Bartram Trail, 1.3 mi NW of Jones Gap parking area at terminus of FS 4522 | New York State Museum | 35.085 N 83.3053 W | NYSM0000194415 |
| PUAP_WHI_04b | Punctelia appalachensis | 2022-08-04 | CAM-NSF-Lichen Genomics Team; Denzer, Eli | USA: Nantahala National Forest, summit of Whiterock Mountain, spur trail 0.3 mi N of jct w/ Bartram Trail, 1.3 mi NW of Jones Gap parking area at terminus of FS 4522 | New York State Museum | 35.085 N 83.3053 W | NYSM0000194414 |
| PUAP_WHI_04a | Punctelia appalachensis | 2022-08-04 | CAM-NSF-Lichen Genomics Team; Denzer, Eli | USA: Nantahala National Forest, summit of Whiterock Mountain, spur trail 0.3 mi N of jct w/ Bartram Trail, 1.3 mi NW of Jones Gap parking area at terminus of FS 4522 | New York State Museum | 35.085 N 83.3053 W | NYSM0000194413 |

| | | | | | | | |
|---|---|---|---|---|---|---|---|
| PUAP_TYS _04c | Punctelia appalachensis | 2022-05-03 | CAM-NSF-Lichen Genomics Team; Allen, Jessica L. | USA: Shenandoah National Park, Central District, Bettys Rock, ~0.3 mi N of Crescent Rock, upper W-facing slopes of Timber Hollow | New York State Museum | 38.56753 N 78.38223 W | NYSM0000194412 |
| PUAP_TYS _04b | Punctelia appalachensis | 2022-05-03 | CAM-NSF-Lichen Genomics Team; Allen, Jessica L. | USA: Shenandoah National Park, Central District, Bettys Rock, ~0.3 mi N of Crescent Rock, upper W-facing slopes of Timber Hollow | New York State Museum | 38.56753 N 78.38223 W | NYSM0000194411 |
| PUAP_TYS _04a | Punctelia appalachensis | 2022-05-03 | CAM-NSF-Lichen Genomics Team; Allen, Jessica L. | USA: Shenandoah National Park, Central District, Bettys Rock, ~0.3 mi N of Crescent Rock, upper W-facing slopes of Timber Hollow | New York State Museum | 38.56753 N 78.38223 W | NYSM0000194410 |
| PUAP_TYS _03c | Punctelia appalachensis | 2022-05-03 | CAM-NSF-Lichen Genomics Team; Allen, Jessica L. | USA: Shenandoah National Park, Central District, Bettys Rock, ~0.3 mi N of Crescent Rock, upper W-facing slopes of Timber Hollow | New York State Museum | 38.56753 N 78.38223 W | NYSM0000194409 |
| PUAP_TYS _03b | Punctelia appalachensis | 2022-05-03 | CAM-NSF-Lichen Genomics Team; Allen, Jessica L. | USA: Shenandoah National Park, Central District, Bettys Rock, ~0.3 mi N of Crescent Rock, upper W-facing slopes of Timber Hollow | New York State Museum | 38.56753 N 78.38223 W | NYSM0000194408 |
| PUAP_TYS _03a | Punctelia appalachensis | 2022-05-03 | CAM-NSF-Lichen Genomics Team; Allen, Jessica L. | USA: Shenandoah National Park, Central District, Bettys Rock, ~0.3 mi N of Crescent Rock, upper W-facing slopes of Timber Hollow | New York State Museum | 38.56753 N 78.38223 W | NYSM0000194407 |

| | | | | | | | |
|---|---|---|---|---|---|---|---|
| PUAP_TYS_02c | Punctelia appalachensis | 2022-05-03 | CAM-NSF-Lichen Genomics Team; Allen, Jessica L. | USA: Shenandoah National Park, Central District, Bettys Rock, ~0.3 mi N of Crescent Rock, upper W-facing slopes of Timber Hollow | New York State Museum | 38.56753 N 78.38223 W | NYSM0000194406 |
| PUAP_TYS_02b | Punctelia appalachensis | 2022-05-03 | CAM-NSF-Lichen Genomics Team; Allen, Jessica L. | USA: Shenandoah National Park, Central District, Bettys Rock, ~0.3 mi N of Crescent Rock, upper W-facing slopes of Timber Hollow | New York State Museum | 38.56753 N 78.38223 W | NYSM0000194405 |
| PUAP_TYS_02a | Punctelia appalachensis | 2022-05-03 | CAM-NSF-Lichen Genomics Team; Allen, Jessica L. | USA: Shenandoah National Park, Central District, Bettys Rock, ~0.3 mi N of Crescent Rock, upper W-facing slopes of Timber Hollow | New York State Museum | 38.56753 N 78.38223 W | NYSM0000194404 |
| PUAP_TYS_01c | Punctelia appalachensis | 2022-05-03 | CAM-NSF-Lichen Genomics Team; Allen, Jessica L. | USA: Shenandoah National Park, Central District, Bettys Rock, ~0.3 mi N of Crescent Rock, upper W-facing slopes of Timber Hollow | New York State Museum | 38.56753 N 78.38223 W | NYSM0000194403 |
| PUAP_TYS_01b | Punctelia appalachensis | 2022-05-03 | CAM-NSF-Lichen Genomics Team; Allen, Jessica L. | USA: Shenandoah National Park, Central District, Bettys Rock, ~0.3 mi N of Crescent Rock, upper W-facing slopes of Timber Hollow | New York State Museum | 38.56753 N 78.38223 W | NYSM0000194402 |
| PUAP_TYS_01a | Punctelia appalachensis | 2022-05-03 | CAM-NSF-Lichen Genomics Team; Allen, Jessica L. | USA: Shenandoah National Park, Central District, Bettys Rock, ~0.3 mi N of Crescent Rock, upper W-facing slopes of Timber Hollow | New York State Museum | 38.56753 N 78.38223 W | NYSM0000194401 |

| | | | | | | | |
|---|---|---|---|---|---|---|---|
| PUAP_TRI_16c | Punctelia appalachensis | 2022-04-28 | CAM-NSF-Lichen Genomics Team; Allen, Jessica L. | USA: Monongahela National Forest, Shavers Mountain, E-facing slopes of Trillium Knob above Old Road Run, S of Glade Run headwaters, Allegheny Trail ~1.5 mi N of FSR27 | New York State Museum | 38.644182 N 79.83029 W | NYSM0000194400 |
| PUAP_TRI_16b | Punctelia appalachensis | 2022-04-28 | CAM-NSF-Lichen Genomics Team; Allen, Jessica L. | USA: Monongahela National Forest, Shavers Mountain, E-facing slopes of Trillium Knob above Old Road Run, S of Glade Run headwaters, Allegheny Trail ~1.5 mi N of FSR27 | New York State Museum | 38.644182 N 79.83029 W | NYSM0000194399 |
| PUAP_TRI_16a | Punctelia appalachensis | 2022-04-28 | CAM-NSF-Lichen Genomics Team; Allen, Jessica L. | USA: Monongahela National Forest, Shavers Mountain, E-facing slopes of Trillium Knob above Old Road Run, S of Glade Run headwaters, Allegheny Trail ~1.5 mi N of FSR27 | New York State Museum | 38.644182 N 79.83029 W | NYSM0000194398 |
| PUAP_TRI_15a | Punctelia appalachensis | 2022-04-28 | CAM-NSF-Lichen Genomics Team; Allen, Jessica L. | USA: Monongahela National Forest, Shavers Mountain, E-facing slopes of Trillium Knob above Old Road Run, S of Glade Run headwaters, Allegheny Trail ~1.5 mi N of FSR27 | New York State Museum | 38.644182 N 79.83029 W | NYSM0000194397 |
| PUAP_TRI_11b | Punctelia appalachensis | 2022-04-28 | CAM-NSF-Lichen Genomics Team; Allen, Jessica L. | USA: Monongahela National Forest, Shavers Mountain, E-facing slopes of Trillium Knob above Old Road Run, S of Glade Run headwaters, Allegheny Trail ~1.5 mi N of FSR27 | New York State Museum | 38.644182 N 79.83029 W | NYSM0000194396 |

| | | | | | | | |
|---|---|---|---|---|---|---|---|
| PUAP_TRI_09b | Punctelia appalachensis | 2022-04-28 | CAM-NSF-Lichen Genomics Team; Allen, Jessica L. | USA: Monongahela National Forest, Shavers Mountain, E-facing slopes of Trillium Knob above Old Road Run, S of Glade Run headwaters, Allegheny Trail ~1.5 mi N of FSR27 | New York State Museum | 38.644182 N 79.83029 W | NYSM0000194395 |
| PUAP_TRI_09a | Punctelia appalachensis | 2022-04-28 | CAM-NSF-Lichen Genomics Team; Allen, Jessica L. | USA: Monongahela National Forest, Shavers Mountain, E-facing slopes of Trillium Knob above Old Road Run, S of Glade Run headwaters, Allegheny Trail ~1.5 mi N of FSR27 | New York State Museum | 38.644182 N 79.83029 W | NYSM0000194394 |
| PUAP_TRI_08c | Punctelia appalachensis | 2022-04-28 | CAM-NSF-Lichen Genomics Team; Allen, Jessica L. | USA: Monongahela National Forest, Shavers Mountain, E-facing slopes of Trillium Knob above Old Road Run, S of Glade Run headwaters, Allegheny Trail ~1.5 mi N of FSR27 | New York State Museum | 38.644182 N 79.83029 W | NYSM0000194393 |
| PUAP_TRI_08b | Punctelia appalachensis | 2022-04-28 | CAM-NSF-Lichen Genomics Team; Allen, Jessica L. | USA: Monongahela National Forest, Shavers Mountain, E-facing slopes of Trillium Knob above Old Road Run, S of Glade Run headwaters, Allegheny Trail ~1.5 mi N of FSR27 | New York State Museum | 38.644182 N 79.83029 W | NYSM0000194392 |
| PUAP_TRI_08a | Punctelia appalachensis | 2022-04-28 | CAM-NSF-Lichen Genomics Team; Allen, Jessica L. | USA: Monongahela National Forest, Shavers Mountain, E-facing slopes of Trillium Knob above Old Road Run, S of Glade Run headwaters, Allegheny Trail ~1.5 mi N of FSR27 | New York State Museum | 38.644182 N 79.83029 W | NYSM0000194391 |

| | | | | | | | |
|---|---|---|---|---|---|---|---|
| PUAP_TRI_06a | Punctelia appalachensis | 2022-04-28 | CAM-NSF-Lichen Genomics Team; Allen, Jessica L. | USA: Monongahela National Forest, Shavers Mountain, E-facing slopes of Trillium Knob above Old Road Run, S of Glade Run headwaters, Allegheny Trail ~1.5 mi N of FSR27 | New York State Museum | 38.644182 N 79.83029 W | NYSM0000194390 |
| PUAP_SNR_11c | Punctelia appalachensis | 2022-04-12 | CAM-NSF-Lichen Genomics Team; Allen, Jessica L. | USA: E-slopes of Pine Mountain, Kentenia State Forest, S of Little Shepherd Trail/SR1679 between Mill Cliff Branch and Hi Lewis Branch | New York State Museum | 36.920858 N 83.230543 W | NYSM0000194389 |
| PUAP_SNR_11b | Punctelia appalachensis | 2022-04-12 | CAM-NSF-Lichen Genomics Team; Allen, Jessica L. | USA: E-slopes of Pine Mountain, Kentenia State Forest, S of Little Shepherd Trail/SR1679 between Mill Cliff Branch and Hi Lewis Branch | New York State Museum | 36.920858 N 83.230543 W | NYSM0000194388 |
| PUAP_SNR_11a | Punctelia appalachensis | 2022-04-12 | CAM-NSF-Lichen Genomics Team; Allen, Jessica L. | USA: E-slopes of Pine Mountain, Kentenia State Forest, S of Little Shepherd Trail/SR1679 between Mill Cliff Branch and Hi Lewis Branch | New York State Museum | 36.920858 N 83.230543 W | NYSM0000194919 |
| PUAP_SNR_03c | Punctelia appalachensis | 2022-04-12 | CAM-NSF-Lichen Genomics Team; Allen, Jessica L. | USA: E-slopes of Pine Mountain, Kentenia State Forest, S of Little Shepherd Trail/SR1679 between Mill Cliff Branch and Hi Lewis Branch | New York State Museum | 36.920858 N 83.230543 W | NYSM0000194387 |

| | | | | | | | |
|---|---|---|---|---|---|---|---|
| PUAP_SNR_03b | Punctelia appalachensis | 2022-04-12 | CAM-NSF-Lichen Genomics Team; Allen, Jessica L. | USA: E-slopes of Pine Mountain, Kentenia State Forest, S of Little Shepherd Trail/SR1679 between Mill Cliff Branch and Hi Lewis Branch | New York State Museum | 36.920858 N 83.230543 W | NYSM0000194386 |
| PUAP_SNR_03a | Punctelia appalachensis | 2022-04-12 | CAM-NSF-Lichen Genomics Team; Allen, Jessica L. | USA: E-slopes of Pine Mountain, Kentenia State Forest, S of Little Shepherd Trail/SR1679 between Mill Cliff Branch and Hi Lewis Branch | New York State Museum | 36.920858 N 83.230543 W | NYSM0000194385 |
| PUAP_SNR_02c | Punctelia appalachensis | 2022-04-12 | CAM-NSF-Lichen Genomics Team; Allen, Jessica L. | USA: E-slopes of Pine Mountain, Kentenia State Forest, S of Little Shepherd Trail/SR1679 between Mill Cliff Branch and Hi Lewis Branch | New York State Museum | 36.920858 N 83.230543 W | NYSM0000194384 |
| PUAP_SNR_02b | Punctelia appalachensis | 2022-04-12 | CAM-NSF-Lichen Genomics Team; Allen, Jessica L. | USA: E-slopes of Pine Mountain, Kentenia State Forest, S of Little Shepherd Trail/SR1679 between Mill Cliff Branch and Hi Lewis Branch | New York State Museum | 36.920858 N 83.230543 W | NYSM0000194383 |
| PUAP_SNR_02a | Punctelia appalachensis | 2022-04-12 | CAM-NSF-Lichen Genomics Team; Allen, Jessica L. | USA: E-slopes of Pine Mountain, Kentenia State Forest, S of Little Shepherd Trail/SR1679 between Mill Cliff Branch and Hi Lewis Branch | New York State Museum | 36.920858 N 83.230543 W | NYSM0000194382 |
| PUAP_RSK_04c | Punctelia appalachensis | 2022-04-26 | CAM-NSF-Lichen Genomics Team; Allen, Jessica L. | USA: Monongahela National Forest, S-facing slopes of Red Spruce Knob, N of Red Lick Mountain, 0.1 mi W of WV150 and FSR451 | New York State Museum | 38.323665 N 80.152203 W | NYSM0000194381 |

| | | | | | | | |
|---|---|---|---|---|---|---|---|
| PUAP_RSK_04b | Punctelia appalachensis | 2022-04-26 | CAM-NSF-Lichen Genomics Team; Allen, Jessica L. | USA: Monongahela National Forest, S-facing slopes of Red Spruce Knob, N of Red Lick Mountain, 0.1 mi W of WV150 and FSR451 | New York State Museum | 38.323665 N 80.152203 W | NYSM0000194380 |
| PUAP_RSK_04a | Punctelia appalachensis | 2022-04-26 | CAM-NSF-Lichen Genomics Team; Allen, Jessica L. | USA: Monongahela National Forest, S-facing slopes of Red Spruce Knob, N of Red Lick Mountain, 0.1 mi W of WV150 and FSR451 | New York State Museum | 38.323665 N 80.152203 W | NYSM0000194379 |
| PUAP_RSK_03c | Punctelia appalachensis | 2022-04-26 | CAM-NSF-Lichen Genomics Team; Allen, Jessica L. | USA: Monongahela National Forest, S-facing slopes of Red Spruce Knob, N of Red Lick Mountain, 0.1 mi W of WV150 and FSR451 | New York State Museum | 38.323665 N 80.152203 W | NYSM0000194378 |
| PUAP_RSK_03b | Punctelia appalachensis | 2022-04-26 | CAM-NSF-Lichen Genomics Team; Allen, Jessica L. | USA: Monongahela National Forest, S-facing slopes of Red Spruce Knob, N of Red Lick Mountain, 0.1 mi W of WV150 and FSR451 | New York State Museum | 38.323665 N 80.152203 W | NYSM0000194377 |
| PUAP_RSK_03a | Punctelia appalachensis | 2022-04-26 | CAM-NSF-Lichen Genomics Team; Allen, Jessica L. | USA: Monongahela National Forest, S-facing slopes of Red Spruce Knob, N of Red Lick Mountain, 0.1 mi W of WV150 and FSR451 | New York State Museum | 38.323665 N 80.152203 W | NYSM0000194376 |
| PUAP_RSK_02c | Punctelia appalachensis | 2022-04-26 | CAM-NSF-Lichen Genomics Team; Allen, Jessica L. | USA: Monongahela National Forest, S-facing slopes of Red Spruce Knob, N of Red Lick Mountain, 0.1 mi W of WV150 and FSR451 | New York State Museum | 38.323665 N 80.152203 W | NYSM0000194375 |

| | | | | | | | |
|---|---|---|---|---|---|---|---|
| PUAP_RSK_02b | Punctelia appalachensis | 2022-04-26 | CAM-NSF-Lichen Genomics Team; Allen, Jessica L. | USA: Monongahela National Forest, S-facing slopes of Red Spruce Knob, N of Red Lick Mountain, 0.1 mi W of WV150 and FSR451 | New York State Museum | 38.323665 N 80.152203 W | NYSM0000194374 |
| PUAP_RSK_02a | Punctelia appalachensis | 2022-04-26 | CAM-NSF-Lichen Genomics Team; Allen, Jessica L. | USA: Monongahela National Forest, S-facing slopes of Red Spruce Knob, N of Red Lick Mountain, 0.1 mi W of WV150 and FSR451 | New York State Museum | 38.323665 N 80.152203 W | NYSM0000194373 |
| PUAP_RSK_01b | Punctelia appalachensis | 2022-04-26 | CAM-NSF-Lichen Genomics Team; Allen, Jessica L. | USA: Monongahela National Forest, S-facing slopes of Red Spruce Knob, N of Red Lick Mountain, 0.1 mi W of WV150 and FSR451 | New York State Museum | 38.323665 N 80.152203 W | NYSM0000194372 |
| PUAP_RSK_01a | Punctelia appalachensis | 2022-04-26 | CAM-NSF-Lichen Genomics Team; Allen, Jessica L. | USA: Monongahela National Forest, S-facing slopes of Red Spruce Knob, N of Red Lick Mountain, 0.1 mi W of WV150 and FSR451 | New York State Museum | 38.323665 N 80.152203 W | NYSM0000194371 |
| PUAP_RON_04b | Punctelia appalachensis | 2022-04-19 | CAM-NSF-Lichen Genomics Team; Allen, Jessica L. | USA: Jefferson National Forest, Round Mountain, vicinity of summit, Round Mountain Rd./VA688, ~4 mi W of US52/Bastian | New York State Museum | 37.154716 N 81.194163 W | NYSM0000194370 |
| PUAP_RON_04a | Punctelia appalachensis | 2022-04-19 | CAM-NSF-Lichen Genomics Team; Allen, Jessica L. | USA: Jefferson National Forest, Round Mountain, vicinity of summit, Round Mountain Rd./VA688, ~4 mi W of US52/Bastian | New York State Museum | 37.154716 N 81.194163 W | NYSM0000194369 |

| | | | | | | | |
|---|---|---|---|---|---|---|---|
| PUAP_RON _03c | Punctelia appalache nsis | 2022-04-19 | CAM-NSF-Lichen Genomics Team; Allen, Jessica L. | USA: Jefferson National Forest, Round Mountain, vicinity of summit, Round Mountain Rd./VA688, ~4 mi W of US52/Bastian | New York State Museum | 37.154716 N 81.194163 W | NYSM0000194368 |
| PUAP_RON _03b | Punctelia appalache nsis | 2022-04-19 | CAM-NSF-Lichen Genomics Team; Allen, Jessica L. | USA: Jefferson National Forest, Round Mountain, vicinity of summit, Round Mountain Rd./VA688, ~4 mi W of US52/Bastian | New York State Museum | 37.154716 N 81.194163 W | NYSM0000194916 |
| PUAP_RON _03a | Punctelia appalache nsis | 2022-04-19 | CAM-NSF-Lichen Genomics Team; Allen, Jessica L. | USA: Jefferson National Forest, Round Mountain, vicinity of summit, Round Mountain Rd./VA688, ~4 mi W of US52/Bastian | New York State Museum | 37.154716 N 81.194163 W | NYSM0000194367 |
| PUAP_RON _02c | Punctelia appalache nsis | 2022-04-19 | CAM-NSF-Lichen Genomics Team; Allen, Jessica L. | USA: Jefferson National Forest, Round Mountain, vicinity of summit, Round Mountain Rd./VA688, ~4 mi W of US52/Bastian | New York State Museum | 37.154716 N 81.194163 W | NYSM0000194366 |
| PUAP_RON _02b | Punctelia appalache nsis | 2022-04-19 | CAM-NSF-Lichen Genomics Team; Allen, Jessica L. | USA: Jefferson National Forest, Round Mountain, vicinity of summit, Round Mountain Rd./VA688, ~4 mi W of US52/Bastian | New York State Museum | 37.154716 N 81.194163 W | NYSM0000194365 |
| PUAP_RON _02a | Punctelia appalache nsis | 2022-04-19 | CAM-NSF-Lichen Genomics Team; Allen, Jessica L. | USA: Jefferson National Forest, Round Mountain, vicinity of summit, Round Mountain Rd./VA688, ~4 mi W of US52/Bastian | New York State Museum | 37.154716 N 81.194163 W | NYSM0000194364 |

| | | | | | | | |
|---|---|---|---|---|---|---|---|
| PUAP_RON_01c | Punctelia appalachensis | 2022-04-19 | CAM-NSF-Lichen Genomics Team; Allen, Jessica L. | USA: Jefferson National Forest, Round Mountain, vicinity of summit, Round Mountain Rd./VA688, ~4 mi W of US52/Bastian | New York State Museum | 37.154716 N 81.194163 W | NYSM0000194363 |
| PUAP_RON_01b | Punctelia appalachensis | 2022-04-19 | CAM-NSF-Lichen Genomics Team; Allen, Jessica L. | USA: Jefferson National Forest, Round Mountain, vicinity of summit, Round Mountain Rd./VA688, ~4 mi W of US52/Bastian | New York State Museum | 37.154716 N 81.194163 W | NYSM0000194362 |
| PUAP_RON_01a | Punctelia appalachensis | 2022-04-19 | CAM-NSF-Lichen Genomics Team; Allen, Jessica L. | USA: Jefferson National Forest, Round Mountain, vicinity of summit, Round Mountain Rd./VA688, ~4 mi W of US52/Bastian | New York State Museum | 37.154716 N 81.194163 W | NYSM0000194915 |
| PUAP_NGA_09a | Punctelia appalachensis | 2022-04-28 | CAM-NSF-Lichen Genomics Team; Allen, Jessica L. | USA: Monongahela National Forest, Shavers Mountain, E-facing slopes of Gaudineer Knob, FSR27 ~1.25 mi N of jct w/ US250 | New York State Museum | 38.611691 N 79.841629 W | NYSM0000194361 |
| PUAP_NGA_08c | Punctelia appalachensis | 2022-04-28 | CAM-NSF-Lichen Genomics Team; Allen, Jessica L. | USA: Monongahela National Forest, Shavers Mountain, E-facing slopes of Gaudineer Knob, FSR27 ~1.25 mi N of jct w/ US250 | New York State Museum | 38.611691 N 79.841629 W | NYSM0000194914 |
| PUAP_NGA_08b | Punctelia appalachensis | 2022-04-28 | CAM-NSF-Lichen Genomics Team; Allen, Jessica L. | USA: Monongahela National Forest, Shavers Mountain, E-facing slopes of Gaudineer Knob, FSR27 ~1.25 mi N of jct w/ US250 | New York State Museum | 38.611691 N 79.841629 W | NYSM0000194360 |

| | | | | | | | |
|---|---|---|---|---|---|---|---|
| PUAP_NGA_08a | Punctelia appalachensis | 2022-04-28 | CAM-NSF-Lichen Genomics Team; Allen, Jessica L. | USA: Monongahela National Forest, Shavers Mountain, E-facing slopes of Gaudineer Knob, FSR27 ~1.25 mi N of jct w/ US250 | New York State Museum | 38.611691 N 79.841629 W | NYSM0000194359 |
| PUAP_NGA_07c | Punctelia appalachensis | 2022-04-28 | CAM-NSF-Lichen Genomics Team; Allen, Jessica L. | USA: Monongahela National Forest, Shavers Mountain, E-facing slopes of Gaudineer Knob, FSR27 ~1.25 mi N of jct w/ US250 | New York State Museum | 38.611691 N 79.841629 W | NYSM0000194358 |
| PUAP_NGA_07b | Punctelia appalachensis | 2022-04-28 | CAM-NSF-Lichen Genomics Team; Allen, Jessica L. | USA: Monongahela National Forest, Shavers Mountain, E-facing slopes of Gaudineer Knob, FSR27 ~1.25 mi N of jct w/ US250 | New York State Museum | 38.611691 N 79.841629 W | NYSM0000194913 |
| PUAP_NGA_07a | Punctelia appalachensis | 2022-04-28 | CAM-NSF-Lichen Genomics Team; Allen, Jessica L. | USA: Monongahela National Forest, Shavers Mountain, E-facing slopes of Gaudineer Knob, FSR27 ~1.25 mi N of jct w/ US250 | New York State Museum | 38.611691 N 79.841629 W | NYSM0000194357 |
| PUAP_NGA_04c | Punctelia appalachensis | 2022-04-28 | CAM-NSF-Lichen Genomics Team; Allen, Jessica L. | USA: Monongahela National Forest, Shavers Mountain, E-facing slopes of Gaudineer Knob, FSR27 ~1.25 mi N of jct w/ US250 | New York State Museum | 38.611691 N 79.841629 W | NYSM0000194356 |
| PUAP_NGA_04b | Punctelia appalachensis | 2022-04-28 | CAM-NSF-Lichen Genomics Team; Allen, Jessica L. | USA: Monongahela National Forest, Shavers Mountain, E-facing slopes of Gaudineer Knob, FSR27 ~1.25 mi N of jct w/ US250 | New York State Museum | 38.611691 N 79.841629 W | NYSM0000194355 |

| | | | | | | | |
|---|---|---|---|---|---|---|---|
| PUAP_NGA_04a | Punctelia appalachensis | 2022-04-28 | CAM-NSF-Lichen Genomics Team; Allen, Jessica L. | USA: Monongahela National Forest, Shavers Mountain, E-facing slopes of Gaudineer Knob, FSR27 ~1.25 mi N of jct w/ US250 | New York State Museum | 38.611691 N 79.841629 W | NYSM0000194354 |
| PUAP_LUE_08c | Punctelia appalachensis | 2022-04-26 | CAM-NSF-Lichen Genomics Team; Allen, Jessica L. | USA: Monongahela National Forest, Spruce Mountain, W-facing slopes below summit of Blue Knob, ~1 mi S of WV39 | New York State Museum | 38.175865 N 80.279981 W | NYSM0000194353 |
| PUAP_LUE_08b | Punctelia appalachensis | 2022-04-26 | CAM-NSF-Lichen Genomics Team; Allen, Jessica L. | USA: Monongahela National Forest, Spruce Mountain, W-facing slopes below summit of Blue Knob, ~1 mi S of WV39 | New York State Museum | 38.175865 N 80.279981 W | NYSM0000194352 |
| PUAP_LUE_08a | Punctelia appalachensis | 2022-04-26 | CAM-NSF-Lichen Genomics Team; Allen, Jessica L. | USA: Monongahela National Forest, Spruce Mountain, W-facing slopes below summit of Blue Knob, ~1 mi S of WV39 | New York State Museum | 38.175865 N 80.279981 W | NYSM0000194351 |
| PUAP_LUE_07a | Punctelia appalachensis | 2022-04-26 | CAM-NSF-Lichen Genomics Team; Allen, Jessica L. | USA: Monongahela National Forest, Spruce Mountain, W-facing slopes below summit of Blue Knob, ~1 mi S of WV39 | New York State Museum | 38.175865 N 80.279981 W | NYSM0000194350 |
| PUAP_LUE_04a | Punctelia appalachensis | 2022-04-26 | CAM-NSF-Lichen Genomics Team; Allen, Jessica L. | USA: Monongahela National Forest, Spruce Mountain, W-facing slopes below summit of Blue Knob, ~1 mi S of WV39 | New York State Museum | 38.175865 N 80.279981 W | NYSM0000194349 |

| | | | | | | | |
|---|---|---|---|---|---|---|---|
| PUAP_LUE_03a | Punctelia appalachensis | 2022-04-26 | CAM-NSF-Lichen Genomics Team; Allen, Jessica L. | USA: Monongahela National Forest, Spruce Mountain, W-facing slopes below summit of Blue Knob, ~1 mi S of WV39 | New York State Museum | 38.175865 N 80.279981 W | NYSM0000194348 |
| PUAP_LUE_02c | Punctelia appalachensis | 2022-04-26 | CAM-NSF-Lichen Genomics Team; Allen, Jessica L. | USA: Monongahela National Forest, Spruce Mountain, W-facing slopes below summit of Blue Knob, ~1 mi S of WV39 | New York State Museum | 38.175865 N 80.279981 W | NYSM0000194347 |
| PUAP_LUE_02b | Punctelia appalachensis | 2022-04-26 | CAM-NSF-Lichen Genomics Team; Allen, Jessica L. | USA: Monongahela National Forest, Spruce Mountain, W-facing slopes below summit of Blue Knob, ~1 mi S of WV39 | New York State Museum | 38.175865 N 80.279981 W | NYSM0000194346 |
| PUAP_LUE_02a | Punctelia appalachensis | 2022-04-26 | CAM-NSF-Lichen Genomics Team; Allen, Jessica L. | USA: Monongahela National Forest, Spruce Mountain, W-facing slopes below summit of Blue Knob, ~1 mi S of WV39 | New York State Museum | 38.175865 N 80.279981 W | NYSM0000194345 |
| PUAP_LUE_01c | Punctelia appalachensis | 2022-04-26 | CAM-NSF-Lichen Genomics Team; Allen, Jessica L. | USA: Monongahela National Forest, Spruce Mountain, W-facing slopes below summit of Blue Knob, ~1 mi S of WV39 | New York State Museum | 38.175865 N 80.279981 W | NYSM0000194344 |
| PUAP_LUE_01b | Punctelia appalachensis | 2022-04-26 | CAM-NSF-Lichen Genomics Team; Allen, Jessica L. | USA: Monongahela National Forest, Spruce Mountain, W-facing slopes below summit of Blue Knob, ~1 mi S of WV39 | New York State Museum | 38.175865 N 80.279981 W | NYSM0000194343 |

| | | | | | | | |
|---|---|---|---|---|---|---|---|
| PUAP_LUE_01a | Punctelia appalachensis | 2022-04-26 | CAM-NSF-Lichen Genomics Team; Allen, Jessica L. | USA: Monongahela National Forest, Spruce Mountain, W-facing slopes below summit of Blue Knob, ~1 mi S of WV39 | New York State Museum | 38.175865 N 80.279981 W | NYSM0000194342 |
| PUAP_LEM_09c | Punctelia appalachensis | 2022-04-13 | CAM-NSF-Lichen Genomics Team; Allen, Jessica L. | USA: Pine Mountain, Bad Branch Nature Preserve, W headwaters of Bad Branch, Pine Mountain Trail ~2.2 mi E of US119 | New York State Museum | 37.09032 N 82.77928 W | NYSM0000194341 |
| PUAP_LEM_09b | Punctelia appalachensis | 2022-04-13 | CAM-NSF-Lichen Genomics Team; Allen, Jessica L. | USA: Pine Mountain, Bad Branch Nature Preserve, W headwaters of Bad Branch, Pine Mountain Trail ~2.2 mi E of US119 | New York State Museum | 37.09032 N 82.77928 W | NYSM0000194340 |
| PUAP_LEM_09a | Punctelia appalachensis | 2022-04-13 | CAM-NSF-Lichen Genomics Team; Allen, Jessica L. | USA: Pine Mountain, Bad Branch Nature Preserve, W headwaters of Bad Branch, Pine Mountain Trail ~2.2 mi E of US119 | New York State Museum | 37.09032 N 82.77928 W | NYSM0000194339 |
| PUAP_LEM_04a | Punctelia appalachensis | 2022-04-13 | CAM-NSF-Lichen Genomics Team; Allen, Jessica L. | USA: Pine Mountain, Bad Branch Nature Preserve, W headwaters of Bad Branch, Pine Mountain Trail ~2.2 mi E of US119 | New York State Museum | 37.09032 N 82.77928 W | NYSM0000194338 |
| PUAP_KLE_04c | Punctelia appalachensis | 2022-04-24 | CAM-NSF-Lichen Genomics Team; Allen, Jessica L. | USA: Monongahela National Forest, Bickle Knob, saddle and W-facing slopes below observation tower, terminus of FSR91A | New York State Museum | 38.93557 N 79.730504 W | NYSM0000194337 |

| | | | | | | | |
|---|---|---|---|---|---|---|---|
| PUAP_KLE_04b | Punctelia appalachensis | 2022-04-24 | CAM-NSF-Lichen Genomics Team; Allen, Jessica L. | USA: Monongahela National Forest, Bickle Knob, saddle and W-facing slopes below observation tower, terminus of FSR91A | New York State Museum | 38.93557 N 79.730504 W | NYSM0000194336 |
| PUAP_KLE_04a | Punctelia appalachensis | 2022-04-24 | CAM-NSF-Lichen Genomics Team; Allen, Jessica L. | USA: Monongahela National Forest, Bickle Knob, saddle and W-facing slopes below observation tower, terminus of FSR91A | New York State Museum | 38.93557 N 79.730504 W | NYSM0000194335 |
| PUAP_KLE_03a | Punctelia appalachensis | 2022-04-24 | CAM-NSF-Lichen Genomics Team; Allen, Jessica L. | USA: Monongahela National Forest, Bickle Knob, saddle and W-facing slopes below observation tower, terminus of FSR91A | New York State Museum | 38.93557 N 79.730504 W | NYSM0000194334 |
| PUAP_KLE_02c | Punctelia appalachensis | 2022-04-24 | CAM-NSF-Lichen Genomics Team; Allen, Jessica L. | USA: Monongahela National Forest, Bickle Knob, saddle and W-facing slopes below observation tower, terminus of FSR91A | New York State Museum | 38.93557 N 79.730504 W | NYSM0000194910 |
| PUAP_KLE_02b | Punctelia appalachensis | 2022-04-24 | CAM-NSF-Lichen Genomics Team; Allen, Jessica L. | USA: Monongahela National Forest, Bickle Knob, saddle and W-facing slopes below observation tower, terminus of FSR91A | New York State Museum | 38.93557 N 79.730504 W | NYSM0000194333 |
| PUAP_KLE_02a | Punctelia appalachensis | 2022-04-24 | CAM-NSF-Lichen Genomics Team; Allen, Jessica L. | USA: Monongahela National Forest, Bickle Knob, saddle and W-facing slopes below observation tower, terminus of FSR91A | New York State Museum | 38.93557 N 79.730504 W | NYSM0000194332 |

| | | | | | | | |
|---|---|---|---|---|---|---|---|
| PUAP_KLE _01c | Punctelia appalachensis | 2022-04-24 | CAM-NSF-Lichen Genomics Team; Allen, Jessica L. | USA: Monongahela National Forest, Bickle Knob, saddle and W-facing slopes below observation tower, terminus of FSR91A | New York State Museum | 38.93557 N 79.730504 W | NYSM0000194331 |
| PUAP_KLE _01b | Punctelia appalachensis | 2022-04-24 | CAM-NSF-Lichen Genomics Team; Allen, Jessica L. | USA: Monongahela National Forest, Bickle Knob, saddle and W-facing slopes below observation tower, terminus of FSR91A | New York State Museum | 38.93557 N 79.730504 W | NYSM0000194330 |
| PUAP_KLE _01a | Punctelia appalachensis | 2022-04-24 | CAM-NSF-Lichen Genomics Team; Allen, Jessica L. | USA: Monongahela National Forest, Bickle Knob, saddle and W-facing slopes below observation tower, terminus of FSR91A | New York State Museum | 38.93557 N 79.730504 W | NYSM0000194329 |
| PUAP_HA W_07c | Punctelia appalachensis | 2022-05-03 | CAM-NSF-Lichen Genomics Team; Allen, Jessica L. | USA: Shenandoah National Park, Central District, SW slopes of Hawksbill Mountain, between Lower Hawksbill Trail and Hawksbill Fire Road ~0.7 mi W of Hawksbill Gap | New York State Museum | 38.55397 N 78.39687 W | NYSM0000194328 |
| PUAP_HA W_07b | Punctelia appalachensis | 2022-05-03 | CAM-NSF-Lichen Genomics Team; Allen, Jessica L. | USA: Shenandoah National Park, Central District, SW slopes of Hawksbill Mountain, between Lower Hawksbill Trail and Hawksbill Fire Road ~0.7 mi W of Hawksbill Gap | New York State Museum | 38.55397 N 78.39687 W | NYSM0000194327 |

| | | | | | | | |
|---|---|---|---|---|---|---|---|
| PUAP_HAW_07a | Punctelia appalachensis | 2022-05-03 | CAM-NSF-Lichen Genomics Team; Allen, Jessica L. | USA: Shenandoah National Park, Central District, SW slopes of Hawksbill Mountain, between Lower Hawksbill Trail and Hawksbill Fire Road ~0.7 mi W of Hawksbill Gap | New York State Museum | 38.55397 N 78.39687 W | NYSM0000194326 |
| PUAP_HAW_03c | Punctelia appalachensis | 2022-05-03 | CAM-NSF-Lichen Genomics Team; Allen, Jessica L. | USA: Shenandoah National Park, Central District, SW slopes of Hawksbill Mountain, between Lower Hawksbill Trail and Hawksbill Fire Road ~0.7 mi W of Hawksbill Gap | New York State Museum | 38.55397 N 78.39687 W | NYSM0000194325 |
| PUAP_HAW_03b | Punctelia appalachensis | 2022-05-03 | CAM-NSF-Lichen Genomics Team; Allen, Jessica L. | USA: Shenandoah National Park, Central District, SW slopes of Hawksbill Mountain, between Lower Hawksbill Trail and Hawksbill Fire Road ~0.7 mi W of Hawksbill Gap | New York State Museum | 38.55397 N 78.39687 W | NYSM0000194324 |
| PUAP_HAW_03a | Punctelia appalachensis | 2022-05-03 | CAM-NSF-Lichen Genomics Team; Allen, Jessica L. | USA: Shenandoah National Park, Central District, SW slopes of Hawksbill Mountain, between Lower Hawksbill Trail and Hawksbill Fire Road ~0.7 mi W of Hawksbill Gap | New York State Museum | 38.55397 N 78.39687 W | NYSM0000194323 |
| PUAP_HAW_01c | Punctelia appalachensis | 2022-05-03 | CAM-NSF-Lichen Genomics Team; Allen, Jessica L. | USA: Shenandoah National Park, Central District, SW slopes of Hawksbill Mountain, between Lower Hawksbill Trail and Hawksbill Fire Road ~0.7 mi W of Hawksbill Gap | New York State Museum | 38.55397 N 78.39687 W | NYSM0000194322 |

| | | | | | | | |
|---|---|---|---|---|---|---|---|
| PUAP_HAW_01b | Punctelia appalachensis | 2022-05-03 | CAM-NSF-Lichen Genomics Team; Allen, Jessica L. | USA: Shenandoah National Park, Central District, SW slopes of Hawksbill Mountain, between Lower Hawksbill Trail and Hawksbill Fire Road ~0.7 mi W of Hawksbill Gap | New York State Museum | 38.55397 N 78.39687 W | NYSM0000194321 |
| PUAP_HAW_01a | Punctelia appalachensis | 2022-05-03 | CAM-NSF-Lichen Genomics Team; Allen, Jessica L. | USA: Shenandoah National Park, Central District, SW slopes of Hawksbill Mountain, between Lower Hawksbill Trail and Hawksbill Fire Road ~0.7 mi W of Hawksbill Gap | New York State Museum | 38.55397 N 78.39687 W | NYSM0000194320 |
| PUAP_GOS_12c | Punctelia appalachensis | 2022-04-11 | CAM-NSF-Lichen Genomics Team; Allen, Jessica L. | USA: E-slopes of Pine Mountain, Kentenia State Forest, vicinity of Goss Park, E of Little Shepherd Trail/SR1679 at jct w/ Putney Lookout Tower Rd., between Spruce Ridge Branch and Hi Lewis Branch | New York State Museum | 36.9243 N 83.22419 W | NYSM0000194912 |
| PUAP_GOS_12b | Punctelia appalachensis | 2022-04-11 | CAM-NSF-Lichen Genomics Team; Allen, Jessica L. | USA: E-slopes of Pine Mountain, Kentenia State Forest, vicinity of Goss Park, E of Little Shepherd Trail/SR1679 at jct w/ Putney Lookout Tower Rd., between Spruce Ridge Branch and Hi Lewis Branch | New York State Museum | 36.9243 N 83.22419 W | NYSM0000194319 |
| PUAP_GOS_12a | Punctelia appalachensis | 2022-04-11 | CAM-NSF-Lichen Genomics Team; Allen, Jessica L. | USA: E-slopes of Pine Mountain, Kentenia State Forest, vicinity of Goss Park, E of Little Shepherd Trail/SR1679 at jct w/ Putney Lookout Tower Rd., between | New York State Museum | 36.9243 N 83.22419 W | NYSM0000194318 |

| | | | | | | | |
|---|---|---|---|---|---|---|---|
| | | | | Spruce Ridge Branch and Hi Lewis Branch | | | |
| PUAP_GOS_06b | Punctelia appalachensis | 2022-04-11 | CAM-NSF-Lichen Genomics Team; Allen, Jessica L. | USA: E-slopes of Pine Mountain, Kentenia State Forest, vicinity of Goss Park, E of Little Shepherd Trail/SR1679 at jct w/ Putney Lookout Tower Rd., between Spruce Ridge Branch and Hi Lewis Branch | New York State Museum | 36.9243 N 83.22419 W | NYSM0000194317 |
| PUAP_GOS_06a | Punctelia appalachensis | 2022-04-11 | CAM-NSF-Lichen Genomics Team; Allen, Jessica L. | USA: E-slopes of Pine Mountain, Kentenia State Forest, vicinity of Goss Park, E of Little Shepherd Trail/SR1679 at jct w/ Putney Lookout Tower Rd., between Spruce Ridge Branch and Hi Lewis Branch | New York State Museum | 36.9243 N 83.22419 W | NYSM0000194316 |
| PUAP_GOS_03c | Punctelia appalachensis | 2022-04-11 | CAM-NSF-Lichen Genomics Team; Allen, Jessica L. | USA: E-slopes of Pine Mountain, Kentenia State Forest, vicinity of Goss Park, E of Little Shepherd Trail/SR1679 at jct w/ Putney Lookout Tower Rd., between Spruce Ridge Branch and Hi Lewis Branch | New York State Museum | 36.9243 N 83.22419 W | NYSM0000194315 |
| PUAP_GOS_03b | Punctelia appalachensis | 2022-04-11 | CAM-NSF-Lichen Genomics Team; Allen, Jessica L. | USA: E-slopes of Pine Mountain, Kentenia State Forest, vicinity of Goss Park, E of Little Shepherd Trail/SR1679 at jct w/ Putney Lookout Tower Rd., between | New York State Museum | 36.9243 N 83.22419 W | NYSM0000194314 |

| | | | | | | | |
|---|---|---|---|---|---|---|---|
| | | | | Spruce Ridge Branch and Hi Lewis Branch | | | |
| PUAP_GOS_03a | Punctelia appalachensis | 2022-04-11 | CAM-NSF-Lichen Genomics Team; Allen, Jessica L. | USA: E-slopes of Pine Mountain, Kentenia State Forest, vicinity of Goss Park, E of Little Shepherd Trail/SR1679 at jct w/ Putney Lookout Tower Rd., between Spruce Ridge Branch and Hi Lewis Branch | New York State Museum | 36.9243 N 83.22419 W | NYSM0000194911 |
| PUAP_GOS_02b | Punctelia appalachensis | 2022-04-11 | CAM-NSF-Lichen Genomics Team; Allen, Jessica L. | USA: E-slopes of Pine Mountain, Kentenia State Forest, vicinity of Goss Park, E of Little Shepherd Trail/SR1679 at jct w/ Putney Lookout Tower Rd., between Spruce Ridge Branch and Hi Lewis Branch | New York State Museum | 36.9243 N 83.22419 W | NYSM0000194313 |
| PUAP_GOS_02a | Punctelia appalachensis | 2022-04-11 | CAM-NSF-Lichen Genomics Team; Allen, Jessica L. | USA: E-slopes of Pine Mountain, Kentenia State Forest, vicinity of Goss Park, E of Little Shepherd Trail/SR1679 at jct w/ Putney Lookout Tower Rd., between Spruce Ridge Branch and Hi Lewis Branch | New York State Museum | 36.9243 N 83.22419 W | NYSM0000194312 |
| PUAP_DBG_14A | Punctelia appalachensis | 2021-10-23 | CAM-NSF-Lichen Genomics Team; Allen, Jessica L. | USA: Great Smoky Mountains National Park, Cataloochee Divide, McClue Ridge, Double Gap Trail ~0.1 mi W of Double Gap | New York State Museum | 35.57191 N 83.1035 W | NYSM0000194311 |

| | | | | | | | |
|---|---|---|---|---|---|---|---|
| PUAP_DBG_12A2 | Punctelia appalachensis | 2021-10-23 | CAM-NSF-Lichen Genomics Team; Allen, Jessica L. | USA: Great Smoky Mountains National Park, Cataloochee Divide, McClue Ridge, Double Gap Trail ~0.1 mi W of Double Gap | New York State Museum | 35.57191 N 83.1035 W | NYSM0000194310 |
| PUAP_DBG_12A1 | Punctelia appalachensis | 2021-10-23 | CAM-NSF-Lichen Genomics Team; Allen, Jessica L. | USA: Great Smoky Mountains National Park, Cataloochee Divide, McClue Ridge, Double Gap Trail ~0.1 mi W of Double Gap | New York State Museum | 35.57191 N 83.1035 W | NYSM0000194309 |
| PUAP_DBG_11A | Punctelia appalachensis | 2021-10-23 | CAM-NSF-Lichen Genomics Team; Allen, Jessica L. | USA: Great Smoky Mountains National Park, Cataloochee Divide, McClue Ridge, Double Gap Trail ~0.1 mi W of Double Gap | New York State Museum | 35.57191 N 83.1035 W | NYSM0000194308 |
| PUAP_DBG_10A | Punctelia appalachensis | 2021-10-23 | CAM-NSF-Lichen Genomics Team; Allen, Jessica L. | USA: Great Smoky Mountains National Park, Cataloochee Divide, McClue Ridge, Double Gap Trail ~0.1 mi W of Double Gap | New York State Museum | 35.57191 N 83.1035 W | NYSM0000194307 |
| PUAP_DBG_09A | Punctelia appalachensis | 2021-10-23 | CAM-NSF-Lichen Genomics Team; Allen, Jessica L. | USA: Great Smoky Mountains National Park, Cataloochee Divide, McClue Ridge, Double Gap Trail ~0.1 mi W of Double Gap | New York State Museum | 35.57191 N 83.1035 W | NYSM0000194306 |
| PUAP_DBG_08A | Punctelia appalachensis | 2021-10-23 | CAM-NSF-Lichen Genomics Team; Allen, Jessica L. | USA: Great Smoky Mountains National Park, Cataloochee Divide, McClue Ridge, Double Gap Trail ~0.1 mi W of Double Gap | New York State Museum | 35.57191 N 83.1035 W | NYSM0000194305 |

| | | | | | | | |
|---|---|---|---|---|---|---|---|
| PUAP_DBG_07A | Punctelia appalachensis | 2021-10-23 | CAM-NSF-Lichen Genomics Team; Allen, Jessica L. | USA: Great Smoky Mountains National Park, Cataloochee Divide, McClue Ridge, Double Gap Trail ~0.1 mi W of Double Gap | New York State Museum | 35.57191 N 83.1035 W | NYSM0000194304 |
| PUAP_DBG_06A | Punctelia appalachensis | 2021-10-23 | CAM-NSF-Lichen Genomics Team; Allen, Jessica L. | USA: Great Smoky Mountains National Park, Cataloochee Divide, McClue Ridge, Double Gap Trail ~0.1 mi W of Double Gap | New York State Museum | 35.57191 N 83.1035 W | NYSM0000194303 |
| PUAP_DBG_03A | Punctelia appalachensis | 2021-10-23 | CAM-NSF-Lichen Genomics Team; Allen, Jessica L. | USA: Great Smoky Mountains National Park, Cataloochee Divide, McClue Ridge, Double Gap Trail ~0.1 mi W of Double Gap | New York State Museum | 35.57191 N 83.1035 W | NYSM0000194302 |
| PUAP_DAD_06b | Punctelia appalachensis | 2022-08-10 | CAM-NSF-Lichen Genomics Team; Denzer, Eli | USA: Grandfather Mountain State Park, Grandfather Mountain, N-slopes N of Profile Trail, ~0.3 mi W of jct w/ Grandfather Trail | New York State Museum | 36.11211 N 81.81832 W | NYSM0000194909 |
| PUAP_DAD_06a | Punctelia appalachensis | 2022-08-10 | CAM-NSF-Lichen Genomics Team; Denzer, Eli | USA: Grandfather Mountain State Park, Grandfather Mountain, N-slopes N of Profile Trail, ~0.3 mi W of jct w/ Grandfather Trail | New York State Museum | 36.11211 N 81.81832 W | NYSM0000194908 |
| PUAP_DAD_05c | Punctelia appalachensis | 2022-08-10 | CAM-NSF-Lichen Genomics Team; Denzer, Eli | USA: Grandfather Mountain State Park, Grandfather Mountain, N-slopes N of Profile Trail, ~0.3 mi W of jct w/ Grandfather Trail | New York State Museum | 36.11211 N 81.81832 W | NYSM0000194907 |

| | | | | | | | |
|---|---|---|---|---|---|---|---|
| PUAP_DAD_05b | Punctelia appalachensis | 2022-08-10 | CAM-NSF-Lichen Genomics Team; Denzer, Eli | USA: Grandfather Mountain State Park, Grandfather Mountain, N-slopes N of Profile Trail, ~0.3 mi W of jct w/ Grandfather Trail | New York State Museum | 36.11211 N 81.81832 W | NYSM0000194906 |
| PUAP_DAD_05a | Punctelia appalachensis | 2022-08-10 | CAM-NSF-Lichen Genomics Team; Denzer, Eli | USA: Grandfather Mountain State Park, Grandfather Mountain, N-slopes N of Profile Trail, ~0.3 mi W of jct w/ Grandfather Trail | New York State Museum | 36.11211 N 81.81832 W | NYSM0000194905 |
| PUAP_BUK_34A | Punctelia appalachensis | 2021-10-24 | CAM-NSF-Lichen Genomics Team; Allen, Jessica L. | USA: Great Smoky Mountains National Park, Cataloochee Divide, N-slope of Little Bald Knob, Hemphill Bald Trail ~3 mi E of Polls Gap | New York State Museum | 35.552276 N 83.120161 W | NYSM0000194296 |
| PUAP_BUK_33A | Punctelia appalachensis | 2021-10-24 | CAM-NSF-Lichen Genomics Team; Allen, Jessica L. | USA: Great Smoky Mountains National Park, Cataloochee Divide, N-slope of Little Bald Knob, Hemphill Bald Trail ~3 mi E of Polls Gap | New York State Museum | 35.552276 N 83.120161 W | NYSM0000194295 |
| PUAP_BUK_32A | Punctelia appalachensis | 2021-10-24 | CAM-NSF-Lichen Genomics Team; Allen, Jessica L. | USA: Great Smoky Mountains National Park, Cataloochee Divide, N-slope of Little Bald Knob, Hemphill Bald Trail ~3 mi E of Polls Gap | New York State Museum | 35.552276 N 83.120161 W | NYSM0000194294 |
| PUAP_BUK_31A | Punctelia appalachensis | 2021-10-24 | CAM-NSF-Lichen Genomics Team; Allen, Jessica L. | USA: Great Smoky Mountains National Park, Cataloochee Divide, N-slope of Little Bald Knob, Hemphill Bald Trail ~3 mi E of Polls Gap | New York State Museum | 35.552276 N 83.120161 W | NYSM0000194293 |

| | | | | | | | |
|---|---|---|---|---|---|---|---|
| PUAP_BUK_30A | Punctelia appalachensis | 2021-10-24 | CAM-NSF-Lichen Genomics Team; Allen, Jessica L. | USA: Great Smoky Mountains National Park, Cataloochee Divide, N-slope of Little Bald Knob, Hemphill Bald Trail ~3 mi E of Polls Gap | New York State Museum | 35.552276 N 83.120161 W | NYSM0000194292 |
| PUAP_BUK_28A | Punctelia appalachensis | 2021-10-24 | CAM-NSF-Lichen Genomics Team; Allen, Jessica L. | USA: Great Smoky Mountains National Park, Cataloochee Divide, N-slope of Little Bald Knob, Hemphill Bald Trail ~3 mi E of Polls Gap | New York State Museum | 35.552276 N 83.120161 W | NYSM0000194291 |
| PUAP_BUK_11A | Punctelia appalachensis | 2021-10-24 | CAM-NSF-Lichen Genomics Team; Allen, Jessica L. | USA: Great Smoky Mountains National Park, Cataloochee Divide, N-slope of Little Bald Knob, Hemphill Bald Trail ~3 mi E of Polls Gap | New York State Museum | 35.552276 N 83.120161 W | NYSM0000194290 |
| PUAP_BUK_10A | Punctelia appalachensis | 2021-10-24 | CAM-NSF-Lichen Genomics Team; Allen, Jessica L. | USA: Great Smoky Mountains National Park, Cataloochee Divide, N-slope of Little Bald Knob, Hemphill Bald Trail ~3 mi E of Polls Gap | New York State Museum | 35.552276 N 83.120161 W | NYSM0000194289 |
| PUAP_BUK_04A | Punctelia appalachensis | 2021-10-24 | CAM-NSF-Lichen Genomics Team; Allen, Jessica L. | USA: Great Smoky Mountains National Park, Cataloochee Divide, N-slope of Little Bald Knob, Hemphill Bald Trail ~3 mi E of Polls Gap | New York State Museum | 35.552276 N 83.120161 W | NYSM0000194288 |
| PUAP_BUK_02A | Punctelia appalachensis | 2021-10-24 | CAM-NSF-Lichen Genomics Team; Allen, Jessica L. | USA: Great Smoky Mountains National Park, Cataloochee Divide, N-slope of Little Bald Knob, Hemphill Bald Trail ~3 mi E of Polls Gap | New York State Museum | 35.552276 N 83.120161 W | NYSM0000194287 |

| | | | | | | | |
|---|---|---|---|---|---|---|---|
| PUAP_BRF_08c | Punctelia appalachensis | 2022-05-03 | CAM-NSF-Lichen Genomics Team; Allen, Jessica L. | USA: Shenandoah National Park, Central District, Bearfence Mountain, W-slopes below Appalachian Trail, 0.3 mi S of jct w/ spur trail to Skyline Drive/Bearfence Parking Area | New York State Museum | 38.4477 N 78.46635 W | NYSM0000194286 |
| PUAP_BRF_08b | Punctelia appalachensis | 2022-05-03 | CAM-NSF-Lichen Genomics Team; Allen, Jessica L. | USA: Shenandoah National Park, Central District, Bearfence Mountain, W-slopes below Appalachian Trail, 0.3 mi S of jct w/ spur trail to Skyline Drive/Bearfence Parking Area | New York State Museum | 38.4477 N 78.46635 W | NYSM0000194285 |
| PUAP_BRF_08a | Punctelia appalachensis | 2022-05-03 | CAM-NSF-Lichen Genomics Team; Allen, Jessica L. | USA: Shenandoah National Park, Central District, Bearfence Mountain, W-slopes below Appalachian Trail, 0.3 mi S of jct w/ spur trail to Skyline Drive/Bearfence Parking Area | New York State Museum | 38.4477 N 78.46635 W | NYSM0000194284 |
| PUAP_BRF_07d | Punctelia appalachensis | 2022-05-03 | CAM-NSF-Lichen Genomics Team; Allen, Jessica L. | USA: Shenandoah National Park, Central District, Bearfence Mountain, W-slopes below Appalachian Trail, 0.3 mi S of jct w/ spur trail to Skyline Drive/Bearfence Parking Area | New York State Museum | 38.4477 N 78.46635 W | NYSM0000194283 |
| PUAP_BRF_07c | Punctelia appalachensis | 2022-05-03 | CAM-NSF-Lichen Genomics Team; Allen, Jessica L. | USA: Shenandoah National Park, Central District, Bearfence Mountain, W-slopes below Appalachian Trail, 0.3 mi S of jct w/ spur trail to Skyline Drive/Bearfence Parking Area | New York State Museum | 38.4477 N 78.46635 W | NYSM0000194904 |

| | | | | | | | |
|---|---|---|---|---|---|---|---|
| PUAP_BRF_07b | Punctelia appalachensis | 2022-05-03 | CAM-NSF-Lichen Genomics Team; Allen, Jessica L. | USA: Shenandoah National Park, Central District, Bearfence Mountain, W-slopes below Appalachian Trail, 0.3 mi S of jct w/ spur trail to Skyline Drive/Bearfence Parking Area | New York State Museum | 38.4477 N 78.46635 W | NYSM0000194282 |
| PUAP_BRF_07a | Punctelia appalachensis | 2022-05-03 | CAM-NSF-Lichen Genomics Team; Allen, Jessica L. | USA: Shenandoah National Park, Central District, Bearfence Mountain, W-slopes below Appalachian Trail, 0.3 mi S of jct w/ spur trail to Skyline Drive/Bearfence Parking Area | New York State Museum | 38.4477 N 78.46635 W | NYSM0000194281 |
| PUAP_BRF_05b | Punctelia appalachensis | 2022-05-03 | CAM-NSF-Lichen Genomics Team; Allen, Jessica L. | USA: Shenandoah National Park, Central District, Bearfence Mountain, W-slopes below Appalachian Trail, 0.3 mi S of jct w/ spur trail to Skyline Drive/Bearfence Parking Area | New York State Museum | 38.4477 N 78.46635 W | NYSM0000194280 |
| PUAP_BRF_05a | Punctelia appalachensis | 2022-05-03 | CAM-NSF-Lichen Genomics Team; Allen, Jessica L. | USA: Shenandoah National Park, Central District, Bearfence Mountain, W-slopes below Appalachian Trail, 0.3 mi S of jct w/ spur trail to Skyline Drive/Bearfence Parking Area | New York State Museum | 38.4477 N 78.46635 W | NYSM0000194279 |
| PUAP_BRF_01a | Punctelia appalachensis | 2022-05-03 | CAM-NSF-Lichen Genomics Team; Allen, Jessica L. | USA: Shenandoah National Park, Central District, Bearfence Mountain, W-slopes below Appalachian Trail, 0.3 mi S of jct w/ spur trail to Skyline Drive/Bearfence Parking Area | New York State Museum | 38.4477 N 78.46635 W | NYSM0000194278 |

| | | | | | | | |
|---|---|---|---|---|---|---|---|
| PUAP_BLU_05c | Punctelia appalachensis | 2022-04-13 | CAM-NSF-Lichen Genomics Team; Allen, Jessica L. | USA: Pine Mountain, Bad Branch Nature Preserve, headwaters of Presley House Branch, Pine Mountain Trail ~1.5 mi E of US119 | New York State Museum | 37.08482 N 82.79308 W | NYSM0000194277 |
| PUAP_BLU_05b | Punctelia appalachensis | 2022-04-13 | CAM-NSF-Lichen Genomics Team; Allen, Jessica L. | USA: Pine Mountain, Bad Branch Nature Preserve, headwaters of Presley House Branch, Pine Mountain Trail ~1.5 mi E of US119 | New York State Museum | 37.08482 N 82.79308 W | NYSM0000194276 |
| PUAP_BLU_05a | Punctelia appalachensis | 2022-04-13 | CAM-NSF-Lichen Genomics Team; Allen, Jessica L. | USA: Pine Mountain, Bad Branch Nature Preserve, headwaters of Presley House Branch, Pine Mountain Trail ~1.5 mi E of US119 | New York State Museum | 37.08482 N 82.79308 W | NYSM0000194275 |
| PUAP_BLU_04a | Punctelia appalachensis | 2022-04-13 | CAM-NSF-Lichen Genomics Team; Allen, Jessica L. | USA: Pine Mountain, Bad Branch Nature Preserve, headwaters of Presley House Branch, Pine Mountain Trail ~1.5 mi E of US119 | New York State Museum | 37.08482 N 82.79308 W | NYSM0000194274 |
| PUAP_BLU_03b | Punctelia appalachensis | 2022-04-13 | CAM-NSF-Lichen Genomics Team; Allen, Jessica L. | USA: Pine Mountain, Bad Branch Nature Preserve, headwaters of Presley House Branch, Pine Mountain Trail ~1.5 mi E of US119 | New York State Museum | 37.08482 N 82.79308 W | NYSM0000194273 |
| PUAP_BLU_03a | Punctelia appalachensis | 2022-04-13 | CAM-NSF-Lichen Genomics Team; Allen, Jessica L. | USA: Pine Mountain, Bad Branch Nature Preserve, headwaters of Presley House Branch, Pine Mountain Trail ~1.5 mi E of US119 | New York State Museum | 37.08482 N 82.79308 W | NYSM0000194272 |

| | | | | | | | |
|---|---|---|---|---|---|---|---|
| PUAP_BLU_02a | Punctelia appalachensis | 2022-04-13 | CAM-NSF-Lichen Genomics Team; Allen, Jessica L. | USA: Pine Mountain, Bad Branch Nature Preserve, headwaters of Presley House Branch, Pine Mountain Trail ~1.5 mi E of US119 | New York State Museum | 37.08482 N 82.79308 W | NYSM0000194271 |
| PUAP_BLU_01c | Punctelia appalachensis | 2022-04-13 | CAM-NSF-Lichen Genomics Team; Allen, Jessica L. | USA: Pine Mountain, Bad Branch Nature Preserve, headwaters of Presley House Branch, Pine Mountain Trail ~1.5 mi E of US119 | New York State Museum | 37.08482 N 82.79308 W | NYSM0000194270 |
| PUAP_BLU_01b | Punctelia appalachensis | 2022-04-13 | CAM-NSF-Lichen Genomics Team; Allen, Jessica L. | USA: Pine Mountain, Bad Branch Nature Preserve, headwaters of Presley House Branch, Pine Mountain Trail ~1.5 mi E of US119 | New York State Museum | 37.08482 N 82.79308 W | NYSM0000194269 |
| PUAP_BLU_01a | Punctelia appalachensis | 2022-04-13 | CAM-NSF-Lichen Genomics Team; Allen, Jessica L. | USA: Pine Mountain, Bad Branch Nature Preserve, headwaters of Presley House Branch, Pine Mountain Trail ~1.5 mi E of US119 | New York State Museum | 37.08482 N 82.79308 W | NYSM0000194268 |
| PUAP_BFF_16A | Punctelia appalachensis | 2021-10-26 | CAM-NSF-Lichen Genomics Team; Allen, Jessica L. | USA: Pisgah National Forest, Bald Mountains, N-Facing slopes of Bluff Mountain summit, NE of Appalachian Trail 1 mi E of Catpen Gap, 3.4 mi E of Lemon Gap | New York State Museum | 35.84115 N 82.90576 W | NYSM0000194267 |

| | | | | | | | |
|---|---|---|---|---|---|---|---|
| PUAP_BFF_14A | Punctelia appalachensis | 2021-10-26 | CAM-NSF-Lichen Genomics Team; Allen, Jessica L. | USA: Pisgah National Forest, Bald Mountains, N-Facing slopes of Bluff Mountain summit, NE of Appalachian Trail 1 mi E of Catpen Gap, 3.4 mi E of Lemon Gap | New York State Museum | 35.84115 N 82.90576 W | NYSM0000194266 |
| PUAP_BFF_13A | Punctelia appalachensis | 2021-10-26 | CAM-NSF-Lichen Genomics Team; Allen, Jessica L. | USA: Pisgah National Forest, Bald Mountains, N-Facing slopes of Bluff Mountain summit, NE of Appalachian Trail 1 mi E of Catpen Gap, 3.4 mi E of Lemon Gap | New York State Museum | 35.84115 N 82.90576 W | NYSM0000194265 |
| PUAP_BFF_10A | Punctelia appalachensis | 2021-10-26 | CAM-NSF-Lichen Genomics Team; Allen, Jessica L. | USA: Pisgah National Forest, Bald Mountains, N-Facing slopes of Bluff Mountain summit, NE of Appalachian Trail 1 mi E of Catpen Gap, 3.4 mi E of Lemon Gap | New York State Museum | 35.84115 N 82.90576 W | NYSM0000194264 |
| PUAP_BFF_08A | Punctelia appalachensis | 2021-10-26 | CAM-NSF-Lichen Genomics Team; Allen, Jessica L. | USA: Pisgah National Forest, Bald Mountains, N-Facing slopes of Bluff Mountain summit, NE of Appalachian Trail 1 mi E of Catpen Gap, 3.4 mi E of Lemon Gap | New York State Museum | 35.84115 N 82.90576 W | NYSM0000194263 |
| PUAP_BFF_07A | Punctelia appalachensis | 2021-10-26 | CAM-NSF-Lichen Genomics Team; Allen, Jessica L. | USA: Pisgah National Forest, Bald Mountains, N-Facing slopes of Bluff Mountain summit, NE of Appalachian Trail 1 mi E of Catpen Gap, 3.4 mi E of Lemon Gap | New York State Museum | 35.84115 N 82.90576 W | NYSM0000194262 |

| | | | | | | | |
|---|---|---|---|---|---|---|---|
| PUAP_BFF_06A | Punctelia appalachensis | 2021-10-26 | CAM-NSF-Lichen Genomics Team; Allen, Jessica L. | USA: Pisgah National Forest, Bald Mountains, N-Facing slopes of Bluff Mountain summit, NE of Appalachian Trail 1 mi E of Catpen Gap, 3.4 mi E of Lemon Gap | New York State Museum | 35.84115 N 82.90576 W | NYSM0000194261 |
| PUAP_BFF_05A | Punctelia appalachensis | 2021-10-26 | CAM-NSF-Lichen Genomics Team; Allen, Jessica L. | USA: Pisgah National Forest, Bald Mountains, N-Facing slopes of Bluff Mountain summit, NE of Appalachian Trail 1 mi E of Catpen Gap, 3.4 mi E of Lemon Gap | New York State Museum | 35.84115 N 82.90576 W | NYSM0000194260 |
| PUAP_BFF_03A | Punctelia appalachensis | 2021-10-26 | CAM-NSF-Lichen Genomics Team; Allen, Jessica L. | USA: Pisgah National Forest, Bald Mountains, N-Facing slopes of Bluff Mountain summit, NE of Appalachian Trail 1 mi E of Catpen Gap, 3.4 mi E of Lemon Gap | New York State Museum | 35.84115 N 82.90576 W | NYSM0000194259 |
| PUAP_BFF_02A | Punctelia appalachensis | 2021-10-26 | CAM-NSF-Lichen Genomics Team; Allen, Jessica L. | USA: Pisgah National Forest, Bald Mountains, N-Facing slopes of Bluff Mountain summit, NE of Appalachian Trail 1 mi E of Catpen Gap, 3.4 mi E of Lemon Gap | New York State Museum | 35.84115 N 82.90576 W | NYSM0000194258 |
| PUAP_BEK_11A | Punctelia appalachensis | 2021-10-26 | CAM-NSF-Lichen Genomics Team; Allen, Jessica L. | USA: Pisgah National Forest, Bald Mountains, N-slope of Roaring Fork Mountain, spur of Buckeye Ridge | New York State Museum | 35.8014 N 82.95096 W | NYSM0000194257 |

| | | | | | | | |
|---|---|---|---|---|---|---|---|
| PUAP_BEK_10A | Punctelia appalachensis | 2021-10-26 | CAM-NSF-Lichen Genomics Team; Allen, Jessica L. | USA: Pisgah National Forest, Bald Mountains, N-slope of Roaring Fork Mountain, spur of Buckeye Ridge | New York State Museum | 35.8014 N 82.95096 W | NYSM0000194256 |
| PUAP_BEK_09A | Punctelia appalachensis | 2021-10-26 | CAM-NSF-Lichen Genomics Team; Allen, Jessica L. | USA: Pisgah National Forest, Bald Mountains, N-slope of Roaring Fork Mountain, spur of Buckeye Ridge | New York State Museum | 35.8014 N 82.95096 W | NYSM0000194255 |
| PUAP_BEK_07A | Punctelia appalachensis | 2021-10-26 | CAM-NSF-Lichen Genomics Team; Allen, Jessica L. | USA: Pisgah National Forest, Bald Mountains, N-slope of Roaring Fork Mountain, spur of Buckeye Ridge | New York State Museum | 35.8014 N 82.95096 W | NYSM0000194254 |
| PUAP_BEK_06A | Punctelia appalachensis | 2021-10-26 | CAM-NSF-Lichen Genomics Team; Allen, Jessica L. | USA: Pisgah National Forest, Bald Mountains, N-slope of Roaring Fork Mountain, spur of Buckeye Ridge | New York State Museum | 35.8014 N 82.95096 W | NYSM0000194253 |
| PUAP_BEK_04A | Punctelia appalachensis | 2021-10-26 | CAM-NSF-Lichen Genomics Team; Allen, Jessica L. | USA: Pisgah National Forest, Bald Mountains, N-slope of Roaring Fork Mountain, spur of Buckeye Ridge | New York State Museum | 35.8014 N 82.95096 W | NYSM0000194252 |
| PUAP_BEK_03A2 | Punctelia appalachensis | 2021-10-26 | CAM-NSF-Lichen Genomics Team; Allen, Jessica L. | USA: Pisgah National Forest, Bald Mountains, N-slope of Roaring Fork Mountain, spur of Buckeye Ridge | New York State Museum | 35.8014 N 82.95096 W | NYSM0000194948 |

| | | | | | | | |
|---|---|---|---|---|---|---|---|
| PUAP_BEK_03A1 | Punctelia appalachensis | 2021-10-26 | CAM-NSF-Lichen Genomics Team; Allen, Jessica L. | USA: Pisgah National Forest, Bald Mountains, N-slope of Roaring Fork Mountain, spur of Buckeye Ridge | New York State Museum | 35.8014 N 82.95096 W | NYSM0000194251 |
| PUAP_BEK_02A | Punctelia appalachensis | 2021-10-26 | CAM-NSF-Lichen Genomics Team; Allen, Jessica L. | USA: Pisgah National Forest, Bald Mountains, N-slope of Roaring Fork Mountain, spur of Buckeye Ridge | New York State Museum | 35.8014 N 82.95096 W | NYSM0000194250 |
| PUAP_BEK_01A1 | Punctelia appalachensis | 2021-10-26 | CAM-NSF-Lichen Genomics Team; Allen, Jessica L. | USA: Pisgah National Forest, Bald Mountains, N-slope of Roaring Fork Mountain, spur of Buckeye Ridge | New York State Museum | 35.8014 N 82.95096 W | NYSM0000194249 |
| USST_WHI_12C | Usnea strigosa | 2022-08-16 | CAM-NSF-Lichen Genomics Team; Denzer, Eli | USA: Nantahala National Forest, summit of Whiterock Mountain, spur trail 0.3 mi N of jct w/ Bartram Trail, 1.3 mi NW of Jones Gap parking area at terminus of FS 4522 | New York State Museum | 35.085 N 83.3053 W | NYSM0000194946 |
| USST_WHI_12B | Usnea strigosa | 2022-08-16 | CAM-NSF-Lichen Genomics Team; Denzer, Eli | USA: Nantahala National Forest, summit of Whiterock Mountain, spur trail 0.3 mi N of jct w/ Bartram Trail, 1.3 mi NW of Jones Gap parking area at terminus of FS 4522 | New York State Museum | 35.085 N 83.3053 W | NYSM0000194945 |

| | | | | | | | |
|---|---|---|---|---|---|---|---|
| USST_WHI_12A | Usnea strigosa | 2022-08-16 | CAM-NSF-Lichen Genomics Team; Denzer, Eli | USA: Nantahala National Forest, summit of Whiterock Mountain, spur trail 0.3 mi N of jct w/ Bartram Trail, 1.3 mi NW of Jones Gap parking area at terminus of FS 4522 | New York State Museum | 35.085 N 83.3053 W | NYSM0000194944 |
| USST_WHI_11C | Usnea strigosa | 2022-08-16 | CAM-NSF-Lichen Genomics Team; Denzer, Eli | USA: Nantahala National Forest, summit of Whiterock Mountain, spur trail 0.3 mi N of jct w/ Bartram Trail, 1.3 mi NW of Jones Gap parking area at terminus of FS 4522 | New York State Museum | 35.085 N 83.3053 W | NYSM0000194943 |
| USST_WHI_07A | Usnea strigosa | 2022-08-04 | CAM-NSF-Lichen Genomics Team; Denzer, Eli | USA: Nantahala National Forest, summit of Whiterock Mountain, spur trail 0.3 mi N of jct w/ Bartram Trail, 1.3 mi NW of Jones Gap parking area at terminus of FS 4522 | New York State Museum | 35.085 N 83.3053 W | NYSM0000194940 |
| USST_WHI_04A | Usnea strigosa | 2022-08-03 | CAM-NSF-Lichen Genomics Team; Denzer, Eli | USA: Nantahala National Forest, summit of Whiterock Mountain, spur trail 0.3 mi N of jct w/ Bartram Trail, 1.3 mi NW of Jones Gap parking area at terminus of FS 4522 | New York State Museum | 35.085 N 83.3053 W | NYSM0000194939 |
| USST_WHI_03A | Usnea strigosa | 2022-08-02 | CAM-NSF-Lichen Genomics Team; Denzer, Eli | USA: Nantahala National Forest, summit of Whiterock Mountain, spur trail 0.3 mi N of jct w/ Bartram Trail, 1.3 mi NW of Jones Gap parking area at terminus of FS 4522 | New York State Museum | 35.085 N 83.3053 W | NYSM0000194938 |

| | | | | | | | |
|---|---|---|---|---|---|---|---|
| USST_WHI_02C | Usnea strigosa | 2022-08-01 | CAM-NSF-Lichen Genomics Team; Denzer, Eli | USA: Nantahala National Forest, summit of Whiterock Mountain, spur trail 0.3 mi N of jct w/ Bartram Trail, 1.3 mi NW of Jones Gap parking area at terminus of FS 4522 | New York State Museum | 35.085 N 83.3053 W | NYSM0000194937 |
| USST_WHI_02B | Usnea strigosa | 2022-07-31 | CAM-NSF-Lichen Genomics Team; Denzer, Eli | USA: Nantahala National Forest, summit of Whiterock Mountain, spur trail 0.3 mi N of jct w/ Bartram Trail, 1.3 mi NW of Jones Gap parking area at terminus of FS 4522 | New York State Museum | 35.085 N 83.3053 W | NYSM0000194936 |
| USST_WHI_02A | Usnea strigosa | 2022-07-30 | CAM-NSF-Lichen Genomics Team; Denzer, Eli | USA: Nantahala National Forest, summit of Whiterock Mountain, spur trail 0.3 mi N of jct w/ Bartram Trail, 1.3 mi NW of Jones Gap parking area at terminus of FS 4522 | New York State Museum | 35.085 N 83.3053 W | NYSM0000194935 |
| USST_SNR_12b | Usnea strigosa | 2022-04-12 | CAM-NSF-Lichen Genomics Team; Allen, Jessica L. | USA: E-slopes of Pine Mountain, Kentenia State Forest, S of Little Shepherd Trail/SR1679 between Mill Cliff Branch and Hi Lewis Branch | New York State Museum | 36.920858 N 83.230543 W | NYSM0000191938 |
| USST_SNR_12a | Usnea strigosa | 2022-04-12 | CAM-NSF-Lichen Genomics Team; Allen, Jessica L. | USA: E-slopes of Pine Mountain, Kentenia State Forest, S of Little Shepherd Trail/SR1679 between Mill Cliff Branch and Hi Lewis Branch | New York State Museum | 36.920858 N 83.230543 W | NYSM0000191937 |

| | | | | | | | |
|---|---|---|---|---|---|---|---|
| USST_SNR_10b | Usnea strigosa | 2022-04-12 | CAM-NSF-Lichen Genomics Team; Allen, Jessica L. | USA: E-slopes of Pine Mountain, Kentenia State Forest, S of Little Shepherd Trail/SR1679 between Mill Cliff Branch and Hi Lewis Branch | New York State Museum | 36.920858 N 83.230543 W | NYSM0000191936 |
| USST_SNR_10a | Usnea strigosa | 2022-04-12 | CAM-NSF-Lichen Genomics Team; Allen, Jessica L. | USA: E-slopes of Pine Mountain, Kentenia State Forest, S of Little Shepherd Trail/SR1679 between Mill Cliff Branch and Hi Lewis Branch | New York State Museum | 36.920858 N 83.230543 W | NYSM0000194934 |
| USST_RON_05c | Usnea strigosa | 2022-04-19 | CAM-NSF-Lichen Genomics Team; Allen, Jessica L. | USA: Jefferson National Forest, Round Mountain, vicinity of summit, Round Mountain Rd./VA688, ~4 mi W of US52/Bastian | New York State Museum | 37.154716 N 81.194163 W | NYSM0000191935 |
| USST_RON_05b | Usnea strigosa | 2022-04-19 | CAM-NSF-Lichen Genomics Team; Allen, Jessica L. | USA: Jefferson National Forest, Round Mountain, vicinity of summit, Round Mountain Rd./VA688, ~4 mi W of US52/Bastian | New York State Museum | 37.154716 N 81.194163 W | NYSM0000191934 |
| USST_RON_05a | Usnea strigosa | 2022-04-19 | CAM-NSF-Lichen Genomics Team; Allen, Jessica L. | USA: Jefferson National Forest, Round Mountain, vicinity of summit, Round Mountain Rd./VA688, ~4 mi W of US52/Bastian | New York State Museum | 37.154716 N 81.194163 W | NYSM0000194926 |
| USST_MAX_10a | Usnea strigosa | 2021-10-26 | CAM-NSF-Lichen Genomics Team; Allen, Jessica L. | USA: Pisgah National Forest, Max Patch Mountain, N-slopes of Max Patch below summit, 0.4 mi E of Max Patch Rd./CR-1182 | New York State Museum | 35.79914 N 82.95679 W | NYSM0000191933 |

| | | | | | | | |
|---|---|---|---|---|---|---|---|
| USST_MAX _09a | Usnea strigosa | 2021-10-26 | CAM-NSF-Lichen Genomics Team; Allen, Jessica L. | USA: Pisgah National Forest, Max Patch Mountain, N-slopes of Max Patch below summit, 0.4 mi E of Max Patch Rd./CR-1182 | New York State Museum | 35.79914 N 82.95679 W | NYSM0000191932 |
| USST_MAX _08a | Usnea strigosa | 2021-10-26 | CAM-NSF-Lichen Genomics Team; Allen, Jessica L. | USA: Pisgah National Forest, Max Patch Mountain, N-slopes of Max Patch below summit, 0.4 mi E of Max Patch Rd./CR-1182 | New York State Museum | 35.79914 N 82.95679 W | NYSM0000191931 |
| USST_MAX _07a | Usnea strigosa | 2021-10-26 | CAM-NSF-Lichen Genomics Team; Allen, Jessica L. | USA: Pisgah National Forest, Max Patch Mountain, N-slopes of Max Patch below summit, 0.4 mi E of Max Patch Rd./CR-1182 | New York State Museum | 35.79914 N 82.95679 W | NYSM0000191930 |
| USST_MAX _06a | Usnea strigosa | 2021-10-26 | CAM-NSF-Lichen Genomics Team; Allen, Jessica L. | USA: Pisgah National Forest, Max Patch Mountain, N-slopes of Max Patch below summit, 0.4 mi E of Max Patch Rd./CR-1182 | New York State Museum | 35.79914 N 82.95679 W | NYSM0000191929 |
| USST_MAX _05a | Usnea strigosa | 2021-10-26 | CAM-NSF-Lichen Genomics Team; Allen, Jessica L. | USA: Pisgah National Forest, Max Patch Mountain, N-slopes of Max Patch below summit, 0.4 mi E of Max Patch Rd./CR-1182 | New York State Museum | 35.79914 N 82.95679 W | NYSM0000191928 |
| USST_MAX _04a | Usnea strigosa | 2021-10-26 | CAM-NSF-Lichen Genomics Team; Allen, Jessica L. | USA: Pisgah National Forest, Max Patch Mountain, N-slopes of Max Patch below summit, 0.4 mi E of Max Patch Rd./CR-1182 | New York State Museum | 35.79914 N 82.95679 W | NYSM0000191927 |

| | | | | | | | |
|---|---|---|---|---|---|---|---|
| USST_MAX _03a | Usnea strigosa | 2021-10-26 | CAM-NSF-Lichen Genomics Team; Allen, Jessica L. | USA: Pisgah National Forest, Max Patch Mountain, N-slopes of Max Patch below summit, 0.4 mi E of Max Patch Rd./CR-1182 | New York State Museum | 35.79914 N 82.95679 W | NYSM0000191926 |
| USST_MAX _02a | Usnea strigosa | 2021-10-26 | CAM-NSF-Lichen Genomics Team; Allen, Jessica L. | USA: Pisgah National Forest, Max Patch Mountain, N-slopes of Max Patch below summit, 0.4 mi E of Max Patch Rd./CR-1182 | New York State Museum | 35.79914 N 82.95679 W | NYSM0000191925 |
| USST_JON_ 06a | Usnea strigosa | 2022-04-19 | CAM-NSF-Lichen Genomics Team; Allen, Jessica L. | USA: Jefferson National Forest, E slopes of Round Mountain, S of Round Mountain Rd./VA688, ~1.5 mi W of US52/Bastian | New York State Museum | 37.170518 N 81.164224 W | NYSM0000191924 |
| USST_JON_ 05b | Usnea strigosa | 2022-04-19 | CAM-NSF-Lichen Genomics Team; Allen, Jessica L. | USA: Jefferson National Forest, E slopes of Round Mountain, S of Round Mountain Rd./VA688, ~1.5 mi W of US52/Bastian | New York State Museum | 37.170518 N 81.164224 W | NYSM0000191923 |
| USST_JON_ 05a | Usnea strigosa | 2022-04-19 | CAM-NSF-Lichen Genomics Team; Allen, Jessica L. | USA: Jefferson National Forest, E slopes of Round Mountain, S of Round Mountain Rd./VA688, ~1.5 mi W of US52/Bastian | New York State Museum | 37.170518 N 81.164224 W | NYSM0000191922 |
| USST_JON_ 03a | Usnea strigosa | 2022-04-19 | CAM-NSF-Lichen Genomics Team; Allen, Jessica L. | USA: Jefferson National Forest, E slopes of Round Mountain, S of Round Mountain Rd./VA688, ~1.5 mi W of US52/Bastian | New York State Museum | 37.170518 N 81.164224 W | NYSM0000191921 |

| | | | | | | | |
|---|---|---|---|---|---|---|---|
| USST_JON_ 02a | Usnea strigosa | 2022-04-19 | CAM-NSF-Lichen Genomics Team; Allen, Jessica L. | USA: Jefferson National Forest, E slopes of Round Mountain, S of Round Mountain Rd./VA688, ~1.5 mi W of US52/Bastian | New York State Museum | 37.170518 N 81.164224 W | NYSM0000191919 |
| USST_JON_ 01d | Usnea strigosa | 2022-04-19 | CAM-NSF-Lichen Genomics Team; Allen, Jessica L. | USA: Jefferson National Forest, E slopes of Round Mountain, S of Round Mountain Rd./VA688, ~1.5 mi W of US52/Bastian | New York State Museum | 37.170518 N 81.164224 W | NYSM0000194921 |
| USST_JON_ 01c | Usnea strigosa | 2022-04-19 | CAM-NSF-Lichen Genomics Team; Allen, Jessica L. | USA: Jefferson National Forest, E slopes of Round Mountain, S of Round Mountain Rd./VA688, ~1.5 mi W of US52/Bastian | New York State Museum | 37.170518 N 81.164224 W | NYSM0000191918 |
| USST_JON_ 01b | Usnea strigosa | 2022-04-19 | CAM-NSF-Lichen Genomics Team; Allen, Jessica L. | USA: Jefferson National Forest, E slopes of Round Mountain, S of Round Mountain Rd./VA688, ~1.5 mi W of US52/Bastian | New York State Museum | 37.170518 N 81.164224 W | NYSM0000191920 |
| USST_JON_ 01a | Usnea strigosa | 2022-04-19 | CAM-NSF-Lichen Genomics Team; Allen, Jessica L. | USA: Jefferson National Forest, Round Mountain, vicinity of summit, Round Mountain Rd./VA688, ~4 mi W of US52/Bastian | New York State Museum | 37.170518 N 81.164224 W | NYSM0000191917 |
| USST_GOS _13c | Usnea strigosa | 2022-04-11 | CAM-NSF-Lichen Genomics Team; Allen, Jessica L. | USA: E-slopes of Pine Mountain, Kentenia State Forest, vicinity of Goss Park, E of Little Shepherd Trail/SR1679 at jct w/ Putney Lookout Tower Rd., between | New York State Museum | 36.9243 N 83.22419 W | NYSM0000191916 |

| | | | | | | | |
|---|---|---|---|---|---|---|---|
| | | | | Spruce Ridge Branch and Hi Lewis Branch | | | |
| USST_GOS _13b | Usnea strigosa | 2022-04-11 | CAM-NSF-Lichen Genomics Team; Allen, Jessica L. | USA: E-slopes of Pine Mountain, Kentenia State Forest, vicinity of Goss Park, E of Little Shepherd Trail/SR1679 at jct w/ Putney Lookout Tower Rd., between Spruce Ridge Branch and Hi Lewis Branch | New York State Museum | 36.9243 N 83.22419 W | NYSM0000191915 |
| USST_GOS _10c | Usnea strigosa | 2022-04-11 | CAM-NSF-Lichen Genomics Team; Allen, Jessica L. | USA: E-slopes of Pine Mountain, Kentenia State Forest, vicinity of Goss Park, E of Little Shepherd Trail/SR1679 at jct w/ Putney Lookout Tower Rd., between Spruce Ridge Branch and Hi Lewis Branch | New York State Museum | 36.9243 N 83.22419 W | NYSM0000191914 |
| USST_GOS _10b | Usnea strigosa | 2022-04-11 | CAM-NSF-Lichen Genomics Team; Allen, Jessica L. | USA: E-slopes of Pine Mountain, Kentenia State Forest, vicinity of Goss Park, E of Little Shepherd Trail/SR1679 at jct w/ Putney Lookout Tower Rd., between Spruce Ridge Branch and Hi Lewis Branch | New York State Museum | 36.9243 N 83.22419 W | NYSM0000191913 |
| USST_GOS _10a | Usnea strigosa | 2022-04-11 | CAM-NSF-Lichen Genomics Team; Allen, Jessica L. | USA: E-slopes of Pine Mountain, Kentenia State Forest, vicinity of Goss Park, E of Little Shepherd Trail/SR1679 at jct w/ Putney Lookout Tower Rd., between | New York State Museum | 36.9243 N 83.22419 W | NYSM0000191912 |

| | | | | | | | |
|---|---|---|---|---|---|---|---|
| | | | | Spruce Ridge Branch and Hi Lewis Branch | | | |
| USST_GOS _09c | Usnea strigosa | 2022-04-11 | CAM-NSF-Lichen Genomics Team; Allen, Jessica L. | USA: E-slopes of Pine Mountain, Kentenia State Forest, vicinity of Goss Park, E of Little Shepherd Trail/SR1679 at jct w/ Putney Lookout Tower Rd., between Spruce Ridge Branch and Hi Lewis Branch | New York State Museum | 36.9243 N 83.22419 W | NYSM0000194920 |
| USST_GOS _09b | Usnea strigosa | 2022-04-11 | CAM-NSF-Lichen Genomics Team; Allen, Jessica L. | USA: E-slopes of Pine Mountain, Kentenia State Forest, vicinity of Goss Park, E of Little Shepherd Trail/SR1679 at jct w/ Putney Lookout Tower Rd., between Spruce Ridge Branch and Hi Lewis Branch | New York State Museum | 36.9243 N 83.22419 W | NYSM0000191911 |
| USST_GOS _09a | Usnea strigosa | 2022-04-11 | CAM-NSF-Lichen Genomics Team; Allen, Jessica L. | USA: E-slopes of Pine Mountain, Kentenia State Forest, vicinity of Goss Park, E of Little Shepherd Trail/SR1679 at jct w/ Putney Lookout Tower Rd., between Spruce Ridge Branch and Hi Lewis Branch | New York State Museum | 36.9243 N 83.22419 W | NYSM0000191910 |
| USST_GOS _04b | Usnea strigosa | 2022-04-11 | CAM-NSF-Lichen Genomics Team; Allen, Jessica L. | USA: E-slopes of Pine Mountain, Kentenia State Forest, vicinity of Goss Park, E of Little Shepherd Trail/SR1679 at jct w/ Putney Lookout Tower Rd., between | New York State Museum | 36.9243 N 83.22419 W | NYSM0000191909 |

| | | | | | | | |
|---|---|---|---|---|---|---|---|
| | | | | Spruce Ridge Branch and Hi Lewis Branch | | | |
| USST_GOS_04a | Usnea strigosa | 2022-04-11 | CAM-NSF-Lichen Genomics Team; Allen, Jessica L. | USA: E-slopes of Pine Mountain, Kentenia State Forest, vicinity of Goss Park, E of Little Shepherd Trail/SR1679 at jct w/ Putney Lookout Tower Rd., between Spruce Ridge Branch and Hi Lewis Branch | New York State Museum | 36.9243 N 83.22419 W | NYSM0000191908 |
| USSU_TRI_05d | Usnea subfusca | 2022-04-28 | CAM-NSF-Lichen Genomics Team; Allen, Jessica L. | USA: Monongahela National Forest, Shavers Mountain, E-facing slopes of Trillium Knob above Old Road Run, S of Glade Run headwaters, Allegheny Trail ~1.5 mi N of FSR27 | New York State Museum | 38.644182 N 79.83029 W | NYSM0000194248 |
| USSU_TRI_05c | Usnea subfusca | 2022-04-28 | CAM-NSF-Lichen Genomics Team; Allen, Jessica L. | USA: Monongahela National Forest, Shavers Mountain, E-facing slopes of Trillium Knob above Old Road Run, S of Glade Run headwaters, Allegheny Trail ~1.5 mi N of FSR27 | New York State Museum | 38.644182 N 79.83029 W | NYSM0000194247 |
| USSU_TRI_05b | Usnea subfusca | 2022-04-28 | CAM-NSF-Lichen Genomics Team; Allen, Jessica L. | USA: Monongahela National Forest, Shavers Mountain, E-facing slopes of Trillium Knob above Old Road Run, S of Glade Run headwaters, Allegheny Trail ~1.5 mi N of FSR27 | New York State Museum | 38.644182 N 79.83029 W | NYSM0000194246 |

| | | | | | | | |
|---|---|---|---|---|---|---|---|
| USSU_TRI_ 05a | Usnea subfusca | 2022-04-28 | CAM-NSF-Lichen Genomics Team; Allen, Jessica L. | USA: Monongahela National Forest, Shavers Mountain, E-facing slopes of Trillium Knob above Old Road Run, S of Glade Run headwaters, Allegheny Trail ~1.5 mi N of FSR27 | New York State Museum | 38.644182 N 79.83029 W | NYSM0000194245 |
| USSU_TRI_ 04a | Usnea subfusca | 2022-04-28 | CAM-NSF-Lichen Genomics Team; Allen, Jessica L. | USA: Monongahela National Forest, Shavers Mountain, E-facing slopes of Trillium Knob above Old Road Run, S of Glade Run headwaters, Allegheny Trail ~1.5 mi N of FSR27 | New York State Museum | 38.644182 N 79.83029 W | NYSM0000194244 |
| USSU_TRI_ 03e | Usnea subfusca | 2022-04-28 | CAM-NSF-Lichen Genomics Team; Allen, Jessica L. | USA: Monongahela National Forest, Shavers Mountain, E-facing slopes of Trillium Knob above Old Road Run, S of Glade Run headwaters, Allegheny Trail ~1.5 mi N of FSR27 | New York State Museum | 38.644182 N 79.83029 W | NYSM0000194243 |
| USSU_TRI_ 03d | Usnea subfusca | 2022-04-28 | CAM-NSF-Lichen Genomics Team; Allen, Jessica L. | USA: Monongahela National Forest, Shavers Mountain, E-facing slopes of Trillium Knob above Old Road Run, S of Glade Run headwaters, Allegheny Trail ~1.5 mi N of FSR27 | New York State Museum | 38.644182 N 79.83029 W | |
| USSU_TRI_ 03c | Usnea subfusca | 2022-04-28 | CAM-NSF-Lichen Genomics Team; Allen, Jessica L. | USA: Monongahela National Forest, Shavers Mountain, E-facing slopes of Trillium Knob above Old Road Run, S of Glade Run headwaters, Allegheny Trail ~1.5 mi N of FSR27 | New York State Museum | 38.644182 N 79.83029 W | NYSM0000191999 |

| | | | | | | | |
|---|---|---|---|---|---|---|---|
| USSU_TRI_03b | Usnea subfusca | 2022-04-28 | CAM-NSF-Lichen Genomics Team; Allen, Jessica L. | USA: Monongahela National Forest, Shavers Mountain, E-facing slopes of Trillium Knob above Old Road Run, S of Glade Run headwaters, Allegheny Trail ~1.5 mi N of FSR27 | New York State Museum | 38.644182 N 79.83029 W | NYSM0000191998 |
| USSU_TRI_03a | Usnea subfusca | 2022-04-28 | CAM-NSF-Lichen Genomics Team; Allen, Jessica L. | USA: Monongahela National Forest, Shavers Mountain, E-facing slopes of Trillium Knob above Old Road Run, S of Glade Run headwaters, Allegheny Trail ~1.5 mi N of FSR27 | New York State Museum | 38.644182 N 79.83029 W | NYSM0000191997 |
| USSU_TRI_02a | Usnea subfusca | 2022-04-28 | CAM-NSF-Lichen Genomics Team; Allen, Jessica L. | USA: Monongahela National Forest, Shavers Mountain, E-facing slopes of Trillium Knob above Old Road Run, S of Glade Run headwaters, Allegheny Trail ~1.5 mi N of FSR27 | New York State Museum | 38.644182 N 79.83029 W | NYSM0000191996 |
| USSU_TRI_01a | Usnea subfusca | 2022-04-28 | CAM-NSF-Lichen Genomics Team; Allen, Jessica L. | USA: Monongahela National Forest, Shavers Mountain, E-facing slopes of Trillium Knob above Old Road Run, S of Glade Run headwaters, Allegheny Trail ~1.5 mi N of FSR27 | New York State Museum | 38.644182 N 79.83029 W | NYSM0000191995 |
| USSU_SHR_09F | Usnea subfusca | 2022-08-04 | CAM-NSF-Lichen Genomics Team; Denzer, Eli | USA: Pisgah National Forest, Balsam Mountains, Shining Rock Wilderness, Shining Rock Ledge, E slopes of Stairs Mountain summit, Art Loeb Trail 1 mi N of Shining Rock Gap | New York State Museum | 35.37817 N 82.86893 W | |

| | | | | | | | |
|---|---|---|---|---|---|---|---|
| USSU_SHR_09E | Usnea subfusca | 2022-08-04 | CAM-NSF-Lichen Genomics Team; Denzer, Eli | USA: Pisgah National Forest, Balsam Mountains, Shining Rock Wilderness, Shining Rock Ledge, E slopes of Stairs Mountain summit, Art Loeb Trail 1 mi N of Shining Rock Gap | New York State Museum | 35.37817 N 82.86893 W | |
| USSU_SHR_09D | Usnea subfusca | 2022-08-04 | CAM-NSF-Lichen Genomics Team; Denzer, Eli | USA: Pisgah National Forest, Balsam Mountains, Shining Rock Wilderness, Shining Rock Ledge, E slopes of Stairs Mountain summit, Art Loeb Trail 1 mi N of Shining Rock Gap | New York State Museum | 35.37817 N 82.86893 W | |
| USSU_SHR_09C | Usnea subfusca | 2022-08-04 | CAM-NSF-Lichen Genomics Team; Denzer, Eli | USA: Pisgah National Forest, Balsam Mountains, Shining Rock Wilderness, Shining Rock Ledge, E slopes of Stairs Mountain summit, Art Loeb Trail 1 mi N of Shining Rock Gap | New York State Museum | 35.37817 N 82.86893 W | NYSM0000191994 |
| USSU_SHR_09B | Usnea subfusca | 2022-08-04 | CAM-NSF-Lichen Genomics Team; Denzer, Eli | USA: Pisgah National Forest, Balsam Mountains, Shining Rock Wilderness, Shining Rock Ledge, E slopes of Stairs Mountain summit, Art Loeb Trail 1 mi N of Shining Rock Gap | New York State Museum | 35.37817 N 82.86893 W | NYSM0000191993 |
| USSU_SHR_09A | Usnea subfusca | 2022-08-04 | CAM-NSF-Lichen Genomics Team; Denzer, Eli | USA: Pisgah National Forest, Balsam Mountains, Shining Rock Wilderness, Shining Rock Ledge, E slopes of Stairs Mountain summit, Art Loeb Trail 1 mi N of Shining Rock Gap | New York State Museum | 35.37817 N 82.86893 W | NYSM0000191992 |

| | | | | | | | |
|---|---|---|---|---|---|---|---|
| USSU_SHR _08A | Usnea subfusca | 2022-08-04 | CAM-NSF-Lichen Genomics Team; Denzer, Eli | USA: Pisgah National Forest, Balsam Mountains, Shining Rock Wilderness, Shining Rock Ledge, E slopes of Stairs Mountain summit, Art Loeb Trail 1 mi N of Shining Rock Gap | New York State Museum | 35.37817 N 82.86893 W | |
| USSU_SHR _06C | Usnea subfusca | 2022-08-04 | CAM-NSF-Lichen Genomics Team; Denzer, Eli | USA: Pisgah National Forest, Balsam Mountains, Shining Rock Wilderness, Shining Rock Ledge, E slopes of Stairs Mountain summit, Art Loeb Trail 1 mi N of Shining Rock Gap | New York State Museum | 35.37817 N 82.86893 W | NYSM0000191991 |
| USSU_SHR _06B | Usnea subfusca | 2022-08-04 | CAM-NSF-Lichen Genomics Team; Denzer, Eli | USA: Pisgah National Forest, Balsam Mountains, Shining Rock Wilderness, Shining Rock Ledge, E slopes of Stairs Mountain summit, Art Loeb Trail 1 mi N of Shining Rock Gap | New York State Museum | 35.37817 N 82.86893 W | NYSM0000191990 |
| USSU_SHR _06A | Usnea subfusca | 2022-08-04 | CAM-NSF-Lichen Genomics Team; Denzer, Eli | USA: Pisgah National Forest, Balsam Mountains, Shining Rock Wilderness, Shining Rock Ledge, E slopes of Stairs Mountain summit, Art Loeb Trail 1 mi N of Shining Rock Gap | New York State Museum | 35.37817 N 82.86893 W | NYSM0000191989 |
| USSU_SHR _05C | Usnea subfusca | 2022-08-04 | CAM-NSF-Lichen Genomics Team; Denzer, Eli | USA: Pisgah National Forest, Balsam Mountains, Shining Rock Wilderness, Shining Rock Ledge, E slopes of Stairs Mountain summit, Art Loeb Trail 1 mi N of Shining Rock Gap | New York State Museum | 35.37817 N 82.86893 W | NYSM0000191988 |

| | | | | | | | |
|---|---|---|---|---|---|---|---|
| USSU_SHR _05B | Usnea subfusca | 2022-08-04 | CAM-NSF-Lichen Genomics Team; Denzer, Eli | USA: Pisgah National Forest, Balsam Mountains, Shining Rock Wilderness, Shining Rock Ledge, E slopes of Stairs Mountain summit, Art Loeb Trail 1 mi N of Shining Rock Gap | New York State Museum | 35.37817 N 82.86893 W | NYSM0000191987 |
| USSU_SHR _05A | Usnea subfusca | 2022-08-04 | CAM-NSF-Lichen Genomics Team; Denzer, Eli | USA: Pisgah National Forest, Balsam Mountains, Shining Rock Wilderness, Shining Rock Ledge, E slopes of Stairs Mountain summit, Art Loeb Trail 1 mi N of Shining Rock Gap | New York State Museum | 35.37817 N 82.86893 W | NYSM0000191986 |
| USSU_SHR _02C | Usnea subfusca | 2022-08-04 | CAM-NSF-Lichen Genomics Team; Denzer, Eli | USA: Pisgah National Forest, Balsam Mountains, Shining Rock Wilderness, Shining Rock Ledge, E slopes of Stairs Mountain summit, Art Loeb Trail 1 mi N of Shining Rock Gap | New York State Museum | 35.37817 N 82.86893 W | NYSM0000191985 |
| USSU_SHR _02B | Usnea subfusca | 2022-08-04 | CAM-NSF-Lichen Genomics Team; Denzer, Eli | USA: Pisgah National Forest, Balsam Mountains, Shining Rock Wilderness, Shining Rock Ledge, E slopes of Stairs Mountain summit, Art Loeb Trail 1 mi N of Shining Rock Gap | New York State Museum | 35.37817 N 82.86893 W | NYSM0000191984 |
| USSU_SHR _02A | Usnea subfusca | 2022-08-04 | CAM-NSF-Lichen Genomics Team; Denzer, Eli | USA: Pisgah National Forest, Balsam Mountains, Shining Rock Wilderness, Shining Rock Ledge, E slopes of Stairs Mountain summit, Art Loeb Trail 1 mi N of Shining Rock Gap | New York State Museum | 35.37817 N 82.86893 W | |

| | | | | | | | |
|---|---|---|---|---|---|---|---|
| USSU_SHR _01C | Usnea subfusca | 2022-08-04 | CAM-NSF-Lichen Genomics Team; Denzer, Eli | USA: Pisgah National Forest, Balsam Mountains, Shining Rock Wilderness, Shining Rock Ledge, E slopes of Stairs Mountain summit, Art Loeb Trail 1 mi N of Shining Rock Gap | New York State Museum | 35.37817 N 82.86893 W | |
| USSU_SHR _01B | Usnea subfusca | 2022-08-04 | CAM-NSF-Lichen Genomics Team; Denzer, Eli | USA: Pisgah National Forest, Balsam Mountains, Shining Rock Wilderness, Shining Rock Ledge, E slopes of Stairs Mountain summit, Art Loeb Trail 1 mi N of Shining Rock Gap | New York State Museum | 35.37817 N 82.86893 W | |
| USSU_SHR _01A | Usnea subfusca | 2022-08-04 | CAM-NSF-Lichen Genomics Team; Denzer, Eli | USA: Pisgah National Forest, Balsam Mountains, Shining Rock Wilderness, Shining Rock Ledge, E slopes of Stairs Mountain summit, Art Loeb Trail 1 mi N of Shining Rock Gap | New York State Museum | 35.37817 N 82.86893 W | NYSM0000191983 |
| USSU_NGA _17b | Usnea subfusca | 2022-04-28 | CAM-NSF-Lichen Genomics Team; Allen, Jessica L. | USA: Monongahela National Forest, Shavers Mountain, E-facing slopes of Gaudineer Knob, FSR27 ~1.25 mi N of jct w/ US250 | New York State Museum | 38.611691 N 79.841629 W | NYSM0000191982 |
| USSU_NGA _17a | Usnea subfusca | 2022-04-28 | CAM-NSF-Lichen Genomics Team; Allen, Jessica L. | USA: Monongahela National Forest, Shavers Mountain, E-facing slopes of Gaudineer Knob, FSR27 ~1.25 mi N of jct w/ US250 | New York State Museum | 38.611691 N 79.841629 W | NYSM0000191981 |

| | | | | | | | |
|---|---|---|---|---|---|---|---|
| USSU_NGA _15a | Usnea subfusca | 2022-04-28 | CAM-NSF-Lichen Genomics Team; Allen, Jessica L. | USA: Monongahela National Forest, Shavers Mountain, E-facing slopes of Gaudineer Knob, FSR27 ~1.25 mi N of jct w/ US250 | New York State Museum | 38.611691 N 79.841629 W | NYSM0000191980 |
| USSU_NGA _14a | Usnea subfusca | 2022-04-28 | CAM-NSF-Lichen Genomics Team; Allen, Jessica L. | USA: Monongahela National Forest, Shavers Mountain, E-facing slopes of Gaudineer Knob, FSR27 ~1.25 mi N of jct w/ US250 | New York State Museum | 38.611691 N 79.841629 W | NYSM0000191977 |
| USSU_NGA _10a | Usnea subfusca | 2022-04-28 | CAM-NSF-Lichen Genomics Team; Allen, Jessica L. | USA: Monongahela National Forest, Shavers Mountain, E-facing slopes of Gaudineer Knob, FSR27 ~1.25 mi N of jct w/ US250 | New York State Museum | 38.611691 N 79.841629 W | NYSM0000191976 |
| USSU_NGA _06b | Usnea subfusca | 2022-04-28 | CAM-NSF-Lichen Genomics Team; Allen, Jessica L. | USA: Monongahela National Forest, Shavers Mountain, E-facing slopes of Gaudineer Knob, FSR27 ~1.25 mi N of jct w/ US250 | New York State Museum | 38.611691 N 79.841629 W | NYSM0000191975 |
| USSU_NGA _06a | Usnea subfusca | 2022-04-28 | CAM-NSF-Lichen Genomics Team; Allen, Jessica L. | USA: Monongahela National Forest, Shavers Mountain, E-facing slopes of Gaudineer Knob, FSR27 ~1.25 mi N of jct w/ US250 | New York State Museum | 38.611691 N 79.841629 W | |
| USSU_NGA _05a | Usnea subfusca | 2022-04-28 | CAM-NSF-Lichen Genomics Team; Allen, Jessica L. | USA: Monongahela National Forest, Shavers Mountain, E-facing slopes of Gaudineer Knob, FSR27 ~1.25 mi N of jct w/ US250 | New York State Museum | 38.611691 N 79.841629 W | |

| | | | | | | | |
|---|---|---|---|---|---|---|---|
| USSU_NAR _11B | Usnea subfusca | 2022-08-04 | CAM-NSF-Lichen Genomics Team; Denzer, Eli | USA: Pisgah National Forest, Balsam Mountains, Shining Rock Wilderness, Shining Rock Ledge, E slopes of Stairs Mountain summit, Art Loeb Trail 1 mi N of Shining Rock Gap | New York State Museum | 35.37817 N 82.86893 W | |
| USSU_NAR _11A | Usnea subfusca | 2022-08-04 | CAM-NSF-Lichen Genomics Team; Denzer, Eli | USA: Pisgah National Forest, Balsam Mountains, Shining Rock Wilderness, Shining Rock Ledge, E slopes of Stairs Mountain summit, Art Loeb Trail 1 mi N of Shining Rock Gap | New York State Museum | 35.37817 N 82.86893 W | |
| USSU_NAR _07C | Usnea subfusca | 2022-08-04 | CAM-NSF-Lichen Genomics Team; Denzer, Eli | USA: Pisgah National Forest, Balsam Mountains, Shining Rock Wilderness, Shining Rock Ledge, E slopes of Stairs Mountain summit, Art Loeb Trail 1 mi N of Shining Rock Gap | New York State Museum | 35.37817 N 82.86893 W | NYSM0000191974 |
| USSU_NAR _07B | Usnea subfusca | 2022-08-04 | CAM-NSF-Lichen Genomics Team; Denzer, Eli | USA: Pisgah National Forest, Balsam Mountains, Shining Rock Wilderness, Shining Rock Ledge, E slopes of Stairs Mountain summit, Art Loeb Trail 1 mi N of Shining Rock Gap | New York State Museum | 35.37817 N 82.86893 W | NYSM0000191973 |
| USSU_NAR _07A | Usnea subfusca | 2022-08-04 | CAM-NSF-Lichen Genomics Team; Denzer, Eli | USA: Pisgah National Forest, Balsam Mountains, Shining Rock Wilderness, Shining Rock Ledge, E slopes of Stairs Mountain summit, Art Loeb Trail 1 mi N of Shining Rock Gap | New York State Museum | 35.37817 N 82.86893 W | NYSM0000191972 |

| | | | | | | | |
|---|---|---|---|---|---|---|---|
| USSU_NAR _06C | Usnea subfusca | 2022-08-04 | CAM-NSF-Lichen Genomics Team; Denzer, Eli | USA: Pisgah National Forest, Balsam Mountains, Shining Rock Wilderness, Shining Rock Ledge, E slopes of Stairs Mountain summit, Art Loeb Trail 1 mi N of Shining Rock Gap | New York State Museum | 35.37817 N 82.86893 W | NYSM0000191971 |
| USSU_NAR _06B | Usnea subfusca | 2022-08-04 | CAM-NSF-Lichen Genomics Team; Denzer, Eli | USA: Pisgah National Forest, Balsam Mountains, Shining Rock Wilderness, Shining Rock Ledge, E slopes of Stairs Mountain summit, Art Loeb Trail 1 mi N of Shining Rock Gap | New York State Museum | 35.37817 N 82.86893 W | NYSM0000191971 |
| USSU_NAR _06A | Usnea subfusca | 2022-08-04 | CAM-NSF-Lichen Genomics Team; Denzer, Eli | USA: Pisgah National Forest, Balsam Mountains, Shining Rock Wilderness, Shining Rock Ledge, E slopes of Stairs Mountain summit, Art Loeb Trail 1 mi N of Shining Rock Gap | New York State Museum | 35.37817 N 82.86893 W | NYSM0000191971 |
| USSU_NAR _03C | Usnea subfusca | 2022-08-04 | CAM-NSF-Lichen Genomics Team; Denzer, Eli | USA: Pisgah National Forest, Balsam Mountains, Shining Rock Wilderness, Shining Rock Ledge, E slopes of Stairs Mountain summit, Art Loeb Trail 1 mi N of Shining Rock Gap | New York State Museum | 35.37817 N 82.86893 W | NYSM0000191971 |
| USSU_NAR _03B | Usnea subfusca | 2022-08-04 | CAM-NSF-Lichen Genomics Team; Denzer, Eli | USA: Pisgah National Forest, Balsam Mountains, Shining Rock Wilderness, Shining Rock Ledge, E slopes of Stairs Mountain summit, Art Loeb Trail 1 mi N of Shining Rock Gap | New York State Museum | 35.37817 N 82.86893 W | NYSM0000191970 |

| | | | | | | | |
|---|---|---|---|---|---|---|---|
| USSU_NAR_03A | Usnea subfusca | 2022-08-04 | CAM-NSF-Lichen Genomics Team; Denzer, Eli | USA: Pisgah National Forest, Balsam Mountains, Shining Rock Wilderness, Shining Rock Ledge, E slopes of Stairs Mountain summit, Art Loeb Trail 1 mi N of Shining Rock Gap | New York State Museum | 35.37817 N 82.86893 W | NYSM0000191969 |
| USSU_NAR_02C | Usnea subfusca | 2022-08-04 | CAM-NSF-Lichen Genomics Team; Denzer, Eli | USA: Pisgah National Forest, Balsam Mountains, Shining Rock Wilderness, Shining Rock Ledge, E slopes of Stairs Mountain summit, Art Loeb Trail 1 mi N of Shining Rock Gap | New York State Museum | 35.37817 N 82.86893 W | NYSM0000191968 |
| USSU_NAR_02B | Usnea subfusca | 2022-08-04 | CAM-NSF-Lichen Genomics Team; Denzer, Eli | USA: Pisgah National Forest, Balsam Mountains, Shining Rock Wilderness, Shining Rock Ledge, E slopes of Stairs Mountain summit, Art Loeb Trail 1 mi N of Shining Rock Gap | New York State Museum | 35.37817 N 82.86893 W | NYSM0000191967 |
| USSU_NAR_02A | Usnea subfusca | 2022-08-04 | CAM-NSF-Lichen Genomics Team; Denzer, Eli | USA: Pisgah National Forest, Balsam Mountains, Shining Rock Wilderness, Shining Rock Ledge, E slopes of Stairs Mountain summit, Art Loeb Trail 1 mi N of Shining Rock Gap | New York State Museum | 35.37817 N 82.86893 W | NYSM0000191966 |
| USSU_NAR_01B | Usnea subfusca | 2022-08-04 | CAM-NSF-Lichen Genomics Team; Denzer, Eli | USA: Pisgah National Forest, Balsam Mountains, Shining Rock Wilderness, Shining Rock Ledge, E slopes of Stairs Mountain summit, Art Loeb Trail 1 mi N of Shining Rock Gap | New York State Museum | 35.37817 N 82.86893 W | |

| | | | | | | | |
|---|---|---|---|---|---|---|---|
| USSU_NAR _01A | Usnea subfusca | 2022-08-04 | CAM-NSF-Lichen Genomics Team; Denzer, Eli | USA: Pisgah National Forest, Balsam Mountains, Shining Rock Wilderness, Shining Rock Ledge, E slopes of Stairs Mountain summit, Art Loeb Trail 1 mi N of Shining Rock Gap | New York State Museum | 35.37817 N 82.86893 W | NYSM0000191965 |
| USSU_COG _14A | Usnea subfusca | 2022-08-01 | CAM-NSF-Lichen Genomics Team; Denzer, Eli | USA: Nantahala National Forest, N-facing slopes above FS4535 0.3 mi W of jct w/ SR1538/Buck Creek Rd. at Cole Gap | New York State Museum | 35.1036 N 83.2075 W | |
| USSU_BUK _36a | Usnea subfusca | 2021-10-24 | CAM-NSF-Lichen Genomics Team; Allen, Jessica L. | USA: Great Smoky Mountains National Park, Cataloochee Divide, N-slope of Little Bald Knob, Hemphill Bald Trail ~3 mi E of Polls Gap | New York State Museum | 35.552276 N 83.120161 W | NYSM0000191964 |
| USSU_BUK _35a | Usnea subfusca | 2021-10-24 | CAM-NSF-Lichen Genomics Team; Allen, Jessica L. | USA: Great Smoky Mountains National Park, Cataloochee Divide, N-slope of Little Bald Knob, Hemphill Bald Trail ~3 mi E of Polls Gap | New York State Museum | 35.552276 N 83.120161 W | NYSM0000191963 |
| USSU_BUK _29a | Usnea subfusca | 2021-10-24 | CAM-NSF-Lichen Genomics Team; Allen, Jessica L. | USA: Great Smoky Mountains National Park, Cataloochee Divide, N-slope of Little Bald Knob, Hemphill Bald Trail ~3 mi E of Polls Gap | New York State Museum | 35.552276 N 83.120161 W | NYSM0000191962 |

| | | | | | | | |
|---|---|---|---|---|---|---|---|
| USSU_BUK_27a | Usnea subfusca | 2021-10-24 | CAM-NSF-Lichen Genomics Team; Allen, Jessica L. | USA: Great Smoky Mountains National Park, Cataloochee Divide, N-slope of Little Bald Knob, Hemphill Bald Trail ~3 mi E of Polls Gap | New York State Museum | 35.552276 N 83.120161 W | NYSM0000191961 |
| USSU_BUK_26a | Usnea subfusca | 2021-10-24 | CAM-NSF-Lichen Genomics Team; Allen, Jessica L. | USA: Great Smoky Mountains National Park, Cataloochee Divide, N-slope of Little Bald Knob, Hemphill Bald Trail ~3 mi E of Polls Gap | New York State Museum | 35.552276 N 83.120161 W | NYSM0000191960 |
| USSU_BUK_23a | Usnea subfusca | 2021-10-24 | CAM-NSF-Lichen Genomics Team; Allen, Jessica L. | USA: Great Smoky Mountains National Park, Cataloochee Divide, N-slope of Little Bald Knob, Hemphill Bald Trail ~3 mi E of Polls Gap | New York State Museum | 35.552276 N 83.120161 W | NYSM0000191959 |
| USSU_BUK_22a | Usnea subfusca | 2021-10-24 | CAM-NSF-Lichen Genomics Team; Allen, Jessica L. | USA: Great Smoky Mountains National Park, Cataloochee Divide, N-slope of Little Bald Knob, Hemphill Bald Trail ~3 mi E of Polls Gap | New York State Museum | 35.552276 N 83.120161 W | NYSM0000191958 |
| USSU_BUK_18a | Usnea subfusca | 2021-10-24 | CAM-NSF-Lichen Genomics Team; Allen, Jessica L. | USA: Great Smoky Mountains National Park, Cataloochee Divide, N-slope of Little Bald Knob, Hemphill Bald Trail ~3 mi E of Polls Gap | New York State Museum | 35.552276 N 83.120161 W | NYSM0000191957 |
| USSU_BUK_06a | Usnea subfusca | 2021-10-24 | CAM-NSF-Lichen Genomics Team; Allen, Jessica L. | USA: Great Smoky Mountains National Park, Cataloochee Divide, N-slope of Little Bald Knob, Hemphill Bald Trail ~3 mi E of Polls Gap | New York State Museum | 35.552276 N 83.120161 W | NYSM0000191956 |

| | | | | | | | |
|---|---|---|---|---|---|---|---|
| USSU_BUK_03a | Usnea subfusca | 2021-10-24 | CAM-NSF-Lichen Genomics Team; Allen, Jessica L. | USA: Great Smoky Mountains National Park, Cataloochee Divide, N-slope of Little Bald Knob, Hemphill Bald Trail ~3 mi E of Polls Gap | New York State Museum | 35.552276 N 83.120161 W | NYSM0000191955 |
| USSU_BFF_29a | Usnea subfusca | 2021-10-26 | CAM-NSF-Lichen Genomics Team; Allen, Jessica L. | USA: Pisgah National Forest, Bald Mountains, N-Facing slopes of Bluff Mountain summit, NE of Appalachian Trail 1 mi E of Catpen Gap, 3.4 mi E of Lemon Gap | New York State Museum | 35.84115 N 82.90576 W | NYSM0000191954 |
| USSU_BFF_27a | Usnea subfusca | 2021-10-26 | CAM-NSF-Lichen Genomics Team; Allen, Jessica L. | USA: Pisgah National Forest, Bald Mountains, N-Facing slopes of Bluff Mountain summit, NE of Appalachian Trail 1 mi E of Catpen Gap, 3.4 mi E of Lemon Gap | New York State Museum | 35.84115 N 82.90576 W | NYSM0000191953 |
| USSU_BFF_26a | Usnea subfusca | 2021-10-26 | CAM-NSF-Lichen Genomics Team; Allen, Jessica L. | USA: Pisgah National Forest, Bald Mountains, N-Facing slopes of Bluff Mountain summit, NE of Appalachian Trail 1 mi E of Catpen Gap, 3.4 mi E of Lemon Gap | New York State Museum | 35.84115 N 82.90576 W | NYSM0000191952 |
| USSU_BFF_25a | Usnea subfusca | 2021-10-26 | CAM-NSF-Lichen Genomics Team; Allen, Jessica L. | USA: Pisgah National Forest, Bald Mountains, N-Facing slopes of Bluff Mountain summit, NE of Appalachian Trail 1 mi E of Catpen Gap, 3.4 mi E of Lemon Gap | New York State Museum | 35.84115 N 82.90576 W | NYSM0000191951 |

| | | | | | | | |
|---|---|---|---|---|---|---|---|
| USSU_BFF_ 24a | Usnea subfusca | 2021-10-26 | CAM-NSF-Lichen Genomics Team; Allen, Jessica L. | USA: Pisgah National Forest, Bald Mountains, N-Facing slopes of Bluff Mountain summit, NE of Appalachian Trail 1 mi E of Catpen Gap, 3.4 mi E of Lemon Gap | New York State Museum | 35.84115 N 82.90576 W | NYSM0000191950 |
| USSU_BFF_ 19a | Usnea subfusca | 2021-10-26 | CAM-NSF-Lichen Genomics Team; Allen, Jessica L. | USA: Pisgah National Forest, Bald Mountains, N-Facing slopes of Bluff Mountain summit, NE of Appalachian Trail 1 mi E of Catpen Gap, 3.4 mi E of Lemon Gap | New York State Museum | 35.84115 N 82.90576 W | NYSM0000191949 |
| USSU_BFF_ 18a | Usnea subfusca | 2021-10-26 | CAM-NSF-Lichen Genomics Team; Allen, Jessica L. | USA: Pisgah National Forest, Bald Mountains, N-Facing slopes of Bluff Mountain summit, NE of Appalachian Trail 1 mi E of Catpen Gap, 3.4 mi E of Lemon Gap | New York State Museum | 35.84115 N 82.90576 W | NYSM0000191948 |
| USSU_BFF_ 15a | Usnea subfusca | 2021-10-26 | CAM-NSF-Lichen Genomics Team; Allen, Jessica L. | USA: Pisgah National Forest, Bald Mountains, N-Facing slopes of Bluff Mountain summit, NE of Appalachian Trail 1 mi E of Catpen Gap, 3.4 mi E of Lemon Gap | New York State Museum | 35.84115 N 82.90576 W | NYSM0000191947 |
| USSU_BFF_ 04a | Usnea subfusca | 2021-10-26 | CAM-NSF-Lichen Genomics Team; Allen, Jessica L. | USA: Pisgah National Forest, Bald Mountains, N-Facing slopes of Bluff Mountain summit, NE of Appalachian Trail 1 mi E of Catpen Gap, 3.4 mi E of Lemon Gap | New York State Museum | 35.84115 N 82.90576 W | NYSM0000191946 |

| | | | | | | | |
|---|---|---|---|---|---|---|---|
| USSU_BEK _29a | Usnea subfusca | 2021-10-26 | CAM-NSF-Lichen Genomics Team; Allen, Jessica L. | USA: Pisgah National Forest, Bald Mountains, N-slope of Roaring Fork Mountain, spur of Buckeye Ridge | New York State Museum | 35.8014 N 82.95096 W | NYSM0000191945 |
| USSU_BEK _28a | Usnea subfusca | 2021-10-26 | CAM-NSF-Lichen Genomics Team; Allen, Jessica L. | USA: Pisgah National Forest, Bald Mountains, N-slope of Roaring Fork Mountain, spur of Buckeye Ridge | New York State Museum | 35.8014 N 82.95096 W | NYSM0000191944 |
| USSU_BEK _26a | Usnea subfusca | 2021-10-26 | CAM-NSF-Lichen Genomics Team; Allen, Jessica L. | USA: Pisgah National Forest, Bald Mountains, N-slope of Roaring Fork Mountain, spur of Buckeye Ridge | New York State Museum | 35.8014 N 82.95096 W | NYSM0000191943 |
| USSU_BEK _22a | Usnea subfusca | 2021-10-26 | CAM-NSF-Lichen Genomics Team; Allen, Jessica L. | USA: Pisgah National Forest, Bald Mountains, N-slope of Roaring Fork Mountain, spur of Buckeye Ridge | New York State Museum | 35.8014 N 82.95096 W | NYSM0000191942 |
| USSU_BEK _19a | Usnea subfusca | 2021-10-26 | CAM-NSF-Lichen Genomics Team; Allen, Jessica L. | USA: Pisgah National Forest, Bald Mountains, N-slope of Roaring Fork Mountain, spur of Buckeye Ridge | New York State Museum | 35.8014 N 82.95096 W | NYSM0000191941 |
| USSU_BEK _18a | Usnea subfusca | 2021-10-26 | CAM-NSF-Lichen Genomics Team; Allen, Jessica L. | USA: Pisgah National Forest, Bald Mountains, N-slope of Roaring Fork Mountain, spur of Buckeye Ridge | New York State Museum | 35.8014 N 82.95096 W | NYSM0000191940 |

| | | | | | | | |
|---|---|---|---|---|---|---|---|
| USSU_BEK _16a | Usnea subfusca | 2021-10-26 | CAM-NSF-Lichen Genomics Team; Allen, Jessica L. | USA: Pisgah National Forest, Bald Mountains, N-slope of Roaring Fork Mountain, spur of Buckeye Ridge | New York State Museum | 35.8014 N 82.95096 W | NYSM0000191939 |
| PURU_WHI _19b | Punctelia rudecta | 2022-08-16 | CAM-NSF-Lichen Genomics Team; Denzer, Eli | USA: Nantahala National Forest, summit of Whiterock Mountain, spur trail 0.3 mi N of jct w/ Bartram Trail, 1.3 mi NW of Jones Gap parking area at terminus of FS 4522 | New York State Museum | 35.085 N 83.3053 W | NYSM0000191674 |
| PURU_WHI _19a | Punctelia rudecta | 2022-08-16 | CAM-NSF-Lichen Genomics Team; Denzer, Eli | USA: Nantahala National Forest, summit of Whiterock Mountain, spur trail 0.3 mi N of jct w/ Bartram Trail, 1.3 mi NW of Jones Gap parking area at terminus of FS 4522 | New York State Museum | 35.085 N 83.3053 W | NYSM0000191673 |
| PURU_WHI _18c | Punctelia rudecta | 2022-08-16 | CAM-NSF-Lichen Genomics Team; Denzer, Eli | USA: Nantahala National Forest, summit of Whiterock Mountain, spur trail 0.3 mi N of jct w/ Bartram Trail, 1.3 mi NW of Jones Gap parking area at terminus of FS 4522 | New York State Museum | 35.085 N 83.3053 W | NYSM0000191672 |
| PURU_WHI _18b | Punctelia rudecta | 2022-08-16 | CAM-NSF-Lichen Genomics Team; Denzer, Eli | USA: Nantahala National Forest, summit of Whiterock Mountain, spur trail 0.3 mi N of jct w/ Bartram Trail, 1.3 mi NW of Jones Gap parking area at terminus of FS 4522 | New York State Museum | 35.085 N 83.3053 W | NYSM0000191671 |

| | | | | | | | |
|---|---|---|---|---|---|---|---|
| PURU_WHI _18a | Punctelia rudecta | 2022-08-16 | CAM-NSF-Lichen Genomics Team; Denzer, Eli | USA: Nantahala National Forest, summit of Whiterock Mountain, spur trail 0.3 mi N of jct w/ Bartram Trail, 1.3 mi NW of Jones Gap parking area at terminus of FS 4522 | New York State Museum | 35.085 N 83.3053 W | NYSM0000191670 |
| PURU_WHI _17a | Punctelia rudecta | 2022-08-16 | CAM-NSF-Lichen Genomics Team; Denzer, Eli | USA: Nantahala National Forest, summit of Whiterock Mountain, spur trail 0.3 mi N of jct w/ Bartram Trail, 1.3 mi NW of Jones Gap parking area at terminus of FS 4522 | New York State Museum | 35.085 N 83.3053 W | NYSM0000191669 |
| PURU_WHI _16c | Punctelia rudecta | 2022-08-16 | CAM-NSF-Lichen Genomics Team; Denzer, Eli | USA: Nantahala National Forest, summit of Whiterock Mountain, spur trail 0.3 mi N of jct w/ Bartram Trail, 1.3 mi NW of Jones Gap parking area at terminus of FS 4522 | New York State Museum | 35.085 N 83.3053 W | NYSM0000191668 |
| PURU_WHI _16a | Punctelia rudecta | 2022-08-16 | CAM-NSF-Lichen Genomics Team; Denzer, Eli | USA: Nantahala National Forest, summit of Whiterock Mountain, spur trail 0.3 mi N of jct w/ Bartram Trail, 1.3 mi NW of Jones Gap parking area at terminus of FS 4522 | New York State Museum | 35.085 N 83.3053 W | NYSM0000191666 |
| PURU_WHI _15c | Punctelia rudecta | 2022-08-16 | CAM-NSF-Lichen Genomics Team; Denzer, Eli | USA: Nantahala National Forest, summit of Whiterock Mountain, spur trail 0.3 mi N of jct w/ Bartram Trail, 1.3 mi NW of Jones Gap parking area at terminus of FS 4522 | New York State Museum | 35.085 N 83.3053 W | NYSM0000191665 |

| | | | | | | | |
|---|---|---|---|---|---|---|---|
| PURU_WHI_15b | Punctelia rudecta | 2022-08-16 | CAM-NSF-Lichen Genomics Team; Denzer, Eli | USA: Nantahala National Forest, summit of Whiterock Mountain, spur trail 0.3 mi N of jct w/ Bartram Trail, 1.3 mi NW of Jones Gap parking area at terminus of FS 4522 | New York State Museum | 35.085 N 83.3053 W | NYSM0000191664 |
| PURU_WHI_14a | Punctelia rudecta | 2022-08-16 | CAM-NSF-Lichen Genomics Team; Denzer, Eli | USA: Nantahala National Forest, summit of Whiterock Mountain, spur trail 0.3 mi N of jct w/ Bartram Trail, 1.3 mi NW of Jones Gap parking area at terminus of FS 4522 | New York State Museum | 35.085 N 83.3053 W | |
| PURU_WHI_13c | Punctelia rudecta | 2022-08-16 | CAM-NSF-Lichen Genomics Team; Denzer, Eli | USA: Nantahala National Forest, summit of Whiterock Mountain, spur trail 0.3 mi N of jct w/ Bartram Trail, 1.3 mi NW of Jones Gap parking area at terminus of FS 4522 | New York State Museum | 35.085 N 83.3053 W | NYSM0000191663 |
| PURU_WHI_13b | Punctelia rudecta | 2022-08-16 | CAM-NSF-Lichen Genomics Team; Denzer, Eli | USA: Nantahala National Forest, summit of Whiterock Mountain, spur trail 0.3 mi N of jct w/ Bartram Trail, 1.3 mi NW of Jones Gap parking area at terminus of FS 4522 | New York State Museum | 35.085 N 83.3053 W | NYSM0000191662 |
| PURU_WHI_13a | Punctelia rudecta | 2022-08-16 | CAM-NSF-Lichen Genomics Team; Denzer, Eli | USA: Nantahala National Forest, summit of Whiterock Mountain, spur trail 0.3 mi N of jct w/ Bartram Trail, 1.3 mi NW of Jones Gap parking area at terminus of FS 4522 | New York State Museum | 35.085 N 83.3053 W | NYSM0000191661 |

| | | | | | | | |
|---|---|---|---|---|---|---|---|
| PURU_WHI _12b | Punctelia rudecta | 2022-08-16 | CAM-NSF-Lichen Genomics Team; Denzer, Eli | USA: Nantahala National Forest, summit of Whiterock Mountain, spur trail 0.3 mi N of jct w/ Bartram Trail, 1.3 mi NW of Jones Gap parking area at terminus of FS 4522 | New York State Museum | 35.085 N 83.3053 W | NYSM0000191660 |
| PURU_WHI _10B | Punctelia rudecta | 2022-08-16 | CAM-NSF-Lichen Genomics Team; Denzer, Eli | USA: Nantahala National Forest, summit of Whiterock Mountain, spur trail 0.3 mi N of jct w/ Bartram Trail, 1.3 mi NW of Jones Gap parking area at terminus of FS 4522 | New York State Museum | 35.085 N 83.3053 W | |
| PURU_WHI _05B | Punctelia rudecta | 2022-07-30 | CAM-NSF-Lichen Genomics Team; Denzer, Eli | USA: Nantahala National Forest, summit of Whiterock Mountain, spur trail 0.3 mi N of jct w/ Bartram Trail, 1.3 mi NW of Jones Gap parking area at terminus of FS 4522 | New York State Museum | 35.085 N 83.3053 W | NYSM0000191659 |
| PURU_WHI _05A | Punctelia rudecta | 2022-07-30 | CAM-NSF-Lichen Genomics Team; Denzer, Eli | USA: Nantahala National Forest, summit of Whiterock Mountain, spur trail 0.3 mi N of jct w/ Bartram Trail, 1.3 mi NW of Jones Gap parking area at terminus of FS 4522 | New York State Museum | 35.085 N 83.3053 W | NYSM0000191658 |
| PURU_WHI _04C | Punctelia rudecta | 2022-07-30 | CAM-NSF-Lichen Genomics Team; Denzer, Eli | USA: Nantahala National Forest, summit of Whiterock Mountain, spur trail 0.3 mi N of jct w/ Bartram Trail, 1.3 mi NW of Jones Gap parking area at terminus of FS 4522 | New York State Museum | 35.085 N 83.3053 W | NYSM0000191657 |

| | | | | | | | |
|---|---|---|---|---|---|---|---|
| PURU_WHI _04B | Punctelia rudecta | 2022-07-30 | CAM-NSF-Lichen Genomics Team; Denzer, Eli | USA: Nantahala National Forest, summit of Whiterock Mountain, spur trail 0.3 mi N of jct w/ Bartram Trail, 1.3 mi NW of Jones Gap parking area at terminus of FS 4522 | New York State Museum | 35.085 N 83.3053 W | NYSM0000191656 |
| PURU_WHI _04A | Punctelia rudecta | 2022-07-30 | CAM-NSF-Lichen Genomics Team; Denzer, Eli | USA: Nantahala National Forest, summit of Whiterock Mountain, spur trail 0.3 mi N of jct w/ Bartram Trail, 1.3 mi NW of Jones Gap parking area at terminus of FS 4522 | New York State Museum | 35.085 N 83.3053 W | NYSM0000191655 |
| PURU_WHI _03C | Punctelia rudecta | 2022-07-30 | CAM-NSF-Lichen Genomics Team; Denzer, Eli | USA: Nantahala National Forest, summit of Whiterock Mountain, spur trail 0.3 mi N of jct w/ Bartram Trail, 1.3 mi NW of Jones Gap parking area at terminus of FS 4522 | New York State Museum | 35.085 N 83.3053 W | NYSM0000191654 |
| PURU_WHI _03B | Punctelia rudecta | 2022-07-30 | CAM-NSF-Lichen Genomics Team; Denzer, Eli | USA: Nantahala National Forest, summit of Whiterock Mountain, spur trail 0.3 mi N of jct w/ Bartram Trail, 1.3 mi NW of Jones Gap parking area at terminus of FS 4522 | New York State Museum | 35.085 N 83.3053 W | NYSM0000191653 |
| PURU_WHI _03A | Punctelia rudecta | 2022-07-30 | CAM-NSF-Lichen Genomics Team; Denzer, Eli | USA: Nantahala National Forest, summit of Whiterock Mountain, spur trail 0.3 mi N of jct w/ Bartram Trail, 1.3 mi NW of Jones Gap parking area at terminus of FS 4522 | New York State Museum | 35.085 N 83.3053 W | NYSM0000191652 |

| | | | | | | | |
|---|---|---|---|---|---|---|---|
| PURU_TYS_04c | Punctelia rudecta | 2022-05-03 | CAM-NSF-Lichen Genomics Team; Allen, Jessica L. | USA: Shenandoah National Park, Central District, Bettys Rock, ~0.3 mi N of Crescent Rock, upper W-facing slopes of Timber Hollow | New York State Museum | 38.56753 N 78.38223 W | NYSM0000191651 |
| PURU_TYS_04b | Punctelia rudecta | 2022-05-03 | CAM-NSF-Lichen Genomics Team; Allen, Jessica L. | USA: Shenandoah National Park, Central District, Bettys Rock, ~0.3 mi N of Crescent Rock, upper W-facing slopes of Timber Hollow | New York State Museum | 38.56753 N 78.38223 W | NYSM0000191650 |
| PURU_TYS_04a | Punctelia rudecta | 2022-05-03 | CAM-NSF-Lichen Genomics Team; Allen, Jessica L. | USA: Shenandoah National Park, Central District, Bettys Rock, ~0.3 mi N of Crescent Rock, upper W-facing slopes of Timber Hollow | New York State Museum | 38.56753 N 78.38223 W | NYSM0000191649 |
| PURU_TYS_03c | Punctelia rudecta | 2022-05-03 | CAM-NSF-Lichen Genomics Team; Allen, Jessica L. | USA: Shenandoah National Park, Central District, Bettys Rock, ~0.3 mi N of Crescent Rock, upper W-facing slopes of Timber Hollow | New York State Museum | 38.56753 N 78.38223 W | NYSM0000191648 |
| PURU_TYS_03b | Punctelia rudecta | 2022-05-03 | CAM-NSF-Lichen Genomics Team; Allen, Jessica L. | USA: Shenandoah National Park, Central District, Bettys Rock, ~0.3 mi N of Crescent Rock, upper W-facing slopes of Timber Hollow | New York State Museum | 38.56753 N 78.38223 W | NYSM0000191647 |
| PURU_TYS_03a | Punctelia rudecta | 2022-05-03 | CAM-NSF-Lichen Genomics Team; Allen, Jessica L. | USA: Shenandoah National Park, Central District, Bettys Rock, ~0.3 mi N of Crescent Rock, upper W-facing slopes of Timber Hollow | New York State Museum | 38.56753 N 78.38223 W | NYSM0000191646 |

| | | | | | | | |
|---|---|---|---|---|---|---|---|
| PURU_TYS_01c | Punctelia rudecta | 2022-05-03 | CAM-NSF-Lichen Genomics Team; Allen, Jessica L. | USA: Shenandoah National Park, Central District, Bettys Rock, ~0.3 mi N of Crescent Rock, upper W-facing slopes of Timber Hollow | New York State Museum | 38.56753 N 78.38223 W | NYSM0000191645 |
| PURU_TYS_01b | Punctelia rudecta | 2022-05-03 | CAM-NSF-Lichen Genomics Team; Allen, Jessica L. | USA: Shenandoah National Park, Central District, Bettys Rock, ~0.3 mi N of Crescent Rock, upper W-facing slopes of Timber Hollow | New York State Museum | 38.56753 N 78.38223 W | NYSM0000191644 |
| PURU_TYS_01a | Punctelia rudecta | 2022-05-03 | CAM-NSF-Lichen Genomics Team; Allen, Jessica L. | USA: Shenandoah National Park, Central District, Bettys Rock, ~0.3 mi N of Crescent Rock, upper W-facing slopes of Timber Hollow | New York State Museum | 38.56753 N 78.38223 W | NYSM0000191643 |
| PURU_TRI_13c | Punctelia rudecta | 2022-04-28 | CAM-NSF-Lichen Genomics Team; Allen, Jessica L. | USA: Monongahela National Forest, Shavers Mountain, E-facing slopes of Trillium Knob above Old Road Run, S of Glade Run headwaters, Allegheny Trail ~1.5 mi N of FSR27 | New York State Museum | 38.644182 N 79.83029 W | NYSM0000191642 |
| PURU_TRI_13b | Punctelia rudecta | 2022-04-28 | CAM-NSF-Lichen Genomics Team; Allen, Jessica L. | USA: Monongahela National Forest, Shavers Mountain, E-facing slopes of Trillium Knob above Old Road Run, S of Glade Run headwaters, Allegheny Trail ~1.5 mi N of FSR27 | New York State Museum | 38.644182 N 79.83029 W | NYSM0000191641 |

| | | | | | | | |
|---|---|---|---|---|---|---|---|
| PURU_TRI_13a | Punctelia rudecta | 2022-04-28 | CAM-NSF-Lichen Genomics Team; Allen, Jessica L. | USA: Monongahela National Forest, Shavers Mountain, E-facing slopes of Trillium Knob above Old Road Run, S of Glade Run headwaters, Allegheny Trail ~1.5 mi N of FSR27 | New York State Museum | 38.644182 N 79.83029 W | NYSM0000191640 |
| PURU_TRI_09b | Punctelia rudecta | 2022-04-28 | CAM-NSF-Lichen Genomics Team; Allen, Jessica L. | USA: Monongahela National Forest, Shavers Mountain, E-facing slopes of Trillium Knob above Old Road Run, S of Glade Run headwaters, Allegheny Trail ~1.5 mi N of FSR27 | New York State Museum | 38.644182 N 79.83029 W | NYSM0000191639 |
| PURU_TRI_09a | Punctelia rudecta | 2022-04-28 | CAM-NSF-Lichen Genomics Team; Allen, Jessica L. | USA: Monongahela National Forest, Shavers Mountain, E-facing slopes of Trillium Knob above Old Road Run, S of Glade Run headwaters, Allegheny Trail ~1.5 mi N of FSR27 | New York State Museum | 38.644182 N 79.83029 W | NYSM0000191638 |
| PURU_TRI_08b | Punctelia rudecta | 2022-04-28 | CAM-NSF-Lichen Genomics Team; Allen, Jessica L. | USA: Monongahela National Forest, Shavers Mountain, E-facing slopes of Trillium Knob above Old Road Run, S of Glade Run headwaters, Allegheny Trail ~1.5 mi N of FSR27 | New York State Museum | 38.644182 N 79.83029 W | NYSM0000191637 |
| PURU_TRI_08a | Punctelia rudecta | 2022-04-28 | CAM-NSF-Lichen Genomics Team; Allen, Jessica L. | USA: Monongahela National Forest, Shavers Mountain, E-facing slopes of Trillium Knob above Old Road Run, S of Glade Run headwaters, Allegheny Trail ~1.5 mi N of FSR27 | New York State Museum | 38.644182 N 79.83029 W | NYSM0000191636 |

| | | | | | | | |
|---|---|---|---|---|---|---|---|
| PURU_TRI_07c | Punctelia rudecta | 2022-04-28 | CAM-NSF-Lichen Genomics Team; Allen, Jessica L. | USA: Monongahela National Forest, Shavers Mountain, E-facing slopes of Trillium Knob above Old Road Run, S of Glade Run headwaters, Allegheny Trail ~1.5 mi N of FSR27 | New York State Museum | 38.644182 N 79.83029 W | NYSM0000191635 |
| PURU_TRI_07b | Punctelia rudecta | 2022-04-28 | CAM-NSF-Lichen Genomics Team; Allen, Jessica L. | USA: Monongahela National Forest, Shavers Mountain, E-facing slopes of Trillium Knob above Old Road Run, S of Glade Run headwaters, Allegheny Trail ~1.5 mi N of FSR27 | New York State Museum | 38.644182 N 79.83029 W | NYSM0000191634 |
| PURU_TRI_07a | Punctelia rudecta | 2022-04-28 | CAM-NSF-Lichen Genomics Team; Allen, Jessica L. | USA: Monongahela National Forest, Shavers Mountain, E-facing slopes of Trillium Knob above Old Road Run, S of Glade Run headwaters, Allegheny Trail ~1.5 mi N of FSR27 | New York State Museum | 38.644182 N 79.83029 W | NYSM0000191633 |
| PURU_TOM_12a | Punctelia rudecta | 2021-10-27 | CAM-NSF-Lichen Genomics Team; Allen, Jessica L. | USA: Mount Mitchell State Park, vicinity of summit of Balsam Cone, Black Mountain Crest Trail ~2 mi N of Mount Mitchell and terminus of NC128 | New York State Museum | 35.7892 N 82.2558 W | NYSM0000191622 |
| PURU_TOM_11a | Punctelia rudecta | 2021-10-27 | CAM-NSF-Lichen Genomics Team; Allen, Jessica L. | USA: Mount Mitchell State Park, vicinity of summit of Balsam Cone, Black Mountain Crest Trail ~2 mi N of Mount Mitchell and terminus of NC128 | New York State Museum | 35.7892 N 82.2558 W | NYSM0000191621 |

| | | | | | | | |
|---|---|---|---|---|---|---|---|
| PURU_TOM _10a | Punctelia rudecta | 2021-10-27 | CAM-NSF-Lichen Genomics Team; Allen, Jessica L. | USA: Mount Mitchell State Park, vicinity of summit of Balsam Cone, Black Mountain Crest Trail ~2 mi N of Mount Mitchell and terminus of NC128 | New York State Museum | 35.7892 N 82.2558 W | NYSM0000191620 |
| PURU_TOM _08a | Punctelia rudecta | 2021-10-27 | CAM-NSF-Lichen Genomics Team; Allen, Jessica L. | USA: Mount Mitchell State Park, vicinity of summit of Balsam Cone, Black Mountain Crest Trail ~2 mi N of Mount Mitchell and terminus of NC128 | New York State Museum | 35.7892 N 82.2558 W | NYSM0000191619 |
| PURU_TOM _07a | Punctelia rudecta | 2021-10-27 | CAM-NSF-Lichen Genomics Team; Allen, Jessica L. | USA: Mount Mitchell State Park, vicinity of summit of Balsam Cone, Black Mountain Crest Trail ~2 mi N of Mount Mitchell and terminus of NC128 | New York State Museum | 35.7892 N 82.2558 W | NYSM0000191618 |
| PURU_TOM _06a | Punctelia rudecta | 2021-10-27 | CAM-NSF-Lichen Genomics Team; Allen, Jessica L. | USA: Mount Mitchell State Park, vicinity of summit of Balsam Cone, Black Mountain Crest Trail ~2 mi N of Mount Mitchell and terminus of NC128 | New York State Museum | 35.7892 N 82.2558 W | NYSM0000191617 |
| PURU_TOM _05a | Punctelia rudecta | 2021-10-27 | CAM-NSF-Lichen Genomics Team; Allen, Jessica L. | USA: Mount Mitchell State Park, vicinity of summit of Balsam Cone, Black Mountain Crest Trail ~2 mi N of Mount Mitchell and terminus of NC128 | New York State Museum | 35.7892 N 82.2558 W | NYSM0000191616 |
| PURU_TOM _04a | Punctelia rudecta | 2021-10-27 | CAM-NSF-Lichen Genomics Team; Allen, Jessica L. | USA: Mount Mitchell State Park, vicinity of summit of Balsam Cone, Black Mountain Crest Trail ~2 mi N of Mount Mitchell and terminus of NC128 | New York State Museum | 35.7892 N 82.2558 W | NYSM0000191615 |

| | | | | | | | |
|---|---|---|---|---|---|---|---|
| PURU_TOM _03a | Punctelia rudecta | 2021-10-27 | CAM-NSF-Lichen Genomics Team; Allen, Jessica L. | USA: Mount Mitchell State Park, vicinity of summit of Balsam Cone, Black Mountain Crest Trail ~2 mi N of Mount Mitchell and terminus of NC128 | New York State Museum | 35.7892 N 82.2558 W | NYSM0000191614 |
| PURU_TOM _01a | Punctelia rudecta | 2021-10-27 | CAM-NSF-Lichen Genomics Team; Allen, Jessica L. | USA: Mount Mitchell State Park, vicinity of summit of Balsam Cone, Black Mountain Crest Trail ~2 mi N of Mount Mitchell and terminus of NC128 | New York State Museum | 35.7892 N 82.2558 W | NYSM0000191613 |
| PURU_TMK _05b | Punctelia rudecta | 2022-04-16 | CAM-NSF-Lichen Genomics Team; Allen, Jessica L. | USA: Jefferson National Forest, Mount Rogers National Recreation Area, Lewis Fork Wilderness, E-facing slopes of Pine Mountain, NW of Appalachian Trail ~0.6 E of Thomas Knob Shelter | New York State Museum | 36.655015 N 81.526364 W | NYSM0000191632 |
| PURU_TMK _05a | Punctelia rudecta | 2022-04-16 | CAM-NSF-Lichen Genomics Team; Allen, Jessica L. | USA: Jefferson National Forest, Mount Rogers National Recreation Area, Lewis Fork Wilderness, E-facing slopes of Pine Mountain, NW of Appalachian Trail ~0.6 E of Thomas Knob Shelter | New York State Museum | 36.655015 N 81.526364 W | NYSM0000191631 |
| PURU_TMK _04a | Punctelia rudecta | 2022-04-16 | CAM-NSF-Lichen Genomics Team; Allen, Jessica L. | USA: Jefferson National Forest, Mount Rogers National Recreation Area, Lewis Fork Wilderness, E-facing slopes of Pine Mountain, NW of Appalachian Trail ~0.6 E of Thomas Knob Shelter | New York State Museum | 36.655015 N 81.526364 W | NYSM0000191630 |

| | | | | | | | |
|---|---|---|---|---|---|---|---|
| PURU_TMK _03b | Punctelia rudecta | 2022-04-16 | CAM-NSF-Lichen Genomics Team; Allen, Jessica L. | USA: Jefferson National Forest, Mount Rogers National Recreation Area, Lewis Fork Wilderness, E-facing slopes of Pine Mountain, NW of Appalachian Trail ~0.6 E of Thomas Knob Shelter | New York State Museum | 36.655015 N 81.526364 W | NYSM0000191629 |
| PURU_TMK _03a | Punctelia rudecta | 2022-04-16 | CAM-NSF-Lichen Genomics Team; Allen, Jessica L. | USA: Jefferson National Forest, Mount Rogers National Recreation Area, Lewis Fork Wilderness, E-facing slopes of Pine Mountain, NW of Appalachian Trail ~0.6 E of Thomas Knob Shelter | New York State Museum | 36.655015 N 81.526364 W | NYSM0000191628 |
| PURU_TMK _02c | Punctelia rudecta | 2022-04-16 | CAM-NSF-Lichen Genomics Team; Allen, Jessica L. | USA: Jefferson National Forest, Mount Rogers National Recreation Area, Lewis Fork Wilderness, E-facing slopes of Pine Mountain, NW of Appalachian Trail ~0.6 E of Thomas Knob Shelter | New York State Museum | 36.655015 N 81.526364 W | NYSM0000191627 |
| PURU_TMK _02b | Punctelia rudecta | 2022-04-16 | CAM-NSF-Lichen Genomics Team; Allen, Jessica L. | USA: Jefferson National Forest, Mount Rogers National Recreation Area, Lewis Fork Wilderness, E-facing slopes of Pine Mountain, NW of Appalachian Trail ~0.6 E of Thomas Knob Shelter | New York State Museum | 36.655015 N 81.526364 W | NYSM0000191626 |

| | | | | | | | |
|---|---|---|---|---|---|---|---|
| PURU_TMK_02a | Punctelia rudecta | 2022-04-16 | CAM-NSF-Lichen Genomics Team; Allen, Jessica L. | USA: Jefferson National Forest, Mount Rogers National Recreation Area, Lewis Fork Wilderness, E-facing slopes of Pine Mountain, NW of Appalachian Trail ~0.6 E of Thomas Knob Shelter | New York State Museum | 36.655015 N 81.526364 W | NYSM0000191625 |
| PURU_TMK_01b | Punctelia rudecta | 2022-04-16 | CAM-NSF-Lichen Genomics Team; Allen, Jessica L. | USA: Jefferson National Forest, Mount Rogers National Recreation Area, Lewis Fork Wilderness, E-facing slopes of Pine Mountain, NW of Appalachian Trail ~0.6 E of Thomas Knob Shelter | New York State Museum | 36.655015 N 81.526364 W | NYSM0000191624 |
| PURU_TMK_01a | Punctelia rudecta | 2022-04-16 | CAM-NSF-Lichen Genomics Team; Allen, Jessica L. | USA: Jefferson National Forest, Mount Rogers National Recreation Area, Lewis Fork Wilderness, E-facing slopes of Pine Mountain, NW of Appalachian Trail ~0.6 E of Thomas Knob Shelter | New York State Museum | 36.655015 N 81.526364 W | NYSM0000191623 |
| PURU_SNR_07c | Punctelia rudecta | 2022-04-12 | CAM-NSF-Lichen Genomics Team; Allen, Jessica L. | USA: E-slopes of Pine Mountain, Kentenia State Forest, S of Little Shepherd Trail/SR1679 between Mill Cliff Branch and Hi Lewis Branch | New York State Museum | 36.920858 N 83.230543 W | NYSM0000191612 |
| PURU_SNR_07b | Punctelia rudecta | 2022-04-12 | CAM-NSF-Lichen Genomics Team; Allen, Jessica L. | USA: E-slopes of Pine Mountain, Kentenia State Forest, S of Little Shepherd Trail/SR1679 between Mill Cliff Branch and Hi Lewis Branch | New York State Museum | 36.920858 N 83.230543 W | NYSM0000191611 |

| | | | | | | | |
|---|---|---|---|---|---|---|---|
| PURU_SNR_07a | Punctelia rudecta | 2022-04-12 | CAM-NSF-Lichen Genomics Team; Allen, Jessica L. | USA: E-slopes of Pine Mountain, Kentenia State Forest, S of Little Shepherd Trail/SR1679 between Mill Cliff Branch and Hi Lewis Branch | New York State Museum | 36.920858 N 83.230543 W | NYSM0000191610 |
| PURU_SNR_05c | Punctelia rudecta | 2022-04-12 | CAM-NSF-Lichen Genomics Team; Allen, Jessica L. | USA: E-slopes of Pine Mountain, Kentenia State Forest, S of Little Shepherd Trail/SR1679 between Mill Cliff Branch and Hi Lewis Branch | New York State Museum | 36.920858 N 83.230543 W | NYSM0000191609 |
| PURU_SNR_05b | Punctelia rudecta | 2022-04-12 | CAM-NSF-Lichen Genomics Team; Allen, Jessica L. | USA: E-slopes of Pine Mountain, Kentenia State Forest, S of Little Shepherd Trail/SR1679 between Mill Cliff Branch and Hi Lewis Branch | New York State Museum | 36.920858 N 83.230543 W | NYSM0000191608 |
| PURU_SNR_05a | Punctelia rudecta | 2022-04-12 | CAM-NSF-Lichen Genomics Team; Allen, Jessica L. | USA: E-slopes of Pine Mountain, Kentenia State Forest, S of Little Shepherd Trail/SR1679 between Mill Cliff Branch and Hi Lewis Branch | New York State Museum | 36.920858 N 83.230543 W | NYSM0000191607 |
| PURU_SNR_04b | Punctelia rudecta | 2022-04-12 | CAM-NSF-Lichen Genomics Team; Allen, Jessica L. | USA: E-slopes of Pine Mountain, Kentenia State Forest, S of Little Shepherd Trail/SR1679 between Mill Cliff Branch and Hi Lewis Branch | New York State Museum | 36.920858 N 83.230543 W | NYSM0000191606 |
| PURU_SNR_04a | Punctelia rudecta | 2022-04-12 | CAM-NSF-Lichen Genomics Team; Allen, Jessica L. | USA: E-slopes of Pine Mountain, Kentenia State Forest, S of Little Shepherd Trail/SR1679 between Mill Cliff Branch and Hi Lewis Branch | New York State Museum | 36.920858 N 83.230543 W | NYSM0000191605 |

| | | | | | | | |
|---|---|---|---|---|---|---|---|
| PURU_SNR_03a | Punctelia rudecta | 2022-04-12 | CAM-NSF-Lichen Genomics Team; Allen, Jessica L. | USA: E-slopes of Pine Mountain, Kentenia State Forest, S of Little Shepherd Trail/SR1679 between Mill Cliff Branch and Hi Lewis Branch | New York State Museum | 36.920858 N 83.230543 W | NYSM0000191604 |
| PURU_SNR_02b | Punctelia rudecta | 2022-04-12 | CAM-NSF-Lichen Genomics Team; Allen, Jessica L. | USA: E-slopes of Pine Mountain, Kentenia State Forest, S of Little Shepherd Trail/SR1679 between Mill Cliff Branch and Hi Lewis Branch | New York State Museum | 36.920858 N 83.230543 W | NYSM0000191603 |
| PURU_SNR_02a | Punctelia rudecta | 2022-04-12 | CAM-NSF-Lichen Genomics Team; Allen, Jessica L. | USA: E-slopes of Pine Mountain, Kentenia State Forest, S of Little Shepherd Trail/SR1679 between Mill Cliff Branch and Hi Lewis Branch | New York State Museum | 36.920858 N 83.230543 W | NYSM0000191602 |
| PURU_SHR_UNK | Punctelia rudecta | 2022-08-04 | CAM-NSF-Lichen Genomics Team; Denzer, Eli | USA: Pisgah National Forest, Balsam Mountains, Shining Rock Wilderness, Shining Rock Ledge, E slopes of Stairs Mountain summit, Art Loeb Trail 1 mi N of Shining Rock Gap | New York State Museum | 35.37817 N 82.86893 W | NYSM0000191601 |
| PURU_SHR_07a | Punctelia rudecta | 2022-08-04 | CAM-NSF-Lichen Genomics Team; Denzer, Eli | USA: Pisgah National Forest, Balsam Mountains, Shining Rock Wilderness, Shining Rock Ledge, E slopes of Stairs Mountain summit, Art Loeb Trail 1 mi N of Shining Rock Gap | New York State Museum | 35.37817 N 82.86893 W | NYSM0000191592 |

| | | | | | | | |
|---|---|---|---|---|---|---|---|
| PURU_SHR _06a | Punctelia rudecta | 2022-08-04 | CAM-NSF-Lichen Genomics Team; Denzer, Eli | USA: Pisgah National Forest, Balsam Mountains, Shining Rock Wilderness, Shining Rock Ledge, E slopes of Stairs Mountain summit, Art Loeb Trail 1 mi N of Shining Rock Gap | New York State Museum | 35.37817 N 82.86893 W | NYSM0000191591 |
| PURU_SHR _03c | Punctelia rudecta | 2022-08-04 | CAM-NSF-Lichen Genomics Team; Denzer, Eli | USA: Pisgah National Forest, Balsam Mountains, Shining Rock Wilderness, Shining Rock Ledge, E slopes of Stairs Mountain summit, Art Loeb Trail 1 mi N of Shining Rock Gap | New York State Museum | 35.37817 N 82.86893 W | NYSM0000191590 |
| PURU_SHR _03b | Punctelia rudecta | 2022-08-04 | CAM-NSF-Lichen Genomics Team; Denzer, Eli | USA: Pisgah National Forest, Balsam Mountains, Shining Rock Wilderness, Shining Rock Ledge, E slopes of Stairs Mountain summit, Art Loeb Trail 1 mi N of Shining Rock Gap | New York State Museum | 35.37817 N 82.86893 W | NYSM0000191589 |
| PURU_SHR _03a | Punctelia rudecta | 2022-08-04 | CAM-NSF-Lichen Genomics Team; Denzer, Eli | USA: Pisgah National Forest, Balsam Mountains, Shining Rock Wilderness, Shining Rock Ledge, E slopes of Stairs Mountain summit, Art Loeb Trail 1 mi N of Shining Rock Gap | New York State Museum | 35.37817 N 82.86893 W | NYSM0000191588 |
| PURU_RSK _04c | Punctelia rudecta | 2022-04-26 | CAM-NSF-Lichen Genomics Team; Allen, Jessica L. | USA: Monongahela National Forest, S-facing slopes of Red Spruce Knob, N of Red Lick Mountain, 0.1 mi W of WV150 and FSR451 | New York State Museum | 38.323665 N 80.152203 W | NYSM0000191587 |

| | | | | | | | |
|---|---|---|---|---|---|---|---|
| PURU_RSK _04b | Punctelia rudecta | 2022-04-26 | CAM-NSF-Lichen Genomics Team; Allen, Jessica L. | USA: Monongahela National Forest, S-facing slopes of Red Spruce Knob, N of Red Lick Mountain, 0.1 mi W of WV150 and FSR451 | New York State Museum | 38.323665 N 80.152203 W | NYSM0000191586 |
| PURU_RSK _04a | Punctelia rudecta | 2022-04-26 | CAM-NSF-Lichen Genomics Team; Allen, Jessica L. | USA: Monongahela National Forest, S-facing slopes of Red Spruce Knob, N of Red Lick Mountain, 0.1 mi W of WV150 and FSR451 | New York State Museum | 38.323665 N 80.152203 W | NYSM0000191585 |
| PURU_RSK _03c | Punctelia rudecta | 2022-04-26 | CAM-NSF-Lichen Genomics Team; Allen, Jessica L. | USA: Monongahela National Forest, S-facing slopes of Red Spruce Knob, N of Red Lick Mountain, 0.1 mi W of WV150 and FSR451 | New York State Museum | 38.323665 N 80.152203 W | NYSM0000191584 |
| PURU_RSK _03b | Punctelia rudecta | 2022-04-26 | CAM-NSF-Lichen Genomics Team; Allen, Jessica L. | USA: Monongahela National Forest, S-facing slopes of Red Spruce Knob, N of Red Lick Mountain, 0.1 mi W of WV150 and FSR451 | New York State Museum | 38.323665 N 80.152203 W | NYSM0000191583 |
| PURU_RSK _03a | Punctelia rudecta | 2022-04-26 | CAM-NSF-Lichen Genomics Team; Allen, Jessica L. | USA: Monongahela National Forest, S-facing slopes of Red Spruce Knob, N of Red Lick Mountain, 0.1 mi W of WV150 and FSR451 | New York State Museum | 38.323665 N 80.152203 W | NYSM0000191582 |
| PURU_RSK _02c | Punctelia rudecta | 2022-04-26 | CAM-NSF-Lichen Genomics Team; Allen, Jessica L. | USA: Monongahela National Forest, S-facing slopes of Red Spruce Knob, N of Red Lick Mountain, 0.1 mi W of WV150 and FSR451 | New York State Museum | 38.323665 N 80.152203 W | NYSM0000191581 |

| | | | | | | | |
|---|---|---|---|---|---|---|---|
| PURU_RSK_02b | Punctelia rudecta | 2022-04-26 | CAM-NSF-Lichen Genomics Team; Allen, Jessica L. | USA: Monongahela National Forest, S-facing slopes of Red Spruce Knob, N of Red Lick Mountain, 0.1 mi W of WV150 and FSR451 | New York State Museum | 38.323665 N 80.152203 W | NYSM0000191580 |
| PURU_RSK_02a | Punctelia rudecta | 2022-04-26 | CAM-NSF-Lichen Genomics Team; Allen, Jessica L. | USA: Monongahela National Forest, S-facing slopes of Red Spruce Knob, N of Red Lick Mountain, 0.1 mi W of WV150 and FSR451 | New York State Museum | 38.323665 N 80.152203 W | NYSM0000191579 |
| PURU_RSK_01c | Punctelia rudecta | 2022-04-26 | CAM-NSF-Lichen Genomics Team; Allen, Jessica L. | USA: Monongahela National Forest, S-facing slopes of Red Spruce Knob, N of Red Lick Mountain, 0.1 mi W of WV150 and FSR451 | New York State Museum | 38.323665 N 80.152203 W | NYSM0000191578 |
| PURU_RSK_01b | Punctelia rudecta | 2022-04-26 | CAM-NSF-Lichen Genomics Team; Allen, Jessica L. | USA: Monongahela National Forest, S-facing slopes of Red Spruce Knob, N of Red Lick Mountain, 0.1 mi W of WV150 and FSR451 | New York State Museum | 38.323665 N 80.152203 W | NYSM0000191577 |
| PURU_RSK_01a | Punctelia rudecta | 2022-04-26 | CAM-NSF-Lichen Genomics Team; Allen, Jessica L. | USA: Monongahela National Forest, S-facing slopes of Red Spruce Knob, N of Red Lick Mountain, 0.1 mi W of WV150 and FSR451 | New York State Museum | 38.323665 N 80.152203 W | NYSM0000191576 |
| PURU_RON_04b | Punctelia rudecta | 2022-04-19 | CAM-NSF-Lichen Genomics Team; Allen, Jessica L. | USA: Jefferson National Forest, Round Mountain, vicinity of summit, Round Mountain Rd./VA688, ~4 mi W of US52/Bastian | New York State Museum | 37.154716 N 81.194163 W | NYSM0000191575 |

| | | | | | | | |
|---|---|---|---|---|---|---|---|
| PURU_RON _04a | Punctelia rudecta | 2022-04-19 | CAM-NSF-Lichen Genomics Team; Allen, Jessica L. | USA: Jefferson National Forest, Round Mountain, vicinity of summit, Round Mountain Rd./VA688, ~4 mi W of US52/Bastian | New York State Museum | 37.154716 N 81.194163 W | NYSM0000191574 |
| PURU_RON _03c | Punctelia rudecta | 2022-04-19 | CAM-NSF-Lichen Genomics Team; Allen, Jessica L. | USA: Jefferson National Forest, Round Mountain, vicinity of summit, Round Mountain Rd./VA688, ~4 mi W of US52/Bastian | New York State Museum | 37.154716 N 81.194163 W | NYSM0000191573 |
| PURU_RON _03b | Punctelia rudecta | 2022-04-19 | CAM-NSF-Lichen Genomics Team; Allen, Jessica L. | USA: Jefferson National Forest, Round Mountain, vicinity of summit, Round Mountain Rd./VA688, ~4 mi W of US52/Bastian | New York State Museum | 37.154716 N 81.194163 W | NYSM0000191572 |
| PURU_RON _03a | Punctelia rudecta | 2022-04-19 | CAM-NSF-Lichen Genomics Team; Allen, Jessica L. | USA: Jefferson National Forest, Round Mountain, vicinity of summit, Round Mountain Rd./VA688, ~4 mi W of US52/Bastian | New York State Museum | 37.154716 N 81.194163 W | NYSM0000191571 |
| PURU_RON _02c | Punctelia rudecta | 2022-04-19 | CAM-NSF-Lichen Genomics Team; Allen, Jessica L. | USA: Jefferson National Forest, Round Mountain, vicinity of summit, Round Mountain Rd./VA688, ~4 mi W of US52/Bastian | New York State Museum | 37.154716 N 81.194163 W | NYSM0000191570 |
| PURU_RON _02b | Punctelia rudecta | 2022-04-19 | CAM-NSF-Lichen Genomics Team; Allen, Jessica L. | USA: Jefferson National Forest, Round Mountain, vicinity of summit, Round Mountain Rd./VA688, ~4 mi W of US52/Bastian | New York State Museum | 37.154716 N 81.194163 W | NYSM0000191569 |

| | | | | | | | |
|---|---|---|---|---|---|---|---|
| PURU_RON_02a | Punctelia rudecta | 2022-04-19 | CAM-NSF-Lichen Genomics Team; Allen, Jessica L. | USA: Jefferson National Forest, Round Mountain, vicinity of summit, Round Mountain Rd./VA688, ~4 mi W of US52/Bastian | New York State Museum | 37.154716 N 81.194163 W | NYSM0000191568 |
| PURU_RON_01c | Punctelia rudecta | 2022-04-19 | CAM-NSF-Lichen Genomics Team; Allen, Jessica L. | USA: Jefferson National Forest, Round Mountain, vicinity of summit, Round Mountain Rd./VA688, ~4 mi W of US52/Bastian | New York State Museum | 37.154716 N 81.194163 W | NYSM0000191567 |
| PURU_RON_01b | Punctelia rudecta | 2022-04-19 | CAM-NSF-Lichen Genomics Team; Allen, Jessica L. | USA: Jefferson National Forest, Round Mountain, vicinity of summit, Round Mountain Rd./VA688, ~4 mi W of US52/Bastian | New York State Museum | 37.154716 N 81.194163 W | NYSM0000191566 |
| PURU_RON_01a | Punctelia rudecta | 2022-04-19 | CAM-NSF-Lichen Genomics Team; Allen, Jessica L. | USA: Jefferson National Forest, Round Mountain, vicinity of summit, Round Mountain Rd./VA688, ~4 mi W of US52/Bastian | New York State Museum | 37.154716 N 81.194163 W | NYSM0000191565 |
| PURU_ROG_06b | Punctelia rudecta | 2022-04-16 | CAM-NSF-Lichen Genomics Team; Allen, Jessica L. | USA: Jefferson National Forest, Mount Rogers National Recreation Area, Lewis Fork Wilderness, S-facing slopes of Mount Rogers, Mount Rogers Spur Trail 0.25 mi E of summit, ~0.25 mi W of jct w/ Appalachian Trail | New York State Museum | 36.659449 N 81.540821 W | NYSM0000191563 |

| | | | | | | | |
|---|---|---|---|---|---|---|---|
| PURU_ROG_06a | Punctelia rudecta | 2022-04-16 | CAM-NSF-Lichen Genomics Team; Allen, Jessica L. | USA: Jefferson National Forest, Mount Rogers National Recreation Area, Lewis Fork Wilderness, S-facing slopes of Mount Rogers, Mount Rogers Spur Trail 0.25 mi E of summit, ~0.25 mi W of jct w/ Appalachian Trail | New York State Museum | 36.659449 N 81.540821 W | NYSM0000191562 |
| PURU_ROG_05a | Punctelia rudecta | 2022-04-16 | CAM-NSF-Lichen Genomics Team; Allen, Jessica L. | USA: Jefferson National Forest, Mount Rogers National Recreation Area, Lewis Fork Wilderness, S-facing slopes of Mount Rogers, Mount Rogers Spur Trail 0.25 mi E of summit, ~0.25 mi W of jct w/ Appalachian Trail | New York State Museum | 36.659449 N 81.540821 W | NYSM0000191561 |
| PURU_ROG_056c | Punctelia rudecta | 2022-04-16 | CAM-NSF-Lichen Genomics Team; Allen, Jessica L. | USA: Jefferson National Forest, Mount Rogers National Recreation Area, Lewis Fork Wilderness, S-facing slopes of Mount Rogers, Mount Rogers Spur Trail 0.25 mi E of summit, ~0.25 mi W of jct w/ Appalachian Trail | New York State Museum | 36.659449 N 81.540821 W | NYSM0000191564 |
| PURU_ROG_04c | Punctelia rudecta | 2022-04-16 | CAM-NSF-Lichen Genomics Team; Allen, Jessica L. | USA: Jefferson National Forest, Mount Rogers National Recreation Area, Lewis Fork Wilderness, S-facing slopes of Mount Rogers, Mount Rogers Spur Trail 0.25 mi E of summit, ~0.25 mi W of jct w/ Appalachian Trail | New York State Museum | 36.659449 N 81.540821 W | NYSM0000191560 |

| | | | | | | | |
|---|---|---|---|---|---|---|---|
| PURU_ROG _04b | Punctelia rudecta | 2022-04-16 | CAM-NSF-Lichen Genomics Team; Allen, Jessica L. | USA: Jefferson National Forest, Mount Rogers National Recreation Area, Lewis Fork Wilderness, S-facing slopes of Mount Rogers, Mount Rogers Spur Trail 0.25 mi E of summit, ~0.25 mi W of jct w/ Appalachian Trail | New York State Museum | 36.659449 N 81.540821 W | NYSM0000191559 |
| PURU_ROG _04a | Punctelia rudecta | 2022-04-16 | CAM-NSF-Lichen Genomics Team; Allen, Jessica L. | USA: Jefferson National Forest, Mount Rogers National Recreation Area, Lewis Fork Wilderness, S-facing slopes of Mount Rogers, Mount Rogers Spur Trail 0.25 mi E of summit, ~0.25 mi W of jct w/ Appalachian Trail | New York State Museum | 36.659449 N 81.540821 W | NYSM0000191558 |
| PURU_ROG _01a | Punctelia rudecta | 2022-04-16 | CAM-NSF-Lichen Genomics Team; Allen, Jessica L. | USA: Jefferson National Forest, Mount Rogers National Recreation Area, Lewis Fork Wilderness, S-facing slopes of Mount Rogers, Mount Rogers Spur Trail 0.25 mi E of summit, ~0.25 mi W of jct w/ Appalachian Trail | New York State Museum | 36.659449 N 81.540821 W | NYSM0000191557 |
| PURU_POK _04a | Punctelia rudecta | 2022-04-24 | CAM-NSF-Lichen Genomics Team; Allen, Jessica L. | USA: Monongahela National Forest, W-facing slopes of Canaan Mountain, E of Pointy Knob, Pointy Knob Trail ~2.5 mi S of FSR13 | New York State Museum | 39.035283 N 79.489141 W | NYSM0000191556 |

| | | | | | | | |
|---|---|---|---|---|---|---|---|
| PURU_POK _03c | Punctelia rudecta | 2022-04-24 | CAM-NSF-Lichen Genomics Team; Allen, Jessica L. | USA: Monongahela National Forest, W-facing slopes of Canaan Mountain, E of Pointy Knob, Pointy Knob Trail ~2.5 mi S of FSR13 | New York State Museum | 39.035283 N 79.489141 W | NYSM0000191555 |
| PURU_POK _03b | Punctelia rudecta | 2022-04-24 | CAM-NSF-Lichen Genomics Team; Allen, Jessica L. | USA: Monongahela National Forest, W-facing slopes of Canaan Mountain, E of Pointy Knob, Pointy Knob Trail ~2.5 mi S of FSR13 | New York State Museum | 39.035283 N 79.489141 W | NYSM0000191554 |
| PURU_POK _03a | Punctelia rudecta | 2022-04-24 | CAM-NSF-Lichen Genomics Team; Allen, Jessica L. | USA: Monongahela National Forest, W-facing slopes of Canaan Mountain, E of Pointy Knob, Pointy Knob Trail ~2.5 mi S of FSR13 | New York State Museum | 39.035283 N 79.489141 W | NYSM0000191553 |
| PURU_POK _02c | Punctelia rudecta | 2022-04-24 | CAM-NSF-Lichen Genomics Team; Allen, Jessica L. | USA: Monongahela National Forest, W-facing slopes of Canaan Mountain, E of Pointy Knob, Pointy Knob Trail ~2.5 mi S of FSR13 | New York State Museum | 39.035283 N 79.489141 W | NYSM0000191552 |
| PURU_POK _02b | Punctelia rudecta | 2022-04-24 | CAM-NSF-Lichen Genomics Team; Allen, Jessica L. | USA: Monongahela National Forest, W-facing slopes of Canaan Mountain, E of Pointy Knob, Pointy Knob Trail ~2.5 mi S of FSR13 | New York State Museum | 39.035283 N 79.489141 W | NYSM0000191551 |
| PURU_POK _02a | Punctelia rudecta | 2022-04-24 | CAM-NSF-Lichen Genomics Team; Allen, Jessica L. | USA: Monongahela National Forest, W-facing slopes of Canaan Mountain, E of Pointy Knob, Pointy Knob Trail ~2.5 mi S of FSR13 | New York State Museum | 39.035283 N 79.489141 W | NYSM0000191550 |

| | | | | | | | |
|---|---|---|---|---|---|---|---|
| PURU_POK _01c | Punctelia rudecta | 2022-04-24 | CAM-NSF-Lichen Genomics Team; Allen, Jessica L. | USA: Monongahela National Forest, W-facing slopes of Canaan Mountain, E of Pointy Knob, Pointy Knob Trail ~2.5 mi S of FSR13 | New York State Museum | 39.035283 N 79.489141 W | NYSM0000191549 |
| PURU_POK _01b | Punctelia rudecta | 2022-04-24 | CAM-NSF-Lichen Genomics Team; Allen, Jessica L. | USA: Monongahela National Forest, W-facing slopes of Canaan Mountain, E of Pointy Knob, Pointy Knob Trail ~2.5 mi S of FSR13 | New York State Museum | 39.035283 N 79.489141 W | NYSM0000191548 |
| PURU_POK _01a | Punctelia rudecta | 2022-04-24 | CAM-NSF-Lichen Genomics Team; Allen, Jessica L. | USA: Monongahela National Forest, W-facing slopes of Canaan Mountain, E of Pointy Knob, Pointy Knob Trail ~2.5 mi S of FSR13 | New York State Museum | 39.035283 N 79.489141 W | NYSM0000191547 |
| PURU_NGA _13c | Punctelia rudecta | 2022-04-28 | CAM-NSF-Lichen Genomics Team; Allen, Jessica L. | USA: Monongahela National Forest, Shavers Mountain, E-facing slopes of Gaudineer Knob, FSR27 ~1.25 mi N of jct w/ US250 | New York State Museum | 38.611691 N 79.841629 W | NYSM0000191546 |
| PURU_NGA _13b | Punctelia rudecta | 2022-04-28 | CAM-NSF-Lichen Genomics Team; Allen, Jessica L. | USA: Monongahela National Forest, Shavers Mountain, E-facing slopes of Gaudineer Knob, FSR27 ~1.25 mi N of jct w/ US250 | New York State Museum | 38.611691 N 79.841629 W | NYSM0000191545 |
| PURU_NGA _13a | Punctelia rudecta | 2022-04-28 | CAM-NSF-Lichen Genomics Team; Allen, Jessica L. | USA: Monongahela National Forest, Shavers Mountain, E-facing slopes of Gaudineer Knob, FSR27 ~1.25 mi N of jct w/ US250 | New York State Museum | 38.611691 N 79.841629 W | NYSM0000191544 |

| | | | | | | | |
|---|---|---|---|---|---|---|---|
| PURU_NGA _12c | Punctelia rudecta | 2022-04-28 | CAM-NSF-Lichen Genomics Team; Allen, Jessica L. | USA: Monongahela National Forest, Shavers Mountain, E-facing slopes of Gaudineer Knob, FSR27 ~1.25 mi N of jct w/ US250 | New York State Museum | 38.611691 N 79.841629 W | NYSM0000191543 |
| PURU_NGA _12b | Punctelia rudecta | 2022-04-28 | CAM-NSF-Lichen Genomics Team; Allen, Jessica L. | USA: Monongahela National Forest, Shavers Mountain, E-facing slopes of Gaudineer Knob, FSR27 ~1.25 mi N of jct w/ US250 | New York State Museum | 38.611691 N 79.841629 W | NYSM0000191542 |
| PURU_NGA _12a | Punctelia rudecta | 2022-04-28 | CAM-NSF-Lichen Genomics Team; Allen, Jessica L. | USA: Monongahela National Forest, Shavers Mountain, E-facing slopes of Gaudineer Knob, FSR27 ~1.25 mi N of jct w/ US250 | New York State Museum | 38.611691 N 79.841629 W | NYSM0000191541 |
| PURU_NGA _11a | Punctelia rudecta | 2022-04-28 | CAM-NSF-Lichen Genomics Team; Allen, Jessica L. | USA: Monongahela National Forest, Shavers Mountain, E-facing slopes of Gaudineer Knob, FSR27 ~1.25 mi N of jct w/ US250 | New York State Museum | 38.611691 N 79.841629 W | NYSM0000191540 |
| PURU_NAR _07b | Punctelia rudecta | 2022-08-04 | CAM-NSF-Lichen Genomics Team; Denzer, Eli | USA: Pisgah National Forest, Balsam Mountains, Shining Rock Wilderness, Shining Rock Ledge, E slopes of Stairs Mountain summit, Art Loeb Trail 1 mi N of Shining Rock Gap | New York State Museum | 35.37817 N 82.86893 W | |

| | | | | | | |
|---|---|---|---|---|---|---|
| PURU_NAR_07a | Punctelia rudecta | 2022-08-04 | CAM-NSF-Lichen Genomics Team; Denzer, Eli | USA: Pisgah National Forest, Balsam Mountains, Shining Rock Wilderness, Shining Rock Ledge, E slopes of Stairs Mountain summit, Art Loeb Trail 1 mi N of Shining Rock Gap | New York State Museum | 35.37817 N 82.86893 W |
| PURU_NAR_05c | Punctelia rudecta | 2022-08-04 | CAM-NSF-Lichen Genomics Team; Denzer, Eli | USA: Pisgah National Forest, Balsam Mountains, Shining Rock Wilderness, Shining Rock Ledge, E slopes of Stairs Mountain summit, Art Loeb Trail 1 mi N of Shining Rock Gap | New York State Museum | 35.37817 N 82.86893 W |
| PURU_NAR_05b | Punctelia rudecta | 2022-08-04 | CAM-NSF-Lichen Genomics Team; Denzer, Eli | USA: Pisgah National Forest, Balsam Mountains, Shining Rock Wilderness, Shining Rock Ledge, E slopes of Stairs Mountain summit, Art Loeb Trail 1 mi N of Shining Rock Gap | New York State Museum | 35.37817 N 82.86893 W |
| PURU_NAR_05a | Punctelia rudecta | 2022-08-04 | CAM-NSF-Lichen Genomics Team; Denzer, Eli | USA: Pisgah National Forest, Balsam Mountains, Shining Rock Wilderness, Shining Rock Ledge, E slopes of Stairs Mountain summit, Art Loeb Trail 1 mi N of Shining Rock Gap | New York State Museum | 35.37817 N 82.86893 W |
| PURU_NAR_04c | Punctelia rudecta | 2022-08-04 | CAM-NSF-Lichen Genomics Team; Denzer, Eli | USA: Pisgah National Forest, Balsam Mountains, Shining Rock Wilderness, Shining Rock Ledge, E slopes of Stairs Mountain summit, Art Loeb Trail 1 mi N of Shining Rock Gap | New York State Museum | 35.37817 N 82.86893 W |

| | | | | | | | |
|---|---|---|---|---|---|---|---|
| PURU_NAR_04b | Punctelia rudecta | 2022-08-04 | CAM-NSF-Lichen Genomics Team; Denzer, Eli | USA: Pisgah National Forest, Balsam Mountains, Shining Rock Wilderness, Shining Rock Ledge, E slopes of Stairs Mountain summit, Art Loeb Trail 1 mi N of Shining Rock Gap | New York State Museum | 35.37817 N 82.86893 W | |
| PURU_NAR_04a | Punctelia rudecta | 2022-08-04 | CAM-NSF-Lichen Genomics Team; Denzer, Eli | USA: Pisgah National Forest, Balsam Mountains, Shining Rock Wilderness, Shining Rock Ledge, E slopes of Stairs Mountain summit, Art Loeb Trail 1 mi N of Shining Rock Gap | New York State Museum | 35.37817 N 82.86893 W | |
| PURU_LUE_04c | Punctelia rudecta | 2022-04-26 | CAM-NSF-Lichen Genomics Team; Allen, Jessica L. | USA: Monongahela National Forest, Spruce Mountain, W-facing slopes below summit of Blue Knob, ~1 mi S of WV39 | New York State Museum | 38.175865 N 80.279981 W | NYSM0000191539 |
| PURU_LUE_04b | Punctelia rudecta | 2022-04-26 | CAM-NSF-Lichen Genomics Team; Allen, Jessica L. | USA: Monongahela National Forest, Spruce Mountain, W-facing slopes below summit of Blue Knob, ~1 mi S of WV39 | New York State Museum | 38.175865 N 80.279981 W | NYSM0000191538 |
| PURU_LUE_04a | Punctelia rudecta | 2022-04-26 | CAM-NSF-Lichen Genomics Team; Allen, Jessica L. | USA: Monongahela National Forest, Spruce Mountain, W-facing slopes below summit of Blue Knob, ~1 mi S of WV39 | New York State Museum | 38.175865 N 80.279981 W | NYSM0000191537 |

| | | | | | | | |
|---|---|---|---|---|---|---|---|
| PURU_LUE_03c | Punctelia rudecta | 2022-04-26 | CAM-NSF-Lichen Genomics Team; Allen, Jessica L. | USA: Monongahela National Forest, Spruce Mountain, W-facing slopes below summit of Blue Knob, ~1 mi S of WV39 | New York State Museum | 38.175865 N 80.279981 W | NYSM0000191536 |
| PURU_LUE_03b | Punctelia rudecta | 2022-04-26 | CAM-NSF-Lichen Genomics Team; Allen, Jessica L. | USA: Monongahela National Forest, Spruce Mountain, W-facing slopes below summit of Blue Knob, ~1 mi S of WV39 | New York State Museum | 38.175865 N 80.279981 W | NYSM0000191535 |
| PURU_LUE_03a | Punctelia rudecta | 2022-04-26 | CAM-NSF-Lichen Genomics Team; Allen, Jessica L. | USA: Monongahela National Forest, Spruce Mountain, W-facing slopes below summit of Blue Knob, ~1 mi S of WV39 | New York State Museum | 38.175865 N 80.279981 W | NYSM0000191534 |
| PURU_LUE_02c | Punctelia rudecta | 2022-04-26 | CAM-NSF-Lichen Genomics Team; Allen, Jessica L. | USA: Monongahela National Forest, Spruce Mountain, W-facing slopes below summit of Blue Knob, ~1 mi S of WV39 | New York State Museum | 38.175865 N 80.279981 W | NYSM0000191533 |
| PURU_LUE_02b | Punctelia rudecta | 2022-04-26 | CAM-NSF-Lichen Genomics Team; Allen, Jessica L. | USA: Monongahela National Forest, Spruce Mountain, W-facing slopes below summit of Blue Knob, ~1 mi S of WV39 | New York State Museum | 38.175865 N 80.279981 W | NYSM0000191532 |
| PURU_LUE_02a | Punctelia rudecta | 2022-04-26 | CAM-NSF-Lichen Genomics Team; Allen, Jessica L. | USA: Monongahela National Forest, Spruce Mountain, W-facing slopes below summit of Blue Knob, ~1 mi S of WV39 | New York State Museum | 38.175865 N 80.279981 W | NYSM0000191531 |

| | | | | | | | |
|---|---|---|---|---|---|---|---|
| PURU_LUE_01c | Punctelia rudecta | 2022-04-26 | CAM-NSF-Lichen Genomics Team; Allen, Jessica L. | USA: Monongahela National Forest, Spruce Mountain, W-facing slopes below summit of Blue Knob, ~1 mi S of WV39 | New York State Museum | 38.175865 N 80.279981 W | NYSM0000191530 |
| PURU_LUE_01b | Punctelia rudecta | 2022-04-26 | CAM-NSF-Lichen Genomics Team; Allen, Jessica L. | USA: Monongahela National Forest, Spruce Mountain, W-facing slopes below summit of Blue Knob, ~1 mi S of WV39 | New York State Museum | 38.175865 N 80.279981 W | NYSM0000191529 |
| PURU_LUE_01a | Punctelia rudecta | 2022-04-26 | CAM-NSF-Lichen Genomics Team; Allen, Jessica L. | USA: Monongahela National Forest, Spruce Mountain, W-facing slopes below summit of Blue Knob, ~1 mi S of WV39 | New York State Museum | 38.175865 N 80.279981 W | NYSM0000191528 |
| PURU_LEM_08c | Punctelia rudecta | 2022-04-13 | CAM-NSF-Lichen Genomics Team; Allen, Jessica L. | USA: Pine Mountain, Bad Branch Nature Preserve, W headwaters of Bad Branch, Pine Mountain Trail ~2.2 mi E of US119 | New York State Museum | 37.09032 N 82.77928 W | NYSM0000191527 |
| PURU_LEM_08b | Punctelia rudecta | 2022-04-13 | CAM-NSF-Lichen Genomics Team; Allen, Jessica L. | USA: Pine Mountain, Bad Branch Nature Preserve, W headwaters of Bad Branch, Pine Mountain Trail ~2.2 mi E of US119 | New York State Museum | 37.09032 N 82.77928 W | NYSM0000191526 |
| PURU_LEM_08a | Punctelia rudecta | 2022-04-13 | CAM-NSF-Lichen Genomics Team; Allen, Jessica L. | USA: Pine Mountain, Bad Branch Nature Preserve, W headwaters of Bad Branch, Pine Mountain Trail ~2.2 mi E of US119 | New York State Museum | 37.09032 N 82.77928 W | NYSM0000191525 |

| | | | | | | | |
|---|---|---|---|---|---|---|---|
| PURU_LEM_05c | Punctelia rudecta | 2022-04-13 | CAM-NSF-Lichen Genomics Team; Allen, Jessica L. | USA: Pine Mountain, Bad Branch Nature Preserve, W headwaters of Bad Branch, Pine Mountain Trail ~2.2 mi E of US119 | New York State Museum | 37.09032 N 82.77928 W | NYSM0000191524 |
| PURU_LEM_05b | Punctelia rudecta | 2022-04-13 | CAM-NSF-Lichen Genomics Team; Allen, Jessica L. | USA: Pine Mountain, Bad Branch Nature Preserve, W headwaters of Bad Branch, Pine Mountain Trail ~2.2 mi E of US119 | New York State Museum | 37.09032 N 82.77928 W | NYSM0000191523 |
| PURU_LEM_05a | Punctelia rudecta | 2022-04-13 | CAM-NSF-Lichen Genomics Team; Allen, Jessica L. | USA: Pine Mountain, Bad Branch Nature Preserve, W headwaters of Bad Branch, Pine Mountain Trail ~2.2 mi E of US119 | New York State Museum | 37.09032 N 82.77928 W | NYSM0000191522 |
| PURU_LEM_04a | Punctelia rudecta | 2022-04-13 | CAM-NSF-Lichen Genomics Team; Allen, Jessica L. | USA: Pine Mountain, Bad Branch Nature Preserve, W headwaters of Bad Branch, Pine Mountain Trail ~2.2 mi E of US119 | New York State Museum | 37.09032 N 82.77928 W | NYSM0000191521 |
| PURU_LEM_03b | Punctelia rudecta | 2022-04-13 | CAM-NSF-Lichen Genomics Team; Allen, Jessica L. | USA: Pine Mountain, Bad Branch Nature Preserve, W headwaters of Bad Branch, Pine Mountain Trail ~2.2 mi E of US119 | New York State Museum | 37.09032 N 82.77928 W | NYSM0000191520 |
| PURU_LEM_03a | Punctelia rudecta | 2022-04-13 | CAM-NSF-Lichen Genomics Team; Allen, Jessica L. | USA: Pine Mountain, Bad Branch Nature Preserve, W headwaters of Bad Branch, Pine Mountain Trail ~2.2 mi E of US119 | New York State Museum | 37.09032 N 82.77928 W | NYSM0000191519 |

| | | | | | | | |
|---|---|---|---|---|---|---|---|
| PURU_LEM _01c | Punctelia rudecta | 2022-04-13 | CAM-NSF-Lichen Genomics Team; Allen, Jessica L. | USA: Pine Mountain, Bad Branch Nature Preserve, W headwaters of Bad Branch, Pine Mountain Trail ~2.2 mi E of US119 | New York State Museum | 37.09032 N 82.77928 W | NYSM0000191518 |
| PURU_LEM _01b | Punctelia rudecta | 2022-04-13 | CAM-NSF-Lichen Genomics Team; Allen, Jessica L. | USA: Pine Mountain, Bad Branch Nature Preserve, W headwaters of Bad Branch, Pine Mountain Trail ~2.2 mi E of US119 | New York State Museum | 37.09032 N 82.77928 W | NYSM0000191517 |
| PURU_LEM _01a | Punctelia rudecta | 2022-04-13 | CAM-NSF-Lichen Genomics Team; Allen, Jessica L. | USA: Pine Mountain, Bad Branch Nature Preserve, W headwaters of Bad Branch, Pine Mountain Trail ~2.2 mi E of US119 | New York State Museum | 37.09032 N 82.77928 W | NYSM0000191516 |
| PURU_KLE _04c | Punctelia rudecta | 2022-04-25 | CAM-NSF-Lichen Genomics Team; Allen, Jessica L. | USA: Monongahela National Forest, Bickle Knob, saddle and W-facing slopes below observation tower, terminus of FSR91A | New York State Museum | 38.93557 N 79.730504 W | NYSM0000191514 |
| PURU_KLE _04b | Punctelia rudecta | 2022-04-25 | CAM-NSF-Lichen Genomics Team; Allen, Jessica L. | USA: Monongahela National Forest, Bickle Knob, saddle and W-facing slopes below observation tower, terminus of FSR91A | New York State Museum | 38.93557 N 79.730504 W | NYSM0000191512 |
| PURU_KLE _04a | Punctelia rudecta | 2022-04-25 | CAM-NSF-Lichen Genomics Team; Allen, Jessica L. | USA: Monongahela National Forest, Bickle Knob, saddle and W-facing slopes below observation tower, terminus of FSR91A | New York State Museum | 38.93557 N 79.730504 W | NYSM0000191510 |

| | | | | | | | |
|---|---|---|---|---|---|---|---|
| PURU_KLE _03c | Punctelia rudecta | 2022-04-25 | CAM-NSF-Lichen Genomics Team; Allen, Jessica L. | USA: Monongahela National Forest, Bickle Knob, saddle and W-facing slopes below observation tower, terminus of FSR91A | New York State Museum | 38.93557 N 79.730504 W | NYSM0000191509 |
| PURU_KLE _03b | Punctelia rudecta | 2022-04-25 | CAM-NSF-Lichen Genomics Team; Allen, Jessica L. | USA: Monongahela National Forest, Bickle Knob, saddle and W-facing slopes below observation tower, terminus of FSR91A | New York State Museum | 38.93557 N 79.730504 W | NYSM0000191508 |
| PURU_KLE _03a | Punctelia rudecta | 2022-04-25 | CAM-NSF-Lichen Genomics Team; Allen, Jessica L. | USA: Monongahela National Forest, Bickle Knob, saddle and W-facing slopes below observation tower, terminus of FSR91A | New York State Museum | 38.93557 N 79.730504 W | NYSM0000191507 |
| PURU_KLE _02c | Punctelia rudecta | 2022-04-25 | CAM-NSF-Lichen Genomics Team; Allen, Jessica L. | USA: Monongahela National Forest, Bickle Knob, saddle and W-facing slopes below observation tower, terminus of FSR91A | New York State Museum | 38.93557 N 79.730504 W | NYSM0000191506 |
| PURU_KLE _02b | Punctelia rudecta | 2022-04-25 | CAM-NSF-Lichen Genomics Team; Allen, Jessica L. | USA: Monongahela National Forest, Bickle Knob, saddle and W-facing slopes below observation tower, terminus of FSR91A | New York State Museum | 38.93557 N 79.730504 W | NYSM0000191505 |
| PURU_KLE _02a | Punctelia rudecta | 2022-04-25 | CAM-NSF-Lichen Genomics Team; Allen, Jessica L. | USA: Monongahela National Forest, Bickle Knob, saddle and W-facing slopes below observation tower, terminus of FSR91A | New York State Museum | 38.93557 N 79.730504 W | NYSM0000191504 |

| | | | | | | | |
|---|---|---|---|---|---|---|---|
| PURU_KLE_01c | Punctelia rudecta | 2022-04-25 | CAM-NSF-Lichen Genomics Team; Allen, Jessica L. | USA: Monongahela National Forest, Bickle Knob, saddle and W-facing slopes below observation tower, terminus of FSR91A | New York State Museum | 38.93557 N 79.730504 W | NYSM0000191503 |
| PURU_KLE_01b | Punctelia rudecta | 2022-04-25 | CAM-NSF-Lichen Genomics Team; Allen, Jessica L. | USA: Monongahela National Forest, Bickle Knob, saddle and W-facing slopes below observation tower, terminus of FSR91A | New York State Museum | 38.93557 N 79.730504 W | NYSM0000191502 |
| PURU_KLE_01a | Punctelia rudecta | 2022-04-25 | CAM-NSF-Lichen Genomics Team; Allen, Jessica L. | USA: Monongahela National Forest, Bickle Knob, saddle and W-facing slopes below observation tower, terminus of FSR91A | New York State Museum | 38.93557 N 79.730504 W | NYSM0000191501 |
| PURU_JON_04c | Punctelia rudecta | 2022-04-19 | CAM-NSF-Lichen Genomics Team; Allen, Jessica L. | USA: Jefferson National Forest, E slopes of Round Mountain, S of Round Mountain Rd./VA688, ~1.5 mi W of US52/Bastian | New York State Museum | 37.170518 N 81.164224 W | NYSM0000191500 |
| PURU_JON_04b | Punctelia rudecta | 2022-04-19 | CAM-NSF-Lichen Genomics Team; Allen, Jessica L. | USA: Jefferson National Forest, E slopes of Round Mountain, S of Round Mountain Rd./VA688, ~1.5 mi W of US52/Bastian | New York State Museum | 37.170518 N 81.164224 W | NYSM0000191499 |
| PURU_JON_04a | Punctelia rudecta | 2022-04-19 | CAM-NSF-Lichen Genomics Team; Allen, Jessica L. | USA: Jefferson National Forest, E slopes of Round Mountain, S of Round Mountain Rd./VA688, ~1.5 mi W of US52/Bastian | New York State Museum | 37.170518 N 81.164224 W | NYSM0000191498 |

| | | | | | | | |
|---|---|---|---|---|---|---|---|
| PURU_JON _03c | Punctelia rudecta | 2022-04-19 | CAM-NSF-Lichen Genomics Team; Allen, Jessica L. | USA: Jefferson National Forest, E slopes of Round Mountain, S of Round Mountain Rd./VA688, ~1.5 mi W of US52/Bastian | New York State Museum | 37.170518 N 81.164224 W | NYSM0000191497 |
| PURU_JON _03b | Punctelia rudecta | 2022-04-19 | CAM-NSF-Lichen Genomics Team; Allen, Jessica L. | USA: Jefferson National Forest, E slopes of Round Mountain, S of Round Mountain Rd./VA688, ~1.5 mi W of US52/Bastian | New York State Museum | 37.170518 N 81.164224 W | NYSM0000191496 |
| PURU_JON _03a | Punctelia rudecta | 2022-04-19 | CAM-NSF-Lichen Genomics Team; Allen, Jessica L. | USA: Jefferson National Forest, E slopes of Round Mountain, S of Round Mountain Rd./VA688, ~1.5 mi W of US52/Bastian | New York State Museum | 37.170518 N 81.164224 W | NYSM0000191495 |
| PURU_JON _02c | Punctelia rudecta | 2022-04-19 | CAM-NSF-Lichen Genomics Team; Allen, Jessica L. | USA: Jefferson National Forest, E slopes of Round Mountain, S of Round Mountain Rd./VA688, ~1.5 mi W of US52/Bastian | New York State Museum | 37.170518 N 81.164224 W | NYSM0000191494 |
| PURU_JON _02b | Punctelia rudecta | 2022-04-19 | CAM-NSF-Lichen Genomics Team; Allen, Jessica L. | USA: Jefferson National Forest, E slopes of Round Mountain, S of Round Mountain Rd./VA688, ~1.5 mi W of US52/Bastian | New York State Museum | 37.170518 N 81.164224 W | NYSM0000191493 |
| PURU_JON _02a | Punctelia rudecta | 2022-04-19 | CAM-NSF-Lichen Genomics Team; Allen, Jessica L. | USA: Jefferson National Forest, E slopes of Round Mountain, S of Round Mountain Rd./VA688, ~1.5 mi W of US52/Bastian | New York State Museum | 37.170518 N 81.164224 W | NYSM0000191492 |

| | | | | | | | |
|---|---|---|---|---|---|---|---|
| PURU_JON_01c | Punctelia rudecta | 2022-04-19 | CAM-NSF-Lichen Genomics Team; Allen, Jessica L. | USA: Jefferson National Forest, E slopes of Round Mountain, S of Round Mountain Rd./VA688, ~1.5 mi W of US52/Bastian | New York State Museum | 37.170518 N 81.164224 W | NYSM0000191491 |
| PURU_JON_01b | Punctelia rudecta | 2022-04-19 | CAM-NSF-Lichen Genomics Team; Allen, Jessica L. | USA: Jefferson National Forest, E slopes of Round Mountain, S of Round Mountain Rd./VA688, ~1.5 mi W of US52/Bastian | New York State Museum | 37.170518 N 81.164224 W | NYSM0000191490 |
| PURU_JON_01a | Punctelia rudecta | 2022-04-19 | CAM-NSF-Lichen Genomics Team; Allen, Jessica L. | USA: Jefferson National Forest, E slopes of Round Mountain, S of Round Mountain Rd./VA688, ~1.5 mi W of US52/Bastian | New York State Museum | 37.170518 N 81.164224 W | NYSM0000191489 |
| PURU_HAW_07b | Punctelia rudecta | 2022-05-03 | CAM-NSF-Lichen Genomics Team; Allen, Jessica L. | USA: Shenandoah National Park, Central District, SW slopes of Hawksbill Mountain, between Lower Hawksbill Trail and Hawksbill Fire Road ~0.7 mi W of Hawksbill Gap | New York State Museum | 38.55397 N 78.39687 W | NYSM0000191488 |
| PURU_HAW_07a | Punctelia rudecta | 2022-05-03 | CAM-NSF-Lichen Genomics Team; Allen, Jessica L. | USA: Shenandoah National Park, Central District, SW slopes of Hawksbill Mountain, between Lower Hawksbill Trail and Hawksbill Fire Road ~0.7 mi W of Hawksbill Gap | New York State Museum | 38.55397 N 78.39687 W | NYSM0000194998 |

| | | | | | | | |
|---|---|---|---|---|---|---|---|
| PURU_HAW_06c | Punctelia rudecta | 2022-05-03 | CAM-NSF-Lichen Genomics Team; Allen, Jessica L. | USA: Shenandoah National Park, Central District, SW slopes of Hawksbill Mountain, between Lower Hawksbill Trail and Hawksbill Fire Road ~0.7 mi W of Hawksbill Gap | New York State Museum | 38.55397 N 78.39687 W | NYSM0000191487 |
| PURU_HAW_06b | Punctelia rudecta | 2022-05-03 | CAM-NSF-Lichen Genomics Team; Allen, Jessica L. | USA: Shenandoah National Park, Central District, SW slopes of Hawksbill Mountain, between Lower Hawksbill Trail and Hawksbill Fire Road ~0.7 mi W of Hawksbill Gap | New York State Museum | 38.55397 N 78.39687 W | NYSM0000191486 |
| PURU_HAW_06a | Punctelia rudecta | 2022-05-03 | CAM-NSF-Lichen Genomics Team; Allen, Jessica L. | USA: Shenandoah National Park, Central District, SW slopes of Hawksbill Mountain, between Lower Hawksbill Trail and Hawksbill Fire Road ~0.7 mi W of Hawksbill Gap | New York State Museum | 38.55397 N 78.39687 W | NYSM0000191485 |
| PURU_HAW_05c | Punctelia rudecta | 2022-05-03 | CAM-NSF-Lichen Genomics Team; Allen, Jessica L. | USA: Shenandoah National Park, Central District, SW slopes of Hawksbill Mountain, between Lower Hawksbill Trail and Hawksbill Fire Road ~0.7 mi W of Hawksbill Gap | New York State Museum | 38.55397 N 78.39687 W | NYSM0000191484 |
| PURU_HAW_05b | Punctelia rudecta | 2022-05-03 | CAM-NSF-Lichen Genomics Team; Allen, Jessica L. | USA: Shenandoah National Park, Central District, SW slopes of Hawksbill Mountain, between Lower Hawksbill Trail and Hawksbill Fire Road ~0.7 mi W of Hawksbill Gap | New York State Museum | 38.55397 N 78.39687 W | NYSM0000191483 |

| | | | | | | | |
|---|---|---|---|---|---|---|---|
| PURU_HAW_05a | Punctelia rudecta | 2022-05-03 | CAM-NSF-Lichen Genomics Team; Allen, Jessica L. | USA: Shenandoah National Park, Central District, SW slopes of Hawksbill Mountain, between Lower Hawksbill Trail and Hawksbill Fire Road ~0.7 mi W of Hawksbill Gap | New York State Museum | 38.55397 N 78.39687 W | NYSM0000191482 |
| PURU_HAW_03c | Punctelia rudecta | 2022-05-03 | CAM-NSF-Lichen Genomics Team; Allen, Jessica L. | USA: Shenandoah National Park, Central District, SW slopes of Hawksbill Mountain, between Lower Hawksbill Trail and Hawksbill Fire Road ~0.7 mi W of Hawksbill Gap | New York State Museum | 38.55397 N 78.39687 W | NYSM0000194997 |
| PURU_HAW_03b | Punctelia rudecta | 2022-05-03 | CAM-NSF-Lichen Genomics Team; Allen, Jessica L. | USA: Shenandoah National Park, Central District, SW slopes of Hawksbill Mountain, between Lower Hawksbill Trail and Hawksbill Fire Road ~0.7 mi W of Hawksbill Gap | New York State Museum | 38.55397 N 78.39687 W | NYSM0000191481 |
| PURU_HAW_03a | Punctelia rudecta | 2022-05-03 | CAM-NSF-Lichen Genomics Team; Allen, Jessica L. | USA: Shenandoah National Park, Central District, SW slopes of Hawksbill Mountain, between Lower Hawksbill Trail and Hawksbill Fire Road ~0.7 mi W of Hawksbill Gap | New York State Museum | 38.55397 N 78.39687 W | NYSM0000191480 |
| PURU_HAW_02c | Punctelia rudecta | 2022-05-03 | CAM-NSF-Lichen Genomics Team; Allen, Jessica L. | USA: Shenandoah National Park, Central District, SW slopes of Hawksbill Mountain, between Lower Hawksbill Trail and Hawksbill Fire Road ~0.7 mi W of Hawksbill Gap | New York State Museum | 38.55397 N 78.39687 W | NYSM0000191479 |

| | | | | | | | |
|---|---|---|---|---|---|---|---|
| PURU_HAW_02b | Punctelia rudecta | 2022-05-03 | CAM-NSF-Lichen Genomics Team; Allen, Jessica L. | USA: Shenandoah National Park, Central District, SW slopes of Hawksbill Mountain, between Lower Hawksbill Trail and Hawksbill Fire Road ~0.7 mi W of Hawksbill Gap | New York State Museum | 38.55397 N 78.39687 W | NYSM0000191478 |
| PURU_HAW_02a | Punctelia rudecta | 2022-05-03 | CAM-NSF-Lichen Genomics Team; Allen, Jessica L. | USA: Shenandoah National Park, Central District, SW slopes of Hawksbill Mountain, between Lower Hawksbill Trail and Hawksbill Fire Road ~0.7 mi W of Hawksbill Gap | New York State Museum | 38.55397 N 78.39687 W | NYSM0000191477 |
| PURU_GRI_03c | Punctelia rudecta | 2022-05-02 | CAM-NSF-Lichen Genomics Team; Allen, Jessica L. | USA: Shenandoah National Park, Central District, S facing slopes of Grindstone Mountain above Deep Run, ~0.2 mi S of summit | New York State Museum | 38.47288 N 78.550732 W | NYSM0000191476 |
| PURU_GRI_03b | Punctelia rudecta | 2022-05-02 | CAM-NSF-Lichen Genomics Team; Allen, Jessica L. | USA: Shenandoah National Park, Central District, S facing slopes of Grindstone Mountain above Deep Run, ~0.2 mi S of summit | New York State Museum | 38.47288 N 78.550732 W | NYSM0000191475 |
| PURU_GRI_03a | Punctelia rudecta | 2022-05-02 | CAM-NSF-Lichen Genomics Team; Allen, Jessica L. | USA: Shenandoah National Park, Central District, S facing slopes of Grindstone Mountain above Deep Run, ~0.2 mi S of summit | New York State Museum | 38.47288 N 78.550732 W | NYSM0000191474 |

| | | | | | | | |
|---|---|---|---|---|---|---|---|
| PURU_GRI_02c | Punctelia rudecta | 2022-05-02 | CAM-NSF-Lichen Genomics Team; Allen, Jessica L. | USA: Shenandoah National Park, Central District, S facing slopes of Grindstone Mountain above Deep Run, ~0.2 mi S of summit | New York State Museum | 38.47288 N 78.550732 W | NYSM0000191473 |
| PURU_GRI_02b | Punctelia rudecta | 2022-05-02 | CAM-NSF-Lichen Genomics Team; Allen, Jessica L. | USA: Shenandoah National Park, Central District, S facing slopes of Grindstone Mountain above Deep Run, ~0.2 mi S of summit | New York State Museum | 38.47288 N 78.550732 W | NYSM0000191472 |
| PURU_GRI_02a | Punctelia rudecta | 2022-05-02 | CAM-NSF-Lichen Genomics Team; Allen, Jessica L. | USA: Shenandoah National Park, Central District, S facing slopes of Grindstone Mountain above Deep Run, ~0.2 mi S of summit | New York State Museum | 38.47288 N 78.550732 W | NYSM0000191471 |
| PURU_GRI_01d | Punctelia rudecta | 2022-05-02 | CAM-NSF-Lichen Genomics Team; Allen, Jessica L. | USA: Shenandoah National Park, Central District, S facing slopes of Grindstone Mountain above Deep Run, ~0.2 mi S of summit | New York State Museum | 38.47288 N 78.550732 W | NYSM0000191470 |
| PURU_GRI_01c | Punctelia rudecta | 2022-05-02 | CAM-NSF-Lichen Genomics Team; Allen, Jessica L. | USA: Shenandoah National Park, Central District, S facing slopes of Grindstone Mountain above Deep Run, ~0.2 mi S of summit | New York State Museum | 38.47288 N 78.550732 W | NYSM0000191469 |
| PURU_GRI_01b | Punctelia rudecta | 2022-05-02 | CAM-NSF-Lichen Genomics Team; Allen, Jessica L. | USA: Shenandoah National Park, Central District, S facing slopes of Grindstone Mountain above Deep Run, ~0.2 mi S of summit | New York State Museum | 38.47288 N 78.550732 W | NYSM0000191468 |

| | | | | | | | |
|---|---|---|---|---|---|---|---|
| PURU_GRI_01a | Punctelia rudecta | 2022-05-02 | CAM-NSF-Lichen Genomics Team; Allen, Jessica L. | USA: Shenandoah National Park, Central District, S facing slopes of Grindstone Mountain above Deep Run, ~0.2 mi S of summit | New York State Museum | 38.47288 N 78.550732 W | NYSM0000191467 |
| PURU_GOS_06b | Punctelia rudecta | 2022-04-11 | CAM-NSF-Lichen Genomics Team; Allen, Jessica L. | USA: E-slopes of Pine Mountain, Kentenia State Forest, vicinity of Goss Park, E of Little Shepherd Trail/SR1679 at jct w/ Putney Lookout Tower Rd., between Spruce Ridge Branch and Hi Lewis Branch | New York State Museum | 36.9243 N 83.22419 W | NYSM0000191466 |
| PURU_GOS_06a | Punctelia rudecta | 2022-04-11 | CAM-NSF-Lichen Genomics Team; Allen, Jessica L. | USA: E-slopes of Pine Mountain, Kentenia State Forest, vicinity of Goss Park, E of Little Shepherd Trail/SR1679 at jct w/ Putney Lookout Tower Rd., between Spruce Ridge Branch and Hi Lewis Branch | New York State Museum | 36.9243 N 83.22419 W | NYSM0000194996 |
| PURU_GOS_05c | Punctelia rudecta | 2022-04-11 | CAM-NSF-Lichen Genomics Team; Allen, Jessica L. | USA: E-slopes of Pine Mountain, Kentenia State Forest, vicinity of Goss Park, E of Little Shepherd Trail/SR1679 at jct w/ Putney Lookout Tower Rd., between Spruce Ridge Branch and Hi Lewis Branch | New York State Museum | 36.9243 N 83.22419 W | NYSM0000191465 |
| PURU_GOS_05b | Punctelia rudecta | 2022-04-11 | CAM-NSF-Lichen Genomics Team; Allen, Jessica L. | USA: E-slopes of Pine Mountain, Kentenia State Forest, vicinity of Goss Park, E of Little Shepherd Trail/SR1679 at jct w/ Putney Lookout Tower Rd., between | New York State Museum | 36.9243 N 83.22419 W | NYSM0000191464 |

| | | | | | | | |
|---|---|---|---|---|---|---|---|
| | | | | Spruce Ridge Branch and Hi Lewis Branch | | | |
| PURU_GOS _05a | Punctelia rudecta | 2022-04-11 | CAM-NSF-Lichen Genomics Team; Allen, Jessica L. | USA: E-slopes of Pine Mountain, Kentenia State Forest, vicinity of Goss Park, E of Little Shepherd Trail/SR1679 at jct w/ Putney Lookout Tower Rd., between Spruce Ridge Branch and Hi Lewis Branch | New York State Museum | 36.9243 N 83.22419 W | NYSM0000191463 |
| PURU_GOS _04b | Punctelia rudecta | 2022-04-11 | CAM-NSF-Lichen Genomics Team; Allen, Jessica L. | USA: E-slopes of Pine Mountain, Kentenia State Forest, vicinity of Goss Park, E of Little Shepherd Trail/SR1679 at jct w/ Putney Lookout Tower Rd., between Spruce Ridge Branch and Hi Lewis Branch | New York State Museum | 36.9243 N 83.22419 W | NYSM0000191462 |
| PURU_GOS _04a | Punctelia rudecta | 2022-04-11 | CAM-NSF-Lichen Genomics Team; Allen, Jessica L. | USA: E-slopes of Pine Mountain, Kentenia State Forest, vicinity of Goss Park, E of Little Shepherd Trail/SR1679 at jct w/ Putney Lookout Tower Rd., between Spruce Ridge Branch and Hi Lewis Branch | New York State Museum | 36.9243 N 83.22419 W | NYSM0000191461 |
| PURU_GOS _03b | Punctelia rudecta | 2022-04-11 | CAM-NSF-Lichen Genomics Team; Allen, Jessica L. | USA: E-slopes of Pine Mountain, Kentenia State Forest, vicinity of Goss Park, E of Little Shepherd Trail/SR1679 at jct w/ Putney Lookout Tower Rd., between | New York State Museum | 36.9243 N 83.22419 W | NYSM0000191460 |

| | | | | | | | |
|---|---|---|---|---|---|---|---|
| | | | | Spruce Ridge Branch and Hi Lewis Branch | | | |
| PURU_GOS _03a | Punctelia rudecta | 2022-04-11 | CAM-NSF-Lichen Genomics Team; Allen, Jessica L. | USA: E-slopes of Pine Mountain, Kentenia State Forest, vicinity of Goss Park, E of Little Shepherd Trail/SR1679 at jct w/ Putney Lookout Tower Rd., between Spruce Ridge Branch and Hi Lewis Branch | New York State Museum | 36.9243 N 83.22419 W | NYSM0000194995 |
| PURU_GOS _02c | Punctelia rudecta | 2022-04-11 | CAM-NSF-Lichen Genomics Team; Allen, Jessica L. | USA: E-slopes of Pine Mountain, Kentenia State Forest, vicinity of Goss Park, E of Little Shepherd Trail/SR1679 at jct w/ Putney Lookout Tower Rd., between Spruce Ridge Branch and Hi Lewis Branch | New York State Museum | 36.9243 N 83.22419 W | NYSM0000191459 |
| PURU_GOS _02b | Punctelia rudecta | 2022-04-11 | CAM-NSF-Lichen Genomics Team; Allen, Jessica L. | USA: E-slopes of Pine Mountain, Kentenia State Forest, vicinity of Goss Park, E of Little Shepherd Trail/SR1679 at jct w/ Putney Lookout Tower Rd., between Spruce Ridge Branch and Hi Lewis Branch | New York State Museum | 36.9243 N 83.22419 W | NYSM0000191458 |
| PURU_GOS _02a | Punctelia rudecta | 2022-04-11 | CAM-NSF-Lichen Genomics Team; Allen, Jessica L. | USA: E-slopes of Pine Mountain, Kentenia State Forest, vicinity of Goss Park, E of Little Shepherd Trail/SR1679 at jct w/ Putney Lookout Tower Rd., between | New York State Museum | 36.9243 N 83.22419 W | NYSM0000191457 |

| | | | | | | | |
|---|---|---|---|---|---|---|---|
| | | | | Spruce Ridge Branch and Hi Lewis Branch | | | |
| PURU_GOS_01c | Punctelia rudecta | 2022-04-11 | CAM-NSF-Lichen Genomics Team; Allen, Jessica L. | USA: E-slopes of Pine Mountain, Kentenia State Forest, vicinity of Goss Park, E of Little Shepherd Trail/SR1679 at jct w/ Putney Lookout Tower Rd., between Spruce Ridge Branch and Hi Lewis Branch | New York State Museum | 36.9243 N 83.22419 W | NYSM0000191456 |
| PURU_GOS_01b | Punctelia rudecta | 2022-04-11 | CAM-NSF-Lichen Genomics Team; Allen, Jessica L. | USA: E-slopes of Pine Mountain, Kentenia State Forest, vicinity of Goss Park, E of Little Shepherd Trail/SR1679 at jct w/ Putney Lookout Tower Rd., between Spruce Ridge Branch and Hi Lewis Branch | New York State Museum | 36.9243 N 83.22419 W | NYSM0000191455 |
| PURU_GOS_01a | Punctelia rudecta | 2022-04-11 | CAM-NSF-Lichen Genomics Team; Allen, Jessica L. | USA: E-slopes of Pine Mountain, Kentenia State Forest, vicinity of Goss Park, E of Little Shepherd Trail/SR1679 at jct w/ Putney Lookout Tower Rd., between Spruce Ridge Branch and Hi Lewis Branch | New York State Museum | 36.9243 N 83.22419 W | NYSM0000191454 |
| PURU_DBG_16a | Punctelia rudecta | 2021-10-23 | CAM-NSF-Lichen Genomics Team; Allen, Jessica L. | USA: Great Smoky Mountains National Park, Cataloochee Divide, McClue Ridge, Double Gap Trail ~0.1 mi W of Double Gap | New York State Museum | 35.57191 N 83.1035 W | NYSM0000191453 |

| | | | | | | | |
|---|---|---|---|---|---|---|---|
| PURU_DBG_15a | Punctelia rudecta | 2021-10-23 | CAM-NSF-Lichen Genomics Team; Allen, Jessica L. | USA: Great Smoky Mountains National Park, Cataloochee Divide, McClue Ridge, Double Gap Trail ~0.1 mi W of Double Gap | New York State Museum | 35.57191 N 83.1035 W | NYSM0000191452 |
| PURU_DBG_14a | Punctelia rudecta | 2021-10-23 | CAM-NSF-Lichen Genomics Team; Allen, Jessica L. | USA: Great Smoky Mountains National Park, Cataloochee Divide, McClue Ridge, Double Gap Trail ~0.1 mi W of Double Gap | New York State Museum | 35.57191 N 83.1035 W | NYSM0000191451 |
| PURU_DBG_09a | Punctelia rudecta | 2021-10-23 | CAM-NSF-Lichen Genomics Team; Allen, Jessica L. | USA: Great Smoky Mountains National Park, Cataloochee Divide, McClue Ridge, Double Gap Trail ~0.1 mi W of Double Gap | New York State Museum | 35.57191 N 83.1035 W | NYSM0000191450 |
| PURU_DBG_07a | Punctelia rudecta | 2021-10-23 | CAM-NSF-Lichen Genomics Team; Allen, Jessica L. | USA: Great Smoky Mountains National Park, Cataloochee Divide, McClue Ridge, Double Gap Trail ~0.1 mi W of Double Gap | New York State Museum | 35.57191 N 83.1035 W | NYSM0000191449 |
| PURU_DBG_06a | Punctelia rudecta | 2021-10-23 | CAM-NSF-Lichen Genomics Team; Allen, Jessica L. | USA: Great Smoky Mountains National Park, Cataloochee Divide, McClue Ridge, Double Gap Trail ~0.1 mi W of Double Gap | New York State Museum | 35.57191 N 83.1035 W | NYSM0000191448 |
| PURU_DBG_05a | Punctelia rudecta | 2021-10-23 | CAM-NSF-Lichen Genomics Team; Allen, Jessica L. | USA: Great Smoky Mountains National Park, Cataloochee Divide, McClue Ridge, Double Gap Trail ~0.1 mi W of Double Gap | New York State Museum | 35.57191 N 83.1035 W | NYSM0000191447 |

| | | | | | | | |
|---|---|---|---|---|---|---|---|
| PURU_DBG_04a | Punctelia rudecta | 2021-10-23 | CAM-NSF-Lichen Genomics Team; Allen, Jessica L. | USA: Great Smoky Mountains National Park, Cataloochee Divide, McClue Ridge, Double Gap Trail ~0.1 mi W of Double Gap | New York State Museum | 35.57191 N 83.1035 W | NYSM0000191446 |
| PURU_DBG_02a | Punctelia rudecta | 2021-10-23 | CAM-NSF-Lichen Genomics Team; Allen, Jessica L. | USA: Great Smoky Mountains National Park, Cataloochee Divide, McClue Ridge, Double Gap Trail ~0.1 mi W of Double Gap | New York State Museum | 35.57191 N 83.1035 W | NYSM0000191445 |
| PURU_DBG_01a | Punctelia rudecta | 2021-10-23 | CAM-NSF-Lichen Genomics Team; Allen, Jessica L. | USA: Great Smoky Mountains National Park, Cataloochee Divide, McClue Ridge, Double Gap Trail ~0.1 mi W of Double Gap | New York State Museum | 35.57191 N 83.1035 W | NYSM0000191444 |
| PURU_COG_15a | Punctelia rudecta | 2022-08-01 | CAM-NSF-Lichen Genomics Team; Denzer, Eli | USA: Nantahala National Forest, N-facing slopes above FS4535 0.3 mi W of jct w/ SR1538/Buck Creek Rd. at Cole Gap | New York State Museum | 35.1036 N 83.2075 W | NYSM0000194994 |
| PURU_CAT_12a | Punctelia rudecta | 2021-10-27 | CAM-NSF-Lichen Genomics Team; Allen, Jessica L. | USA: Pisgah National Forest, summit of Cattail Peak, Black Mountain Crest Trail 3 mi N of Mount Mitchell, ~8 mi S of US19E | New York State Museum | 35.7964 N 82.2569 W | NYSM0000191443 |
| PURU_CAT_11a | Punctelia rudecta | 2021-10-27 | CAM-NSF-Lichen Genomics Team; Allen, Jessica L. | USA: Pisgah National Forest, summit of Cattail Peak, Black Mountain Crest Trail 3 mi N of Mount Mitchell, ~8 mi S of US19E | New York State Museum | 35.7964 N 82.2569 W | NYSM0000191442 |

| | | | | | | | |
|---|---|---|---|---|---|---|---|
| PURU_CAT _10a | Punctelia rudecta | 2021-10-27 | CAM-NSF-Lichen Genomics Team; Allen, Jessica L. | USA: Pisgah National Forest, summit of Cattail Peak, Black Mountain Crest Trail 3 mi N of Mount Mitchell, ~8 mi S of US19E | New York State Museum | 35.7964 N 82.2569 W | NYSM0000191441 |
| PURU_CAT _09a | Punctelia rudecta | 2021-10-27 | CAM-NSF-Lichen Genomics Team; Allen, Jessica L. | USA: Pisgah National Forest, summit of Cattail Peak, Black Mountain Crest Trail 3 mi N of Mount Mitchell, ~8 mi S of US19E | New York State Museum | 35.7964 N 82.2569 W | NYSM0000191440 |
| PURU_CAT _08a | Punctelia rudecta | 2021-10-27 | CAM-NSF-Lichen Genomics Team; Allen, Jessica L. | USA: Pisgah National Forest, summit of Cattail Peak, Black Mountain Crest Trail 3 mi N of Mount Mitchell, ~8 mi S of US19E | New York State Museum | 35.7964 N 82.2569 W | NYSM0000191439 |
| PURU_CAT _05a | Punctelia rudecta | 2021-10-27 | CAM-NSF-Lichen Genomics Team; Allen, Jessica L. | USA: Pisgah National Forest, summit of Cattail Peak, Black Mountain Crest Trail 3 mi N of Mount Mitchell, ~8 mi S of US19E | New York State Museum | 35.7964 N 82.2569 W | NYSM0000191438 |
| PURU_CAT _04a | Punctelia rudecta | 2021-10-27 | CAM-NSF-Lichen Genomics Team; Allen, Jessica L. | USA: Pisgah National Forest, summit of Cattail Peak, Black Mountain Crest Trail 3 mi N of Mount Mitchell, ~8 mi S of US19E | New York State Museum | 35.7964 N 82.2569 W | NYSM0000191437 |
| PURU_CAT _03a | Punctelia rudecta | 2021-10-27 | CAM-NSF-Lichen Genomics Team; Allen, Jessica L. | USA: Pisgah National Forest, summit of Cattail Peak, Black Mountain Crest Trail 3 mi N of Mount Mitchell, ~8 mi S of US19E | New York State Museum | 35.7964 N 82.2569 W | NYSM0000191436 |

| | | | | | | | |
|---|---|---|---|---|---|---|---|
| PURU_CAT _02a | Punctelia rudecta | 2021-10-27 | CAM-NSF-Lichen Genomics Team; Allen, Jessica L. | USA: Pisgah National Forest, summit of Cattail Peak, Black Mountain Crest Trail 3 mi N of Mount Mitchell, ~8 mi S of US19E | New York State Museum | 35.7964 N 82.2569 W | NYSM0000191435 |
| PURU_CAT _01a | Punctelia rudecta | 2021-10-27 | CAM-NSF-Lichen Genomics Team; Allen, Jessica L. | USA: Pisgah National Forest, summit of Cattail Peak, Black Mountain Crest Trail 3 mi N of Mount Mitchell, ~8 mi S of US19E | New York State Museum | 35.7964 N 82.2569 W | NYSM0000191434 |
| PURU_BUK _20a | Punctelia rudecta | 2021-10-24 | CAM-NSF-Lichen Genomics Team; Allen, Jessica L. | USA: Great Smoky Mountains National Park, Cataloochee Divide, N-slope of Little Bald Knob, Hemphill Bald Trail ~3 mi E of Polls Gap | New York State Museum | 35.552276 N 83.120161 W | NYSM0000191433 |
| PURU_BUK _19a | Punctelia rudecta | 2021-10-24 | CAM-NSF-Lichen Genomics Team; Allen, Jessica L. | USA: Great Smoky Mountains National Park, Cataloochee Divide, N-slope of Little Bald Knob, Hemphill Bald Trail ~3 mi E of Polls Gap | New York State Museum | 35.552276 N 83.120161 W | NYSM0000191432 |
| PURU_BUK _17a | Punctelia rudecta | 2021-10-24 | CAM-NSF-Lichen Genomics Team; Allen, Jessica L. | USA: Great Smoky Mountains National Park, Cataloochee Divide, N-slope of Little Bald Knob, Hemphill Bald Trail ~3 mi E of Polls Gap | New York State Museum | 35.552276 N 83.120161 W | NYSM0000191431 |
| PURU_BUK _16a | Punctelia rudecta | 2021-10-24 | CAM-NSF-Lichen Genomics Team; Allen, Jessica L. | USA: Great Smoky Mountains National Park, Cataloochee Divide, N-slope of Little Bald Knob, Hemphill Bald Trail ~3 mi E of Polls Gap | New York State Museum | 35.552276 N 83.120161 W | NYSM0000191430 |

| | | | | | | | |
|---|---|---|---|---|---|---|---|
| PURU_BUK_15a | Punctelia rudecta | 2021-10-24 | CAM-NSF-Lichen Genomics Team; Allen, Jessica L. | USA: Great Smoky Mountains National Park, Cataloochee Divide, N-slope of Little Bald Knob, Hemphill Bald Trail ~3 mi E of Polls Gap | New York State Museum | 35.552276 N 83.120161 W | NYSM0000191429 |
| PURU_BUK_14a | Punctelia rudecta | 2021-10-24 | CAM-NSF-Lichen Genomics Team; Allen, Jessica L. | USA: Great Smoky Mountains National Park, Cataloochee Divide, N-slope of Little Bald Knob, Hemphill Bald Trail ~3 mi E of Polls Gap | New York State Museum | 35.552276 N 83.120161 W | NYSM0000191428 |
| PURU_BUK_10a | Punctelia rudecta | 2021-10-24 | CAM-NSF-Lichen Genomics Team; Allen, Jessica L. | USA: Great Smoky Mountains National Park, Cataloochee Divide, N-slope of Little Bald Knob, Hemphill Bald Trail ~3 mi E of Polls Gap | New York State Museum | 35.552276 N 83.120161 W | NYSM0000191427 |
| PURU_BUK_05a | Punctelia rudecta | 2021-10-24 | CAM-NSF-Lichen Genomics Team; Allen, Jessica L. | USA: Great Smoky Mountains National Park, Cataloochee Divide, N-slope of Little Bald Knob, Hemphill Bald Trail ~3 mi E of Polls Gap | New York State Museum | 35.552276 N 83.120161 W | NYSM0000191426 |
| PURU_BUK_04a | Punctelia rudecta | 2021-10-24 | CAM-NSF-Lichen Genomics Team; Allen, Jessica L. | USA: Great Smoky Mountains National Park, Cataloochee Divide, N-slope of Little Bald Knob, Hemphill Bald Trail ~3 mi E of Polls Gap | New York State Museum | 35.552276 N 83.120161 W | NYSM0000191425 |
| PURU_BUK_01a | Punctelia rudecta | 2021-10-24 | CAM-NSF-Lichen Genomics Team; Allen, Jessica L. | USA: Great Smoky Mountains National Park, Cataloochee Divide, N-slope of Little Bald Knob, Hemphill Bald Trail ~3 mi E of Polls Gap | New York State Museum | 35.552276 N 83.120161 W | NYSM0000191424 |

| | | | | | | | |
|---|---|---|---|---|---|---|---|
| PURU_BRF_04c | Punctelia rudecta | 2022-05-03 | CAM-NSF-Lichen Genomics Team; Allen, Jessica L. | USA: Shenandoah National Park, Central District, Bearfence Mountain, W-slopes below Appalachian Trail, 0.3 mi S of jct w/ spur trail to Skyline Drive/Bearfence Parking Area | New York State Museum | 38.4477 N 78.46635 W | NYSM0000191412 |
| PURU_BRF_04b | Punctelia rudecta | 2022-05-03 | CAM-NSF-Lichen Genomics Team; Allen, Jessica L. | USA: Shenandoah National Park, Central District, Bearfence Mountain, W-slopes below Appalachian Trail, 0.3 mi S of jct w/ spur trail to Skyline Drive/Bearfence Parking Area | New York State Museum | 38.4477 N 78.46635 W | NYSM0000191411 |
| PURU_BRF_04a | Punctelia rudecta | 2022-05-03 | CAM-NSF-Lichen Genomics Team; Allen, Jessica L. | USA: Shenandoah National Park, Central District, Bearfence Mountain, W-slopes below Appalachian Trail, 0.3 mi S of jct w/ spur trail to Skyline Drive/Bearfence Parking Area | New York State Museum | 38.4477 N 78.46635 W | NYSM0000191410 |
| PURU_BRF_03c | Punctelia rudecta | 2022-05-03 | CAM-NSF-Lichen Genomics Team; Allen, Jessica L. | USA: Shenandoah National Park, Central District, Bearfence Mountain, W-slopes below Appalachian Trail, 0.3 mi S of jct w/ spur trail to Skyline Drive/Bearfence Parking Area | New York State Museum | 38.4477 N 78.46635 W | NYSM0000191409 |
| PURU_BRF_03b | Punctelia rudecta | 2022-05-03 | CAM-NSF-Lichen Genomics Team; Allen, Jessica L. | USA: Shenandoah National Park, Central District, Bearfence Mountain, W-slopes below Appalachian Trail, 0.3 mi S of jct w/ spur trail to Skyline Drive/Bearfence Parking Area | New York State Museum | 38.4477 N 78.46635 W | NYSM0000191408 |

| | | | | | | | |
|---|---|---|---|---|---|---|---|
| PURU_BRF _03a | Punctelia rudecta | 2022-05-03 | CAM-NSF-Lichen Genomics Team; Allen, Jessica L. | USA: Shenandoah National Park, Central District, Bearfence Mountain, W-slopes below Appalachian Trail, 0.3 mi S of jct w/ spur trail to Skyline Drive/Bearfence Parking Area | New York State Museum | 38.4477 N 78.46635 W | NYSM0000191407 |
| PURU_BRF _02c | Punctelia rudecta | 2022-05-03 | CAM-NSF-Lichen Genomics Team; Allen, Jessica L. | USA: Shenandoah National Park, Central District, Bearfence Mountain, W-slopes below Appalachian Trail, 0.3 mi S of jct w/ spur trail to Skyline Drive/Bearfence Parking Area | New York State Museum | 38.4477 N 78.46635 W | NYSM0000191406 |
| PURU_BRF _02b | Punctelia rudecta | 2022-05-03 | CAM-NSF-Lichen Genomics Team; Allen, Jessica L. | USA: Shenandoah National Park, Central District, Bearfence Mountain, W-slopes below Appalachian Trail, 0.3 mi S of jct w/ spur trail to Skyline Drive/Bearfence Parking Area | New York State Museum | 38.4477 N 78.46635 W | NYSM0000191405 |
| PURU_BRF _02a | Punctelia rudecta | 2022-05-03 | CAM-NSF-Lichen Genomics Team; Allen, Jessica L. | USA: Shenandoah National Park, Central District, Bearfence Mountain, W-slopes below Appalachian Trail, 0.3 mi S of jct w/ spur trail to Skyline Drive/Bearfence Parking Area | New York State Museum | 38.4477 N 78.46635 W | NYSM0000191404 |
| PURU_BRF _01c | Punctelia rudecta | 2022-05-03 | CAM-NSF-Lichen Genomics Team; Allen, Jessica L. | USA: Shenandoah National Park, Central District, Bearfence Mountain, W-slopes below Appalachian Trail, 0.3 mi S of jct w/ spur trail to Skyline Drive/Bearfence Parking Area | New York State Museum | 38.4477 N 78.46635 W | NYSM0000191403 |

| | | | | | | | |
|---|---|---|---|---|---|---|---|
| PURU_BRF _01b | Punctelia rudecta | 2022-05-03 | CAM-NSF-Lichen Genomics Team; Allen, Jessica L. | USA: Shenandoah National Park, Central District, Bearfence Mountain, W-slopes below Appalachian Trail, 0.3 mi S of jct w/ spur trail to Skyline Drive/Bearfence Parking Area | New York State Museum | 38.4477 N 78.46635 W | NYSM0000191402 |
| PURU_BRF _01a | Punctelia rudecta | 2022-05-03 | CAM-NSF-Lichen Genomics Team; Allen, Jessica L. | USA: Shenandoah National Park, Central District, Bearfence Mountain, W-slopes below Appalachian Trail, 0.3 mi S of jct w/ spur trail to Skyline Drive/Bearfence Parking Area | New York State Museum | 38.4477 N 78.46635 W | NYSM0000191401 |
| PURU_BLU _06b | Punctelia rudecta | 2022-04-13 | CAM-NSF-Lichen Genomics Team; Allen, Jessica L. | USA: Pine Mountain, Bad Branch Nature Preserve, headwaters of Presley House Branch, Pine Mountain Trail ~1.5 mi E of US119 | New York State Museum | 37.08482 N 82.79308 W | NYSM0000191423 |
| PURU_BLU _06a | Punctelia rudecta | 2022-04-13 | CAM-NSF-Lichen Genomics Team; Allen, Jessica L. | USA: Pine Mountain, Bad Branch Nature Preserve, headwaters of Presley House Branch, Pine Mountain Trail ~1.5 mi E of US119 | New York State Museum | 37.08482 N 82.79308 W | NYSM0000191422 |
| PURU_BLU _03c | Punctelia rudecta | 2022-04-13 | CAM-NSF-Lichen Genomics Team; Allen, Jessica L. | USA: Pine Mountain, Bad Branch Nature Preserve, headwaters of Presley House Branch, Pine Mountain Trail ~1.5 mi E of US119 | New York State Museum | 37.08482 N 82.79308 W | NYSM0000191421 |

| | | | | | | | |
|---|---|---|---|---|---|---|---|
| PURU_BLU _03b | Punctelia rudecta | 2022-04-13 | CAM-NSF-Lichen Genomics Team; Allen, Jessica L. | USA: Pine Mountain, Bad Branch Nature Preserve, headwaters of Presley House Branch, Pine Mountain Trail ~1.5 mi E of US119 | New York State Museum | 37.08482 N 82.79308 W | NYSM0000191420 |
| PURU_BLU _03a | Punctelia rudecta | 2022-04-13 | CAM-NSF-Lichen Genomics Team; Allen, Jessica L. | USA: Pine Mountain, Bad Branch Nature Preserve, headwaters of Presley House Branch, Pine Mountain Trail ~1.5 mi E of US119 | New York State Museum | 37.08482 N 82.79308 W | NYSM0000191419 |
| PURU_BLU _02c | Punctelia rudecta | 2022-04-13 | CAM-NSF-Lichen Genomics Team; Allen, Jessica L. | USA: Pine Mountain, Bad Branch Nature Preserve, headwaters of Presley House Branch, Pine Mountain Trail ~1.5 mi E of US119 | New York State Museum | 37.08482 N 82.79308 W | NYSM0000191418 |
| PURU_BLU _02b | Punctelia rudecta | 2022-04-13 | CAM-NSF-Lichen Genomics Team; Allen, Jessica L. | USA: Pine Mountain, Bad Branch Nature Preserve, headwaters of Presley House Branch, Pine Mountain Trail ~1.5 mi E of US119 | New York State Museum | 37.08482 N 82.79308 W | NYSM0000191417 |
| PURU_BLU _02a | Punctelia rudecta | 2022-04-13 | CAM-NSF-Lichen Genomics Team; Allen, Jessica L. | USA: Pine Mountain, Bad Branch Nature Preserve, headwaters of Presley House Branch, Pine Mountain Trail ~1.5 mi E of US119 | New York State Museum | 37.08482 N 82.79308 W | NYSM0000191416 |
| PURU_BLU _01c | Punctelia rudecta | 2022-04-13 | CAM-NSF-Lichen Genomics Team; Allen, Jessica L. | USA: Pine Mountain, Bad Branch Nature Preserve, headwaters of Presley House Branch, Pine Mountain Trail ~1.5 mi E of US119 | New York State Museum | 37.08482 N 82.79308 W | NYSM0000191415 |

| | | | | | | | |
|---|---|---|---|---|---|---|---|
| PURU_BLU _01b | Punctelia rudecta | 2022-04-13 | CAM-NSF-Lichen Genomics Team; Allen, Jessica L. | USA: Pine Mountain, Bad Branch Nature Preserve, headwaters of Presley House Branch, Pine Mountain Trail ~1.5 mi E of US119 | New York State Museum | 37.08482 N 82.79308 W | NYSM0000191414 |
| PURU_BLU _01a | Punctelia rudecta | 2022-04-13 | CAM-NSF-Lichen Genomics Team; Allen, Jessica L. | USA: Pine Mountain, Bad Branch Nature Preserve, headwaters of Presley House Branch, Pine Mountain Trail ~1.5 mi E of US119 | New York State Museum | 37.08482 N 82.79308 W | NYSM0000191413 |
| PURU_BFF _17a | Punctelia rudecta | 2021-10-26 | CAM-NSF-Lichen Genomics Team; Allen, Jessica L. | USA: Pisgah National Forest, Bald Mountains, N-Facing slopes of Bluff Mountain summit, NE of Appalachian Trail 1 mi E of Catpen Gap, 3.4 mi E of Lemon Gap | New York State Museum | 35.84115 N 82.90576 W | NYSM0000191400 |
| PURU_BFF _16a | Punctelia rudecta | 2021-10-26 | CAM-NSF-Lichen Genomics Team; Allen, Jessica L. | USA: Pisgah National Forest, Bald Mountains, N-Facing slopes of Bluff Mountain summit, NE of Appalachian Trail 1 mi E of Catpen Gap, 3.4 mi E of Lemon Gap | New York State Museum | 35.84115 N 82.90576 W | NYSM0000191399 |
| PURU_BFF _13a | Punctelia rudecta | 2021-10-26 | CAM-NSF-Lichen Genomics Team; Allen, Jessica L. | USA: Pisgah National Forest, Bald Mountains, N-Facing slopes of Bluff Mountain summit, NE of Appalachian Trail 1 mi E of Catpen Gap, 3.4 mi E of Lemon Gap | New York State Museum | 35.84115 N 82.90576 W | NYSM0000191398 |

| | | | | | | | |
|---|---|---|---|---|---|---|---|
| PURU_BFF_11a | Punctelia rudecta | 2021-10-26 | CAM-NSF-Lichen Genomics Team; Allen, Jessica L. | USA: Pisgah National Forest, Bald Mountains, N-Facing slopes of Bluff Mountain summit, NE of Appalachian Trail 1 mi E of Catpen Gap, 3.4 mi E of Lemon Gap | New York State Museum | 35.84115 N 82.90576 W | NYSM0000191397 |
| PURU_BFF_10a | Punctelia rudecta | 2021-10-26 | CAM-NSF-Lichen Genomics Team; Allen, Jessica L. | USA: Pisgah National Forest, Bald Mountains, N-Facing slopes of Bluff Mountain summit, NE of Appalachian Trail 1 mi E of Catpen Gap, 3.4 mi E of Lemon Gap | New York State Museum | 35.84115 N 82.90576 W | NYSM0000191396 |
| PURU_BFF_09a | Punctelia rudecta | 2021-10-26 | CAM-NSF-Lichen Genomics Team; Allen, Jessica L. | USA: Pisgah National Forest, Bald Mountains, N-Facing slopes of Bluff Mountain summit, NE of Appalachian Trail 1 mi E of Catpen Gap, 3.4 mi E of Lemon Gap | New York State Museum | 35.84115 N 82.90576 W | NYSM0000191395 |
| PURU_BFF_08a | Punctelia rudecta | 2021-10-26 | CAM-NSF-Lichen Genomics Team; Allen, Jessica L. | USA: Pisgah National Forest, Bald Mountains, N-Facing slopes of Bluff Mountain summit, NE of Appalachian Trail 1 mi E of Catpen Gap, 3.4 mi E of Lemon Gap | New York State Museum | 35.84115 N 82.90576 W | NYSM0000191394 |
| PURU_BFF_03a | Punctelia rudecta | 2021-10-26 | CAM-NSF-Lichen Genomics Team; Allen, Jessica L. | USA: Pisgah National Forest, Bald Mountains, N-Facing slopes of Bluff Mountain summit, NE of Appalachian Trail 1 mi E of Catpen Gap, 3.4 mi E of Lemon Gap | New York State Museum | 35.84115 N 82.90576 W | NYSM0000191393 |

| | | | | | | | |
|---|---|---|---|---|---|---|---|
| PURU_BFF_02a | Punctelia rudecta | 2021-10-26 | CAM-NSF-Lichen Genomics Team; Allen, Jessica L. | USA: Pisgah National Forest, Bald Mountains, N-Facing slopes of Bluff Mountain summit, NE of Appalachian Trail 1 mi E of Catpen Gap, 3.4 mi E of Lemon Gap | New York State Museum | 35.84115 N 82.90576 W | NYSM0000191392 |
| PURU_BFF_01a | Punctelia rudecta | 2021-10-26 | CAM-NSF-Lichen Genomics Team; Allen, Jessica L. | USA: Pisgah National Forest, Bald Mountains, N-Facing slopes of Bluff Mountain summit, NE of Appalachian Trail 1 mi E of Catpen Gap, 3.4 mi E of Lemon Gap | New York State Museum | 35.84115 N 82.90576 W | NYSM0000191391 |
| PURU_BEK_11a | Punctelia rudecta | 2021-10-26 | CAM-NSF-Lichen Genomics Team; Allen, Jessica L. | USA: Pisgah National Forest, Bald Mountains, N-slope of Roaring Fork Mountain, spur of Buckeye Ridge | New York State Museum | 35.8014 N 82.95096 W | NYSM0000191390 |
| PURU_BEK_10a | Punctelia rudecta | 2021-10-26 | CAM-NSF-Lichen Genomics Team; Allen, Jessica L. | USA: Pisgah National Forest, Bald Mountains, N-slope of Roaring Fork Mountain, spur of Buckeye Ridge | New York State Museum | 35.8014 N 82.95096 W | NYSM0000191389 |
| PURU_BEK_08a | Punctelia rudecta | 2021-10-26 | CAM-NSF-Lichen Genomics Team; Allen, Jessica L. | USA: Pisgah National Forest, Bald Mountains, N-slope of Roaring Fork Mountain, spur of Buckeye Ridge | New York State Museum | 35.8014 N 82.95096 W | NYSM0000191388 |

| | | | | | | | |
|---|---|---|---|---|---|---|---|
| PURU_BEK _07a | Punctelia rudecta | 2021-10-26 | CAM-NSF-Lichen Genomics Team; Allen, Jessica L. | USA: Pisgah National Forest, Bald Mountains, N-slope of Roaring Fork Mountain, spur of Buckeye Ridge | New York State Museum | 35.8014 N 82.95096 W | NYSM0000191387 |
| PURU_BEK _06a | Punctelia rudecta | 2021-10-26 | CAM-NSF-Lichen Genomics Team; Allen, Jessica L. | USA: Pisgah National Forest, Bald Mountains, N-slope of Roaring Fork Mountain, spur of Buckeye Ridge | New York State Museum | 35.8014 N 82.95096 W | NYSM0000191386 |
| PURU_BEK _04a | Punctelia rudecta | 2021-10-26 | CAM-NSF-Lichen Genomics Team; Allen, Jessica L. | USA: Pisgah National Forest, Bald Mountains, N-slope of Roaring Fork Mountain, spur of Buckeye Ridge | New York State Museum | 35.8014 N 82.95096 W | NYSM0000191384 |
| PURU_BEK _03b | Punctelia rudecta | 2021-10-26 | CAM-NSF-Lichen Genomics Team; Allen, Jessica L. | USA: Pisgah National Forest, Bald Mountains, N-slope of Roaring Fork Mountain, spur of Buckeye Ridge | New York State Museum | 35.8014 N 82.95096 W | NYSM0000191385 |
| PURU_BEK _03a | Punctelia rudecta | 2021-10-26 | CAM-NSF-Lichen Genomics Team; Allen, Jessica L. | USA: Pisgah National Forest, Bald Mountains, N-slope of Roaring Fork Mountain, spur of Buckeye Ridge | New York State Museum | 35.8014 N 82.95096 W | NYSM0000191383 |
| PURU_BEK _02a | Punctelia rudecta | 2021-10-26 | CAM-NSF-Lichen Genomics Team; Allen, Jessica L. | USA: Pisgah National Forest, Bald Mountains, N-slope of Roaring Fork Mountain, spur of Buckeye Ridge | New York State Museum | 35.8014 N 82.95096 W | NYSM0000191382 |

| | | | | | | | |
|---|---|---|---|---|---|---|---|
| PURU_BEK _01a | Punctelia rudecta | 2021-10-26 | CAM-NSF-Lichen Genomics Team; Allen, Jessica L. | USA: Pisgah National Forest, Bald Mountains, N-slope of Roaring Fork Mountain, spur of Buckeye Ridge | New York State Museum | 35.8014 N 82.95096 W | NYSM0000191381 |
| LELA_TOM _2a | Lepraria lanata | 2021-10-27 | CAM-NSF-Lichen Genomics Team | USA: Mount Mitchell State Park, vicinity of summit of Balsam Cone, Black Mountain Crest Trail ~2 mi N of Mount Mitchell and terminus of NC128 | New York State Museum | 35.7892 N 82.2558 W | NYSM0000194833 |
| LELA_TOM _19c | Lepraria lanata | 2021-10-27 | CAM-NSF-Lichen Genomics Team | USA: Mount Mitchell State Park, vicinity of summit of Balsam Cone, Black Mountain Crest Trail ~2 mi N of Mount Mitchell and terminus of NC128 | New York State Museum | 35.7892 N 82.2558 W | NYSM0000194823 |
| LELA_TOM _19b | Lepraria lanata | 2021-10-27 | CAM-NSF-Lichen Genomics Team | USA: Mount Mitchell State Park, vicinity of summit of Balsam Cone, Black Mountain Crest Trail ~2 mi N of Mount Mitchell and terminus of NC128 | New York State Museum | 35.7892 N 82.2558 W | NYSM0000194824 |
| LELA_TOM _19a | Lepraria lanata | 2021-10-29 | CAM-NSF-Lichen Genomics Team | USA: Mount Mitchell State Park, vicinity of summit of Balsam Cone, Black Mountain Crest Trail ~2 mi N of Mount Mitchell and terminus of NC128 | New York State Museum | 35.7892 N 82.2558 W | NYSM0000194825 |
| LELA_TOM _17b | Lepraria lanata | 2021-10-28 | CAM-NSF-Lichen Genomics Team | USA: Mount Mitchell State Park, vicinity of summit of Balsam Cone, Black Mountain Crest Trail ~2 mi N of Mount Mitchell and terminus of NC128 | New York State Museum | 35.7892 N 82.2558 W | NYSM0000194826 |

| | | | | | | | |
|---|---|---|---|---|---|---|---|
| LELA_TOM _17a | Lepraria lanata | 2021-10-27 | CAM-NSF-Lichen Genomics Team | USA: Mount Mitchell State Park, vicinity of summit of Balsam Cone, Black Mountain Crest Trail ~2 mi N of Mount Mitchell and terminus of NC128 | New York State Museum | 35.7892 N 82.2558 W | NYSM0000194827 |
| LELA_TOM _16a | Lepraria lanata | 2021-10-27 | CAM-NSF-Lichen Genomics Team | USA: Mount Mitchell State Park, vicinity of summit of Balsam Cone, Black Mountain Crest Trail ~2 mi N of Mount Mitchell and terminus of NC128 | New York State Museum | 35.7892 N 82.2558 W | NYSM0000194828 |
| LELA_TOM _14b | Lepraria lanata | 2021-10-27 | CAM-NSF-Lichen Genomics Team | USA: Mount Mitchell State Park, vicinity of summit of Balsam Cone, Black Mountain Crest Trail ~2 mi N of Mount Mitchell and terminus of NC128 | New York State Museum | 35.7892 N 82.2558 W | NYSM0000194829 |
| LELA_TOM _14a | Lepraria lanata | 2021-10-27 | CAM-NSF-Lichen Genomics Team | USA: Mount Mitchell State Park, vicinity of summit of Balsam Cone, Black Mountain Crest Trail ~2 mi N of Mount Mitchell and terminus of NC128 | New York State Museum | 35.7892 N 82.2558 W | NYSM0000194830 |
| LELA_TOM _13b | Lepraria lanata | 2021-10-27 | CAM-NSF-Lichen Genomics Team | USA: Mount Mitchell State Park, vicinity of summit of Balsam Cone, Black Mountain Crest Trail ~2 mi N of Mount Mitchell and terminus of NC128 | New York State Museum | 35.7892 N 82.2558 W | NYSM0000194831 |
| LELA_TOM _13a | Lepraria lanata | 2021-10-27 | CAM-NSF-Lichen Genomics Team | USA: Mount Mitchell State Park, vicinity of summit of Balsam Cone, Black Mountain Crest Trail ~2 mi N of Mount Mitchell and terminus of NC128 | New York State Museum | 35.7892 N 82.2558 W | NYSM0000194832 |

| | | | | | | | |
|---|---|---|---|---|---|---|---|
| LELA_SPF_01c | Lepraria lanata | 2022-08-06 | CAM-NSF-Lichen Genomics Team | USA: Pisgah National Forest, Roan Mountain, S slope of Roan High Bluff, ~0.2 mi E of overlook & ~0.3 mi W of parking area | New York State Museum | 36.09362 N 82.14485 W | NYSM0000191907 |
| LELA_SPF_01b | Lepraria lanata | 2022-08-06 | CAM-NSF-Lichen Genomics Team | USA: Pisgah National Forest, Roan Mountain, S slope of Roan High Bluff, ~0.2 mi E of overlook & ~0.3 mi W of parking area | New York State Museum | 36.09362 N 82.14485 W | NYSM0000191906 |
| LELA_SPF_01a | Lepraria lanata | 2022-08-06 | CAM-NSF-Lichen Genomics Team | USA: Pisgah National Forest, Roan Mountain, S slope of Roan High Bluff, ~0.2 mi E of overlook & ~0.3 mi W of parking area | New York State Museum | 36.09362 N 82.14485 W | NYSM0000191905 |
| LELA_RBB_02f | Lepraria lanata | 2022-08-05 | CAM-NSF-Lichen Genomics Team | USA: Chattahoochee National Forest, Rabun Bald, jct of Bartram Trail ~1.7 mi S of Beegum Gap | New York State Museum | 34.96532 N 83.2999 W | NYSM0000194834 |
| LELA_RBB_02e | Lepraria lanata | 2022-08-05 | CAM-NSF-Lichen Genomics Team | USA: Chattahoochee National Forest, Rabun Bald, jct of Bartram Trail ~1.7 mi S of Beegum Gap | New York State Museum | 34.96532 N 83.2999 W | NYSM0000194835 |
| LELA_RBB_02d | Lepraria lanata | 2022-08-05 | CAM-NSF-Lichen Genomics Team | USA: Chattahoochee National Forest, Rabun Bald, jct of Bartram Trail ~1.7 mi S of Beegum Gap | New York State Museum | 34.96532 N 83.2999 W | NYSM0000194836 |

| | | | | | | | |
|---|---|---|---|---|---|---|---|
| LELA_RBB_02c | Lepraria lanata | 2022-08-05 | CAM-NSF-Lichen Genomics Team | USA: Chattahoochee National Forest, Rabun Bald, jct of Bartram Trail ~1.7 mi S of Beegum Gap | New York State Museum | 34.96532 N 83.2999 W | NYSM0000194837 |
| LELA_RBB_02b | Lepraria lanata | 2022-08-05 | CAM-NSF-Lichen Genomics Team | USA: Chattahoochee National Forest, Rabun Bald, jct of Bartram Trail ~1.7 mi S of Beegum Gap | New York State Museum | 34.96532 N 83.2999 W | NYSM0000194838 |
| LELA_RBB_02a | Lepraria lanata | 2022-08-05 | CAM-NSF-Lichen Genomics Team | USA: Chattahoochee National Forest, Rabun Bald, jct of Bartram Trail ~1.7 mi S of Beegum Gap | New York State Museum | 34.96532 N 83.2999 W | NYSM0000194839 |
| LELA_RBB_01c | Lepraria lanata | 2022-08-05 | CAM-NSF-Lichen Genomics Team | USA: Chattahoochee National Forest, Rabun Bald, jct of Bartram Trail ~1.7 mi S of Beegum Gap | New York State Museum | 34.96532 N 83.2999 W | NYSM0000194840 |
| LELA_RBB_01b | Lepraria lanata | 2022-08-05 | CAM-NSF-Lichen Genomics Team | USA: Chattahoochee National Forest, Rabun Bald, jct of Bartram Trail ~1.7 mi S of Beegum Gap | New York State Museum | 34.96532 N 83.2999 W | NYSM0000194841 |
| LELA_RBB_01a | Lepraria lanata | 2022-08-05 | CAM-NSF-Lichen Genomics Team | USA: Chattahoochee National Forest, Rabun Bald, jct of Bartram Trail ~1.7 mi S of Beegum Gap | New York State Museum | 34.96532 N 83.2999 W | NYSM0000194842 |

| | | | | | | | |
|---|---|---|---|---|---|---|---|
| LELA_NAL _03c | Lepraria lanata | 2022-08-02 | CAM-NSF-Lichen Genomics Team | USA: Great Smoky Mountains National Park, Mount LeConte, N-facing slopes above Trillium Gap Trail, 0.4 mi N of jct w/ Rainbow Falls Trail/Boulevard Trail at LeConte Lodge | New York State Museum | 35.65878 N 83.43787 W | |
| LELA_NAL _03b | Lepraria lanata | 2022-08-02 | CAM-NSF-Lichen Genomics Team | USA: Great Smoky Mountains National Park, Mount LeConte, N-facing slopes above Trillium Gap Trail, 0.4 mi N of jct w/ Rainbow Falls Trail/Boulevard Trail at LeConte Lodge | New York State Museum | 35.65878 N 83.43787 W | NYSM0000194843 |
| LELA_NAL _03a | Lepraria lanata | 2022-08-02 | CAM-NSF-Lichen Genomics Team | USA: Great Smoky Mountains National Park, Mount LeConte, N-facing slopes above Trillium Gap Trail, 0.4 mi N of jct w/ Rainbow Falls Trail/Boulevard Trail at LeConte Lodge | New York State Museum | 35.65878 N 83.43787 W | NYSM0000194844 |
| LELA_NAL _02c | Lepraria lanata | 2022-08-02 | CAM-NSF-Lichen Genomics Team | USA: Great Smoky Mountains National Park, Mount LeConte, N-facing slopes above Trillium Gap Trail, 0.4 mi N of jct w/ Rainbow Falls Trail/Boulevard Trail at LeConte Lodge | New York State Museum | 35.65878 N 83.43787 W | NYSM0000194845 |
| LELA_NAL _02b | Lepraria lanata | 2022-08-02 | CAM-NSF-Lichen Genomics Team | USA: Great Smoky Mountains National Park, Mount LeConte, N-facing slopes above Trillium Gap Trail, 0.4 mi N of jct w/ Rainbow Falls Trail/Boulevard Trail at LeConte Lodge | New York State Museum | 35.65878 N 83.43787 W | NYSM0000194846 |

| | | | | | | | |
|---|---|---|---|---|---|---|---|
| LELA_NAL_02a | Lepraria lanata | 2022-08-02 | CAM-NSF-Lichen Genomics Team | USA: Great Smoky Mountains National Park, Mount LeConte, N-facing slopes above Trillium Gap Trail, 0.4 mi N of jct w/ Rainbow Falls Trail/Boulevard Trail at LeConte Lodge | New York State Museum | 35.65878 N 83.43787 W | NYSM0000194847 |
| LELA_NAL_01c | Lepraria lanata | 2022-08-02 | CAM-NSF-Lichen Genomics Team | USA: Great Smoky Mountains National Park, Mount LeConte, N-facing slopes above Trillium Gap Trail, 0.4 mi N of jct w/ Rainbow Falls Trail/Boulevard Trail at LeConte Lodge | New York State Museum | 35.65878 N 83.43787 W | NYSM0000194848 |
| LELA_NAL_01b | Lepraria lanata | 2022-08-02 | CAM-NSF-Lichen Genomics Team | USA: Great Smoky Mountains National Park, Mount LeConte, N-facing slopes above Trillium Gap Trail, 0.4 mi N of jct w/ Rainbow Falls Trail/Boulevard Trail at LeConte Lodge | New York State Museum | 35.65878 N 83.43787 W | NYSM0000194849 |
| LELA_NAL_01a | Lepraria lanata | 2022-08-02 | CAM-NSF-Lichen Genomics Team | USA: Great Smoky Mountains National Park, Mount LeConte, N-facing slopes above Trillium Gap Trail, 0.4 mi N of jct w/ Rainbow Falls Trail/Boulevard Trail at LeConte Lodge | New York State Museum | 35.65878 N 83.43787 W | NYSM0000194850 |
| LELA_LCN_09c | Lepraria lanata | 2022-08-02 | CAM-NSF-Lichen Genomics Team | USA: Great Smoky Mountains National Park, Mount LeConte, N-facing slopes above Trillium Gap Trail, 0.4 mi N of jct w/ Rainbow Falls Trail/Boulevard Trail at LeConte Lodge | New York State Museum | 35.65878 N 83.43787 W | NYSM0000194851 |

| | | | | | | | |
|---|---|---|---|---|---|---|---|
| LELA_LCN_09b | Lepraria lanata | 2022-08-02 | CAM-NSF-Lichen Genomics Team | USA: Great Smoky Mountains National Park, Mount LeConte, N-facing slopes above Trillium Gap Trail, 0.4 mi N of jct w/ Rainbow Falls Trail/Boulevard Trail at LeConte Lodge | New York State Museum | 35.65878 N 83.43787 W | NYSM0000194852 |
| LELA_LCN_09a | Lepraria lanata | 2022-08-02 | CAM-NSF-Lichen Genomics Team | USA: Great Smoky Mountains National Park, Mount LeConte, N-facing slopes above Trillium Gap Trail, 0.4 mi N of jct w/ Rainbow Falls Trail/Boulevard Trail at LeConte Lodge | New York State Museum | 35.65878 N 83.43787 W | NYSM0000194853 |
| LELA_LCN_08c | Lepraria lanata | 2022-08-02 | CAM-NSF-Lichen Genomics Team | USA: Great Smoky Mountains National Park, Mount LeConte, N-facing slopes above Trillium Gap Trail, 0.4 mi N of jct w/ Rainbow Falls Trail/Boulevard Trail at LeConte Lodge | New York State Museum | 35.65878 N 83.43787 W | NYSM0000194854 |
| LELA_LCN_08b | Lepraria lanata | 2022-08-02 | CAM-NSF-Lichen Genomics Team | USA: Great Smoky Mountains National Park, Mount LeConte, N-facing slopes above Trillium Gap Trail, 0.4 mi N of jct w/ Rainbow Falls Trail/Boulevard Trail at LeConte Lodge | New York State Museum | 35.65878 N 83.43787 W | NYSM0000194855 |
| LELA_LCN_08a | Lepraria lanata | 2022-08-02 | CAM-NSF-Lichen Genomics Team | USA: Great Smoky Mountains National Park, Mount LeConte, N-facing slopes above Trillium Gap Trail, 0.4 mi N of jct w/ Rainbow Falls Trail/Boulevard Trail at LeConte Lodge | New York State Museum | 35.65878 N 83.43787 W | NYSM0000194856 |

| | | | | | | | |
|---|---|---|---|---|---|---|---|
| LELA_LCN _07c | Lepraria lanata | 2022-08-02 | CAM-NSF-Lichen Genomics Team | USA: Great Smoky Mountains National Park, Mount LeConte, N-facing slopes above Trillium Gap Trail, 0.4 mi N of jct w/ Rainbow Falls Trail/Boulevard Trail at LeConte Lodge | New York State Museum | 35.65878 N 83.43787 W | NYSM0000194857 |
| LELA_LCN _07b | Lepraria lanata | 2022-08-02 | CAM-NSF-Lichen Genomics Team | USA: Great Smoky Mountains National Park, Mount LeConte, N-facing slopes above Trillium Gap Trail, 0.4 mi N of jct w/ Rainbow Falls Trail/Boulevard Trail at LeConte Lodge | New York State Museum | 35.65878 N 83.43787 W | NYSM0000194858 |
| LELA_LCN _07a | Lepraria lanata | 2022-08-02 | CAM-NSF-Lichen Genomics Team | USA: Great Smoky Mountains National Park, Mount LeConte, N-facing slopes above Trillium Gap Trail, 0.4 mi N of jct w/ Rainbow Falls Trail/Boulevard Trail at LeConte Lodge | New York State Museum | 35.65878 N 83.43787 W | NYSM0000194859 |
| LELA_LCN _06c | Lepraria lanata | 2022-08-02 | CAM-NSF-Lichen Genomics Team | USA: Great Smoky Mountains National Park, Mount LeConte, N-facing slopes above Trillium Gap Trail, 0.4 mi N of jct w/ Rainbow Falls Trail/Boulevard Trail at LeConte Lodge | New York State Museum | 35.65878 N 83.43787 W | NYSM0000194860 |
| LELA_LCN _06b | Lepraria lanata | 2022-08-02 | CAM-NSF-Lichen Genomics Team | USA: Great Smoky Mountains National Park, Mount LeConte, N-facing slopes above Trillium Gap Trail, 0.4 mi N of jct w/ Rainbow Falls Trail/Boulevard Trail at LeConte Lodge | New York State Museum | 35.65878 N 83.43787 W | NYSM0000194861 |

| | | | | | | | |
|---|---|---|---|---|---|---|---|
| LELA_LCN_06a | Lepraria lanata | 2022-08-02 | CAM-NSF-Lichen Genomics Team | USA: Great Smoky Mountains National Park, Mount LeConte, N-facing slopes above Trillium Gap Trail, 0.4 mi N of jct w/ Rainbow Falls Trail/Boulevard Trail at LeConte Lodge | New York State Museum | 35.65878 N 83.43787 W | NYSM0000194862 |
| LELA_LCN_05c | Lepraria lanata | 2022-08-02 | CAM-NSF-Lichen Genomics Team | USA: Great Smoky Mountains National Park, Mount LeConte, N-facing slopes above Trillium Gap Trail, 0.4 mi N of jct w/ Rainbow Falls Trail/Boulevard Trail at LeConte Lodge | New York State Museum | 35.65878 N 83.43787 W | NYSM0000194863 |
| LELA_LCN_05b | Lepraria lanata | 2022-08-02 | CAM-NSF-Lichen Genomics Team | USA: Great Smoky Mountains National Park, Mount LeConte, N-facing slopes above Trillium Gap Trail, 0.4 mi N of jct w/ Rainbow Falls Trail/Boulevard Trail at LeConte Lodge | New York State Museum | 35.65878 N 83.43787 W | NYSM0000194864 |
| LELA_LCN_05a | Lepraria lanata | 2022-08-02 | CAM-NSF-Lichen Genomics Team | USA: Great Smoky Mountains National Park, Mount LeConte, N-facing slopes above Trillium Gap Trail, 0.4 mi N of jct w/ Rainbow Falls Trail/Boulevard Trail at LeConte Lodge | New York State Museum | 35.65878 N 83.43787 W | NYSM0000194865 |
| LELA_LCN_04c | Lepraria lanata | 2022-08-02 | CAM-NSF-Lichen Genomics Team | USA: Great Smoky Mountains National Park, Mount LeConte, N-facing slopes above Trillium Gap Trail, 0.4 mi N of jct w/ Rainbow Falls Trail/Boulevard Trail at LeConte Lodge | New York State Museum | 35.65878 N 83.43787 W | NYSM0000194866 |

| | | | | | | | |
|---|---|---|---|---|---|---|---|
| LELA_LCN _04b | Lepraria lanata | 2022-08-02 | CAM-NSF-Lichen Genomics Team | USA: Great Smoky Mountains National Park, Mount LeConte, N-facing slopes above Trillium Gap Trail, 0.4 mi N of jct w/ Rainbow Falls Trail/Boulevard Trail at LeConte Lodge | New York State Museum | 35.65878 N 83.43787 W | NYSM0000194867 |
| LELA_LCN _04a | Lepraria lanata | 2022-08-02 | CAM-NSF-Lichen Genomics Team | USA: Great Smoky Mountains National Park, Mount LeConte, N-facing slopes above Trillium Gap Trail, 0.4 mi N of jct w/ Rainbow Falls Trail/Boulevard Trail at LeConte Lodge | New York State Museum | 35.65878 N 83.43787 W | NYSM0000194868 |
| LELA_LCN _03c | Lepraria lanata | 2022-08-02 | CAM-NSF-Lichen Genomics Team | USA: Great Smoky Mountains National Park, Mount LeConte, N-facing slopes above Trillium Gap Trail, 0.4 mi N of jct w/ Rainbow Falls Trail/Boulevard Trail at LeConte Lodge | New York State Museum | 35.65878 N 83.43787 W | NYSM0000194869 |
| LELA_LCN _03b | Lepraria lanata | 2022-08-02 | CAM-NSF-Lichen Genomics Team | USA: Great Smoky Mountains National Park, Mount LeConte, N-facing slopes above Trillium Gap Trail, 0.4 mi N of jct w/ Rainbow Falls Trail/Boulevard Trail at LeConte Lodge | New York State Museum | 35.65878 N 83.43787 W | NYSM0000194870 |
| LELA_LCN _03a | Lepraria lanata | 2022-08-02 | CAM-NSF-Lichen Genomics Team | USA: Great Smoky Mountains National Park, Mount LeConte, N-facing slopes above Trillium Gap Trail, 0.4 mi N of jct w/ Rainbow Falls Trail/Boulevard Trail at LeConte Lodge | New York State Museum | 35.65878 N 83.43787 W | NYSM0000194871 |

| | | | | | | | |
|---|---|---|---|---|---|---|---|
| LELA_LCN _02c | Lepraria lanata | 2022-08-02 | CAM-NSF-Lichen Genomics Team | USA: Great Smoky Mountains National Park, Mount LeConte, N-facing slopes above Trillium Gap Trail, 0.4 mi N of jct w/ Rainbow Falls Trail/Boulevard Trail at LeConte Lodge | New York State Museum | 35.65878 N 83.43787 W | NYSM0000194872 |
| LELA_LCN _02b | Lepraria lanata | 2022-08-02 | CAM-NSF-Lichen Genomics Team | USA: Great Smoky Mountains National Park, Mount LeConte, N-facing slopes above Trillium Gap Trail, 0.4 mi N of jct w/ Rainbow Falls Trail/Boulevard Trail at LeConte Lodge | New York State Museum | 35.65878 N 83.43787 W | NYSM0000194873 |
| LELA_LCN _02a | Lepraria lanata | 2022-08-02 | CAM-NSF-Lichen Genomics Team | USA: Great Smoky Mountains National Park, Mount LeConte, N-facing slopes above Trillium Gap Trail, 0.4 mi N of jct w/ Rainbow Falls Trail/Boulevard Trail at LeConte Lodge | New York State Museum | 35.65878 N 83.43787 W | NYSM0000194874 |
| LELA_LCN _01c | Lepraria lanata | 2022-08-02 | CAM-NSF-Lichen Genomics Team | USA: Great Smoky Mountains National Park, Mount LeConte, N-facing slopes above Trillium Gap Trail, 0.4 mi N of jct w/ Rainbow Falls Trail/Boulevard Trail at LeConte Lodge | New York State Museum | 35.65878 N 83.43787 W | NYSM0000194875 |
| LELA_LCN _01b | Lepraria lanata | 2022-08-02 | CAM-NSF-Lichen Genomics Team | USA: Great Smoky Mountains National Park, Mount LeConte, N-facing slopes above Trillium Gap Trail, 0.4 mi N of jct w/ Rainbow Falls Trail/Boulevard Trail at LeConte Lodge | New York State Museum | 35.65878 N 83.43787 W | NYSM0000194876 |

| | | | | | | | |
|---|---|---|---|---|---|---|---|
| LELA_LCN _01a | Lepraria lanata | 2022-08-02 | CAM-NSF-Lichen Genomics Team | USA: Great Smoky Mountains National Park, Mount LeConte, N-facing slopes above Trillium Gap Trail, 0.4 mi N of jct w/ Rainbow Falls Trail/Boulevard Trail at LeConte Lodge | New York State Museum | 35.65878 N 83.43787 W | NYSM0000194877 |
| LELA_DAD _03d | Lepraria lanata | 2022-08-10 | CAM-NSF-Lichen Genomics Team | USA: Grandfather Mountain State Park, Grandfather Mountain, N-slopes N of Profile Trail, ~0.3 mi W of jct w/ Grandfather Trail | New York State Museum | 36.11211 N 81.81832 W | NYSM0000194878 |
| LELA_DAD _03c | Lepraria lanata | 2022-08-10 | CAM-NSF-Lichen Genomics Team | USA: Grandfather Mountain State Park, Grandfather Mountain, N-slopes N of Profile Trail, ~0.3 mi W of jct w/ Grandfather Trail | New York State Museum | 36.11211 N 81.81832 W | NYSM0000194879 |
| LELA_DAD _03b | Lepraria lanata | 2022-08-10 | CAM-NSF-Lichen Genomics Team | USA: Grandfather Mountain State Park, Grandfather Mountain, N-slopes N of Profile Trail, ~0.3 mi W of jct w/ Grandfather Trail | New York State Museum | 36.11211 N 81.81832 W | NYSM0000194880 |
| LELA_DAD _03a | Lepraria lanata | 2022-08-10 | CAM-NSF-Lichen Genomics Team | USA: Grandfather Mountain State Park, Grandfather Mountain, N-slopes N of Profile Trail, ~0.3 mi W of jct w/ Grandfather Trail | New York State Museum | 36.11211 N 81.81832 W | NYSM0000194881 |

| | | | | | | | |
|---|---|---|---|---|---|---|---|
| LELA_DAD_02c | Lepraria lanata | 2022-08-10 | CAM-NSF-Lichen Genomics Team | USA: Grandfather Mountain State Park, Grandfather Mountain, N-slopes N of Profile Trail, ~0.3 mi W of jct w/ Grandfather Trail | New York State Museum | 36.11211 N 81.81832 W | NYSM0000194882 |
| LELA_DAD_02b | Lepraria lanata | 2022-08-10 | CAM-NSF-Lichen Genomics Team | USA: Grandfather Mountain State Park, Grandfather Mountain, N-slopes N of Profile Trail, ~0.3 mi W of jct w/ Grandfather Trail | New York State Museum | 36.11211 N 81.81832 W | NYSM0000194883 |
| LELA_DAD_02a | Lepraria lanata | 2022-08-10 | CAM-NSF-Lichen Genomics Team | USA: Grandfather Mountain State Park, Grandfather Mountain, N-slopes N of Profile Trail, ~0.3 mi W of jct w/ Grandfather Trail | New York State Museum | 36.11211 N 81.81832 W | NYSM0000194884 |
| LELA_DAD_01c | Lepraria lanata | 2022-08-10 | CAM-NSF-Lichen Genomics Team | USA: Grandfather Mountain State Park, Grandfather Mountain, N-slopes N of Profile Trail, ~0.3 mi W of jct w/ Grandfather Trail | New York State Museum | 36.11211 N 81.81832 W | NYSM0000194885 |
| LELA_DAD_01b | Lepraria lanata | 2022-08-10 | CAM-NSF-Lichen Genomics Team | USA: Grandfather Mountain State Park, Grandfather Mountain, N-slopes N of Profile Trail, ~0.3 mi W of jct w/ Grandfather Trail | New York State Museum | 36.11211 N 81.81832 W | NYSM0000194886 |
| LELA_DAD_01a | Lepraria lanata | 2022-08-10 | CAM-NSF-Lichen Genomics Team | USA: Grandfather Mountain State Park, Grandfather Mountain, N-slopes N of Profile Trail, ~0.3 mi W of jct w/ Grandfather Trail | New York State Museum | 36.11211 N 81.81832 W | NYSM0000194887 |

| | | | | | | | |
|---|---|---|---|---|---|---|---|
| LELA_CAT _17c | Lepraria lanata | 2021-10-27 | CAM-NSF-Lichen Genomics Team | USA: Pisgah National Forest, summit of Cattail Peak, Black Mountain Crest Trail 3 mi N of Mount Mitchell, ~8 mi S of US19E | New York State Museum | 35.7964 N 82.2569 W | NYSM0000194889 |
| LELA_CAT _17b | Lepraria lanata | 2021-10-27 | CAM-NSF-Lichen Genomics Team | USA: Pisgah National Forest, summit of Cattail Peak, Black Mountain Crest Trail 3 mi N of Mount Mitchell, ~8 mi S of US19E | New York State Museum | 35.7964 N 82.2569 W | NYSM0000194890 |
| LELA_CAT _17a | Lepraria lanata | 2021-10-27 | CAM-NSF-Lichen Genomics Team | USA: Pisgah National Forest, summit of Cattail Peak, Black Mountain Crest Trail 3 mi N of Mount Mitchell, ~8 mi S of US19E | New York State Museum | 35.7964 N 82.2569 W | NYSM0000194891 |
| LELA_CAT _15c | Lepraria lanata | 2021-10-27 | CAM-NSF-Lichen Genomics Team | USA: Pisgah National Forest, summit of Cattail Peak, Black Mountain Crest Trail 3 mi N of Mount Mitchell, ~8 mi S of US19E | New York State Museum | 35.7964 N 82.2569 W | NYSM0000194892 |
| LELA_CAT _15b | Lepraria lanata | 2021-10-27 | CAM-NSF-Lichen Genomics Team | USA: Pisgah National Forest, summit of Cattail Peak, Black Mountain Crest Trail 3 mi N of Mount Mitchell, ~8 mi S of US19E | New York State Museum | 35.7964 N 82.2569 W | NYSM0000194893 |
| LELA_CAT _15a | Lepraria lanata | 2021-10-27 | CAM-NSF-Lichen Genomics Team | USA: Pisgah National Forest, summit of Cattail Peak, Black Mountain Crest Trail 3 mi N of Mount Mitchell, ~8 mi S of US19E | New York State Museum | 35.7964 N 82.2569 W | NYSM0000194888 |

| | | | | | | | |
|---|---|---|---|---|---|---|---|
| LELA_CAT_14e | Lepraria lanata | 2021-10-27 | CAM-NSF-Lichen Genomics Team | USA: Pisgah National Forest, summit of Cattail Peak, Black Mountain Crest Trail 3 mi N of Mount Mitchell, ~8 mi S of US19E | New York State Museum | 35.7964 N 82.2569 W | NYSM0000194894 |
| LELA_CAT_14d | Lepraria lanata | 2021-10-27 | CAM-NSF-Lichen Genomics Team | USA: Pisgah National Forest, summit of Cattail Peak, Black Mountain Crest Trail 3 mi N of Mount Mitchell, ~8 mi S of US19E | New York State Museum | 35.7964 N 82.2569 W | NYSM0000194895 |
| LELA_CAT_14c | Lepraria lanata | 2021-10-27 | CAM-NSF-Lichen Genomics Team | USA: Pisgah National Forest, summit of Cattail Peak, Black Mountain Crest Trail 3 mi N of Mount Mitchell, ~8 mi S of US19E | New York State Museum | 35.7964 N 82.2569 W | NYSM0000194896 |
| LELA_CAT_14b | Lepraria lanata | 2021-10-27 | CAM-NSF-Lichen Genomics Team | USA: Pisgah National Forest, summit of Cattail Peak, Black Mountain Crest Trail 3 mi N of Mount Mitchell, ~8 mi S of US19E | New York State Museum | 35.7964 N 82.2569 W | NYSM0000194897 |
| LELA_CAT_14a | Lepraria lanata | 2021-10-27 | CAM-NSF-Lichen Genomics Team | USA: Pisgah National Forest, summit of Cattail Peak, Black Mountain Crest Trail 3 mi N of Mount Mitchell, ~8 mi S of US19E | New York State Museum | 35.7964 N 82.2569 W | NYSM0000194898 |
| LELA_CAT_13c | Lepraria lanata | 2021-10-27 | CAM-NSF-Lichen Genomics Team | USA: Pisgah National Forest, summit of Cattail Peak, Black Mountain Crest Trail 3 mi N of Mount Mitchell, ~8 mi S of US19E | New York State Museum | 35.7964 N 82.2569 W | NYSM0000194899 |

| | | | | | | | |
|---|---|---|---|---|---|---|---|
| LELA_CAT_13b | Lepraria lanata | 2021-10-27 | CAM-NSF-Lichen Genomics Team | USA: Pisgah National Forest, summit of Cattail Peak, Black Mountain Crest Trail 3 mi N of Mount Mitchell, ~8 mi S of US19E | New York State Museum | 35.7964 N 82.2569 W | NYSM0000194900 |
| LELA_CAT_13a | Lepraria lanata | 2021-10-27 | CAM-NSF-Lichen Genomics Team | USA: Pisgah National Forest, summit of Cattail Peak, Black Mountain Crest Trail 3 mi N of Mount Mitchell, ~8 mi S of US19E | New York State Museum | 35.7964 N 82.2569 W | NYSM0000194901 |
| LELA_ASG_02f | Lepraria lanata | 2022-08-06 | CAM-NSF-Lichen Genomics Team | USA: Cherokee National Forest, Roan Mountain, Appalachian Trail 0.1 mi E of Cloudland Hotel Site, just N of Toll House Gap | New York State Museum | 36.10455 N 82.1293 W | NYSM0000194902 |
| LELA_ASG_02e | Lepraria lanata | 2022-08-06 | CAM-NSF-Lichen Genomics Team | USA: Cherokee National Forest, Roan Mountain, Appalachian Trail 0.1 mi E of Cloudland Hotel Site, just N of Toll House Gap | New York State Museum | 36.10455 N 82.1293 W | NYSM0000191883 |
| LELA_ASG_02d | Lepraria lanata | 2022-08-06 | CAM-NSF-Lichen Genomics Team | USA: Cherokee National Forest, Roan Mountain, Appalachian Trail 0.1 mi E of Cloudland Hotel Site, just N of Toll House Gap | New York State Museum | 36.10455 N 82.1293 W | NYSM0000191882 |
| LELA_ASG_02c | Lepraria lanata | 2022-08-06 | CAM-NSF-Lichen Genomics Team | USA: Cherokee National Forest, Roan Mountain, Appalachian Trail 0.1 mi E of Cloudland Hotel Site, just N of Toll House Gap | New York State Museum | 36.10455 N 82.1293 W | NYSM0000191881 |

| | | | | | | | |
|---|---|---|---|---|---|---|---|
| LELA_ASG _02b | Lepraria lanata | 2022-08-06 | CAM-NSF-Lichen Genomics Team | USA: Cherokee National Forest, Roan Mountain, Appalachian Trail 0.1 mi E of Cloudland Hotel Site, just N of Toll House Gap | New York State Museum | 36.10455 N 82.1293 W | NYSM0000191880 |
| LELA_ASG _02a | Lepraria lanata | 2022-08-06 | CAM-NSF-Lichen Genomics Team | USA: Cherokee National Forest, Roan Mountain, Appalachian Trail 0.1 mi E of Cloudland Hotel Site, just N of Toll House Gap | New York State Museum | 36.10455 N 82.1293 W | NYSM0000191879 |
| LELA_ASG _01e | Lepraria lanata | 2022-08-06 | CAM-NSF-Lichen Genomics Team | USA: Cherokee National Forest, Roan Mountain, Appalachian Trail 0.1 mi E of Cloudland Hotel Site, just N of Toll House Gap | New York State Museum | 36.10455 N 82.1293 W | NYSM0000194903 |
| LELA_ASG _01d | Lepraria lanata | 2022-08-06 | CAM-NSF-Lichen Genomics Team | USA: Cherokee National Forest, Roan Mountain, Appalachian Trail 0.1 mi E of Cloudland Hotel Site, just N of Toll House Gap | New York State Museum | 36.10455 N 82.1293 W | NYSM0000191878 |
| LELA_ASG _01c | Lepraria lanata | 2022-08-06 | CAM-NSF-Lichen Genomics Team | USA: Cherokee National Forest, Roan Mountain, Appalachian Trail 0.1 mi E of Cloudland Hotel Site, just N of Toll House Gap | New York State Museum | 36.10455 N 82.1293 W | NYSM0000191877 |
| LELA_ASG _01b | Lepraria lanata | 2022-08-06 | CAM-NSF-Lichen Genomics Team | USA: Cherokee National Forest, Roan Mountain, Appalachian Trail 0.1 mi E of Cloudland Hotel Site, just N of Toll House Gap | New York State Museum | 36.10455 N 82.1293 W | NYSM0000191876 |

| | | | | | | | |
|---|---|---|---|---|---|---|---|
| LELA_ASG _01a | Lepraria lanata | 2022-08-06 | CAM-NSF-Lichen Genomics Team | USA: Cherokee National Forest, Roan Mountain, Appalachian Trail 0.1 mi E of Cloudland Hotel Site, just N of Toll House Gap | New York State Museum | 36.10455 N 82.1293 W | NYSM0000191875 |
| LELA_ALM _07c | Lepraria lanata | 2022-08-02 | CAM-NSF-Lichen Genomics Team | USA: Great Smoky Mountains National Park, Mount LeConte, N-facing slopes above Trillium Gap Trail, 0.4 mi N of jct w/ Rainbow Falls Trail/Boulevard Trail at LeConte Lodge | New York State Museum | 35.65878 N 83.43787 W | NYSM0000191904 |
| LELA_ALM _07b | Lepraria lanata | 2022-08-02 | CAM-NSF-Lichen Genomics Team | USA: Great Smoky Mountains National Park, Mount LeConte, N-facing slopes above Trillium Gap Trail, 0.4 mi N of jct w/ Rainbow Falls Trail/Boulevard Trail at LeConte Lodge | New York State Museum | 35.65878 N 83.43787 W | NYSM0000191903 |
| LELA_ALM _07a | Lepraria lanata | 2022-08-02 | CAM-NSF-Lichen Genomics Team | USA: Great Smoky Mountains National Park, Mount LeConte, N-facing slopes above Trillium Gap Trail, 0.4 mi N of jct w/ Rainbow Falls Trail/Boulevard Trail at LeConte Lodge | New York State Museum | 35.65878 N 83.43787 W | NYSM0000191902 |
| LELA_ALM _06c | Lepraria lanata | 2022-08-02 | CAM-NSF-Lichen Genomics Team | USA: Great Smoky Mountains National Park, Mount LeConte, N-facing slopes above Trillium Gap Trail, 0.4 mi N of jct w/ Rainbow Falls Trail/Boulevard Trail at LeConte Lodge | New York State Museum | 35.65878 N 83.43787 W | NYSM0000191901 |

| | | | | | | | |
|---|---|---|---|---|---|---|---|
| LELA_ALM_06b | Lepraria lanata | 2022-08-02 | CAM-NSF-Lichen Genomics Team | USA: Great Smoky Mountains National Park, Mount LeConte, N-facing slopes above Trillium Gap Trail, 0.4 mi N of jct w/ Rainbow Falls Trail/Boulevard Trail at LeConte Lodge | New York State Museum | 35.65878 N 83.43787 W | NYSM0000191900 |
| LELA_ALM_06a | Lepraria lanata | 2022-08-02 | CAM-NSF-Lichen Genomics Team | USA: Great Smoky Mountains National Park, Mount LeConte, N-facing slopes above Trillium Gap Trail, 0.4 mi N of jct w/ Rainbow Falls Trail/Boulevard Trail at LeConte Lodge | New York State Museum | 35.65878 N 83.43787 W | NYSM0000191899 |
| LELA_ALM_05c | Lepraria lanata | 2022-08-02 | CAM-NSF-Lichen Genomics Team | USA: Great Smoky Mountains National Park, Mount LeConte, N-facing slopes above Trillium Gap Trail, 0.4 mi N of jct w/ Rainbow Falls Trail/Boulevard Trail at LeConte Lodge | New York State Museum | 35.65878 N 83.43787 W | NYSM0000191898 |
| LELA_ALM_05b | Lepraria lanata | 2022-08-02 | CAM-NSF-Lichen Genomics Team | USA: Great Smoky Mountains National Park, Mount LeConte, N-facing slopes above Trillium Gap Trail, 0.4 mi N of jct w/ Rainbow Falls Trail/Boulevard Trail at LeConte Lodge | New York State Museum | 35.65878 N 83.43787 W | NYSM0000191897 |
| LELA_ALM_05a | Lepraria lanata | 2022-08-02 | CAM-NSF-Lichen Genomics Team | USA: Great Smoky Mountains National Park, Mount LeConte, N-facing slopes above Trillium Gap Trail, 0.4 mi N of jct w/ Rainbow Falls Trail/Boulevard Trail at LeConte Lodge | New York State Museum | 35.65878 N 83.43787 W | NYSM0000191896 |

| | | | | | | | |
|---|---|---|---|---|---|---|---|
| LELA_ALM _04c | Lepraria lanata | 2022-08-02 | CAM-NSF-Lichen Genomics Team | USA: Great Smoky Mountains National Park, Mount LeConte, N-facing slopes above Trillium Gap Trail, 0.4 mi N of jct w/ Rainbow Falls Trail/Boulevard Trail at LeConte Lodge | New York State Museum | 35.65878 N 83.43787 W | NYSM0000191895 |
| LELA_ALM _04b | Lepraria lanata | 2022-08-02 | CAM-NSF-Lichen Genomics Team | USA: Great Smoky Mountains National Park, Mount LeConte, N-facing slopes above Trillium Gap Trail, 0.4 mi N of jct w/ Rainbow Falls Trail/Boulevard Trail at LeConte Lodge | New York State Museum | 35.65878 N 83.43787 W | NYSM0000191894 |
| LELA_ALM _04a | Lepraria lanata | 2022-08-02 | CAM-NSF-Lichen Genomics Team | USA: Great Smoky Mountains National Park, Mount LeConte, N-facing slopes above Trillium Gap Trail, 0.4 mi N of jct w/ Rainbow Falls Trail/Boulevard Trail at LeConte Lodge | New York State Museum | 35.65878 N 83.43787 W | NYSM0000191893 |
| LELA_ALM _03c | Lepraria lanata | 2022-08-02 | CAM-NSF-Lichen Genomics Team | USA: Great Smoky Mountains National Park, Mount LeConte, N-facing slopes above Trillium Gap Trail, 0.4 mi N of jct w/ Rainbow Falls Trail/Boulevard Trail at LeConte Lodge | New York State Museum | 35.65878 N 83.43787 W | NYSM0000191892 |
| LELA_ALM _03b | Lepraria lanata | 2022-08-02 | CAM-NSF-Lichen Genomics Team | USA: Great Smoky Mountains National Park, Mount LeConte, N-facing slopes above Trillium Gap Trail, 0.4 mi N of jct w/ Rainbow Falls Trail/Boulevard Trail at LeConte Lodge | New York State Museum | 35.65878 N 83.43787 W | NYSM0000191891 |

| | | | | | | | |
|---|---|---|---|---|---|---|---|
| LELA_ALM_03a | Lepraria lanata | 2022-08-02 | CAM-NSF-Lichen Genomics Team | USA: Great Smoky Mountains National Park, Mount LeConte, N-facing slopes above Trillium Gap Trail, 0.4 mi N of jct w/ Rainbow Falls Trail/Boulevard Trail at LeConte Lodge | New York State Museum | 35.65878 N 83.43787 W | NYSM0000191890 |
| LELA_ALM_02c | Lepraria lanata | 2022-08-02 | CAM-NSF-Lichen Genomics Team | USA: Great Smoky Mountains National Park, Mount LeConte, N-facing slopes above Trillium Gap Trail, 0.4 mi N of jct w/ Rainbow Falls Trail/Boulevard Trail at LeConte Lodge | New York State Museum | 35.65878 N 83.43787 W | NYSM0000191889 |
| LELA_ALM_02b | Lepraria lanata | 2022-08-02 | CAM-NSF-Lichen Genomics Team | USA: Great Smoky Mountains National Park, Mount LeConte, N-facing slopes above Trillium Gap Trail, 0.4 mi N of jct w/ Rainbow Falls Trail/Boulevard Trail at LeConte Lodge | New York State Museum | 35.65878 N 83.43787 W | NYSM0000191888 |
| LELA_ALM_02a | Lepraria lanata | 2022-08-02 | CAM-NSF-Lichen Genomics Team | USA: Great Smoky Mountains National Park, Mount LeConte, N-facing slopes above Trillium Gap Trail, 0.4 mi N of jct w/ Rainbow Falls Trail/Boulevard Trail at LeConte Lodge | New York State Museum | 35.65878 N 83.43787 W | NYSM0000191887 |
| LELA_ALM_01c | Lepraria lanata | 2022-08-02 | CAM-NSF-Lichen Genomics Team | USA: Great Smoky Mountains National Park, Mount LeConte, N-facing slopes above Trillium Gap Trail, 0.4 mi N of jct w/ Rainbow Falls Trail/Boulevard Trail at LeConte Lodge | New York State Museum | 35.65878 N 83.43787 W | NYSM0000191886 |

| | | | | | | | |
|---|---|---|---|---|---|---|---|
| LELA_ALM_01b | Lepraria lanata | 2022-08-02 | CAM-NSF-Lichen Genomics Team | USA: Great Smoky Mountains National Park, Mount LeConte, N-facing slopes above Trillium Gap Trail, 0.4 mi N of jct w/ Rainbow Falls Trail/Boulevard Trail at LeConte Lodge | New York State Museum | 35.65878 N 83.43787 W | NYSM0000191885 |
| LELA_ALM_01a | Lepraria lanata | 2022-08-02 | CAM-NSF-Lichen Genomics Team | USA: Great Smoky Mountains National Park, Mount LeConte, N-facing slopes above Trillium Gap Trail, 0.4 mi N of jct w/ Rainbow Falls Trail/Boulevard Trail at LeConte Lodge | New York State Museum | 35.65878 N 83.43787 W | NYSM0000191884 |

**Table S13.**
Voucher specimen information.

| Species | Subset | Individuals (pre-filter) | Variants (pre-filter) | Individuals (post-filter) | Variants (post-filter) | | |
|---|---|---|---|---|---|---|---|
| *Lepraria finkii* | Mycobiont - Noncoding | 238 | 3,364,960 | 227 | 17,416 | | |
| *Lepraria lanata* | Mycobiont - Noncoding | 114 | 2,649,775 | 81 | 12,692 | | |
| *Punctelia appalachensis* | Mycobiont - Noncoding | 187 | 3,175,490 | 181 | 18,495 | **Max. Variant Count (pre-filter):** | 3,770,852 |
| *Punctelia rudecta* | Mycobiont - Noncoding | 297 | 3,006,924 | 291 | 18,612 | **Min. Variant Count (pre-filter):** | 2,649,775 |
| *Usnea strigosa* | Mycobiont - Noncoding | 45 | 3,763,258 | 45 | 12,250 | **Max. Variant Count (post-filter):** | 18,612 |
| *Usnea subfusca* | Mycobiont - Noncoding | 81 | 3,770,852 | 76 | 14,151 | **Min. Variant Count (post-filter):** | 12,250 |
| | **Total Individuals (pre-filter):** | **962** | **Total Individuals (post-filter):** | **901** | | | |
| *Lepraria finkii* | Photobiont - Pop Gen. | 238 | 14,948 | 237 | 3,085 | | |
| *Lepraria lanata* | Photobiont - Pop Gen. | 114 | 1,188 | 81 | 1,104 | | |
| *Punctelia appalachensis* | Photobiont - Pop Gen. | 187 | 15,671 | 184 | 5,234 | **Max. Variant Count (pre-filter):** | 22,971 |
| *Punctelia rudecta* | Photobiont - Pop Gen. | 297 | 17,068 | 289 | 5,224 | **Min. Variant Count (pre-filter):** | 1,188 |
| *Usnea strigosa* | Photobiont - | 45 | 17,421 | 45 | 4,464 | **Max. Variant** | 5,904 |

| | | | | | | | |
|---|---|---|---|---|---|---|---|
| | Pop Gen. | | | | | **Count (post-filter):** | |
| *Usnea subfusca* | Photobiont - Pop Gen. | 81 | 22,971 | 78 | 5,904 | **Min. Variant Count (post-filter):** | 1,104 |
| | **Total Individuals (pre-filter):** | **962** | **Total Individuals (post-filter):** | **914** | | | |
| *Lepraria finkii* | Mycobont - Coding | 238 | 1,175,991 | 202 | 22,668 | | |
| *Lepraria lanata* | Mycobont - Coding | 114 | 755,604 | 60 | 4,322 | | |
| *Punctelia appalachensis* | Mycobont - Coding | 187 | 1,204,953 | 126 | 17,525 | **Max. Variant Count (pre-filter):** | 2,330,686 |
| *Punctelia rudecta* | Mycobont - Coding | 297 | 1,610,697 | 219 | 20,001 | **Min. Variant Count (pre-filter):** | 755,604 |
| *Usnea strigosa* | Mycobont - Coding | 45 | 2,330,686 | 44 | 26,396 | **Max. Variant Count (post-filter):** | 26,396 |
| *Usnea subfusca* | Mycobont - Coding | 81 | 1,441,249 | 74 | 24,850 | **Min. Variant Count (post-filter):** | 4,322 |
| | **Total Individuals (pre-filter):** | **962** | **Total Individuals (post-filter):** | **725** | | | |
| *Lepraria finkii* | Photobiont - LFMM | 238 | 14,948 | 237 | 257 | | |
| *Lepraria lanata* | Photobiont - LFMM | 114 | 1,188 | 114 | 710 | | |

| | | | | | | | |
|---|---|---|---|---|---|---|---|
| *Punctelia appalachensis* | Photobiont - LFMM | 187 | 15,671 | 145 | 800 | **Max. Variant Count (pre-filter):** | 22,971 |
| *Punctelia rudecta* | Photobiont - LFMM | 297 | 17,068 | 229 | 874 | **Min. Variant Count (pre-filter):** | 1,188 |
| *Usnea strigosa* | Photobiont - LFMM | 45 | 17,421 | 45 | 1,797 | **Max. Variant Count (post-filter):** | 1797 |
| *Usnea subfusca* | Photobiont - LFMM | 81 | 22,971 | 51 | 1,248 | **Min. Variant Count (post-filter):** | 257 |
| | **Total Individuals (pre-filter):** | **962** | **Total Individuals (post-filter):** | **821** | | | |

**Table S14.**
Variant counts for all datasets before and after filtering.

| Species | Subset | Site | Mean Allelic Richness (Ar) | Population-specific $F_{ST}$ |
|---|---|---|---|---|
| *Lepraria finkii* | Mycobiont | BEK | 1.201408 | 0.1843865 |
| | | BFF | 1.220111 | 0.1095744 |
| | | BLU | 1.188992 | 0.2349695 |
| | | BRF | 1.214647 | 0.131644 |
| | | BUK | 1.208311 | 0.1572266 |
| | | CAT | 1.16473 | 0.3325582 |
| | | COG | 1.160962 | 0.345468 |
| | | GOS | 1.189165 | 0.2335191 |
| | | GRI | 1.220495 | 0.107588 |
| | | HAW | 1.210549 | 0.1480284 |
| | | JON | 1.198102 | 0.1982925 |
| | | KLE | 1.197696 | 0.1996521 |
| | | LEM | 1.190582 | 0.228399 |
| | | LUE | 1.207514 | 0.1597273 |
| | | NGA | 1.206581 | 0.1643264 |
| | | POK | 1.179587 | 0.2731833 |
| | | ROG | 1.178124 | 0.2785621 |
| | | RON | 1.207568 | 0.1601501 |
| | | RSK | 1.206873 | 0.1630422 |
| | | SHR | 1.183971 | 0.2550503 |
| | | SNR | 1.211581 | 0.1440487 |
| | | TMK | 1.188222 | 0.2379072 |
| | | TOM | 1.19519 | 0.2099396 |
| | | TRI | 1.193413 | 0.2170274 |
| | | TYS | 1.19887 | 0.1952826 |
| | | WHI | 1.173819 | 0.296424 |
| | | **Mean** | 1.196040885 | 0.20638375 |
| *Lepraria lanata* | Noncoding | ALM | 1 | 0.09303902 |

| | | | | |
|---|---|---|---|---|
| | | CAT | 1 | 0.4509759 |
| | | LCN | 1 | 0.03377004 |
| | | NAL | 1 | 0.10580479 |
| | | TOM | 1 | 0.41481628 |
| | | **Mean** | 1 | 0.219681206 |
| *Punctelia appalachensis* | Mycobiont | BEK | 1.092693 | NaN |
| | | BFF | 1.096207 | 0.16510303 |
| | | BLU | 1.094743 | 0.17760852 |
| | | BRF | 1.090385 | 0.21262306 |
| | | BUK | 1.09426 | 0.18286965 |
| | | DAD | 1.087925 | 0.22407365 |
| | | DBG | 1.101154 | 0.12815129 |
| | | GOS | 1.106051 | 0.08005878 |
| | | HAW | 1.069981 | 0.38280261 |
| | | KLE | 1.088749 | 0.22801667 |
| | | LEM | 1.060367 | 0.45579395 |
| | | LUE | 1.071074 | 0.37263537 |
| | | NGA | 1.104621 | 0.09353399 |
| | | RON | 1.072722 | 0.35964 |
| | | RSK | 1.093554 | 0.18569973 |
| | | SNR | 1.096078 | 0.1681604 |
| | | TRI | 1.056279 | 0.5050352 |
| | | TYS | 1.069496 | NaN |
| | | WHI | 1.095716 | 0.17240064 |
| | | **Mean** | 1.086423947 | 0.2408356788 |
| *Punctelia rudecta* | Mycobiont | BEK | 1.0612 | 0.183313202 |
| | | BFF | 1.067003 | 0.106154223 |
| | | BLU | 1.06837 | 0.087923339 |
| | | BRF | 1.070854 | 0.054784717 |

| | | | | |
|---|---|---|---|---|
| | | BUK | 1.068904 | 0.080801106 |
| | | CAT | 1.055377 | 0.261025708 |
| | | DBG | 1.067786 | 0.095708701 |
| | | GOS | 1.065758 | 0.122773057 |
| | | GRI | 1.050699 | 0.321787658 |
| | | HAW | 1.060332 | 0.195145754 |
| | | JON | 1.062395 | 0.167376868 |
| | | KLE | 1.070153 | 0.064131399 |
| | | LEM | 1.047739 | 0.362362018 |
| | | LUE | 1.068915 | 0.080655084 |
| | | NAR | 1.059247 | 0.209385196 |
| | | NGA | 1.068548 | 0.08526168 |
| | | POK | 1.047674 | 0.363618378 |
| | | ROG | 1.056875 | 0.241268097 |
| | | RON | 1.069659 | 0.070726126 |
| | | RSK | 1.074459 | 0.006688795 |
| | | SHR | 1.061894 | 0.174310541 |
| | | SNR | 1.062686 | 0.163488048 |
| | | TMK | 1.064345 | 0.141619069 |
| | | TOM | 1.054367 | 0.274307615 |
| | | TRI | 1.069516 | 0.072639195 |
| | | TYS | 1.028091 | 0.621529624 |
| | | WHI | 1.072529 | 0.032438513 |
| | | **Mean** | 1.062050926 | 0.1718971745 |
| *Usnea strigosa* | Mycobiont | GOS | 1.310823 | 0.1796578 |
| | | JON | 1.291814 | 0.2290684 |
| | | MAX | 1.308074 | 0.1900963 |
| | | RON | 1.230375 | 0.3830101 |
| | | SNR | 1.259088 | 0.3093637 |
| | | WHI | 1.292188 | 0.2319005 |

| | | | | |
|---|---|---|---|---|
| | | **Mean** | 1.282060333 | 0.2538494667 |
| *Usnea subfusca* | Mycobiont | BEK | 1.19768 | 0.08860078 |
| | | BFF | 1.198523 | 0.08129445 |
| | | BUK | 1.199339 | 0.08051829 |
| | | NAR | 1.205831 | 0.05534028 |
| | | NGA | 1.186957 | 0.12617281 |
| | | SHR | 1.209062 | 0.0370096 |
| | | TRI | 1.202397 | 0.05010078 |
| | | **Mean** | 1.199969857 | 0.07414814143 |
| *Lepraria finkii* | Photobiont | BEK | 1.383938 | -0.12852643 |
| | | BFF | 1.356525 | -0.04795236 |
| | | BLU | 1.353357 | -0.03863962 |
| | | BRF | 1.352636 | -0.03651903 |
| | | BUK | 1.360062 | -0.05834908 |
| | | CAT | 1.35588 | -0.04605524 |
| | | COG | 1.373937 | -0.0991322 |
| | | GOS | 1.357564 | -0.05100385 |
| | | GRI | 1.346747 | -0.01920915 |
| | | HAW | 1.351027 | -0.03179147 |
| | | JON | 1.352171 | -0.03515454 |
| | | KLE | 1.360812 | -0.06055187 |
| | | LEM | 1.357955 | -0.05215315 |
| | | LUE | 1.358728 | -0.05442707 |
| | | NGA | 1.352004 | -0.03466166 |
| | | POK | 1.354522 | -0.04206247 |
| | | ROG | 1.366531 | -0.07736321 |
| | | RON | 1.348346 | -0.02390997 |
| | | RSK | 1.352868 | -0.03720085 |
| | | SHR | 1.375836 | -0.10485584 |

| | | | | |
|---|---|---|---|---|
| | | SNR | 1.355865 | -0.04601233 |
| | | TMK | 1.376121 | -0.1055499 |
| | | TOM | 1.359945 | -0.05800436 |
| | | TRI | 1.362643 | -0.06593475 |
| | | TYS | 1.35178 | -0.03400258 |
| | | WHI | 1.357767 | -0.05160308 |
| | | **Mean** | 1.359060269 | -0.05540869462 |
| *Lepraria lanata* | Photobiont | ALM | 1.228384 | 0.44613257 |
| | | ASG | 1.199578 | 0.42753993 |
| | | CAT | 1.299585 | 0.26665334 |
| | | DAD | 1.327446 | -0.07688372 |
| | | LCN | 1.045237 | 0.85460742 |
| | | NAL | 1.045884 | 0.84900182 |
| | | RBB | 1.540376 | -0.78454527 |
| | | SPF | 1.135878 | 0.49794495 |
| | | TOM | 1.462116 | -0.29009062 |
| | | **Mean** | 1.253831556 | 0.24337338 |
| *Punctelia appalachensis* | Photobiont | BEK | 1.040489 | 0.44417551 |
| | | BFF | 1.053208 | 0.26956774 |
| | | BLU | 1.047767 | 0.34425968 |
| | | BRF | 1.062516 | 0.14178571 |
| | | BUK | 1.037249 | 0.48864743 |
| | | DAD | 1.074932 | -0.02864854 |
| | | DBG | 1.036787 | 0.49498846 |
| | | GOS | 1.036172 | 0.5034302 |
| | | HAW | 1.080836 | -0.10970627 |
| | | KLE | 1.109238 | -0.49959759 |
| | | LEM | 1.120086 | -0.64852251 |
| | | LUE | 1.020631 | 0.71678289 |

| | | | | |
|---|---|---|---|---|
| | | NGA | 1.084828 | -0.16451016 |
| | | RON | 1.098241 | -0.34863845 |
| | | RSK | 1.090206 | -0.23833553 |
| | | SNR | 1.031683 | 0.56505786 |
| | | TRI | 1.120325 | -0.65180085 |
| | | TYS | 1.105144 | -0.44339279 |
| | | WHI | 1.035389 | 0.5141851 |
| | | **Mean** | 1.067669842 | 0.07103831 |
| *Punctelia rudecta* | Photobiont | BEK | 1.193861 | -0.132034084 |
| | | BFF | 1.041882 | 0.766916267 |
| | | BLU | 1.163062 | 0.132168423 |
| | | BRF | 1.220902 | -0.417959814 |
| | | BUK | 1.074832 | 0.630703135 |
| | | CAT | 1.165402 | 0.088525995 |
| | | DBG | 1.039947 | 0.77997439 |
| | | GOS | 1.148021 | 0.240617338 |
| | | GRI | 1.223011 | -0.314334768 |
| | | HAW | 1.223236 | -0.353525979 |
| | | JON | 1.207528 | -0.340146539 |
| | | KLE | 1.202883 | -0.367406888 |
| | | LEM | 1.206986 | -0.183559694 |
| | | LUE | 1.168297 | 0.143301879 |
| | | NAR | 1.038396 | 0.785265735 |
| | | NGA | 1.160789 | 0.130899759 |
| | | POK | 1.178828 | 0.003806816 |
| | | ROG | 1.237971 | -0.380211145 |
| | | RON | 1.178531 | 0.0415925 |
| | | RSK | 1.20127 | -0.197960066 |
| | | SHR | 1.02263 | 0.849734866 |
| | | SNR | 1.174562 | 0.082884526 |

| | | | | |
|---|---|---|---|---|
| | | TMK | 1.186432 | -0.005025765 |
| | | TOM | 1.179474 | -0.062929329 |
| | | TRI | 1.227596 | -0.363725761 |
| | | TYS | 1.187406 | -0.053454273 |
| | | WHI | 1.078296 | 0.6030682 |
| | | **Mean** | 1.160445593 | 0.0780439157 |
| *Usnea strigosa* | Photobiont | GOS | 2.210147 | 0.049458816 |
| | | JON | 2.542267 | -0.051934394 |
| | | MAX | 2.352976 | 0.002662306 |
| | | RON | 2.028898 | -0.021417204 |
| | | SNR | 2.478839 | -0.082236597 |
| | | WHI | 2.032847 | 0.097130962 |
| | | **Mean** | 2.274329 | -0.0010560185 |
| *Usnea subfusca* | Photobiont | BEK | 1.296276 | 0.023333767 |
| | | BFF | 1.298904 | 0.120957666 |
| | | BUK | 1.351395 | -0.104804338 |
| | | NAR | 1.328529 | 0.033481693 |
| | | NGA | 1.282666 | 0.045921641 |
| | | SHR | 1.350607 | -0.065344672 |
| | | TRI | 1.323518 | -0.006388669 |
| | | **Mean** | 1.318842143 | 0.006736726857 |

**Table S15.**
Allelic richness and population-specific $F_{ST}$ for all mycobiont and photobiont site.
